# *Ba*Cas12a3 represents a new subtype of type V CRISPR effector with collateral activity toward tRNA

**DOI:** 10.64898/2026.01.16.699850

**Authors:** Shiqi Ji, Xin Li, Chenwei Wu, Jinshan Guo, Ruyi Zheng, Jingwen Kong, Lei Du, Qunxin She

**Author notes:** **Authors for correspondence:** Shiqi Ji, Qunxin She. Both authors equally contribute to this work (Shiqi Ji, Xin Li).

## Abstract

The CRISPR-Cas12 family encompasses diverse RNA-guided nucleases with both DNA-targeting and RNA-targeting subtypes. They can trigger antiviral activities mainly through either direct elimination invading nucleic acids, or activating broad collateral cleavage to induce abortive infection. Here, we report a novel type V CRISPR effector *Ba*Cas12a3 that causes growth inhibition through a unique tRNA-cleavage mechanism. Plasmid interference assays indicated that *Ba*Cas12a3 inhibits host growth arrest without invoking the DNA damage response, suggesting that the immune responses may not involve double strand breaks of DNA. Indeed, biochemical characterization of the *Ba*Cas12a3-crRNA ribonucleoprotein (RNP) unraveled that the effector is an RNA-activating nuclease that cleaves 3′ terminus of tRNAs. Cryo-EM structures of *Ba*Cas12a3 reveal a conserved bilobed architecture featuring a unique nucleic acid-loading (NL) domain adjacent to the RuvC catalytic center. Structural and mutagenesis analyses show that the NL domain, together with a zinc ribbon domain, form a gated substrate groove. Target RNA binding induces conformational changes that open this groove and expose the RuvC active site, enabling specific tRNA cleavage while preventing other non-specific degradation. Our findings identified the NL domain aside the RuvC active site responsible for the tRNA recognition in *Ba*Cas12a3, expanding the functional diversity of CRISPR immunity.

## Introduction

The CRISPR-Cas (clustered regularly interspaced short palindromic repeats and CRISPR-associated proteins) system constitutes the prokaryotic adaptive immunity, enabling bacteria and archaea to neutralize viral invaders through RNA-guided nucleic acid cleavage (1, 2). These antiviral systems are broadly classified into Class 1 systems with multi-subunit protein effectors and Class 2 systems utilizing a single-unit protein effector (2–4). Three Class 2 types are known among which Type V CRISPR-Cas employing nucleases of the Cas12 family as the effector have emerged as versatile tools for genome engineering and diagnostics due to their programmable DNA/RNA-targeting activities and collateral nuclease behaviors (5–10). Evolutionarily related to TnpB transposases, Cas12 proteins share a conserved C-terminal RuvC endonuclease domain (5, 9, 11). To date, more than 16 Type V subtypes (Cas12a–Cas12o) have been characterized, most displaying dual cleavage activities: *cis*-cleavage of target nucleic acids and *trans*-degradation (collateral cleavage) of non-target nucleic acids (7, 8, 12–16).

Most Type V effectors are classified as DNA-targeting nucleases (7, 8). They target dsDNA as well as ss DNA, and induce target-specific cleavage and nucleic acid collateral cleavage (17–19). Nevertheless, the characterization of Cas12a2 revealed unexpected functional plasticity within the Cas12 family. Cas12a2 effector is an RNA-guided effector capable of inducing abortive infection—a defense mechanism in which cells undergo dormancy or death to limit viral spread (20–22). Upon recognizing RNA target, Cas12a2 unleashes promiscuous cleavage of ssRNA, ssDNA, and dsDNA, a mechanism distinct from the ssRNA-specific collateral activity of Type VI Cas13 enzymes (20, 21, 23).

Structural comparisons across Cas12 subtypes reveal diversification in N-terminal domains, which govern guide RNA and target nucleic acid recognition, and in C-terminal domains, which count for functional divergence. In dsDNA targeting subtypes (*e.g.*, Cas12a, b, e, f, j, m), various accessory domains (*e.g.*, TNB/NUC, BH, Zinc-finger, TSL) facilitate dsDNA unwinding and substrate loading into the RuvC catalytic sites (18, 24–28). By contrast, RNA-targeting variants such as Cas12a2 and Cas12g retain a zinc ribbon (ZR) domain crucial for nucleic acid accommodation (21, 29). These structural variations underscore the evolutionary adaptability of the RuvC domain associated domains and suggest further mechanistic diversity within the Cas12 family remains unexplored.

We further investigated the diversity of Cas12 systems by searching for Cas12 nucleases in different metagenomic datasets, and a new clade, Cas12a3 was revealed, including several such sequences encoded by gut bacteria. One representative, *Ba*Cas12a3, was characterized. We demonstrate that this Cas12 subtype possesses unique structural and functional features, including a novel nucleic acid-loading (NL) domain. By integrating cryo-electron microscopy (cryo-EM), phylogenetic analysis, and biochemical assays, we provide insights into the evolutionary divergence, substrate specificity, and antiviral mechanisms of *Ba*Cas12a3. These findings expand our understanding of the antiviral repertoire of CRISPR systems.

## Materials and methods

### Identification and phylogenetic analysis of Cas12a3 nucleases

Cas12a3 sequences were identified through sequence-based searches using Cas12a (AcCpf1) as a query against the metagenome databases (24, 30, 31). Initial hits served as seeds for subsequent BLAST searches against the NCBI non-redundant database to retrieve entries encoding full-length proteins. Among these, The *Ba*Cas12a3 CRISPR locus was one of these annotated transposases identified from a gut metagenome assembled genome of a *Bacteroidales* bacterium isolate RGIG7249 (GenBank: JAFYNK010000087).

To localize the RuvC domain and its conserved residues, *Ba*Cas12a3 orthologs (Supplementary File 1) were aligned using Clustal Omega (32, 33), and the *Ba*Cas12a3 structure was predicted with AlphaFold3 (34). A multiple sequence alignment of these orthologs and related transposases (Supplementary File 1) was generated using Clustal Omega, followed by phylogenetic analysis performed with the Neighbor-Joining method in MEGA7 and TreeViewer (v2.2.0) (35–37), and the resulting tree in Figure 1 was visualized and annotated using and tvBOT (38).

**Figure 1.**
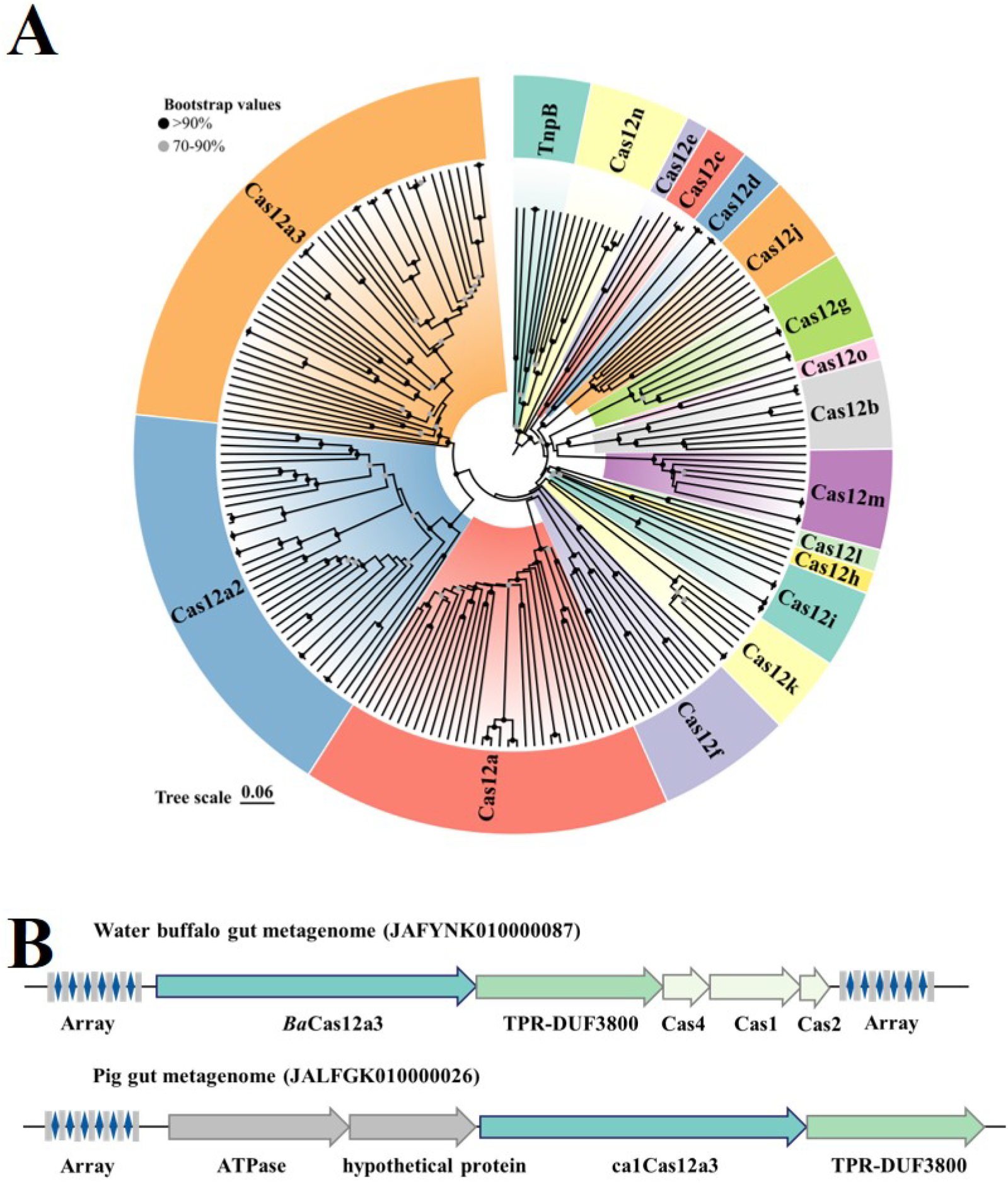
Cas12a3 is a newly identified subtype within the type V CRISPR-Cas effectors. (A), Phylogenetic tree of Cas12 subtypes and their ancestral TnpB nucleases. Distinct subtypes are color-coded. Sequences used for tree construction are provided in Supplementary File 1. (B), Gene arrangements of two representative CRISPR-Cas12a3 systems.

### Plasmid construction

The gene encoding *Ba*Cas12a3 was codon-optimized for *Escherichia coli* expression and synthesized by Tsingke Biotechnology Co., Ltd. (Beijing, China). This optimized sequence was cloned into pCDFDuet-1 vector downstream of the first T7 promoter to generate the expression plasmid pCDFDuet-*Ba*Cas12a3. To enable co-expression of the nuclease and its cognate crRNA, a minimal CRISPR array containing five identical spacer repeats was synthesized and inserted downstream of the second T7 promoter in the pCDFDuet-*Ba*Cas12a3 backbone, yielding the plasmid pCas12a3. In both constructs, the nuclease and the crRNA are expressed from independent T7 promoters. For the plasmid interference assays, reporter vectors were constructed in two steps. First, a gene encoding green fluorescent protein (GFP) was inserted into the pBAD plasmid, yielding pBAD-G. Subsequently, DNA fragments containing either a protospacer flanking site (PFS) with its complementary cRNA sequence or the PFS alone (as a non-targeting control) were cloned between the *ara*BAD promoter and the GFP gene in pBAD-G, yielding plasmids pTarget and pNTarget. Following the same approach, the plasmids pTNoPfs (containing the target but lacking a PFS) and pTNoPr (containing the target but lacking its promoter) were also constructed. All plasmids used in the study are listed in Supplementary Tables S1, and all primers are are provided in Supplementary Table S2.

### Plasmid interference assay

The plasmid interference assay was performed as described previously (20). To evaluate the function of the *Ba*Cas12a3 effector complex, we conducted two parallel assays: the nuclease-target selection assay (tns) and a modified version termed the nuclease selection assay (ns). Electrocompetent cells were prepared as follows. For strains expressing the nuclease, the plasmid pCas12a3 (encoding *Ba*Cas12a3 and crRNA) was chemically transformed into *E. coli* BL21(DE3). The strain was cultured in LB medium containing 34 *µ*g/mL streptomycin to OD₆₀₀ about 0.25, induced with 0.05 mM isopropyl-*β*-D-thiogalactoside (IPTG), and harvested at an OD₆₀₀ of 0.6–0.8 for electrocompetent cell preparation. For strains harboring the reporter plasmids, pTarget or pNTarget was separately transformed into BL21(DE3). These strains were grown in LB medium with 34 *µ*g/mL kanamycin to OD₆₀₀ 0.6–0.8 before harvesting for competent cell preparation.

For the nuclease and target selection assays, 50 *µ*L of the prepared electrocompetent cells containing pCas12a3 were mixed with 100 ng of one of the following reporter plasmids: pTarget, pNTarget, pTNoPfs, or pTNoPr. Electroporation was performed using a 1 mm cuvette (Bio-Rad) with a Gene Pulser II system (Bio-Rad) (1800 V, 200 Ω, 25 *µ*F). Immediately after pulsing, 920 *µ*L of SOC recovery medium containing 0.05 mM IPTG was added. Cells were recovered for 60 minutes at 37 °C with shaking (200 rpm), serially diluted, and plated on LB agar plates containing 0.05 mM IPTG, 34 *µ*g/mL streptomycin, 34 *µ*g/mL kanamycin, and 0.2% L-arabinose to select for colonies carrying both the nuclease-expression plasmid and the reporter plasmid. After overnight incubation at 37 °C, transformation efficiency was calculated by colony counts. All nuclease and target selection experiments were performed in triplicate.

To test whether the *Ba*Cas12a3 effector complex exhibits additional *in vivo* activity that could affect bacterial growth, we performed the nuclease selection assay. In this assay, 50 *µ*L of electrocompetent cells containing either pTarget or pNTarget were mixed on ice with 100 ng of the pCas12a3 plasmid. Electroporation was performed under the same conditions described above. After recovery, cells were plated on selective LB agar containing 34 *µ*g/mL streptomycin, 0.2% L-arabinose, and 0.05 mM IPTG (no kanamycin). Plates were incubated overnight at 37 °C, colonies were counted, and transformation efficiency was calculated.

To assess growth under nuclease-targeting conditions, *E. coli* BL21(DE3) cells were co-transformed with pCas12a3 and either the target or non-target reporter plasmid. Transformants were recovered in SOC medium supplemented with 0.2% glucose to suppress expression of the nuclease and crRNA. After overnight incubation, cells were pelleted (5, 000 × g, 2 min), washed, and resuspended in LB medium to an initial OD₆₀₀ of 0.1. This suspension was used to inoculate LB medium containing kanamycin (34 *µ*g/mL), streptomycin (34 *µ*g/mL), 0.05 mM IPTG, and 0.2% (w/v) L-arabinose, which was dispensed into 48-well plates. Growth was monitored at 37 °C by measuring OD₆₀₀ every 10 min in a plate reader.

### Protein expression and purification

The recombinant plasmid pCas12a3 was transformed into *E. coli* BL21(DE3) competent cells. Transformants were selected on LB agar plates containing streptomycin (34 *µ*g/mL) and incubation at 37 °C for 20 h. A single colony was inoculated into LB medium with the same antibiotic and grown overnight. This pre-culture was diluted 1:100 into 1L of fresh LB medium and grown at 37 °C until OD_600_ reached 0.6-0.8. Protein expression was induced with 0.2 mM IPTG for 5 h at 37℃. Cells were harvested, resuspended in Buffer A (25 mM Tris-HCl pH 8.0, 500 mM NaCl, 2 mM MgCl_2_, 10% glycerol), and lysed by sonication. After centrifugation, the supernatant was filtered through a 0.22 *µ*m membrane and loaded onto a Ni-NTA affinity column. The column was washed with Buffer A, followed by a wash with 8% Buffer B (Buffer A supplemented with 500 mM imidazole). Bound protein was eluted using a linear gradient of 0-100% Buffer B. Peak fractions were pooled and concentrated. The concentrated sample was further purified by size-exclusion chromatography (SEC) using a Superdex 200 Increase 10/200 GL column (Cytiva) equilibrated with Buffer C (25 mM Tris-HCl pH 8.0, 150 mM NaCl, 2 mM MgCl_2_, 5% glycerol). Chromatography was performed on an ÄKTA purifier system (Cytiva). The main peak fractions were collected, analyzed by SDS-PAGE and western blot, concentrated, and stored at −80 °C. The same purification procedure was used for the *Ba*Cas12a3-crRNA-target RNA complex and all mutant proteins.

### *In vitro* nucleic acid cleavage assays

Cleavage assays were performed to characterize the substrate specificity and collateral activity of the *Ba*Cas12a3-crRNA complex. Reactions (10 *µ*L) were prepared in cleavage buffer (50 mM Tris-HCl, pH 8.0, 10 mM MgCl₂) containing specified concentrations of effector complex and substrate. Substrates included FAM-labeled ssDNA, dsDNA, or ssRNA oligonucleotides, that were either complementary or non-complementary to the crRNA. Fluorescent tRNA substrates (5′-Cy5, 3′-Cy2) were obtained commercially (Tsingke Biotechnology Co., Ltd.). All oligonucleotide sequences are listed in Supplementary Tables S3. For target-activated cleavage, reactions containing 50-500 nM *Ba*Cas12a3-crRNA and 100 nM oligonucleotide were incubated at 37 °C for 60 min, stopped by addition of 10×DNA loading dye (Vazyme), and analyzed by 12% PAGE. Gels were visualized using fluorescence imaging (Amersham ImageQuant 800, Cytiva). To assess collateral activity, 10 *µ*L reactions containing 100 nM *Ba*Cas12a3-crRNA, 100 nM cognate target RNA, and 100 nM fluorescence-labeled substrates (ssDNA, dsDNA, or ssRNA) in cleavage buffer were incubated under the same conditions and analyzed as described.

For tRNA cleavage assays, 3 tRNA substrate types were tested: fluorescently labeled tRNA, *in vitro*-transcribed tRNA and total tRNA enriched from *E. coli* according to the reported method (39). To generate *in vitro*-transcribed tRNA, tRNA genes from *E. coli* BL21 were transcribed with a T7 promoter sequence (5′-TAATACGACTCACTATAGGG-3′) introduced upstream of each tRNA sequence. Due to the short length of these genes (75-90 bp), each full-length sequence was amplified using two overlapping primers (∼20 bp overlap; see Supplementary Table S2 and S3). The resulting PCR products, containing the T7 promoter, were purified by ethanol precipitation and used as templates for *in vitro* transcription with a commercial kit (JT101-02, TransGen Biotech) according to the manufacturer instructions. Transcription from the T7 promoter added a 5′-GGG overhang to each tRNA, which did not alter the predicted secondary structure of the transcripts. Reactions contained 1 *μ*M *Ba*Cas12a3, 1 *μ*M target RNA, and 1 *μ*M of tRNA in cleavage buffer, incubated at 37 °C for 60 min, and stopped with 2 × RNA loading dye (New England Biolabs). Products were separated on a 16% TBE-urea polyacrylamide gel, stained with ethidium bromide, and visualized using a gel imaging system (JIAPENG ZF-288, Shanghai Jiapeng Technology Co., Ltd.). For assays with fluorescently labeled tRNA (5′-Cy5, 3′-Cy2), reactions were performed under identical conditions and visualized by fluorescence imaging (Amersham ImageQuant 800, Cytiva).

### crRNA and transcriptome sequencing

The *Ba*Cas12a3-crRNA complex was purified as described, and total RNA was extracted using TRIzol reagent (Invitrogen), followed by Turbo DNase (Life Technologies) treatment to remove DNA. RNA-seq libraries were prepared and analyzed by GENEWIZ, Azenta Life Sciences on an Illumina NovaSeq 6000 platform. Sequencing libraries were prepared from the extracted RNA, and then sequenced and analyzed by GENEWIZ from Azenta Life Sciences on an Illumina Novaseq600 platform. To assess *Ba*Cas12a3 cleavage specificity, we performed RNA-seq on *E. coli* BL21(DE3) strains expressing *Ba*Cas12a3 containing pCas12a3 plasmid and either a target or non-target plasmid. Cells were grown in LB medium (34 μg/mL kanamycin and streptomycin) to OD600 of 0.6, chilled on ice, and resuspended in fresh LB (37 °C). Nuclease and target expression were induced with 0.1 mM IPTG and 0.4% (w/v) L-arabinose. After 3 h, samples were collected, and their transcriptomes were sequenced (Shanghai Majorbio Bio-pharm Technology).

### PFS specificity profiling

To identify DNA motifs in the PFS required for *Ba*Cas12a3 interference, a high-throughput plasmid depletion assay was performed. A library of potential target sequences was constructed by inserting a randomized 5-nucleotide (5′-NNNNN-3′) region into the PFS. Complementary oligonucleotides (Target-Flank-F and Target-Flank-R) containing the random region were annealed (95 °C for 5 min, followed by gradual cooling to 4 °C at 0.1 °C per 15 s) and ligated into a pBAD vector downstream of the *ara*BAD promoter. The ligation product was transformed into *E. coli* DH5*α*, and plasmid DNA was extracted from >10⁴ colonies to generate the unselected input library. For positive selection, the input library was electroporated into *E. coli* BL21 cells expressing *Ba*Cas12a3 and its cognate crRNA from plasmid pCas12a3. Transformants were selected on LB agar containing 0.05 mM IPTG, 0.2 % (w/v) L-arabinose, kanamycin, and streptomycin. Plasmid DNA was recovered from >10⁴ surviving colonies to generate the output library following *Ba*Cas12a3-mediated depletion of functional PFS sequences. The randomized region from both libraries was amplified and deep-sequenced. The proportion of each PFS variant in the input and output libraries was calculated and log_10_-transformed. The difference between these log-values served as a quantitative measure of selection: a positive value indicates enrichment (functional PFS), while a negative value indicates depletion (impaired function). Sequence enrichment and depletion patterns were visualized as a heatmap and as sequence logos. In this analysis, a positive difference (>0) indicates that the PFS likely possesses functional activity, whereas a negative difference (<0) suggests functional impairment, with zero serving as the statistical threshold for functional classification. Sequence enrichment and depletion patterns were analyzed and visualized using Heatmapper (40) and sequence logos generated with WebLogo (41).

### Site-directed mutagenesis

Specific substitutions were introduced into the target plasmid using site-directed mutagenesis. Complementary primers containing the desired mutations (Supplementary Table S2) were used for whole-plasmid PCR amplification with a high-fidelity DNA polymerase (Phanta Max Super-Fidelity DNA Polymerase, Vazyme Biotech Co., Ltd.). The PCR product was treated with *DpnI* (Thermo Fisher Scientific, Waltham) at 37 °C for 1 h to digest the methylated template DNA. Then circularized using a recombinase assembly mix (ABclonal Technology Co., Ltd.). The reaction was transformed into chemically competent *E. coli* DH5*α* cells. Transformants were selected on LB agar plates containing streptomycin and incubated overnight at 37 °C. Individual colonies were cultured, and plasmids were extracted for verification by Sanger sequencing (Tsingke Biotechnology Co., Ltd.).

### SOS response induction assay

To determine whether *Ba*Cas12a3 activates the DNA damage mediated bacterial SOS response, *E. coli* BL21 was co-transformed with the pCas12a3 plasmid and either a target pTarget or non-target pNTarget reporter plasmid. Transformants were recovered overnight in SOC medium supplemented with 0.2% glucose to repress nuclease and crRNA expression. Cells were harvested, resuspended in LB medium with appropriate antibiotics, and grown to an OD_600_ of ∼0.6. After chilling and washing, cultures were resuspended in antibiotic-free LB. Nuclease and crRNA expression was induced with 0.1 mM IPTG for 30 min, followed by induction of the target plasmid with 0.4% L-arabinose. Samples were collected immediately before and at 1 h intervals for up to 5 h after arabinose addition. Cells were pelleted, resuspended in PBS (8.0 g/L NaCl, 0.2 g/L KCl, 3.63 g/L Na_2_HPO_4_·12H_2_O, 0.24 g/L KH_2_PO_4_, pH 7.4), and lysed by sonication. Total protein concentration in the lysate supernatant was determined using the Bradford assay. (42). Equal amounts of protein from each time point were subjected to western blot analysis. RecA was monitored using an anti-RecA antibody (Abcam plc, Cambridge, UK); an antibody against the RNA polymerase *α*-subunit (BioLegend, San Diego, CA, USA) served as the loading control.

### Cryo-electron microscopy

The *Ba*Cas12a3-crRNA binary complex was overexpressed in *E. coli* BL21 (DE3) cells containing the pCas12a3 plasmid, followed by purification via nickel-affinity and SEC. To form the ternary BaCas12a3–crRNA–target RNA complex, the purified binary complex was incubated with a 41-nt synthetic target RNA (Tsingke) at a 1:1.5 molar ratio for 1 h at 15 °C. The mixture was further purified by SEC on a Superdex 200 Increase 10/300 column (Cytiva) equilibrated with cryo-EM buffer (25 mM Tris-HCl pH 8.0, 150 mM NaCl, 2 mM MgCl₂, 5% glycerol).

For grid preparation, 4 *μ*L of each complex (0.3 mg/mL for binary, 0.5 mg/mL for ternary) was applied to glow-discharged Quantifoil R1.2/1.3 200-mesh copper grids. After 4 s blotting, grids were plunge-frozen in liquid ethane using a Vitrobot Mark IV (Thermo Scientific). Cryo-EM data were collected on a Glacios (Thermo Scientific) microscope operated at 200 kV. Images were acquired using EPU software at a nominal magnification of ×130, 000 (calibrated pixel size 0.89 Å/px), with a defocus range of - 0.6 to −1.6 *µ*m. Movies were recorded on a Falcon 4i direct electron detector equipped with a Selectris X energy filter (10 eV slit width), at a dose rate of 10 e⁻ /px/s over 3.35 s, yielding 40 frames and a total dose of ∼40 e^−^/Å^2^. A total of 5, 823 and 4, 581 movies were collected for the binary and ternary complexes, respectively (Supplementary Fig. S7, S8).

### Image processing, model building, refinement, and validation

Data processing was performed in cryoSPARC (v4.5.3). (43). For the *Ba*Cas12a3–crRNA complex, extracted particles underwent multiple rounds of 2D classification to remove false picks. Particles from selected classes were used to generate six ab initio models. Following three rounds of heterogeneous refinement, one class (Class 1) exhibited well-defined, high-resolution features and was selected for further 3D refinement (43, 44). The other classes showed poorly resolved density (Extended Data Fig. S7). A final set of 484, 846 particles from Class 1 yielded a reconstruction at 2.91 Å resolution. For the *Ba*Cas12a3–crRNA–target RNA complex, a similar processing strategy was applied. A final set of 530, 902 particles was refined to a global resolution of 2.87 Å (Extended Data Fig. S8). These particles subsequently underwent 3D refinement and 3D classification (43, 44), yielding five classes that were analyzed for conformational heterogeneity. Atomic models for both complexes were built *de novo* in Coot, using predicted structures from AlphaFold 3 as initial guides (34). The models were refined against their respective cryo-EM maps using real-space refinement in Phenix (45, 46) and validated with MolProbity (47). Structural figures were prepared using UCSF ChimeraX (version 1.10) (48). Atomic coordinates have been deposited in the Protein Data Bank. A detailed summary of cryo-EM data collection and processing is provided in Supplementary Table S4.

## Results

### Identification and phylogenetic analysis of Cas12a3

The discovery of Cas12a3 originated from sequence-based mining of metagenomic databases. Using the well-characterized DNA-targeting nuclease Cas12a (also known as AcCpf1) as an initial query, researchers performed searches against two expansive metagenome databases (24, 30, 31). Promising initial hits from these searches were then used as seeds for iterative BLAST searches against the NCBI non-redundant protein database (49). This bioinformatic approach successfully identified several distinct CRISPR loci associated with a putative *cas12* gene, which were often annotated as transposases or hypothetical proteins, suggesting they could represent novel, uncharacterized CRISPR-Cas systems. Construction of phylogenetic trees with these nuclease sequences and representative sequences retrieved from other Cas12 subtypes revealed that Cas12a3 sequences form an independent evolutionary clade, sharing only 10–20% sequence identity with either Cas12a (DNA-targeting) or Cas12a2 (RNA-targeting) (Fig. 1A and Fig. S1). This divergence underscores the unique evolutionary trajectory and functional specialization of Cas12a3. Interestingly, several Cas12a3 were identified in ruminant gut microbiome, and a representative Cas12a3 encoded by a *Bacteroidales* (*Ba*Cas12a3) was chosen for further study.

Cas12 subtypes exhibit an unprecedented diversity not only in sequence but also in mechanisms of action, and they are believed to derive from their TnpB ancestors present in IS elements of IS605 and IS607 families (7, 8, 11, 16, 50). A unifying feature conserved across all characterized Cas12 proteins and their TnpB ancestor is the presence of a RuvC nuclease domain, which forms the catalytic core (8). Consistent with this, AlphaFold3 structural modeling confirms that the RuvC domain of *Ba*Cas12a3 closely resembles that of the RNA-targeting Cas12a2. Notably, 3 catalytical amino acid residues are conserved both in the primary sequence alignment (Fig. S2A) and in their relative three-dimensional positions within the predicted structure of *Ba*Cas12a3 (Fig. S2B), strongly suggesting a shared enzymatic mechanism. Intriguingly, bioinformatic analysis revealed that several identified Cas12a3 members (marked in Fig. S1) possess an additional domain of unknown function that is completely absent from all other known Cas12 subtypes (Fig. S2), suggesting functional/mechanistic innovation of this unique group of Cas12a3. Examination of the genomic loci encoding CRISPR-Cas12a3 systems shows they also contain core adaptation genes *cas1*, *cas2*, and often *cas4*, and CRISPR arrays, suggesting these CRISPR systems are active in spacer acquisition (51). These Cas12a3 systems are further associated with other accessory genes, including TPR-DUF3800 proteins or Cas12a systems (Fig. 1B). This genetic architecture supports their role as functional immune systems. Collectively, these results suggested that Cas12a3 systems could represent a new Cas12 subtype, exhibiting mechanistically diversified antiviral mechanisms.

### *Ba*Cas12a3 induces immune activity with RNA-targeting

In order to test if this CRISPR-Cas12a3 could be functional in *E. coli*, a heterologous host, we constructed an expression plasmid (pCas12a3) harboring both the *BaCas12a3* gene and a CRISPR array. Plasmid-borne expression of the *cas* gene and CRISPR array in *E. coli* yielded *Ba*Cas12a3-crRNA ribonucleoprotein complexes (RNPs) that were purified by SEC (Fig. S3). The RNA components were then extracted from the RNPs and characterized by small RNA sequencing, and this revealed mature crRNAs containing a 19-nt repeat linked to a ∼23-nt spacer, indicating the *Ba*Cas12a3 pre-crRNAs were processed in *E. coli* cells probably by the *Ba*Casl2a3 protein itself (Fig. S3).

*Ba*Cas12a3 was implicated in mediating antiviral defense by RNA-targeting since its RuvC domain resembles that of Cas12a2, an RNA-targeting system that executing immunity in PFS-dependent fashion (20, 21). We employed PFS-depletion assay (Fig. S4A) to identify functional PFS motifs for *Ba*Cas12a3. A DNA ligation was set up with pBAD and diverse protospacer DNA fragments carrying 5 random nucleotides at the predicted PFS position (3′-NNNNN-target-5′) (Fig. 2A). Transformation of *E. coli* DH5*α* with the ligation yielded transformants that were used for two experiments: (a) the bacterial cells were employed for extracting plasmid DNA to generate a library of protospacer plasmids (the unselected input library), and (b) the bacterial cells were transformed with pCas12a3, and cultured for 8 hours during which plasmid-borne expression of the *Ba*Cas12a3 RNPs and target RNAs would induce immunity against the host cells. As a result, PFS-carrying plasmids were selectively depleted from the culture. Extraction of plasmids from the bacterial cells yielded the output library. DNA fragments carrying the PFS region were amplified from the input and output libraries and sequenced by high-throughput sequencing. Analysis of their sequence reads revealed 54% (550/1024) of DNA motifs at the PFS position in the protospacer plasmid library were effectively depleted by the nuclease (Fig. S4B). Analysis of the depleted sequences identified a consensus motif with invariable AA nucleotides at −2 and −3 positions relative to the protospacer (Fig. 2B and Fig. S4B).

**Figure 2.**
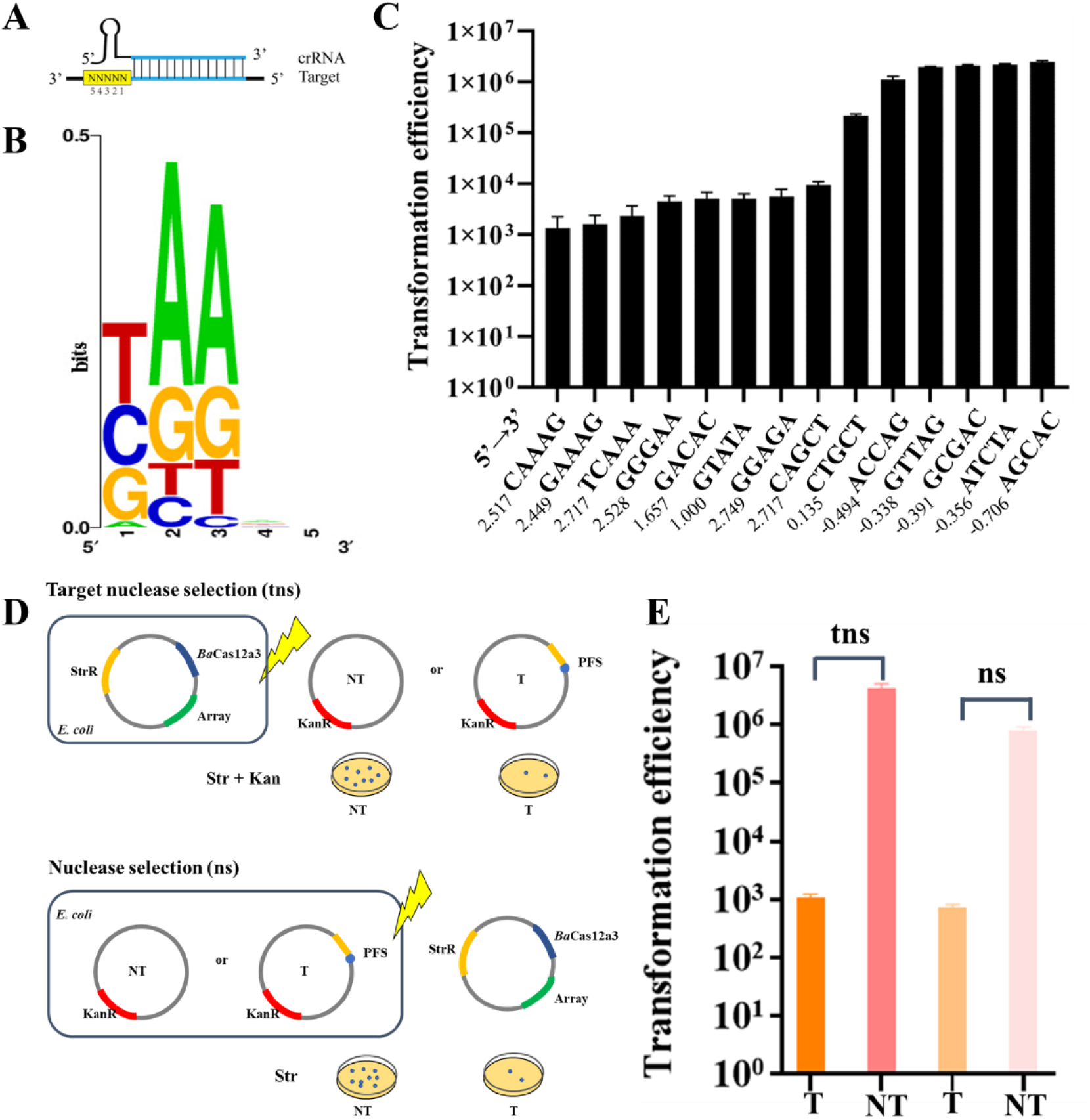
PFS-depleting screening and immune activity assessment by plasmid interference assays. (A), Schematic of the randomized 5-nucleotide region at the predicted PFS position. (B), Sequence logo of the depleted PFS motif, generated from screening data. The logo, generated from the depletion data, reveals a marked enrichment for adenine (A) at the −2 and −3 positions, defining the core specificity determinant for *Ba*Cas12a3. (C), Validation of interference for selected PFSs. Plasmids containing PFS motifs with varying depletion values (from Fig. S4B) were tested individually. Sequences containing the characteristic AA motif at the −2/−3 positions (high positive values) conferred strong interference (reduced transformation), whereas sequences lacking this motif (low negative values) showed no significant interference. All depletion values were shown in Fig. S4B. Data represent mean ± s.d. of three biological replicates. (D), Schematic of the target nuclease selection (tns) and nuclease selection (ns) plasmid interference assays. T, plasmid encoding the target RNA (pTarget); NT, reference plasmid with no target sequence (pNTarget); Kan, kanamycin; Str, streptomycin. (E), Plasmid transformation efficiency for *Ba*Cas12a3 under target plasmid selection versus nuclease plasmid selection.

To validate the PFS-depleting results, DNA motifs with high depletion values (Fig. S4B) were selected for testing their capability of mediating plasmid interference assays in *E. coli*. We found that several highly depleted DNA motifs (*e.g.* 5′-CAAAG-3′, 5′-GAAAG-3′, 5′-TCAAA-3′, 5′-GGAGA-3′) conferred significantly higher levels of transformation fold reduction compared to non-depleted ones (Fig. 2C). One of such motifs, 5′-GAAAG-3′ was chosen as an optimal PFS motif for Cas12a3 and employed in all subsequent *in vivo* and *in vitro* experimental assays.

To gain a further insight into the antiviral defense by the *Ba*Cas12a3 system, two plasmids were employed: pCas12a3 carrying *cas12a3* gene and a cognate mini-CRISPR array and pTarget containing a corresponding protospacer for expressing the cognate target RNA complementary to the spacer in the mini-CRISPR array (Fig. 2D). Introduction of each plasmid into *E. coli* yielded two strains for interference assay: one carrying pCas12a3 and the other containing pTarget. The former was then transformed with pTarget while the latter, with pCas12a3. Scoring for both plasmids in the host expressing *Ba*Cas12a3 upon pTarget transformation in the first experiment revealed > 10^3^-fold lower rate of transformation than that of a reference plasmid pNTarget (Fig. 2E). Since survivals should carry both plasmids, only bacterial cells with escape mutation of the *Ba*Cas12a3 immunity, *i.e.*, mutation in *Bacas12a3* gene, spacers or target sequence, can form colonies on the selective plates. Thus, *Ba*Cas12a3 executes strong immunity against pTarget. Transformation of pTarget-carrying cells with pCas12a3 upon showed the transformation efficiency was comparable to the occurrence of escape mutation in the first experiment (Fig. 2E), suggesting Cas12a3 operates through influencing host cell viability rather than specific plasmid targeting, as shown for Cas12a2 (20).

To further determine the genetic determinants for activating the *Ba*Cas12a3 system in *E. coli*, two additional plasmids were constructed, including pTNoPfs (producing target RNAs with 5′-AGCAC −3′, a non-functional motif), and pTNoPr (not producing target RNA), and these plasmids were employed in plasmid interference assay along with pTarget (producing target RNAs carrying the 5′-GAAAG-3′ PFS), pNTarget (without target with 5′-GAAAG-3′ PFS) (Fig.3A). We found that a drastic reduction in transformation efficiency by > 1, 000-fold only for pTarget whereas the absence of any element i.e., a non-functional PFS, a non-target sequence, or a missing promoter, restored the transformation efficiency, demonstrating an absolute requirement for the cognate target RNA for *Ba*Cas12a3 immune responses (Fig. 3B). Growth experiments showed that, in contrast to normal growth occurred for the reference cultures, robust growth inhibition was observed exclusively for cultures containing both the *Ba*Cas12a3 nuclease and the complete target plasmid (Fig. 3C), consistent with the results obtained with plasmid interference assay (Fig. 3B).

**Figure 3.**
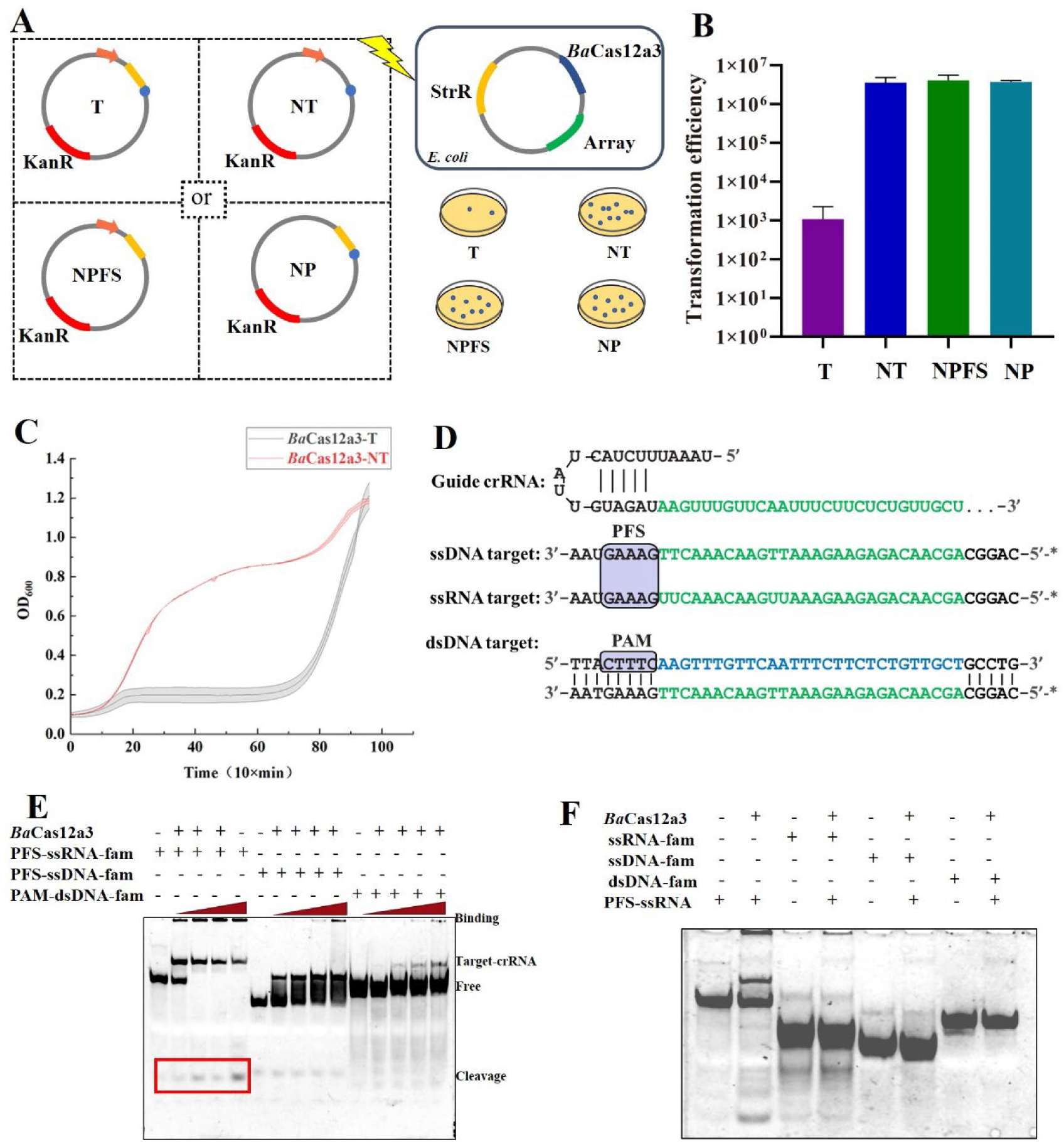
RNA-targeting activated *Ba*Cas12a3. (A), Experimental design to identify requirements for *Ba*Cas12a3 activation. StrR, streptomycin resistance; KanR, kanamycin resistance. Plasmids used: T (pTarget, contains target RNA, PFS, and promoter); NT (pNTarget, contains PFS and promoter but no target); NPFS (pTNoPfs, contains target and promoter but no PFS); NP (pTNoPr, contains target and PFS but no promoter). (B), Plasmid transformation efficiencies for *Ba*Cas12a3 upon introduction of each plasmid. (C), Bacterial growth curves during induction of plasmid interference. (D), Schematics of the nucleic acid targets used for in vitro assays. The asterisk denotes a 5′ FAM label. (E), Direct cleavage of FAM-labeled substrates by purified *Ba*Cas12a3–crRNA complex. The red box highlights cleavage products. (F), Assay for collateral cleavage activity. FAM-labeled non-target substrates were incubated with *Ba*Cas12a3–crRNA complex in the presence of target RNA.

Different fluorescently labeled nucleic acid substrates, including single-stranded RNA (ssRNA), single-stranded DNA (ssDNA), and double-stranded DNA (dsDNA), were employed to test the nuclease activities of the immune system using purified *Ba*Cas12a3 RNP. Each substrate contained a target strand that is complementary to the crRNA guide carrying the identified PFS sequence (Fig. 3D), and cleavage of nucleic acids yielding fragments carrying 5′-FAM fluorophores that can be detected by fluorometer. We found, *Ba*Cas12a3 cleaved its target RNA but failed to cleave ssDNA or dsDNA. To test if *Ba*Cas12a3 could exhibit any collateral nuclease activity, the *Ba*Cas12a3 enzyme was incubated with FAM-labeled ssRNA, ssDNA, or dsDNA in the presence of the cognate target RNA (containing PFS), but we failed to detect any activity towards these nucleic acid substrates (Fig. 3F).

### Collateral activity towards tRNA

CRISPR-Cas12a2, along with Type III and Type VI systems, is characterized by an RNA-targeting immune response (20, 23, 52, 53). In these systems, recognition of a cognate RNA guide activates the effector nuclease. This activation typically induces potent, indiscriminate nuclease activity that degrades cellular nucleic acids, triggering a dramatic abortive infection response to protect the bacterial population.

Building upon the established mechanism of the related effector Cas12a2—which, upon target recognition, exhibits collateral cleavage of double-stranded DNA, thereby triggering the RecA-dependent SOS response and genomic damage (20)—we sought to precisely differentiate. *E. coli* BL21 co-transformed with pCas12a3 and pTarget plasmids was grown to OD600 ∼0.6 and induced to activate *Ba*Cas12a3. Samples were collected hourly for up to 5 h for Western blot analysis. As a positive control, induction of the Cas12a2 system resulted in a significant increase in RecA expression, confirming the expected DNA damage response (Fig. 4A). In contrast, activation of *Ba*Cas12a3 by its target RNA did not alter RecA expression levels, indicating no detectable DNA damage during its immune reaction (Fig. 2C). This clear distinction suggests that *Ba*Cas12a3 employs a immune mechanism distinct from Cas12a2.

**Figure 4.**
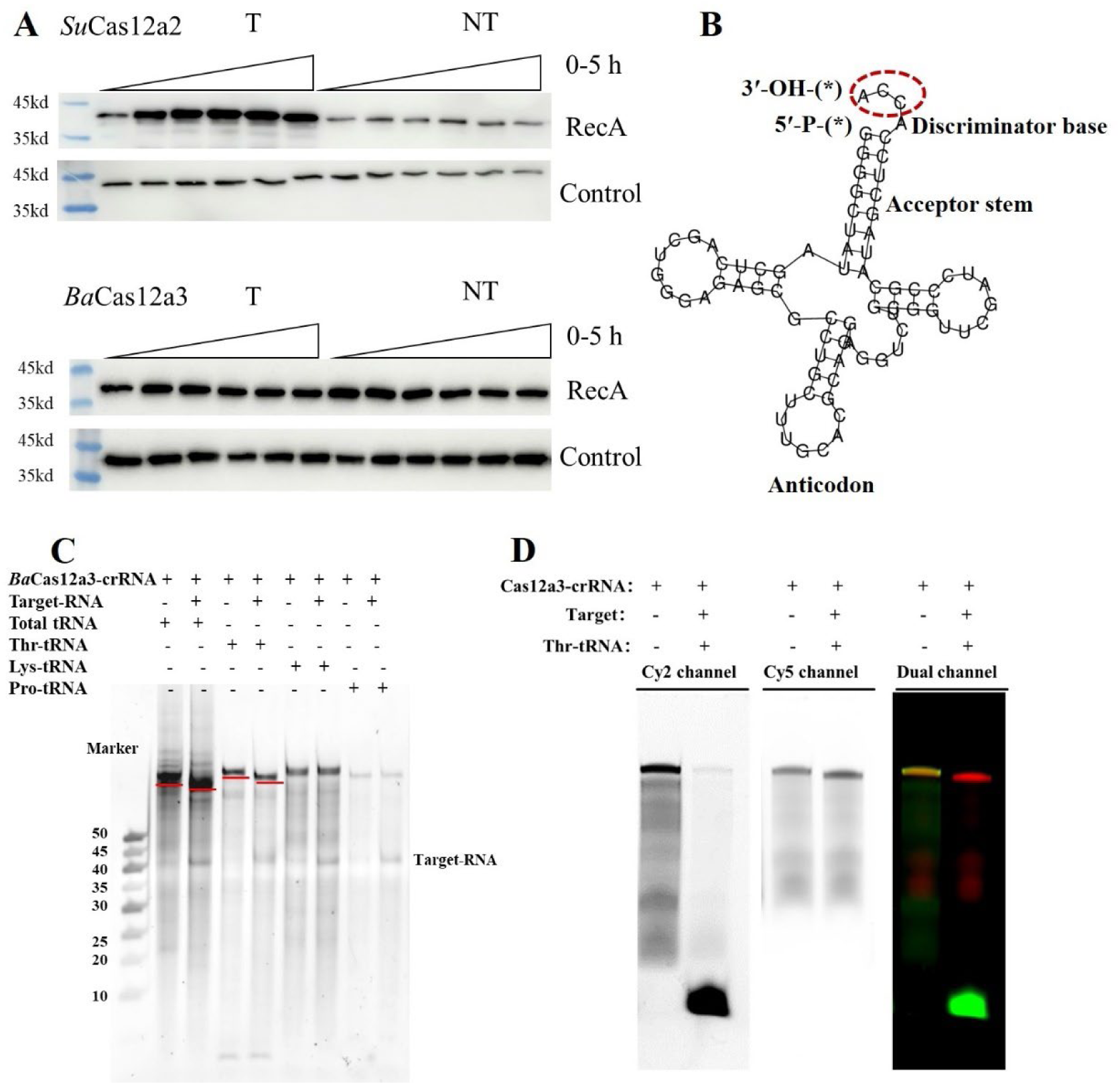
Western blot analysis of RecA expression and collateral cleavage of tRNAs. (A), Western blot analysis of RecA protein levels in strains expressing *Ba*Cas12a3 and *Su*Cas12a2 after induction of interference. The RNA polymerase *α* subunit serves as the internal reference. Data are representative of three independent experiments. (B), Schematic of a tRNA substrate. The asterisk indicates the position of a fluorescent label. (C), Detection of tRNA collateral cleavage by gel electrophoresis. The red line indicates the band shift corresponding to cleaved product. (D), Collateral cleavage of a dual-labeled Thr-tRNA. The substrate was labeled with Cy5 at the 5′ end and Cy2 at the 3′ end. Cleavage products were visualized by single-channel and merged fluorescence imaging. The Cy2 channel detects the 3′ fragment, the Cy5 channel detects the 5′ fragment, and the merged image shows co-localization of the intact tRNA (pre-cleavage) and separation of the fragments (post-cleavage).

To gain an insight into how *Ba*Cas12a3 could inhibit the growth of the host, cell samples were taken from cultures of activated versus non-activated *Ba*Cas12a3 from which total RNAs were prepared and analyzed by RNA-seq. We found that SOS genes were not activated in *Ba*Cas12a3-expressing cells, which is in contrast to the data of Cas12a2 (20). Strikingly, the transcription levels of several tRNAs increased over 300-fold within 3 hours of induction (Fig. S5). This massive upregulation indicates a severe disruption of tRNA homeostasis, suggesting BaCas12a3 may have impaired the function of these tRNAs in vivo.

(Some of) these tRNA species were then tested as potential substrates for activated *Ba*Cas12a3, electrophoretic analysis revealed a slight but consistent band shift for some species, suggesting possible tRNA degradation/cleavage (Fig. 4C and Fig. S6). To precisely map the cleavage site, we employed fluorescently labeled Thr-tRNA substrates. This assay definitively demonstrated that *Ba*Cas12a3 efficiently cleaves the tRNA molecule at its 3′ end (Fig. 4D). This cleavage is highly consequential, as the integrity of the 3′-CCA terminus is essential for tRNA aminoacylation by aminoacyl-tRNA synthetases and for subsequent ribosomal interactions during translation (54). By cleaving off this terminal sequence, activated *Ba*Cas12a3 irreversibly inactivates the tRNA. This provides a direct mechanistic explanation for the profound growth inhibition observed *in vivo* upon *Ba*Cas12a3 activation (Fig. 3C). The cleavage of tRNAs effectively blocked the cellular translation, linking the targeted recognition of tRNA molecules to a fatal disruption of an essential housekeeping process and thereby executing an immune function.

### Structural features of *Ba*Cas12a3

Using cryo-EM, we determined the structures of the *Ba*Cas12a3–crRNA binary complex at 2.91 Å resolution and the *Ba*Cas12a3–crRNA–target RNA ternary complex at 2.87 Å resolution (Figs. S7 and S8). Atomic models were built based on the two corresponding maps (Fig. 5). The domain organization of *Ba*Cas12a3, as revealed by these structural models, is presented in Fig. 5A. *Ba*Cas12a3 adopts the conserved bilobed architecture characteristic of the Cas12 protein family, featuring an N-terminal recognition (REC) lobe and a C-terminal nuclease (NUC) lobe, connected by a flexible linker (residues 694–709) (9, 55). In both complexes, the crRNA and the crRNA–target RNA duplex are positioned within the central channel formed between the two lobes (Figs. 5B and 5C). The REC lobe comprises of the REC1, REC2, wedge (WED), and protospacer-flanking site-interacting (PI) domains. REC1 and REC2 domains primarily facilitate crRNA binding and are responsible for the initial recognition and verification of the target RNA sequence (21), while the WED and PI domains are critical for recognizing both the crRNA and the PFS (Fig. 5C). Compared to the Cas12a2 ortholog, *Ba*Cas12a3 incorporated a streamlined REC2 domain (Fig. 6A). Notably, in the binary complex of *Ba*Cas12a3, the PI domain mediates extensive interactions with the crRNA through hydrogen bonding and *π*–*π* stacking involving key residues such as Arg473, Gln476, Glu517, and Phe518 (Fig. S9). These interactions result in an extended, ordered conformation of the crRNA 3′ region. In contrast, comparative analysis with the Cas12a2 binary complex (PDB: 8D49) reveals that neither a structurally defined PI domain nor an ordered 3′ crRNA extension is present in that structure (21) (Fig. S9). Despite these architectural differences in the REC2 and PI domains, the REC lobe of *Ba*Cas12a3 maintains functional parallels with SuCas12a2, as both orthologs preserve conserved mechanisms for crRNA and target RNA binding (21).

**Figure 5.**
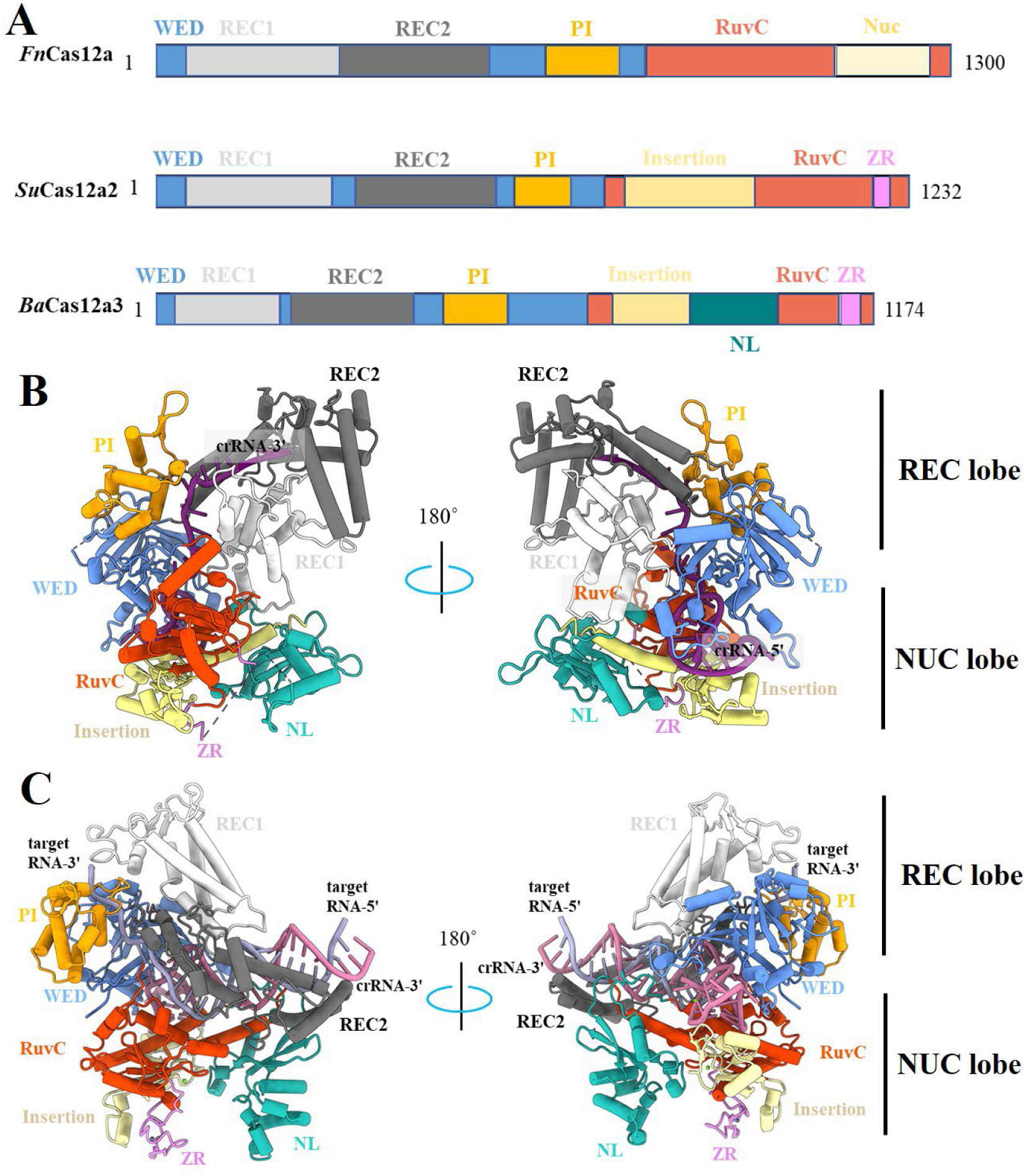
Overall structure of the *Ba*Cas12a3 binary and ternary complexes. (A), Schematic of domain organizations of *Fn*Cas12a, *Su*Cas12a2 and *Ba*Cas12a3 (21, 24). (B), Atomic model of the *Ba*Cas12a3–crRNA complex in two views. Domains are colored as follows: RuvC (orange red), WED (blue), REC1 (white), REC2 (gray), PI (orange), Insertion (yellow), ZR (pink), and the new NL domain (sea green). The crRNA is shown in purple. (C), Atomic model of the *Ba*Cas12a3–crRNA–target RNA complex. The model was showed and colored as in (B), with the target RNA in magenta.

**Figure 6.**
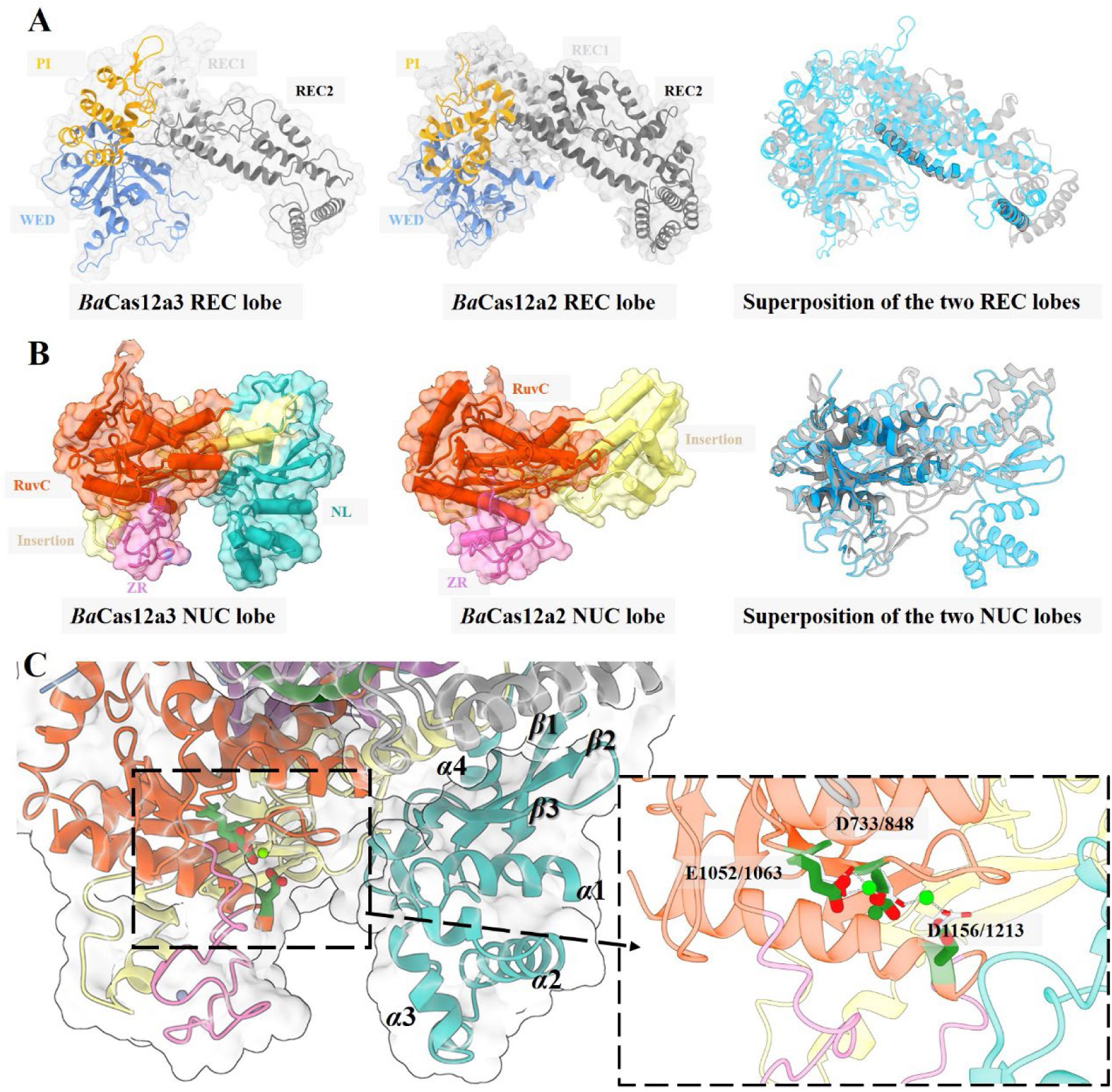
Structural comparison of *Ba*Cas12a3 with *Su*Cas12a2 (21). (A) REC lobes of *Ba*Cas12a3 and *Su*Cas12a2, and their structure superposition. (B) The NUC lobes of *Ba*Cas12a3 and *Su*Cas12a2, and their structure superposition. (C) The catalytic residues in the RuvC domain of *Ba*Cas12a3 and their equivalents in *Su*Cas12a2, shown in stick representation. The carbon bond for *Ba*Cas12a3 catalytic residues was colored in green and their equivalents in *Su*Cas12a2 in white. The secondary structural elements in the NL domain were marked. Domains are colored as in Fig.5, with a transparent surface overlaid.

The C-terminal NUC lobe of *Ba*Cas12a3 contains the essential catalytic core and several auxiliary domains critical for its unique function. Central to this lobe is the RuvC domain (Fig. 6B), which adopts the canonical RNase H-like fold that is characteristic of Cas12 nucleases (56). Embedded within this fold is the conserved catalytic triad (D733, E1052, and D1156), which is indispensable for magnesium ion coordination and phosphodiester bond cleavage (Fig. 6C). Flanking the RuvC domain are several key structural modules. The insertion domain, positioned adjacent to RuvC, exhibits distinct structural adaptations in *Ba*Cas12a3, including an N-terminal extension that wraps behind the ZR domain to provide critical structural reinforcement and stability to the entire lobe (Fig. 6C). Notably, *Ba*Cas12a3 possesses a novel nucleic acid-loading (NL) domain. This domain, composed of 3 antiparallel *β*-strands and 4 *α*-helices, extends from the insertion domain to the periphery of the RuvC domain, forming a distinctive architectural feature (Fig. 6C). Furthermore, the RuvC domain is associated with the ZR domain incorporating two conserved zinc-finger motifs (CXXC/CXXXC), which is also occurring in Cas12a2/f/e/m/g orthologs (18, 21, 27–29). Spatially, the ZR and NL domains are positioned on opposite sides of the RuvC domain. Together, they create a continuous, electropositive groove that is structurally optimized for binding and accommodating nucleic acid substrates. Combined with the previous results, we concluded that the groove is a defining feature of *Ba*Cas12a3 and is critical for loading target RNA and collateral tRNA molecules to the catalytic center for cleavage (Fig. 6C).

### Activation and allosteric gating of *Ba*Cas12a3 nuclease

Structural analysis of the binary complex revealed that the 5′ repeat region of the crRNA (19 nt) forms a pseudoknot. Comparative structural studies indicated that this 5′ region remains largely static upon target RNA binding (Fig. 7B; Supplementary Video 1). In contrast, the 3′ region of the crRNA undergoes a pronounced conformational rearrangement (Fig. 7B; Supplementary Video 1). Upon formation of the *Ba*Cas12a3–crRNA–target RNA ternary complex, PFS recognition is mediated by residues from the PI, WED, and REC1 domains (Fig. 7A), while the remainder of the target RNA is primarily recognized by the RuvC and REC2 domains. Concurrently, substantial conformational rearrangements occur in the REC1 (2–65 Å), REC2 (1–68 Å), NL (1–25 Å), and PI (1–14 Å) domains (Fig. 7B; Supplementary Video 1). Notably, unlike its Cas12a2 ortholog—where the C-terminal insertion domain undergoes marked conformational change upon target RNA binding—the homologous α-helix in the *Ba*Cas12a3 insertion domain remains conformationally rigid (Fig. 7B). The presence of the NL domain facilitates the formation of a positively charged groove with the adjacent REC2, RuvC, and ZR domains (Fig. 6C). In the binary complex, this groove is too narrow to accommodate nucleic acid substrates, and the catalytic site of the RuvC domain is buried and obstructed by the NL domain (Fig. 5B). Target RNA binding triggers a backward movement of the NL domain, thereby exposing the RuvC catalytic center (Figs. 7B and 7C; Supplementary Video 1). In contrast to the highly exposed RuvC active site observed in the Cas12a2 ternary complex (PDB: 8D4A) (Fig. 6B) (21), the NL domain in *Ba*Cas12a3 likely restricts substrate access and consequently limits its collateral nuclease activity.

**Figure 7.**
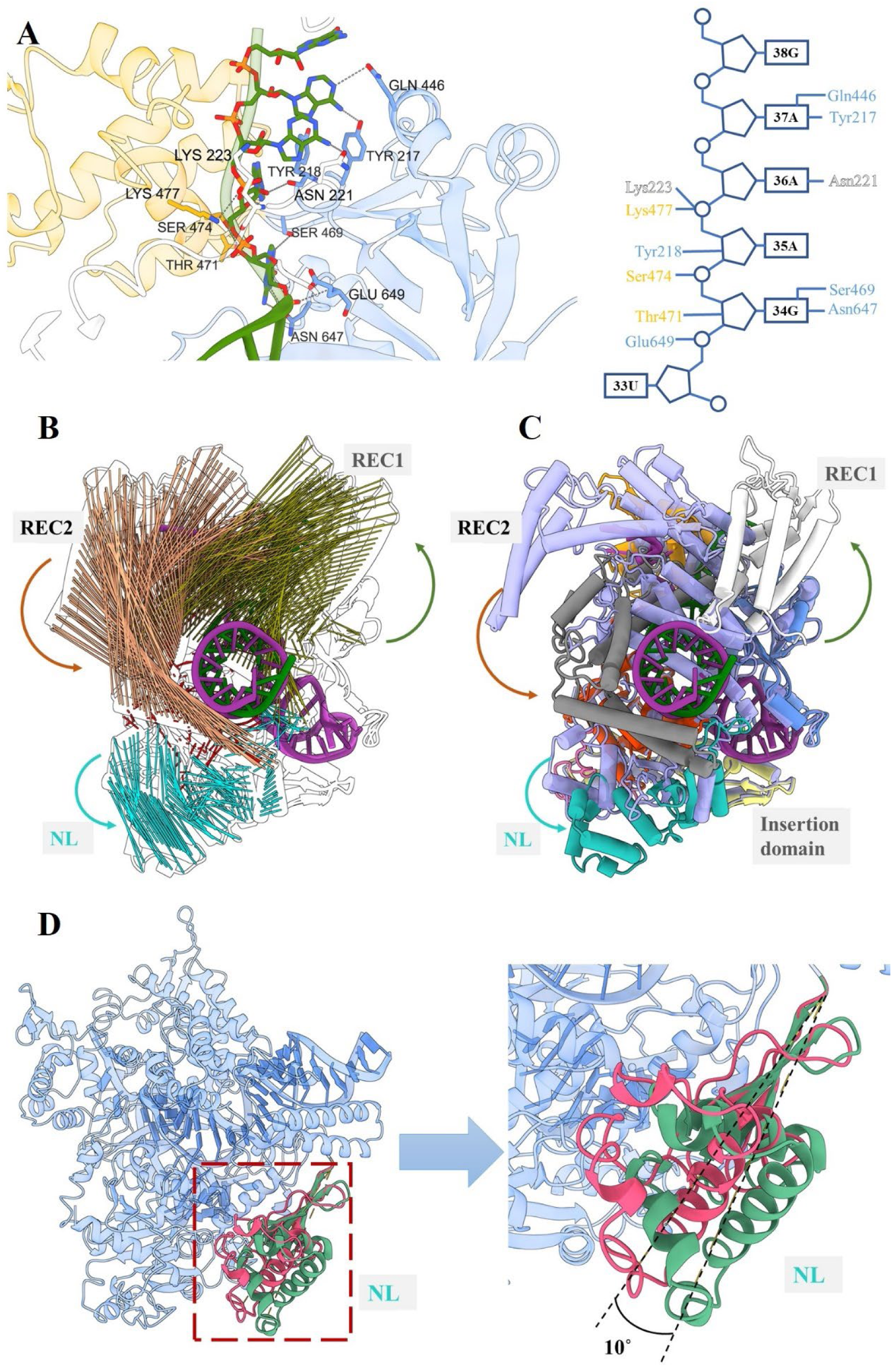
Target RNA binding induces structural rearrangements for activation. (A), Recognition of the PFS by residues from the PI, WED, and REC1 domains. (B), Motion vector map depicting conformational changes following ternary complex formation. Vectors are colored by domains: REC1(light green), REC2 (brown), NL (teal), and RuvC (orange). (C), Domain reorganization of *Ba*Cas12a3 showed in cylinder/stubs. The binary complex is shown in blue-magenta; domains in the ternary structural domains are colored as in Fig. 5. Related conformational changes are shown in Supplementary Video 1. (D), Conformational change of the NL domain, resolved by 3D classification of the ternary structure (Supplementary Video 2).

While dsDNA-targeting Cas12 subtypes rely on domains adjacent to RuvC—such as the TNB, Nuc, or TSL domains—to load both DNA strands into the catalytic pocket (13, 25, 28, 57–59), RNA-targeting subtypes utilize minimal zinc-binding modules for nucleic acid binding, such as the 37-aa Nuc domain in Cas12g or the 47-aa ZR domain in Cas12a2 (21, 29). In contrast, *Ba*Cas12a3 employs an additional NL domain. The NL and ZR domains, positioned on either side of the catalytic site, cooperate to position the tRNA for RuvC-mediated cleavage. To elucidate how tRNA substrates are loaded into the active site and cleaved in *trans*, we attempted to resolve complex structures using multiple strategies, including glutaraldehyde immobilization and testing different tRNAs. However, none of these approaches succeeded, likely due to the transient or dynamic nature of this interaction. Nevertheless, a 3D classification analysis of the existing ternary complex data revealed insightful conformational details (Fig. S8). This analysis revealed a significant and reproducible movement of the NL domain, which undergoes a rotational shift of approximately 10° relative to the core catalytic body (Fig. 7D; Supplementary Video 2). The observed “open” state likely represents a transient conformation that facilitates the initial entry and positioning of the bulky tRNA substrate. Following cleavage, a reversal of this motion—a “closing” state— may promote product release, thereby enabling catalytic turnover. Thus, the detected structural plasticity provides a compelling mechanistic framework for the tRNA substrate handling cycle of the enzyme.

To validate the enzymatic activity and immune mechanism of *Ba*Cas12a3, we performed site-directed mutagenesis. Initial investigations focused on the conserved catalytic triad (D733, E1052, and D1156) within the RuvC nuclease domain (Fig. S2 and Fig. 8B). To trigger immune activity, *E.coli* BL21 strains harboring both a mutant pCas12a3 plasmid (pCas12a3^D733A^, pCas12a3^E1052A^, or pCas12a3^D1156A^) (Table S1) and the pTarget plasmid were induced with IPTG and L-arabinose, and their growth curves were monitored. The results revealed that mutations in any of the 3 catalytic residues nearly abolished the inhibition of bacterial cell growth (Fig. 8C). Consistent with this cellular phenotype, *in vitro* cleavage assays demonstrated a complete loss of collateral tRNA cleavage activity for all 3 RuvC mutants (Fig. 8E). These results demonstrate that *Ba*Cas12a3 strictly depends on its RuvC domain for function and inhibits cell growth via a tRNA disabled mechanism, thus establishing a functional paradigm distinct from all other characterized Cas12 systems.

**Figure 8.**
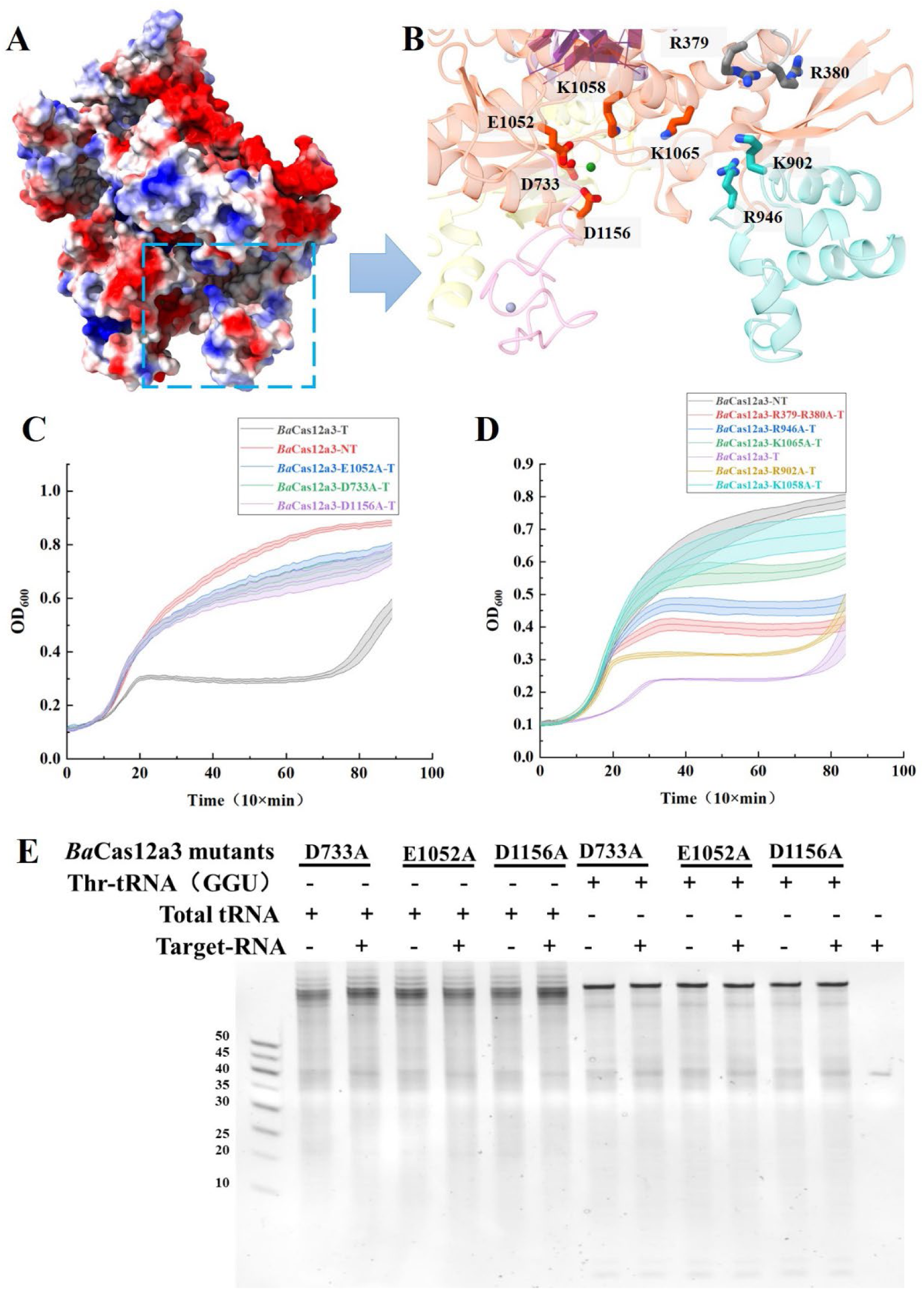
Validation of enzyme activity and immune mechanism by site-directed mutagenesis. (A), Surface electrostatic potential of ternary complex. (B), Key residues within the groove formed by the NL domain and its adjacent REC2, RuvC and ZR domains. Domains were colored as in Fig. 5. Key residues are shown as sticks. (C), Growth curves of RuvC catalytic mutants (D733A, E1052A, and D1156A). (D), Growth curves of substrate binding groove mutants (R379A-R380A, K902A, R946A, K1058A, and K1065A). (E), *In vitro* tRNA collateral cleavage assay of the RuvC mutants.

To further elucidate the functional role of the positively charged groove observed in our structural model, we conducted a mutational analysis. This distinctive channel is formed by the NL domain and adjacent regions of the REC2, RuvC, and ZR domains, creating an interface well-suited for tRNA binding (Fig. 8A). We selected 6 solvent-exposed positively charged residues (R379, R380, K902, R946, K1058, and K1065) lining this groove—which we suspected to be involved in substrate recognition—and substituted each with alanine. Correspondingly, 5 mutant pCas12a3 plasmids were generated (pCas12a3^R379-R380A^, pCas12a3^R946A^, pCas12a3^K902A^, pCas12a3^K1065A^, and pCas12a3^K1058A^) (Table S1). The functional impact of these mutations was assessed using a growth curve assay, which measures immune-mediated growth inhibition upon activation by IPTG and arabinose induction. Strikingly, all 6 alanine mutants exhibited a significant growth inhibition compared to cells expressing the wild-type protein (Fig. 8D). The degree of functional impairment varied among the mutants, suggesting that substrate binding is likely a multivalent process involving several weak interactions. This characteristic is consistent with the ability of *Ba*Cas12a3 to cleave multiple tRNAs with diverse stem-loop structures (Fig. 4C and Fig. S6), as a multivalent binding interface could accommodate such structural variation.

## Discussion

The discovery of Cas12a3 presented in this study expands the functional diversity of Type V CRISPR-Cas systems. While most Cas12 effectors target DNA, the emergence of RNA-targeting subtypes like Cas12a2 and now Cas12a3 underscores the remarkable evolutionary adaptability of the RuvC catalytic core (8, 9, 20). Our data establish Cas12a3 as a phylogenetically and mechanistically distinct entity that induces bacterial growth arrest not through broad collateral nucleic acid cleavage, but via a precise, tRNA collateral activity.

The tRNA cleavage activity of Cas12a3 shares conceptual parallels with a broader biological strategy observed in certain bacterial toxin-antitoxin systems and eukaryotic stress responses, suggesting a convergent evolutionary strategy to regulate cellular processes (60–65). For instance, bacterial defense systems like PARIS deploy nuclease AriB to cleave specific tRNAs within their anticodon stem-loops upon phage detection, halting translation to abort infection (61, 66). Phages counter by encoding immune-resistant tRNAs, driving a molecular arms race (61, 66). While enzymes like Angiogenin primarily target the anticodon loop to generate regulatory tRNA-derived fragments, cleavage at the conserved 3′-CCA terminus provides another layer of control (64, 67). The CCA-end cleavage inactivates tRNAs by preventing aminoacylation, ensuring a robust and sustained blockade of translation (64). Thus, tRNA cleavage serves as a fundamental host regulation strategy, utilizing both precise anticodon scission and terminal CCA inactivation to inhibit translation. The convergent evolution of tRNA-cleavage in CRISPR-Cas systems underscores the potency of disrupting translation as an antiviral strategy. It highlights how different CRISPR lineages have independently evolved effectors that, despite different ancestral structures (RuvC nuclease vs. HEPN nuclease), have been selected to disrupt protein synthesis via tRNA damage (68). Cas13a from *Leptotrichia shahii* cleaves tRNAs within the anticodon loop, particularly those with uridine-rich anticodons, leading to translation inhibition (68). In contrast, *Ba*Cas12a3 targets the conserved 3′-CCA aminoacylation site. This difference in cleavage site suggests distinct modes of translational arrest: Cas13a may impair codon recognition, while Cas12a3 directly prevents tRNA aminoacylation and entry into the ribosomal cycle (68). The depletion of functional tRNA pools by either mechanism would globally impair translation, providing a potent means to halt viral replication without the potentially mutagenic DNA damage.

The structural basis for *Ba*Cas12a3’s unique function is explained by its cryo-EM structures. The presence of the novel NL domain is a defining characteristic (Fig. 6). In the binary complex, the NL domain partially blocked the RuvC active site and contributes to a narrow groove. Target RNA recognition and PFS verification trigger large-scale conformational changes, including a retraction of the NL domain. This opens the groove and expose the catalytic triad, likely forming a dedicated binding channel for the tRNA substrate (Fig. 7). The cooperative action of the NL domain and the ZR domain appears to guide the tRNA 3′ end into the RuvC active site for cleavage. This “gated channel” mechanism explains the enzyme’s specificity: the architecture may sterically or electrostatically favor the structural features of tRNA over other nucleic acids. This model contrasts with Cas12a2, whose more exposed active site in the ternary complex allows promiscuous substrate access. Cas13 enzymes undergo a different activation, reorienting HEPN domains to create a nonspecific RNase site that subsequently selects tRNA anticodon loops (21, 69, 70). While both *Ba*Cas12a3 and *Ba*Cas12a3 activation opens a pre-formed, specific channel—governed by the NL domain—that directly positions the tRNA 3′ end into the repurposed RuvC site. More importantly, the detailed structural basis of tRNA recognition and cleavage need elucidation. Capturing a quaternary complex of Cas12a3 with a tRNA substrate remains a key challenge.

From a biotechnology perspective, *Ba*Cas12a3 possesses a unique combination of properties. Its RNA-triggered activation and highly specific tRNA collateral activity could enable novel diagnostic applications, such as specific RNA detection assays with a tRNA cleavage readout, potentially offering in sensitivity or signal amplification. Furthermore, its ability to induce a potent, DNA-damage-free growth arrest may be useful for safe human gene therapy or for antimicrobial strategies that selectively trigger pathogenic bacteria inhibition upon detection of specific RNA signatures.

In conclusion, we have identified and characterized Cas12a3, a novel RNA-targeting subtype within the Type V CRISPR-Cas12 family. This system demonstrates functional diversification by employing a unique tRNA-directed collateral nuclease activity to mediate bacterial growth arrest without causing DNA damage. Its distinct mechanism, governed by the novel NL domain that forms a gated substrate channel, provides new insights into the evolution of CRISPR immunity. The functional convergence with Cas13a on tRNA cleavage, despite distinct evolutionary origins and molecular architectures, underscores the critical role of disabling translation as a robust antiviral strategy in prokaryotes. Cas12a3 thus offers a novel and valuable scaffold for developing RNA-responsive biotechnological tools.

During the preparation of this manuscript, the article “RNA-triggered Cas12a3 cleaves tRNA tails to execute bacterial immunity” has been published. Dmytrenko, O., Yuan, B., Crosby, K.T. et al. (2026) RNA-triggered Cas12a3 cleaves tRNA tails to execute bacterial immunity. *Nature*. https://doi.org/10.1038/s41586-025-09852-9.

## Data availability

Sequencing data from PFS depletion assays, crRNA sequencing, and post-interference transcriptomics are available under NCBI GEO accession codes PRJNA1400466, PRJNA1400313, and PRJNA1400230, respectively. The atomic models of *Ba*Cas12a3 binary and tertiary complexes have been deposited into the PDB with accession codes 21WE and 21WJ, and the corresponding maps have been deposited into the Electron Microscopy Data Bank with codes EMD-68046 and EMD-68050, respectively.

## Acknowledgments

The authors thank Emeritus Professor David W. Rice (Sheffield University) and Professor John van der Oost (Wageningen University) for their valuable insights during the mechanistic investigation of Cas12a3. We also thank Xiaoju Li, Haiyan Sui, and Xueyun Geng from Shandong University Core Facilities for Life and Environmental Sciences for their help with the Cryo-EM. Molecular graphics and analyses performed with UCSF ChimeraX, developed by the Resource for Biocomputing, Visualization, and Informatics at the University of California, San Francisco, with support from National Institutes of Health R01-GM129325 and the Office of Cyber Infrastructure and Computational Biology, National Institute of Allergy and Infectious Diseases.

