## Supplementary material for "*Ba*Cas12a3 represents a new subtype of type V CRISPR effector with collateral activity toward tRNA": Figs. S1-S9

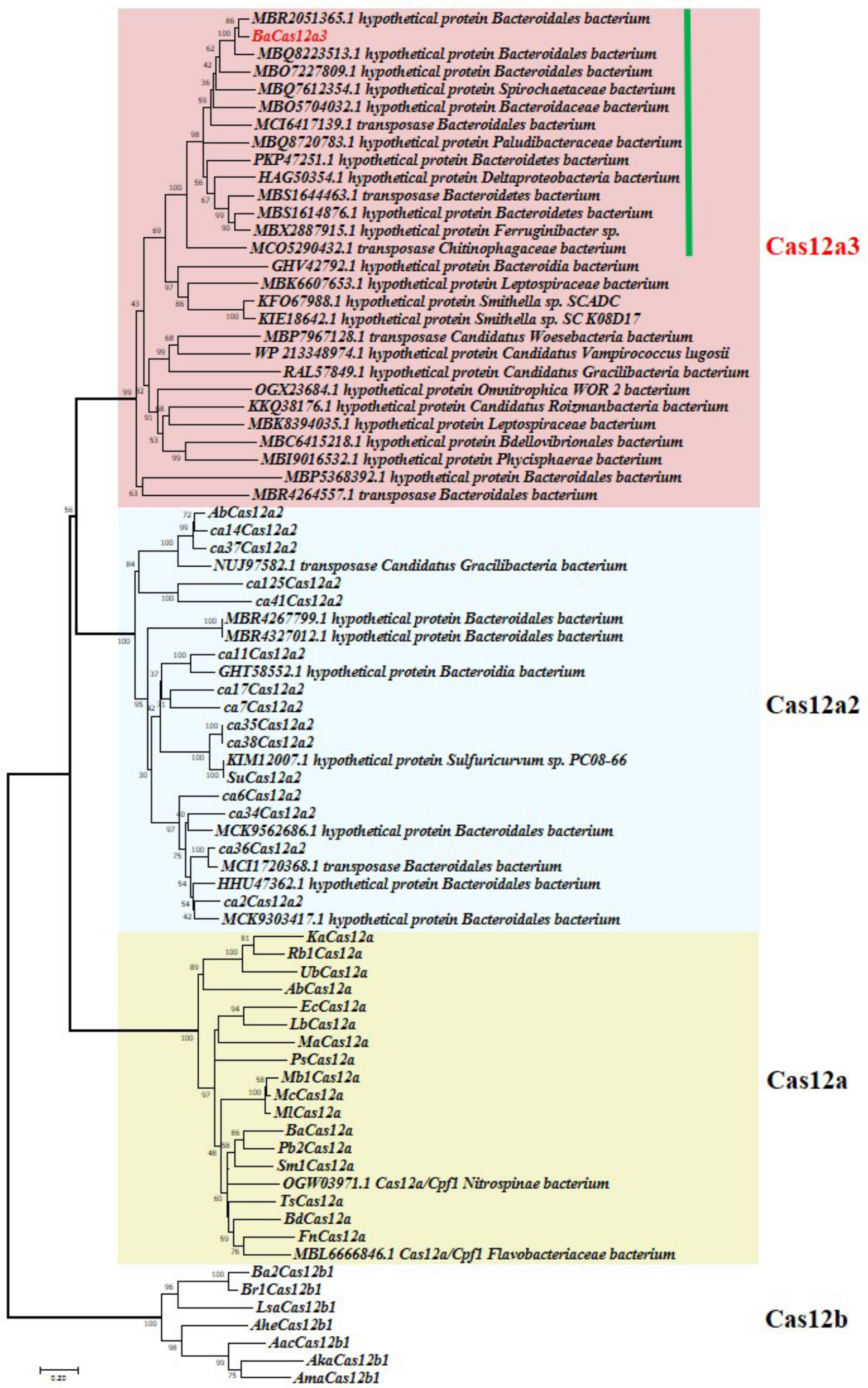


**Extended Data Figure S1.** Phylogenetic analysis of Cas12a3, Cas12a2, Cas12a, and Cas12b nucleases. A neighbor-joining phylogenetic tree was constructed using Cas12b as an outgroup. Clades are shaded to designate Cas12a3 (red), Cas12a2 (blue), Cas12a (yellow), and Cas12b orthologs. Sequences without an NCBI accession number were obtained from a prior Cas12a2 study (1). The green line highlights members that contain an additional domain of unknown function. The evolutionary history was inferred using the Neighbor-Joining method. The percentage of replicate trees (1000 bootstrap replicates) in which associated taxa clustered together is shown next to branches (2, 3). The tree is drawn to scale, with branch lengths proportional to evolutionary distances, which were computed using the Poisson correction method and are expressed as the number of amino acid substitutions per site (4). All positions containing gaps or missing data were eliminated. Evolutionary analyses were conducted in MEGA7 (5).

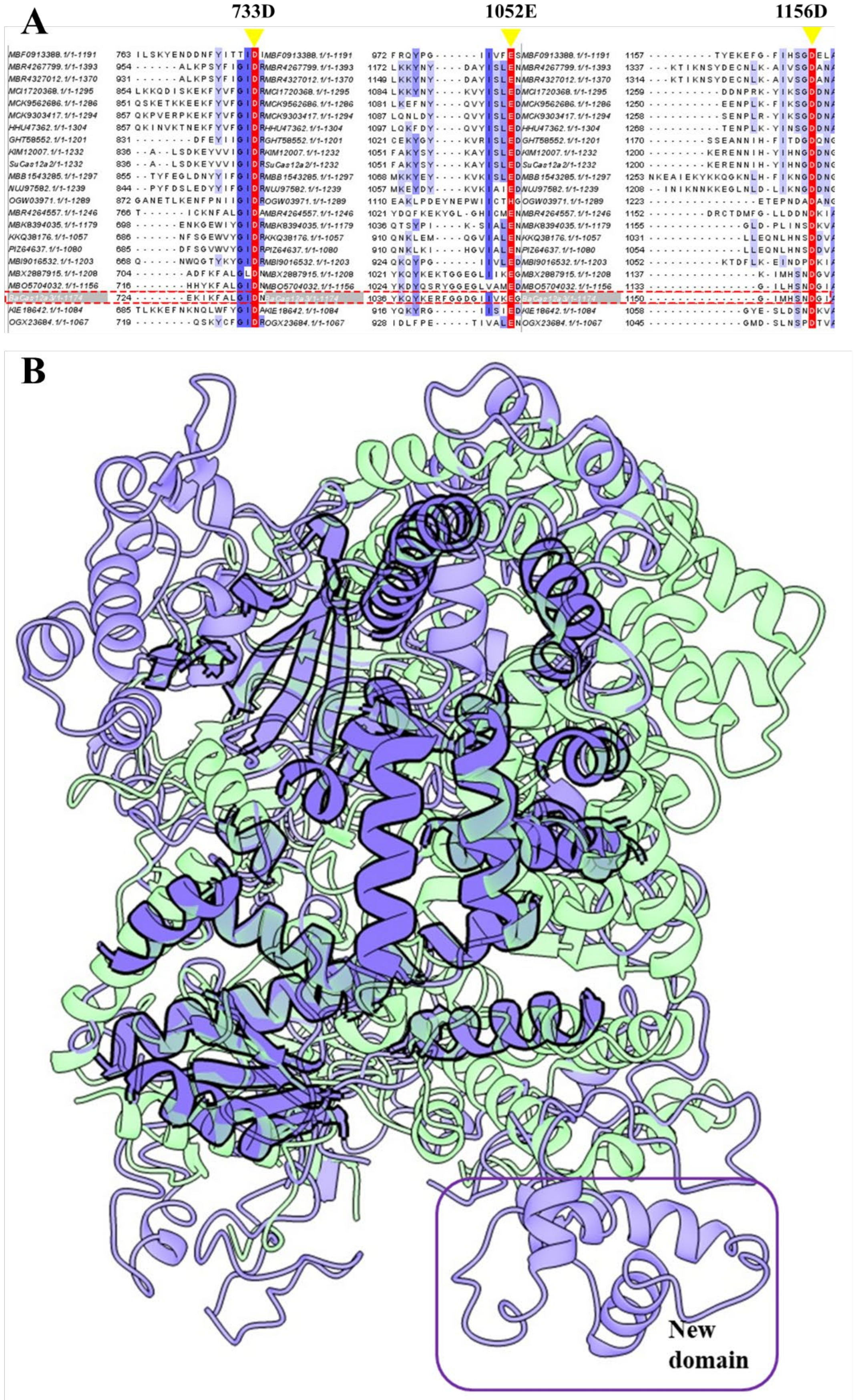


**Extended Data Figure S2.** Sequence and predicted structure alignments of *Ba*Cas12a3

and *Su*Cas12a2. (A), Sequence alignment of the RuvC nuclease domains. The 3 key catalytic residues are highlighted. (B), Structural comparison of the binary complexes *Ba*Cas12a3–crRNA and *Su*Cas12a2–crRNA (PDB: 8D49). *Ba*Cas12a3 is shown in purple and SuCas12a2 in green (cartoon representation). Regions of structural alignment are highlighted. The RMSD is 1.2 Å over 141 pruned atom pairs (compared to 18.3 Å over 789 pairs prior to pruning). A distinct domain present in *Ba*Cas12a3 is indicated within the box.

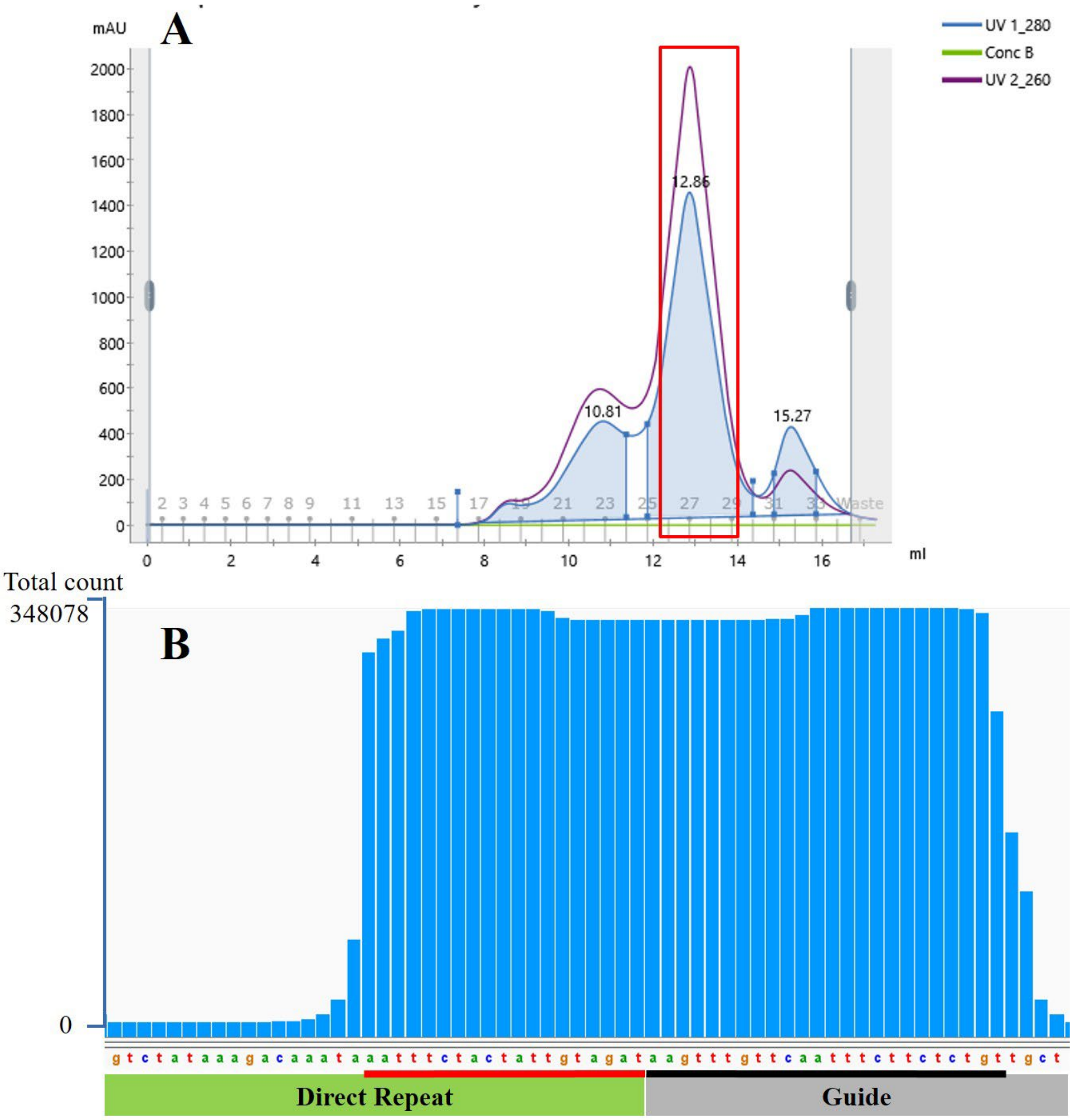


**Extended Data Figure S3.** Purification of the binary complex and small RNA sequencing of the crRNA. (A), Purification profile of the BaCas12a3-crRNA binary complex. The elevated $A_{260}$ relative to $A_{280}$ (highlighted in the red box) confirms the presence of bound nucleic acid (crRNA) in the complex. (B), Sequence of the crRNA extracted from the purified complex. The repeat region is underlined in red, and the guide region is underlined in black.

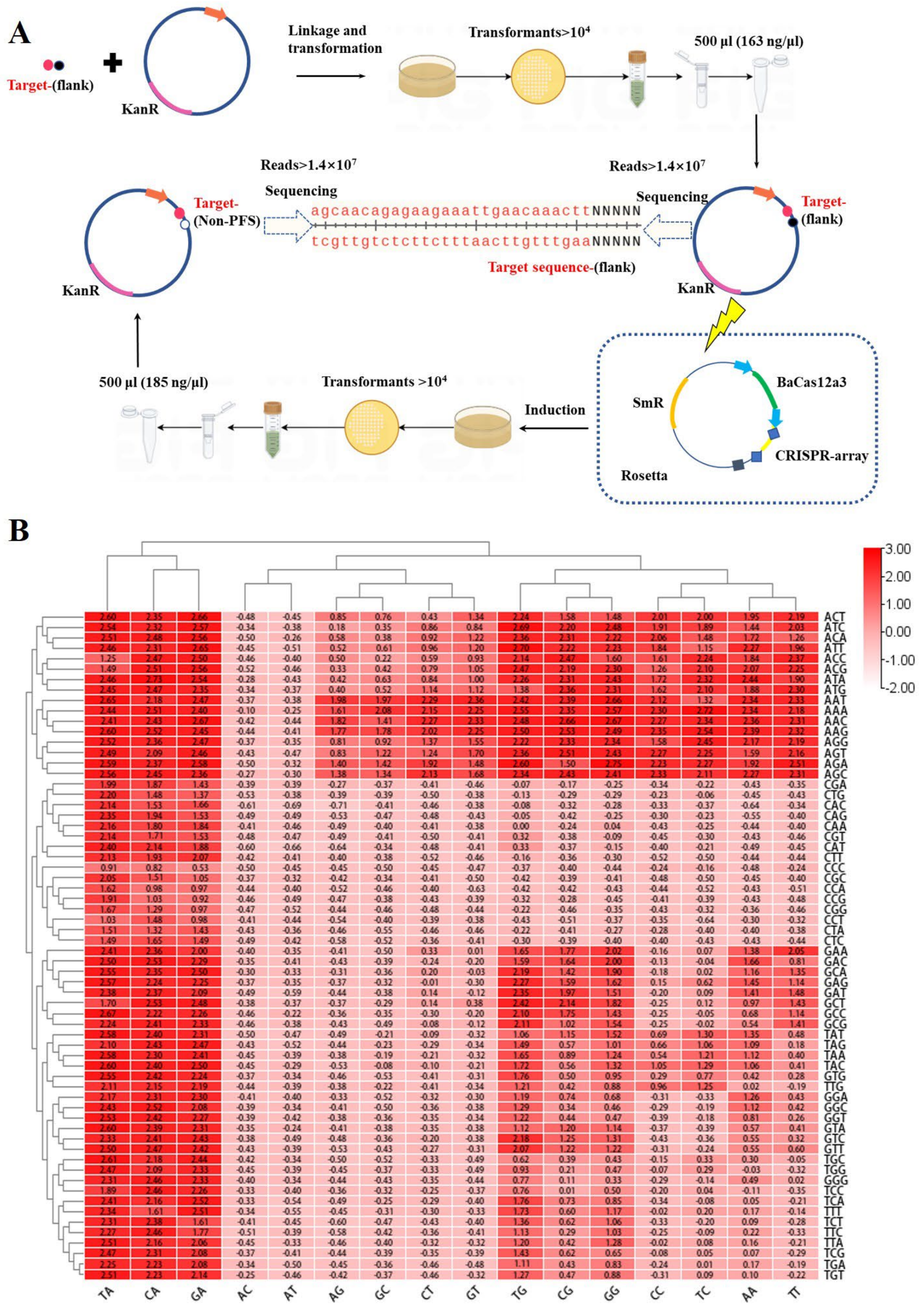


**Extended Data Figure S4.** PFS depletion screen for *Ba*Cas12a3. (A), Schematic of the PFS depletion assay. A plasmid library containing a target sequence flanked by a fully randomized 5-nt region (5′-NNNNN-3′) was transformed into *E.coli* expressing the *Ba*Cas12a3 effector complex. Plasmids bearing a functional PFS are depleted via

CRISPR interference. The PFS region was deep-sequenced from both the initial library and the surviving "escape" library for comparative analysis. (B), Heatmap of PFS depletion values. The heatmap displays the $log_2$-transformed depletion ratio (escape vs. initial library) for all 1024 possible 5-nt sequences. Red indicates positive values (depleted sequences, corresponding to putative functional PFS motifs), whereas light red indicates negative values (enriched or neutral sequences, corresponding to non-functional PFS).

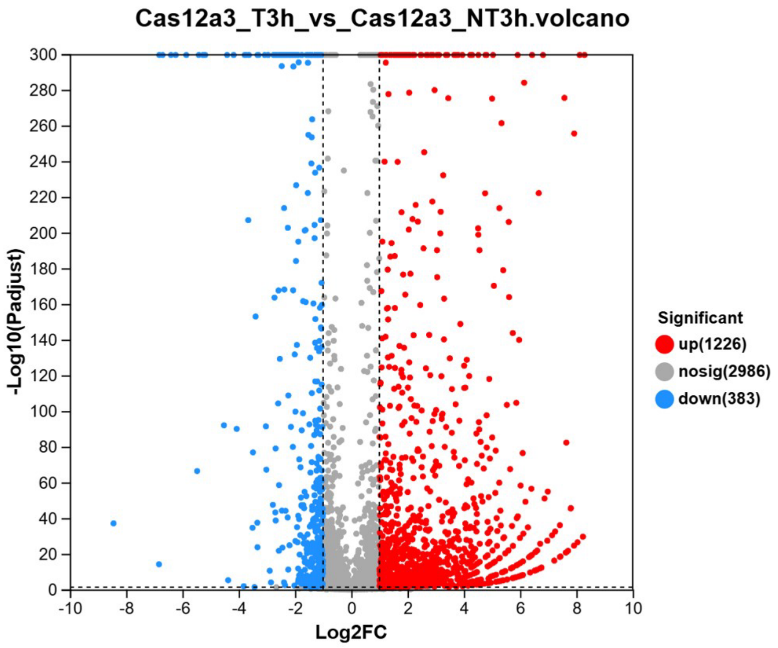


| Gene_id | Gene name | Gene description | FC | Log2FC | Pvalue | Padjust | Significant | Regulate |
|---|---|---|---|---|---|---|---|---|
| HO396_RS19140 | HO396_RS19140 | tRNA-Ala | 313.654 | 8.293028019 | 0 | 0 | yes | up |
| HO396_RS15970 | HO396_RS15970 | tRNA-Ala | 301.035 | 8.233786413 | 4.45E-31 | 2.44E-30 | yes | up |
| HO396_RS01310 | ecpR | EcpR | 278.079 | 8.119353018 | 0 | 0 | yes | up |
| HO396_RS01020 | HO396_RS01020 | tRNA-Ala | 242.732 | 7.92322259 | 1.16E-257 | 3.09E-256 | yes | up |

**Extended Data Figure S5.** Transcription analysis of the targeting and non-targeting strains after 3h induction. The upper panel is volcano plot of differential expression. The x axis represents the fold change (FC) in gene expression between the two sample groups. The x axis represents the statistical test value of the difference in gene expression levels, i.e., the *p*-value. The higher the *p*-value, the more significant the expression difference. Both the horizontal and y axis values have been log-transformed. Each point in the figure represents a specific gene: red points indicate significantly upregulated genes, green points indicate significantly downregulated genes, and gray points represent non-significant differential genes. The lower panel shows the 4 genes exhibiting the greatest upregulation in expression levels.

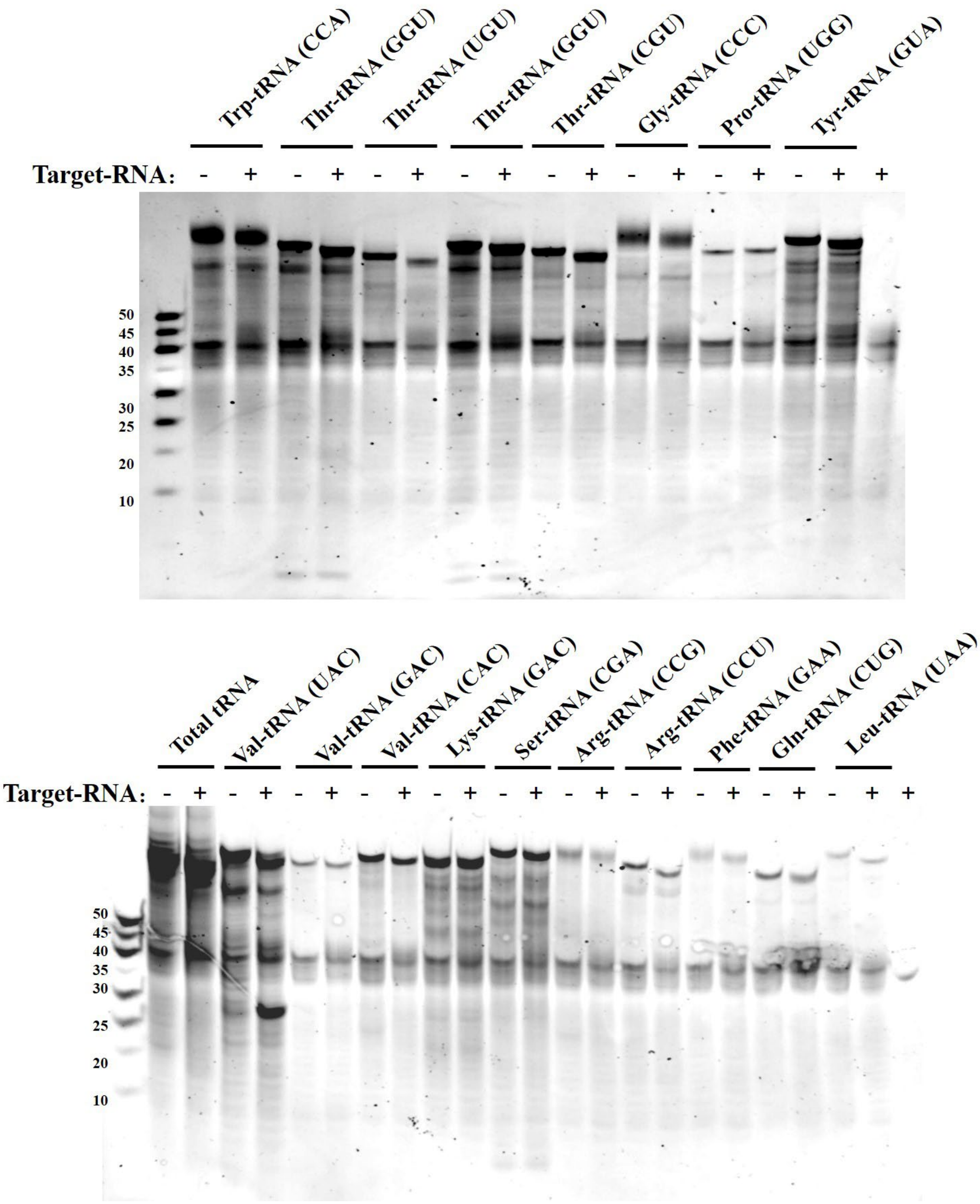


**Extended Data Figure S6.** Various tRNA cleavage assay by *Ba*Cas12a3. *Ba*Cas12a3-crRNA was added in every lane except the markers, and target RNA was added to active the collateral cleavage of tRNAs.

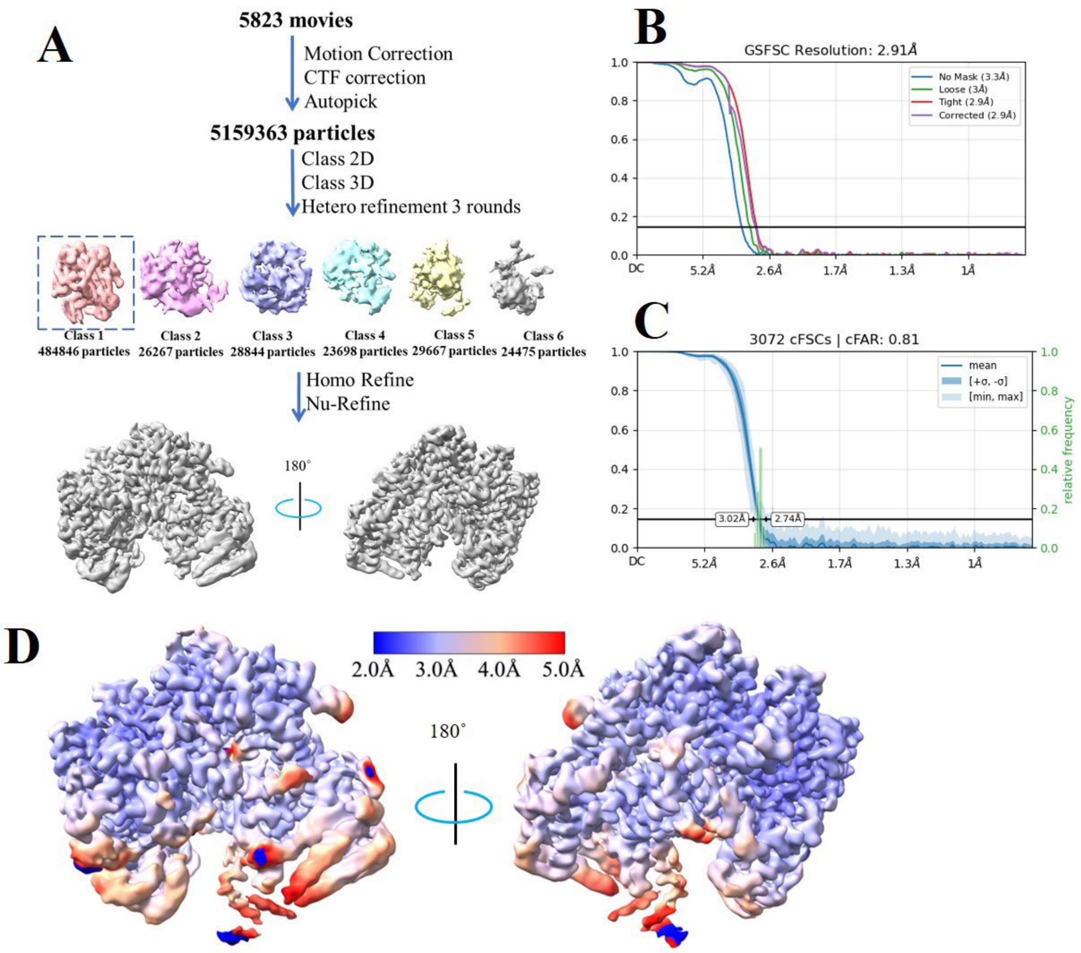


**Extended Data Figure S7.** Cryo-EM analysis of the binary complex, related to Figure 5. (A), Single-particle cryo-EM image processing workflow. (B), Gold Standard Fourier Shell Correlation (GSFSC) plot for the final refinement iteration. (C), The cFSC plot using the auto-tightened mask with a conical FSC Area Ratio (cFAR) of 0.81. (D), Local resolution of the density map with two views.

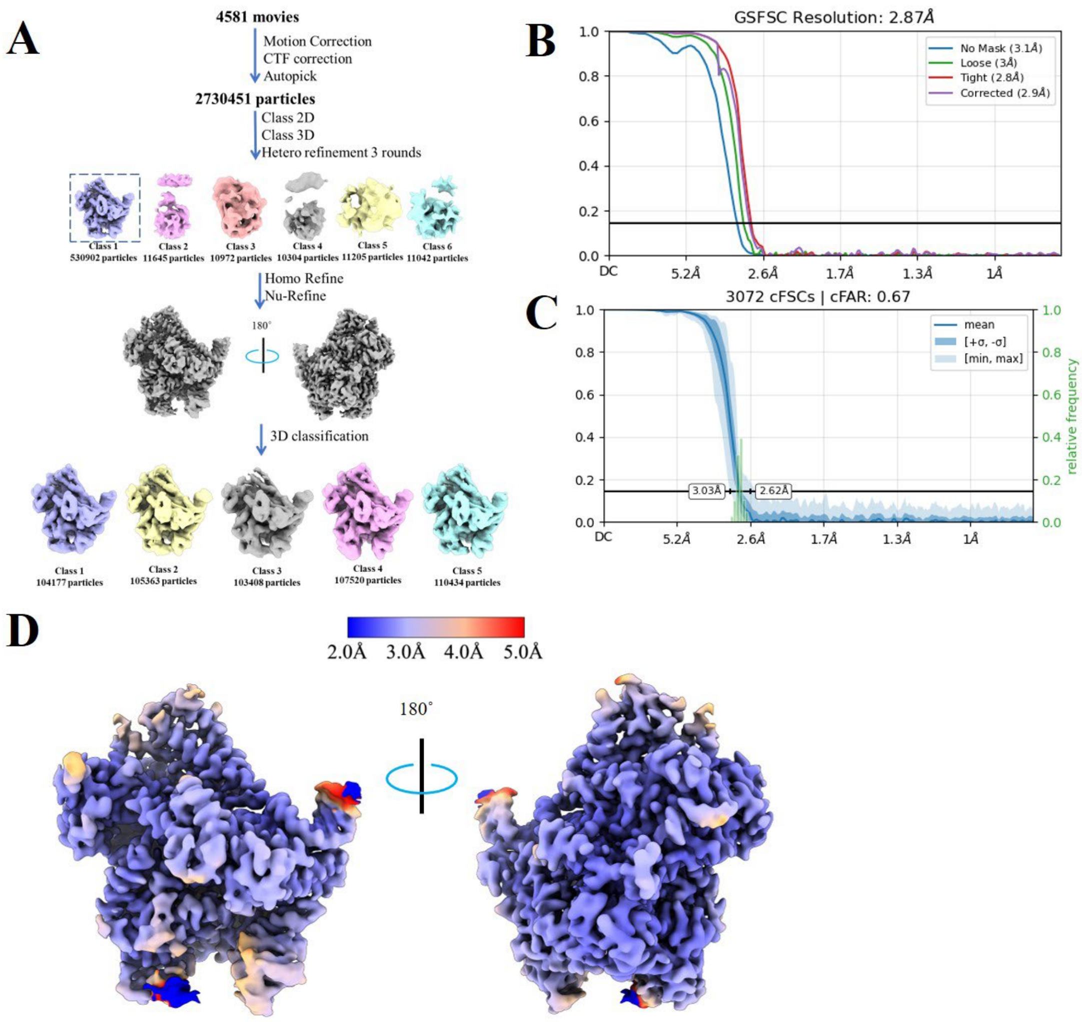


**Extended Data Figure S8.** Cryo-EM analysis of the ternary complex, related to Figure 5. (A), Single-particle cryo-EM image processing workflow. (B), Gold Standard Fourier Shell Correlation (GSFSC) plot for the final refinement iteration. (C), The cFSC plot using the auto-tightened mask with a conical FSC Area Ratio (cFAR) of 0.67. (D), Local resolution of the density map with two views.

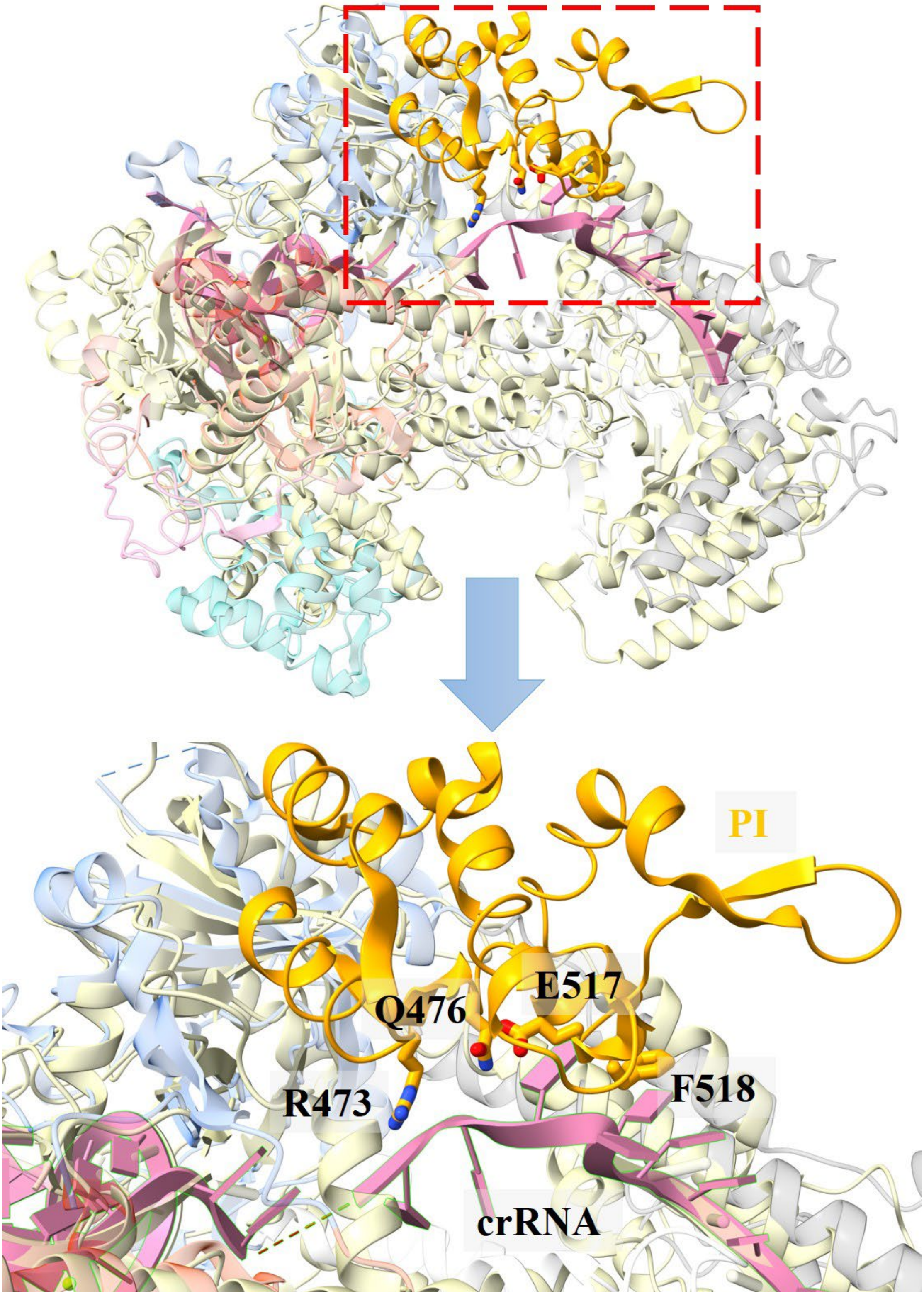


**Extended Data Figure S9.** Structural superposition of the *Ba*Cas12a3 and Cas12a2 binary complexes. The structure of Cas12a2 (PDB: 8D49) is shown for comparison. (Upper panel) Overall view of the superimposed complexes. (Lower panel) Close-up view of the region indicated by the red rectangle in the upper panel. *Ba*Cas12a3 domains are colored as in Fig. 5, and Cas12a2 is shown as a light green cartoon. The crRNA of BaCas12a3 is depicted in pink (cartoon and sticks). Key residues from the PI domain that interact with the crRNA are displayed as sticks.
