## Supplementary material for "*Ba*Cas12a3 represents a new subtype of type V CRISPR effector with collateral activity toward tRNA": Tables S1-S4

**Extended Data Table S1. The strains and plasmids used in this study.**

| Strain/Plasmid | Feature | Source |
|---|---|---|
| *E.coli* DH5*α* | Cloning strain | Lab stock |
| *E.coli* BL21 | Expression strain | Lab stock |
| pCDFDuet-1 | This plasmid contains two T7 promoter-lac operator-MCS (multiple cloning site) cassettes and carries a streptomycin resistance gene. | Lab stock |
| pBAD-G | The plasmid carries an araBAD promoter-MCS-gfp (green fluorescent protein gene) expression cassette and confers kanamycin resistance. | Lab stock |
| pCDFDuet-*Ba*Cas12a3 | Based on the pCDFDuet-1 vector, this construct enables IPTG-inducible co-expression in *E. coli* of *Ba*Cas12a3 protein with a C-terminal 6×His tag | This study |
| pCas12a3 | Based on the pCDFDuet-*Ba*Cas12a3 vector, this construct enables IPTG-inducible co-expression in *E. coli* of *Ba*Cas12a3 protein with a C-terminal 6×His tag and transcription of pre-crRNA. | |
| pCas12a3$^{D733A}$ | This construct was generated from the pCDFDuet-1 vector to enable IPTG-inducible co-expression of the *Ba*Cas12a3D733A protein mutant and transcription of pre-crRNA in *E. coli*. | This study |
| pCas12a3$^{E1052A}$ | This construct was generated from the pCDFDuet-1 vector to enable IPTG-inducible co-expression of the *Ba*Cas12a3E1052A protein mutant and transcription of pre-crRNA in *E. coli*. | This study |
| pCas12a3$^{D1156A}$ | This construct was generated from the pCDFDuet-1 vector to enable IPTG-inducible co-expression of the *Ba*Cas12a3D1156A protein mutant and transcription of pre-crRNA in *E. coli*. | This study |
| pCas12a3$^{R379\text{-}R380A}$ | This construct was generated from the pCDFDuet-1 vector to enable IPTG-inducible co-expression of the *Ba*Cas12a3R379-R380A protein mutant and transcription of pre-crRNA in *E. coli*. | This study |
| pCas12a3$^{R946A}$ | This construct was generated from the pCDFDuet-1 vector to enable IPTG-inducible co-expression of the *Ba*Cas12a3R946A protein mutant and transcription of pre-crRNA in *E. coli*. | This study |
| pCas12a3$^{K902A}$ | This construct was generated from the pCDFDuet-1 vector to enable IPTG-inducible co-expression of the *Ba*Cas12a3K902A protein mutant and transcription of | This study |

| | | |
|---|---|---|
| | pre-crRNA in *E. coli*. | |
| pCas12a3$^{K1065A}$ | This construct was generated from the pCDFDuet-1 vector to enable IPTG-inducible co-expression of the *Ba*Cas12a3K1065A protein mutant and transcription of pre-crRNA in *E. coli*. | This study |
| pCas12a3$^{K1058A}$ | This construct was generated from the pCDFDuet-1 vector to enable IPTG-inducible co-expression of the *Ba*Cas12a3K1058A protein mutant and transcription of pre-crRNA in *E. coli*. | This study |
| pTarget | This construct, based on the pBAD vector, contains a protospacer sequence with a 5'-CAAAG-3' protospacer-flanking site (PFS) and allows for L-arabinose-inducible transcription of the protospacer transcript. | This study |
| pNTarget | Based on the pBAD-Target backbone, this control construct lacks a protospacer sequence. | This study |
| pTNoPr | Based on the pBAD-Target backbone, this control construct lacks the L-arabinose inducible promoter, thereby abolishing transcription of the protospacer sequence. | This study |
| pTNoPfs | This construct was generated from the pBAD-Target backbone by substituting the protospacer-flanking site (PFS) sequence from CAAAG to CACGA. | This study |
| pTarget-(flank) | For unbiased screening of PFS preferences, a plasmid library was constructed based on the pBAD-Target backbone, in which the defined PFS (CAAAG) was replaced with a fully randomized sequence (NNNNN). | This study |

**Extended Data Table S2. Primers used in this study.**

| Primer | Sequence (5'-3') |
|---|---|
| *Ba*Cas12a3-C-His-tag-F | GTTTAACTTTAATAAGGAGATATACCATGGATTTTATTAA GACAACCAAAGC |
| *Ba*Cas12a3-C-His-tag-R | ATTATGCGGCCGTGTACAATATCAGTGGTGGTGGTGGTG GTGCTCGAGTGC |
| CRISPR array-F | CGGATAACAATTCCCCATCTTAGTAACAATTCCCCTCTAG AG TCT |
| CRISPR array-R | TTAAGCTGCGCTAGTAGACGAGTGGTGCTCGAGATCTAC |
| *Ba*Cas12a3D733A-SOE-F | TTCGCTTTGGGTATCGCAAATGGCGAGGTGGAACTTT |
| *Ba*Cas12a3D733A-SOE-R | CACCTCGCCATTTGCGATACCCAAAGCGAACT |
| *Ba*Cas12a3E1051A-SOE-F | GGCATTATCGTAAAAGCTGGTTTCGACAGTGCTAAGG |
| *Ba*Cas12a3E1051A-SOE-R | ACTGTCGAAACCAGCTTTTACGATAATGCCATCGC |
| *Ba*Cas12a3D1156A-SOE-F | AAGGGAATTATGCATTCCAACGCTGGCATTGCTGGTTAC AAT |

| | |
|---|---|
| *Ba*Cas12a3D1156A-SOE-R | ACCTCTCTTAGCTATATTGTAACCAGCAATGCCAGCGTTGGA ATGCATAATTCCC |
| *Ba*Cas12a3R379-380A-SOE-R | GCCGCAGATGCTGCTACGAAGGAAAAATTCAAGACATAT |
| *Ba*Cas12a3R379-380A-SOE-F | TTCGTAGCAGCATCTGCGGCATTTATGAGAGAA |
| *Ba*Cas12a3K902A-SOE-R | CGACTATCTAAATGGTGCTAATAAAAAATACGAAGATTTGTCTG |
| *Ba*Cas12a3K902A-SOE-F | CTTCGTATTTTTTATTAGCACCATTTAGATAGTCGATGAACT |
| *Ba*Cas12a3R946A-SOE-R | AAGTAAAGGTCAAGCTGTAGGAGATTACTTGAATAATGTTCTTG |
| *Ba*Cas12a3R946A-SOE-F | CAAGTAATCTCCTACAGCTTGACCTTTACTTATTTCGGC |
| *Ba*Cas12a3K1058A-SOE-R | CGACAGTGCTGCTGTTGAAAGCGATAGAGAG |
| *Ba*Cas12a3K1058A-SOE-F | GCTTTCAACAGCAGCACTGTCGAAACCTTC |
| *Ba*Cas12a3K1065A-SOE-R | GAAAGCGATAGAGAGGCTTTCTCAGGTAACATATACAG |
| *Ba*Cas12a3K1065A-SOE-F | TGAGAAAGCCTCTCTATCGCTTTCAACTTTAGC |
| Target-F | CATGGAGCAACAGAGAAGAAATTGAACAAACTTCAAAGGAA |
| Target-R | AGCTTTCCTTTGAAGTTTGTTCAATTTCTTCTCTGTTGCTC |
| pBad-Nonpromoter-F | CGGTACCTGATAACCGGCGCTACGATAC |
| pBad-Nonpromoter-R | CATGGTATCGTAGCGCCGGTTATCAGGTACCGAGCT |
| Target-(flank)-F | CATGGAGCAACAGAGAAGAAATTGAACAAACTTNNNNNGA A |
| Target-(flank)-R | AGCTTTCNNNNNAAGTTTGTTCAATTTCTTCTCTGTTGCTC |
| NonTarget-R | AGCTTAGATACCCTGGAGGGAAACCAGACTTCATGTGGTCTT TTCCTCCTTGC |
| NonTarget-F | CATGGCAAGGAGGAAAAGACCACATGAAGTCTGGTTTCCCT CCAGGGTATCTA |
| NonPFS-Target-F | AATTGAACAAACTTAGCACGAAAGCTTTGAAAAAAAAATATTTGGAG |
| NonPFS-Target-R | AGCTTTCGTGCTAAGTTTGTTCAATTTCTTCTCTGTTGC |
| tRNA-Arg（CCG)-F | TAATACGACTCACTATAGGCGCCCGTAGCTCAGCTGGATAGAGCGCTGCCCT |
| tRNA-Arg（CCG)-R | TGGCGCGCCCGACAGGATTCGAACCTGAGACCTCTGCCTCCGGAGGGCAGCGCTCTATC |
| tRNA-Arg(CCU)-F | TAATACGACTCACTATAGGTCCTCTTAGTTAAATGGATATAACGAGCCCCTCC |
| tRNA-Arg(CCU)-R | TGGTGTCCCCTGCAGGAATCGAACCTGCAATTAGCCCTTAGGAGGGGCTCGTTATATCC |
| tRNA-Gln(CUG)-F | TAATACGACTCACTATAGTGGGGTATCGCCAAGCGGTAAGGCACCGGATTCTG |
| tRNA-Gln(CUG)-R | TGGCTGGGGTACGAGGATTCGAACCTCGGAATGCCGGA |

| | |
|---|---|
| | ATCAGAATCCGGTGCCTTACC |
| tRNA-Gly(CCC)-F | TAATACGACTCACTATAGGCGGGCGTAGTTCAATGGTAGAACGAGAGCTTCCCA |
| tRNA-Gly(CCC)-R | TGGAGCGGGCGAAGGGAATCGAACCCTCGTATAGAGCTTGGGAAGCTCTCGTTCTACCA |
| tRNA-Leu(UAA)-F | TAATACGACTCACTATAGGCCCGGATGGTGGAATCGGTAGACACAAGGGATTTAAAATC |
| tRNA-Leu(UAA)-R | TGGTACCCGGAGCGGGACTTGAACCCGCACAGCGCGAACGCCGAGGGATTTTAAATCCCTTGTGTCTAC |
| tRNA-Lys(UUU)-F | TAATACGACTCACTATAGGGGTCGTTAGCTCAGTTGGTAGAGCAGTTGACTTTTAATC |
| tRNA-Lys(UUU)-R | TGGTGGGTCGTGCAGGATTCGAACCTGCGACCAATTGATTAAAAGTCAACTGCTCTACC |
| tRNA-Phe(GAA)-F | TAATACGACTCACTATAGGCCCGGATAGCTCAGTCGGTAGAGCAGGGGATTGAAA |
| tRNA-Phe(GAA)-R | TGGTGCCCGGACTCGGAATCGAACCAAGGACACGGGGATTTTCAATCCCCTGCTCTACC |
| tRNA-Pro(UGG)-F | TAATACGACTCACTATAGCGGCGAGTAGCGCAGCTTGGTAGCGCAACTGGTTTGG |
| tRNA-Pro(UGG)-R | TGGTCGGCGAGAGAGGATTCGAACCTCCGACCCACTGGTCCCAAACCAGTTGCGCTACC |
| tRNA-Ser(CGA)-F | TAATACGACTCACTATAGGGTGAGGTGTCCGAGTGGCTGAAGGAGCACGCCTGGAAAGT |
| tRNA-Ser(CGA)-R | TGGCGGTGAGGGGGGGATTCGAACCCCCGATACGTTGCCGTATACACACTTTCCAGGCGTGCTCCTTC |
| tRNA-Thr(GGU)-F | TAATACGACTCACTATAGGCTGATATAGCTCAGTTGGTAGAGCGCACCCTTG |
| tRNA-Thr(GGU)-R | TGGTGCTGATAGGCAGATTCGAACTGCCGACCTCACCCTTACCAAGGGTGCGCTCTACC |
| tRNA-Thr(UGU)-F | TAATACGACTCACTATAGGCCGACTTAGCTCAGTAGGTAGAGCAACTGACTTGTAATC |
| tRNA-Thr(UGU)-R | TGGTGCCGACTACCGGAATCGAACTGGTGACCTACTGATTACAAGTCAGTTGCTCTACC |
| tRNA-Thr(GGU)-F | TAATACGACTCACTATAGGCTGATATGGCTCAGTTGGTAGAGCGCACCCTTGG |
| tRNA-Thr(GGU)-R | TGGTGCTGATACCCAGAGTCGAACTGGGGACCTCACCCTTACCAAGGGTGCGCTCTACC |
| tRNA-Thr(CGU)-F | TAATACGACTCACTATAGGCCGATATAGCTCAGTTGGTAGAGCAGCGCATTC |
| tRNA-Thr(CGU)-R | TGGTGCCGATAATAGGAGTCGAACCTACGACCTTCGCATTACGAATGCGCTGCTCTACC |
| tRNA-Trp(CCA)-F | TAATACGACTCACTATAGAGGGGCGTAGTTCAATTGGTAGAGCACCGGTCTCC |
| tRNA-Trp(CCA)-R | TGGCAGGGGCGGAGAGACTCGAACTCCCAACACCCGG |

| | |
|---|---|
| | TTTTGGAGACCGGTGCTCTACC |
| tRNA-Tyr(GUA)-F | TAATACGACTCACTATAGGGTGGGGTTCCCGAGCGGCCA AAGGGAGCAGACTGTAAATCTGCCG |
| tRNA-Tyr(GUA)-R | TGGTGGTGGGGGAAGGATTCGAACCTTCGAAGTCGATG ACGGCAGATTTACAGTCTGCT |
| tRNA-Val(UAC)-F | TAATACGACTCACTATAGGGGTGATTAGCTCAGCTGGGA GAGCACCTCCCTTAC |
| tRNA-Val(UAC)-R | TGGTGGGTGATGACGGGATCGAACCGCCGACCCCCTCC TTGTAAGGGAGGTGCTCTCCC |
| tRNA-Val(GAC)-F | TAATACGACTCACTATAGGCGTCCGTAGCTCAGTTGGTT AGAGCACCACCTTGACA |
| tRNA-Val(GAC)-R | TGGTGCGTCCGAGTGGACTCGAACCACCGACCCCCACC ATGTCAAGGTGGTGCTCTAAC |
| tRNA-Val(GAC)-F | TAATACGACTCACTATAGGCGTTCATAGCTCAGTTGGTTA GAGCACCACCTTGACATG |
| tRNA-Val(GAC)-R | TGGTGCGTTCAATTGGACTCGAACCAACGACCCCCACC ATGTCAAGGTGGTGCTCTAAC |

**Extended Data Table S3. Nucleic acid substrates.**

| Substrate | Sequence (5'-3') | Length (nt) |
|---|---|---|
| Target ssRNA | CAGGCAGCAACAGAGAAGAAAUUGAACAAACUUGA AAGUAA | 41 |
| Target ssDNA | CAGGCAGCAACAGAGAAGAAATTGAACAA ACTTGAAAGTAA | 41 |
| Target dsDNA | CAGGCAGCAACAGAGAAGAAATTGAACAAAC TTGAAAGTAA | 41 |
| | TTACTTTCAAGTTTGTTCAATTTCTTCTCTGTTG CTGCCTG | |
| Collateral ssRNA | ACGACGAACAGUCAUGAAAGUCUUA | 25 |
| Collateral ssDNA | TAAGACTTTCATGACTGTTCGTCGT | 25 |
| Collateral dsDNA | GACGCTGCCGAATTCTTCTGGATTGCTAGG CCTAGCAATCCAGAAGAATTCGGCAGCGTC | 30 |
| Ala-tRNA（UGC） | GGGGCUAUAGCUCAAGCUGGGAGAGCGCCUGCUUU GCACGCAGGAGGUCUGCGGUUCGAUCCCGCAUAGC UCCACCA | 76 |
| tRNA-Arg（CCG） | GCGCCCGUAGCUCAAGCUGGAUAGAGCGCUGCCCUC CGGAGGCAGAGGUCUCAGGUUCGAAUCCUGUCGGG CGCGCCA | 77 |
| tRNA-Arg (CCU) | GUCCUCUUAGUUAAAUGGAUAUAACGAGCCCCUCC UAAGGGCCAAUUGCAGGUUCGAUUCCUGCAGGGGA | 75 |

| | | |
|---|---|---|
| | CACCA | |
| tRNA-Gln (CUG) | UGGGGUAUCGCCAAGCGGUAAGGCACCGGAUUCUG AUUCCGGCAUUCCGAGGUUCGAAUCCUCGUACCCCA GCCA | 75 |
| tRNA-Gly (CCC) | GCGGGCGUAGUUCAAUGGUAGAACGAGAGCUUCCC AAAGCUCUAUACGAGGGUUCGAUUCCCUUCCCCCGC TCCA | 74 |
| tRNA-Leu (UAA) | GCCCGGAGUGGUGGAAUCGGUAGACACAAGGGAUU UAAAAUCCCUCGGGCGUUCGCGCUGUGCGGGUUCA AGUCCCGCCCCGGGUACCA | 87 |
| tRNA-Lys (UUU) | GGGUCGUUAGCUCAAGUUGGUAGAGCAGUUGACUU UUAAUCAAUUGGUCGCAGGUUCGAAUCCUGCACGA CCCACCA | 76 |
| tRNA-Phe (GAA) | GCCCGGAGAGCUCAAGUCGGUAGAGCAGGGGAUUG AAAAUCCCCGUGUCCUUGGUUCGAUUCCGAGUCCG GGCACCA | 76 |
| tRNA-Pro (UGG) | CGGCGAGUAGCGCAAGCUGGUAGCGCAACUGGUUU GGGACCAGUGGGUCGGAGGUUCGAAUCCUCUCUCG CCGACCA | 77 |
| tRNA-Ser (CGA) | GGUGAGGUGUCCGAGUGGCUGAGGAGGCACGCCUG GAAAGUGUGUAUACGGCAAAGUAUCGGGGGUUCGA AUCCCCCCCCUCACCGCCA | 88 |
| tRNA-Thr (GGU) | GCUGAUAUAGCUCAAGUUGGUAGAGCGACCCUUGG UAAGGGUGAGGUCGGCAGUUCGAAUCUGCCUAUCA GCACCA | 76 |
| tRNA-Thr (UGU) | GCCGACUUAGCUCAAGAGGUAGAGCAGCUGACUUG UAAUCAGUAGGUCACCAGUUCGAUUCCGGUAGUCG GCACCA | 76 |
| tRNA-Thr (GGU) | GCUGAUAUGGCUCAAGUUGGUAGAGCGCACCCUUG GUAAGGGUGAGGUCCCCAGUUCGACUCUGGGUAUC AGCACCA | 76 |
| tRNA-Thr (CGU) | GCCGAUAUAGCUCAAGUUGGUAGAGCAGCGCAUUC GUAAUGCGAAGGUCGUAGGUUCGACUCCUAUUAUC CGGCACCA | 76 |
| tRNA-Trp (CCA) | AGGGGCGUAGUUCAAUUGGUAGAGCACCCGUCUCC AAAACCCGGGUGUUGGGAGUUCGAGUCUCUCCGCC CCUGCCA | 76 |
| tRNA-Tyr (GUA) | GGUGGGGUUCCCGAGCGGCCAAAGGGAGCAGACUG UAAAUCUGCCGUCAUCGACUUCGAAGGUUCGAAUC CUUCCCCCCACCACCA | 85 |
| tRNA-Val (UAC) | GGGUGAUUAGCUCAAGCUGGGAGAGCACCUCCCUU ACAAGGAGGGGGUCGGCGGUUCGAUCCCGUCAUCA CCCACCA | 76 |
| tRNA-Val (GAC) | GCGUCCGUAGCUCAAGUUGGUUAGAGCACCACCUU | 77 |

| | | |
|---|---|---|
| | GACAUGGUGGGGGUCGGUGGUUCGAGUCCACUCCG ACGCACCA | |
| tRNA-Val (GAC) | GCGUUCAUAGCUCAAGUUGGUUAGAGCACCACCUU GACAUGGUGGGGGUCGUUGGUUCGAGUCCAAUUGA AUGCACCA | 77 |

**Extended Data Table S4. Cryo-EM data collection and model validation statistics.**

| Data collection and procession | Binary complex (PDB, 21WE, EMD-68046) | Ternary complex (PDB, 21WJ, EMD-68050) |
|---|---|---|
| Voltage (kV) | 200 | 200 |
| Electron exposure($e^{-}$/Å$^2$) | 40 | 40 |
| Defocus range (μm) | | |
| Pixel size (Å) | 0.89 | 0.89 |
| Symmetry imposed initial particle images (no.) | 5159363 | 2730451 |
| Final particle images (no.) | 484846 | 530902 |
| Map resolution (Å) | 2.91 | 2.87 |
| FSC threshold | 0.143 | 0.143 |
| Refinement | | |
| Intial model used | *De novo* | *De novo* |
| Model resolution (Å) | 3.0 | 3.0 |
| FSC threshold | 0.5 | 0.5 |
| Model composition | | |
| Non-hydrogen atoms | 10191 | 11026 |
| Protein residues | 1167 | 1166 |
| Nucleotides | 31 | 69 |
| Ligands | Mg:1 | Zn:1, Mg:2 |
| Mean B factors (Å$^2$) | | |
| Protein | 171.6 | 122.7 |
| Nucleotides | 149.6 | 103.2 |
| Ligand | 110.4 | 152.1 |
| R.m.s.deviations | | |
| Bond lengths (Å) | 0.003 | 0.004 |
| Bond angles (°) | 0.607 | 0.647 |
| Validation | | |
| MolProbity score | 2.17 | 2.31 |
| Clashscore | 9.16 | 9.08 |
| Poor rotamers (%) | 3.07 | 4.27 |
| Ramachandran plot | | |
| Favored (%) | 95.44 | 94.92 |
| Allowed (%%) | 4.56 | 4.99 |

| | | |
|---|---|---|
| Disallowed (%) | 0.00 | 0.09 |
| Map CC (mask) | 0.87 | 0.90 |
