## Supplementary File 1 for "*Ba*Cas12a3 represents a new subtype of type V CRISPR effector with collateral activity toward tRNA"

>AacCas12b1

SMAVKSIKVKLRLDDMPEIRAGLWKLHKEVNAGVRYYTEWLSLLRQENLYRRSPNGDGEQ

ECDKTAEECKAELLERLRARQVENGHRGPAGSDDELLQLARQLYELLVPQAIGAKGDAQQ

IARKFLSPLADKDAVGGLGIAKAGNKPRWVRMREAGEPGWEEEKEKAETRKSADRTADVL

RALADFGLKPLMRVYTDSEMSSVEWKPLRKGQAVRTWDRDMFQQAIERMMSWESWNQRVG

QEYAKLVEQKNRFEQKNFVGQEHLVHLVNQLQQDMKEASPGLESKEQTAHYVTGRALRGS

DKVFEKWGKLAPDAPFDLYDAEIKNVQRRNTRRFGSHDLFAKLAEPEYQALWREDASFLT

RYAVYNSILRKLNHAKMFATFTLPDATAHPIWTRFDKLGGNLHQYTFLFNEFGERRHAIR

FHKLLKVENGVAREVDDVTVPISMSEQLDNLLPRDPNEPIALYFRDYGAEQHFTGEFGGA

KIQCRRDQLAHMHRRRGARDVYLNVSVRVQSQSEARGERRPPYAAVFRLVGDNHRAFVHF

DKLSDYLAEHPDDGKLGSEGLLSGLRVMSVALGLRTSASISVFRVARKDELKPNSKGRVP

FFFPIKGNDNLVAVHERSQLLKLPGETESKDLRAIREERQRTLRQLRTQLAYLRLLVRCG

SEDVGRRERSWAKLIEQPVDAANHMTPDWREAFENELQKLKSLHGICSDKEWMDAVYESV

RRVWRHMGKQVRDWRKDVRSGERPKIRGYAKDVVGGNSIEQIEYLERQYKFLKSWSFFGK

VSGQVIRAEKGSRFAITLREHIDHAKEDRLKKLADRIIMEALGYVYALDERGKGKWVAKY

PPCQLILLAELSEYQFNNDRPPSENNQLMQWSHRGVFQELINQAQVHDLLVGTMYAAFSS

RFDARTGAPGIRCRRVPARCTQEHNPEPFPWWLNKFVVEHTLDACPLRADDLIPTGEGEI

FVSPFSAEEGDFHQIHAALNAAQNLQQRLWSDFDISQIRLRCDWGEVDGELVLIPRLTGK

RTADSYSNKVFYTNTGVTYYERERGKKRRKVFAQEKLSEEEAELLVEADEAREKSVVLMR

DPSGIINRGNWTRQKEFWSMVNQRIEGYLVKQIRSRVPLQDSACENTGDI

>AbCas12a

MKSDFFAPFTNLYPIQKTLRRELKPLNENFQHDPALTALRNSEIPQRDQQREKDYQAIKP

LLDELHNQFITESLQSLETQDRSDFVLFYQTYQKKKKNKADFSEKELKSLDEEFESRTKS

LRTAIGNSYAKLAELRKSNPDYVNEKGKPFLDQKSYKILTEAGVLGLLEKKYASDLEKLA

LIRRFGNFFTYFTGFNQNRENYYTTDEKATAVAYRAINENLLTFANNCELFEKLAVLGLS

ELEKKIFNPDSYSEYLTQSGIDFYNEMLANIRSKANLYTQEHKTKLPQPKLLYKQIGSPR

GNTIPFDLINNDEEFKKMLHTMIAETDQRIPEFNKLLESIFEENADLSQIFLSKTSLNII

SNRYFSSWNTLLEKGVELKLFKFKKNDEESFKLPTYLSLAELKELLEAVPFQTAEKSDTA

EGTHHQVSLLKLQRENLHLESSHSNWELLLKSMKSDFESFWTGEGEFGSYTSAKKALQSL

SSLQSTNQEHKNLIKMLLDNTLYAYRMLKWFKVDTSKLGFVPEGEFYPSLDQLLQDYPLP

KWYDMIRNYLTRKPYSQAKLKLNFDCSTLLNGRDKNKETQNLSVILRKDGKFYLAIMKKD

QNKFFEKSALYEGNLGTMEKMDYKLLPGANKMLPKCLMPGSDKKKYGASDQVLELYAKGS

FKKSEKSFNLADLHTLIDFYKSALPKYEDWKVFNFQFQATENYQDISQFYREVEQQGYLL

NRRKVNEELIKQGIKDGSLFLFQISSKDFEGKSKTPDLQTLYWCQLFESPTNVKLNGEAE

IFFRLGSMKKEKKTLKVDNYDVFKHKRYTEDKILFHVPITLGFGNNEVSASAPSKFNQKL

NQELIIPHFDDLHVIGVDRGEKHLAFYSVVSVKTGKIVKQGTLNLLNGTDYEAKLSQKAE

NRLHARQNRDTIEKIADLKNGYISQVVNKLVELVLEYNAVIVFEDLNAGFKRGRQKIEQS

VYQKLELALAKKLNFIVKKEKAVVEPGSVTSAYQLAPQINTFGDIKGKQRGIMLYTRANY

TSVTDPLTGRRKQYYFKKGSSEEMKAQFFKSFKNLTRDAGQKAYIFDDGTWQLYSNVERR

RGKRGDHRERTQIKYDPSLELDTLFSKYQIEKSDSLLDQLKNRDLPQTFWTSFFRILDLI

MQIRNTDDEGRDIILSPIGNAQERFDSRKRYDQLPRDEKGVVIEESAFEYPTSGDANGAY

NIARKGVMMLERIKENTEKPDLLIRDAEWDKKITN

>AbCas12a2

MVNLESFKNLYEVRKTVRFGLNQPNKKSNINKTHGQLKDLVDLSFEREKKLINNEKNQVL

IDSEKALIEKLQQYVNGLEVQLENWEGTYQRYDLIAINKDYYKILARKAKFDAMWEISKF

DKKSNKYIKVKQPQASQISLSSLKIGNRSDSIIQYWGEIIEKTDYLLNIFKPKLEQYERA

INDANSTHIKPDSIDFRKIFLQLLKLTKEFLQPLLDRSIIFEFSKKKVSQEIEKISEFAG

EKNNTKIYNVLKNGEELRQYFEANGSQVPYGRVSLNYYTAVQKPNNFDQEIKKAIDDLGI

INFLKKRDSQIIDYLHQGSKQKIKLLLTSKSPYSIELLQLFKVKPIPFSVKYNLAKFIEK

NYKNEINLSYEEILDKLNLLGRAIDIANDFKNSNNQNNFSLDEYPVKLAFDYAWENTARS

LKRTIPFPKEVCKQFLKDNFDVDVNVDNADFKLYANLLFIADNLATIEYNNPNNEAELIN

EIKQVFDSIDFSFDKERYGGYKNDVLVLLNKAKPQRDYSTILKAKQELGLLRGGLKNKIK

KYRDLTQRLIDKKDSHFGIASFVGKTLAKIRDRLKEENELNKISHYGVILEDKNQDKYLL

ISQLDGKDTREKISQKFGNGDIKVYQVNSFTSKALNKFIKNPLSEDAKKFHGDFRYKHKE

VSIYDEKGKWTGYQESFLSHIKKCLIDSEISREQNWEAFGWNFAGCNTYEEIEKEVDSKG

YQLTENFISIGNLESLEKDEGCLLFPIINQDISSQKQENKNIFTLDLEKVFEGKECRIHP

EFSIFYRKPMEEHKKENKSGIINRFGRLQLLANLGIEFIPRNSSFKTKKEQNRIAIDQKK

QNQLVQEFNQEKVNTYFEGLDNYYIFGIDRGIKQLATLCITNKNGVIQDFDIYTKHFNSE

SKKWEYKFHRKDGILDLTNLKIESDKSGNKYIVDISLFQAKDEDGNPTGTNKQNIQLKKL

AYIRKLQYQMSANEEGVLSFLEKYKNKEEREQNMKELITPYKEGKNFVDLPMDIFQEMFE

NYYRLKTDQNLSESEKKNLMKITTELDASESLKKGVVANMIGVIYYLMKKYEYKVKISLE

DLSNAWFFSKDGLSGDVVLNTKNDGTMDLKKQDNLALAGVGTYHFFEMQLLKKLFKISTE

EGILHLVPSFGSVKNYTEIFKDKGKYVYKQFGIVYFVDPRNTSKKCPVCLNTQTTGKKEG

IPIINRNYKKSNIFYCERCGFQSIHSHCQEENIKDSNGFHYSVEEVERIEMKNKEAIEKY

KKQGKNLHFIKNGDDNGAYNIGEKIRELPKKSDVKNT

>AheCas12b1

MTVRSIRVKLAVGSPQYRDVRRGLWKTHEIMNQGVRYYCEWLVLMRQEPIYDEDEHGLTV

VQRTREDIQAELLSRLRTLQSAHQHSGDMGTDEELLSLMRQLYEQLVPSSVDKNKSGDAR

MIARNFFNPLTNPNSQGGLGISNAGRKPKWLLKKLSGDPTWEEDYKKAMEQKQESSVSFL

LLELRRFGLHPIFLPYTDTVLEVSWAPKKARQWVRKWDYDLFQQSIERMLSWESWTRRVK

ERFEKLVESEKKFYDENFATDPEFIKLAETLEGELQASSQGFVAVDEHAFQIRPRSMRGF

DRVADEWCKLADDAPIEEYEAAIKRVQARLGRNFGSYVLFAHLAKPEYWSLWRSDPTKIL

RFARLRALQRAVARAKRHARLTLPDAIHHPIWIRYDAKGKNIYSYRLLIPEKRSKRYYVE

FSSLIMPDGENRWAEHRNIRVPLAFSRQWERLHFSIMEDGSLCVQYRDPGVDEPLRAELG

GAKIQFDRRYLIRRSSTLSAGECGPVYLNVSVDVNPAHRPDVQVLQSAKLVSVSRDTNRI

YLRPENLSAYWKSQGDGTLPLRVMSVDLGVRSSAAVVICRLEHRDSVVSSGRRTATIYRI

AGTDEFVAVQERAFLLRLPGEGKGTNEDAPLRDVYAQLGTIRQGIQILRSLLRLCDTKTP

DERQEALHGLAQSLEPSGAWKDELHPHLVMLQGVVHDSVDNWKQKVISVHRQMERILGHA

VREWKVARKNAGKPPIRRGAGGLSLRRIRQLEQERRTLVAWSNHAREPGQVVRIKRGTQV

AQWLVERVNHLKEDRLKKLADLLIMTALGYVYDETKPSGHKWDKRYPPCQIILMEDLSRY

RFQSDRPPSENSQLMAWSHRRLLEILKLQADLHKLIVGTVFPAFSSRFDAQSGAPGVRCR

SVKKQDIENAAQGKGWLARELQRLNWTLEWLQPNDLIPTGDGELFVTPACCDRQKGIKIV

HADLNAAQNLQRRFWGGHAESLCRVTCDVVERDGRRYAVPRISNAFADSFYKVFGQGVFV

STDEEDVYRWMVGEKISSRGRSRGRTSDEEAEAETWIDEAREQQGKVIALFRDASGQIHG

GDWLVAKVFWGWVERLVTARLLSRMSEREAAAHKE

>AkaCas12b1

MAVKSIKVKLRLSECPDILAGMWQLHRATNAGVRYYTEWVSLMRQEILYSRGPDGGQQCY

MTAEDCQRELLRRLRNRQLHNGRQDQPGTDADLLAISRRLYEILVLQSIGKRGDAQQIAS

SFLSPLVDPNSKGGRGEAKSGRKPAWQKMRDQGDPRWVAAREKYEQRKAVDPSKEILNSL

DALGLRPLFAVFTETYRSGVDWKPLGKSQGVRTWDRDMFQQALERLMSWESWNRRVGEEY

ARLFQQKMKFEQEHFAEQSHLVKLARALEADMRAASQGFEAKRGTAHQITRRALRGADRV

FEIWKSIPEEALFSQYDEVIRQVQAEKRRDFGSHDLFAKLAEPKYQPLWRADETFLTRYA

LYNGVLRDLEKARQFATFTLPDACVNPIWTRFESSQGSNLHKYEFLFDHLGPGRHAVRFQ

RLLVVESEGAKERDSVVVPVAPSGQLDKLVLREEEKSSVALHLHDTARPDGFMAEWAGAK

LQYERSTLARKARRDKQGMRSWRRQPSMLMSAAQMLEDAKQAGDVYLNISVRVKSPSEVR

GQRRPPYAALFRIDDKQRRVTVNYNKLSAYLEEHPDKQIPGAPGLLSGLRVMSVDLGLRT

SASISVFRVAKKEEVEALGDGRPPHYYPIHGTDDLVAVHERSHLIQMPGETETKQLRKLR

EERQAVLRPLFAQLALLRLLVRCGAADERIRTRSWQRLTKQGREFTKRLTPSWREALELE

LTRLEAYCGRVPDDEWSRIVDRTVIALWRRMGKQVRDWRKQVKSGAKVKVKGYQLDVVGG

NSLAQIDYLEQQYKFLRRWSFFARASGLVVRADRESHFAVALRQHIENAKRDRLKKLADR

ILMEALGYVYEASGPREGQWTAQHPPCQLIILEELSAYRFSDDRPPSENSKLMAWGHRGI

LEELVNQAQVHDVLVGTVYAAFSSRFDARTGAPGVRCRRVPARFVGATVDDSLPLWLTEF

LDKHRLDKNLLRPDDVIPTGEGEFLVSPCGEEAARVRQVHADINAAQNLQRRLWQNFDIT

ELRLRCDVKMGGEGTVLVPRVNNARAKQLFGKKVLVSQDGVTFFERSQTGGKPHSEKQTD

LTDKELELIAEADEARAKSVVLFRDPSGHIGKGHWIRQREFWSLVKQRIESHTAERIRVR

GVGSSLD

>AmaCas12b1

MKSIKVKLMLGHLPEIREGLWHLHEAVNLGVRYYTEWLALLRQGNLYRRGKDGAQECYMT

AEQCRQELLVRLRDRQKRNGHTGDPGTDEELLGVARRLYELLVPQSVGKKGQAQMLASGF

LSPLADPKSEGGKGTSKSGRKPAWMGMKEAGDSRWVEAKARYEANKAKDPTKQVIASLEM

YGLRPLFDVFTETYKTIRWMPLGKHQGVRAWDRDMFQQSLERLMSWESWNERVGAEFARL

VDRRDRFREKHFTGQEHLVALAQRLEQEMKEASPGFESKSSQAHRITKRALRGADGIIDD

WLKLSEGEPVDRFDEILRKRQAQNPRRFGSHDLFLKLAEPVFQPLWREDPSFLSRWASYN

EVLNKLEDAKQFATFTLPSPCSNPVWARFENAEGTNIFKYDFLFDHFGKGRHGVRFQRMI

VMRDGVPTEVEGIVVPIAPSRQLDALAPNDAASPIDVFVGDPAAPGAFRGQFGGAKIQYR

RSALVRKGRREEKAYLCGFRLPSQRRTGTPADDAGEVFLNLSLRVESQSEQAGRRNPPYA

AVFHISDQTRRVIVRYGEIERYLAEHPDTGIPGSRGLTSGLRVMSVDLGLRTSAAISVFR

VAHRDELTPDAHGRQPFFFPIHGMDHLVALHERSHLIRLPGETESKKVRSIREQRLDRLN

RLRSQMASLRLLVRTGVLDEQKRDRNWERLQSSMERGGERMPSDWWDLFQAQVRYLAQHR

DASGEAWGRMVQAAVRTLWRQLAKQVRDWRKEVRRNADKVKIRGIARDVPGGHSLAQLDY

LERQYRFLRSWSAFSVQAGQVVRAERDSRFAVALREHIDNGKKDRLKKLADRILMEALGY

VYVTDGRRAGQWQAVYPPCQLVLLEELSEYRFSNDRPPSENSQLMVWSHRGVLEELIHQA

QVHDVLVGTIPAAFSSRFDARTGAPGIRCRRVPSIPLKDAPSIPIWLSHYLKQTERDAAA

LRPGELIPTGDGEFLVTPAGRGASGVRVVHADINAAHNLQRRLWENFDLSDIRVRCDRRE

GKDGTVVLIPRLTNQRVKERYSGVIFTSEDGVSFTVGDAKTRRRSSASQGEGDDLSDEEQ

ELLAEADDARERSVVLFRDPSGFVNGGRWTAQRAFWGMVHNRIETLLAERFSVSGAAEKV

RG

>AsCas12a

MTQFEGFTNLYQVSKTLRFELIPQGKTLKHIQEQGFIEEDKARNDHYKELKPIIDRIYKT

YADQCLQLVQLDWENLSAAIDSYRKEKTEETRNALIEEQATYRNAIHDYFIGRTDNLTDA

INKRHAEIYKGLFKAELFNGKVLKQLGTVTTTEHENALLRSFDKFTTYFSGFYENRKNVF

SAEDISTAIPHRIVQDNFPKFKENCHIFTRLITAVPSLREHFENVKKAIGIFVSTSIEEV

FSFPFYNQLLTQTQIDLYNQLLGGISREAGTEKIKGLNEVLNLAIQKNDETAHIIASLPH

RFIPLFKQILSDRNTLSFILEEFKSDEEVIQSFCKYKTLLRNENVLETAEALFNELNSID

LTHIFISHKKLETISSALCDHWDTLRNALYERRISELTGKITKSAKEKVQRSLKHEDINL

QEIISAAGKELSEAFKQKTSEILSHAHAALDQPLPTTLKKQEEKEILKSQLDSLLGLYHL

LDWFAVDESNEVDPEFSARLTGIKLEMEPSLSFYNKARNYATKKPYSVEKFKLNFQMPTL

ASGWDVNKEKNNGAILFVKNGLYYLGIMPKQKGRYKALSFEPTEKTSEGFDKMYYDYFPD

AAKMIPKCSTQLKAVTAHFQTHTTPILLSNNFIEPLEITKEIYDLNNPEKEPKKFQTAYA

KKTGDQKGYREALCKWIDFTRDFLSKYTKTTSIDLSSLRPSSQYKDLGEYYAELNPLLYH

ISFQRIAEKEIMDAVETGKLYLFQIYNKDFAKGHHGKPNLHTLYWTGLFSPENLAKTSIK

LNGQAELFYRPKSRMKRMAHRLGEKMLNKKLKDQKTPIPDTLYQELYDYVNHRLSHDLSD

EARALLPNVITKEVSHEIIKDRRFTSDKFFFHVPITLNYQAANSPSKFNQRVNAYLKEHP

ETPIIGIDRGERNLIYITVIDSTGKILEQRSLNTIQQFDYQKKLDNREKERVAARQAWSV

VGTIKDLKQGYLSQVIHEIVDLMIHYQAVVVLENLNFGFKSKRTGIAEKAVYQQFEKMLI

DKLNCLVLKDYPAEKVGGVLNPYQLTDQFTSFAKMGTQSGFLFYVPAPYTSKIDPLTGFV

DPFVWKTIKNHESRKHFLEGFDFLHYDVKTGDFILHFKMNRNLSFQRGLPGFMPAWDIVF

EKNETQFDAKGTPFIAGKRIVPVIENHRFTGRYRDLYPANELIALLEEKGIVFRDGSNIL

PKLLENDDSHAIDTMVALIRSVLQMRNSNAATGEDYINSPVRDLNGVCFDSRFQNPEWPM

DADANGAYHIALKGQLLLNHLKESKDLKLQNGISNQDWLAYIQELRN

>Ba2Cas12b1

MAIRSIKLKLKTHTGPEAQNLRKGIWRTHRLLNEGVAYYMKMLLLFRQESTGERPKEELQ

EELICHIREQQQRNQADKNTQALPLDKALEALRQLYELLVPSSVGQSGDAQIISRKFLSP

LVDPNSEGGKGTSKAGAKPTWQKKKEANDPTWEQDYEKWKKRREEDPTASVITTLEEYGI

RPIFPLYTNTVTDIAWLPLQSNQFVRTWDRDMLQQAIERLLSWESWNKRVQEEYAKLKEK

MAQLNEQLEGGQEWISLLEQYEENRERELRENMTAANDKYRITKRQMKGWNELYELWSTF

PASASHEQYKEALKRVQQRLRGRFGDAHFFQYLMEEKNRLIWKGNPQRIHYFVARNELTK

RLEEAKQSATMTLPNARKHPLWVRFDARGGNLQDYYLTAEADKPRSRRFVTFSQLIWPSE

SGWMEKKDVEVELALSRQFYQQVKLLKNDKGKQKIEFKDKGSGSTFNGHLGGAKLQLERG

DLEKEEKNFEDGEIGSVYLNVVIDFEPLQEVKNGRVQAPYGQVLQLIRRPNEFPKVTTYK

SEQLVEWIKASPQHSAGVESLASGFRVMSIDLGLRAAAATSIFSVEESSDKNAADFSYWI

EGTPLVAVHQRSYMLRLPGEQVEKQVMEKRDERFQLHQRVKFQIRVLAQIMRMANKQYGD

RWDELDSLKQAVEQKKSPLDQTDRTFWEGIVCDLTKVLPRNEADWEQAVVQIHRKAEEYV

GKAVQAWRKRFAADERKGIAGLSMWNIEELEGLRKLLISWSRRTRNPQEVNRFERGHTSH

QRLLTHIQNVKEDRLKQLSHAIVMTALGYVYDERKQEWCAEYPACQVILFENLSQYRSNL

DRSTKENSTLMKWAHRSIPKYVHMQAEPYGIQIGDVRAEYSSRFYAKTGTPGIRCKKVRG

QDLQGRRFENLQKRLVNEQFLTEEQVKQLRPGDIVPDDSGELFMTLTDGSGSKEVVFLQA

DINAAHNLQKRFWQRYNELFKVSCRVIVRDEEEYLVPKTKSVQAKLGKGLFVKKSDTAWK

DVYVWDSQAKLKGKTTFTEESESPEQLEDFQEIIEEAEEAKGTYRTLFRDPSGVFFPESV

WYPQKDFWGEVKRKLYGKLRERFLTKAR

>BaCas12a

MESPTTQLKKFTNLYQLSKTLRFELKPVGKTKEHIETKGILKKDEERAVNYKLIKKIIDG

FHKHFIELAMQQVKLSKLDELAELYNASAERKKEESYKKELEQVQAALRKEIVKGFNIGE

AKEIFSKIDKKELFTELLDEWVKNLEEKKLVDDFKTFTTYFTGFHENRKNMYTDKAQSTA

IAYRLVHENLPKFLDNTKIFKQIETKFEASKIEEIETKLEPIIQGTSLSEIFTLDYYNHA

LTQAGIDFINNIIGGYTEDEGKKKIQGLNEYINLYNQKQEKKNRIPKLKILYKQILSDRD

SISFLPDAFEDSQEVLNAIQNYYQTNLIDFKPKDKEETENVLEETKKLLTELFSNELSKI

YIRNDKAITDISQALFNDWGVFKSALEYKFIQDLELGTKELSKKQENEKEKYLKQAYFSI

AEIENALFAYQNETDVLNEIKENSHPIADYFTKHFKAKKKVDTSTSSVEKDFDLIANIDA

KYSCIKGILNTDYPKDKKLNQEKKTIDDLKVFLDSLMELLHFVKPLALPNDSILEKDENF

YSHFESYYEQLELLIPLYNKVRNYAAKKPYSTEKFKLNFENATLLKGWDKNKEIDNTSVI

LRKRGLYYLAIMPQDNKNVFKKSPNLKNNESCFEKMDYKQMALPMGFGAFVRKCFGTAFQ

LGWNCPKSCINEEDKIIIKEDEVKNNRAEIIDCYKDFLNIYEKDGFQYKEYGFNFKESKE

YESLREFFIDVEQKGYKIEFQNISENYIHQLVNEGKLYLFQIYNKDFSSYSKGKPNMHTM

YWKALFDPENLKDVVYKLNGQAEVFYRKKSIEDKNIITHKANEPIENKNPKAKKTQSTFE

YDLIKDKRYTVDKFHFHVPITINFKATGNNYINQQVLDHLKNNTDVNIIGLDRGERHLIY

LTLINQKGEILLQESLNTIVNKKFDIETPYHTLLQNKEDERAKARENWGVIENIKELKEG

YLSQVVHKIAKLMVDYNAIVVMEDLNTGFKRGRFKVEKQVYQKLEKMLIDKLNYLVFKDK

DPNEVGGLYNALQLTNKFESFSKMGKQSGFLFYVPAWNTSKIDPTTGFVNLFYAKYESIP

KAQDFFTKFKSIRYNSDENYFEFAFDYNDFTTRAEGTKSDWTVCTYGDRIKTFRNPEKNN

QWDNQEVNLIEQFEAFFGKHNITYGDGNCIKKQLIEQDKKEFFEELFHLFKLTLQMRNSI

TNSEIDYLISPVKNSKKEFYDSRKADSTLPKDADANGAYHIAKKGLMWLEKINSFKGSDW

KKLDLDKTNKTWLNFVQETASEKHKKLQTV

>BdCas12a

MIYNVKLKKLTNSSDVFSLFSRKYSLSKTLKFELKPTAETKQHLQDFIVSDTKRAKEYKE

LKKIIDEFHKDYIELTLSHKNILDKDKLTDFCKLWSDNNLLDEIKEKHNLKEKTKEEVIK

KMEQQFRKDIVEQFKTLNPLLERFFKTADEEQLLGFLKKLDNKLWTKFKEKKEKALLDEY

LQEPIKQKNIEKSILFSSELIKYLLPVWLKNSQLTDKEDKQKIIKNFGRFTTYLTGFNEN

RKNMYSVEEQGTAIAHRIINENLMKFLGNLQAYEKIKSSHPELQQSFKNMKSDFKEEFDY

FSLENIEDLFKPEFFNSCLSQKEIDFYNTLLGGKTVEDGKKLQGINEYINLYRQKKKSED

IDIKIKYSNKNLPTMELLYKQILSDRESHSFILAEFENKKELLEALKSFWSVLFEEKTYQ

DYFHTKKKTSLWNNLHSLLTGRSYYDLEGAYFKSSELNKLSHNLFKDYRIITEALNENYD

KIKVTLESYKKVLKKEDKKEFEKNIKSLKGFQTEWKKILSFEKNKEELKFSDEILASYFE

FPIDKKTKKKNQKDFYSLQEIKDHIELYSKESDELKEDLDNLKKKLKDYKKDNIISSWFK

YQFESKQNLDFFLQRNHEKENEDKREQNKNSLFSHIENHYQKIQNLSLDEKEFKKEEVED

IKFFLDLILHVLHLMKPFYLEGKASNLLDKGINNEIEVLYKKLKPIEKLYNQTRNYIAKK

KSRYNKVKINFEDSTLLDGWDLNKEKDNLAVLLRKKDAVIGWKYYLGVMNKTTKDLFDYH

IKLDDSEKVKQKKEELQDLILHRENDGNFYEKMNYKLLPDPSKMLPKVFFSKKNLDSFKP

SEEIVRIRDNKTYAKNGGQDFSKADCHKFIDFYKESLKKHKEWNDFFKFNFSPTSQYNDI

SDFFQEVKNQGYNLQFDKIKSSYIEEKVKTGELFLFEIYSKDFSTKSKNRKNSKDNLHTI

YFKGLFEKENLKDTVLKLNGKAEIFYRKATKKFNITHKKNTELENKNKNNPNKHSIFNYD

LIKDKRFTEDKFFFHFPISLNFNSKGMKSYLFNQEVLKCLKGNKEVNIIGIDRGERHLAY

YTIINQKREVLSQGSFNKMESSYKDNSGKEVKIEKDYHELLESKEKERDKSRKEWNKIEN

IKELKSGYLSYLVHKISKLMIEHKAIVIFEDLNLGFKRGRFKFEKQVYQKLEKALIDKLN

YLVFKDKKSSELGGYLNAYQLTAPFESFQRMGKQTGYLFYVPAYYTSKVCPLTGFINLIY

PKYKNVKESQQFFEKFDRIYFDKNKNYFVFEYQDKKVNPSKKTESIETLWKVCTHGEERY

KWDVKDRKMIEVNVTENLKKLFEEHKIEYRQIADLKSTITKQEKKDFFSKLIDGLKITLQ

LRHINPDSKDEKEKDFILSPVADESGRFFDSRKAKEGEPKNADANGAYHIALKGLRTLEN

ITSDKKDKLKLQAITNKDWFSFLKENSNKKIPKVG

>Br1Cas12b1

MAIRSIKLKLKTRTGPEAQNLRKGIWRTHRLLNEGVAYYMKMLLLFRQESTGGQTKKELQ

EELVRHIREQQQKNRADKNTQALPLDKALEALRQLYELLVPSSIGQSGDAQIISRKFLSP

LVDPNSEGGKGTSKAGAKPTWQKKKEANDPTWEQDYEKWKKRREEDPTASVITTLEEYGI

RPIFPLYTNTVADIAWLPLQSNQFVRTWDRDMLQQAIERLLSWESWNKRVQEEYSKLQEK

MTQLNEQLEGGQEWISLLEQYEEQREQELIENMTAANDKYRITKRQMKGWNELYEQWSTV

LPNASHEQYREALKRVQQRLRGRFGDAHFFQYLMKEEHHLIWKGNPQRIHYFVARNELKK

RLEEAKQNATMTLPDARKHPLWVRFDARGGNLQDYYLTAEADNPRSRRFVTFSQLIWPNE

SGWMEKQDVEVELALSKQFYQQVTLQKNDKGKQEIEFKDKGSGSTFSGHLGGAKLQLERG

DLEKEEKDFEGGEIGSVYLNIVIDFEPLQEVKNGRLQSPYGQVLQLVRRPNEFPKVTTYK

SEELVEWMKASQNHSSGVESLESGFRVMSIDLGLRTAAATSIFSVEESNDANAAGFSYWI

EGTPLVAVHKRSYMLKLPGEQVEKQVREKRDERQDQQRRVRFQIRILSQVIRMAKKQNRE

RADELDHLSQALEKQKSLLDQTDRTFWNGIVCDLTDALREKEGGWEQAVVQIHRKAEEHV

GKVVQAWRKRFDADERKGIAGLSMWSIEELDSLRKLLISWSRRTRNPQEINRFEQGHTSH

QRLLTHIQNVKEDRLKQLSHAIVMTALGYVYDEKKLEWFAKYPACQVILFENLSQYRSNM

DRSTKENSTLMKWAHRSIPKYVHMQAEPYGIQVGDVRAEYSSRFHAKTGTPGIRCKMVSG

HDLQGRRFENLQKRLISEQFLTEEQVKQLRPGDIVPDDSGEWFVTLTNGSGDKEVVLLQA

DINAAQNLQKRFWQRYNELFKVSCRVLIRGEEEYLIPKAKSVQAKLGKGLFVKKTDTVMK

DVYVWDSQAKLKGKTTFTEESESPEQLEDFQEIIEEAEEAKGTYRTLFRDPSGVFFPEFV

WNTQKDFWSEVKRRLYGKLRERFLMKTR

>ca2Cas12a2

MNSIKNEYQLSKTLRFGLTKKKKLLKDDCNEIIYESHTELKELVLISEKKIMESVYINQK

AKLDLSVDQIDTCLSSIKNFIDSWKGIYPRADQIAIDKDYYKILCKKITFDGFWIDEKTK

TKKPQSRTILLSELSKKDASGKERKQHILDYWKNNIFSAIEKYEVVSRELKQFQKALKIQ

RTDNKPNEVELRKLFLSLANIILDILKPLVNGQICFPKIEKLDISKTDNKNLIDFATNHK

FQSDLLNEIAELQHYFEENGSNVPFCRASLNPKTIIKSKLSTDNNIDKEIKQLGLDRILN

EYLSAPYFDNSIIHLSAKEKLNKIEDKKENYITRGLLFKYKPIQIMLHHEIAKTLSKEIG

KSEENIIEFLGNIGQIKSPAKDYEVSKEDFNINNYPLKVAFDFAWENVARNLYHTDTHAP

IDECRKFLADNFDIKIEDNNLKLYANLLELNALLSTLKYGKPKDETSIKQNIKDLLNKIS

WNEIGKSGQKNKTNIENWLNNKDKIDNQNGIENAKKQIGLFRGSLKNKVPKYYKLTETYK

DISMKMGKIFATMRDKITDEAELNKVSHYAMIVEDDNKDKYILLQEFTDKKEECIYSKTQ

THNSDFTTYSVNSITSSAIAKMIRKVKAEELRKNQYNKDTFSIEETKEEKENRIIKEWKQ

FLKDKQWDYEFNLDTKNKNFEELKKEIDSKCYKLNISYIDKKTITDLVENKNCLLLPIIN

QDLSKEEKTQNNQFTKDWDAIFSQNTPWRLTPEFRISYRKPTPNYPISDKGDKRYSRFQM

IGHFLCDYIPQSNTYISNREQIANYKDNEKQEQAVQCFHDKLLGKTEKEAKNEKLIALQA

KFGSISRTNITQEKKKEKFYVFGIDRGQKELATLCVIDQDKKIIDDFDIYTRSFNSKTKQ

WDHTFLEKRAIMDLSNLRVETTISIDGKTEKKKVLVDLSKVKVKDKQGHYSKPDKMQIKM

QQLAYIRKLQFQIQTNPDVVLAWYSDNNTQDLILENFVRKDDNDNKGLVSFYGAAVEELK

DTLPIEEILNMLKQFKELKEKEKAGENVKYEIDRLIQLEPVDNLKTGVVANMVGVIAFLL

EKFNYQVYISLEDLSQPFDNKINGGITGVPIKTNKESGRMADVEKYAGLGLYNFFEMQLL

KKLFRIQQKSTTILHLVPAFRAQKNYDHVTVGQDNIKGQFGIVFFVNANATSKTCPICGA

NNSEKPDKNKYPNAHKELAKDGKEVWIERDKSNGNDIIRCFVCGFDTTKTYEDNPAKFIK

SGDDNAAYLISVSAIKAYELATILAIEKYK

>ca6Cas12a2

MNHIYNNYQVSKTLRFGLTQKQKIRRPGYTGELYESHKVLKELVKISEEKVKNLIVPAKN

EELLSSLDSVKWTLTEIREFLDQWRYIYNKSNQIALDKSYYLILSKKLGFNDEKKSRVIK

MIEIKDDIKEKIINYWAFNLNESNQKLLMVNEMVNTQLKALEINRTDHKINEIELRKALQ

SLFNTVLDILKPLVYREISFINLEKIEKDSKNSLLEKFATDFQRKIDLLEKIRSLKTHFS

ENGGNVSFCRATFNPKTAIKNPKSNDNSILKEIKKLGIKDILENNENVFYFEKKLAEITA

KEKLEYIIKDSESFLIRSLLFKYISIPAFLHHGIATELAPIISKEKNDLINFMISIGQIK

SPAKDYADIPNKNDFNVNAYPIKVAFDYAWETVAKSQYHHDINAPVSMCKTFLDENFENC

TKTKYFTLYSDLLELHTLLSTLDYGNPSMEDSIIDKANKIIAKIDNKEHKTKDKDLDKDI

DKYKETIKNRLNHKNFNDKQRYSDAKKELSQFRGKLKNENDIYRKLTESYKKIAMNTGKI

FAEMRDKISNASEQNKISHHALIIEDHNKDRYLFLQEFTTDKEKQIESICNDQAGQYIVY

WVNSITSKSISKMLSKKRIEKLKQKKIINNSIKTSILSDAEKEARDIKEWVSFIKEKGWD

IDFNLDLQNKNLEEIKKEVDAKAYKLKETLISQKTLSNLVKEGNCLLFPIINKDLVKKVK

TEKNQFTKDWNSIFKKDNLWRLTPEFRVSYRQATPGYPTSDIGTKRYSRFQMTAHFLCDF

LPQGTKYISNREQIENYKSSEKQKEAVEIFHQQIENDNNNVISTQSLNHLARHFGSKNIK

KKHNTIEKKFYVFGIDRGQKELATLCIIDQDKKIEGPFKIYTRSFNTKTKQWEHQFYEER

YILDISNLRVETSISIDGKPDQQKILVDLSYYKEGEKFIKLPKMQVKLQQLAYIRKLQYQ

MQRNPETVLDWSYKNTDDKSILENFVDKPNGEKGLVSFYGAAVIELKDTLPLSEIKDMLE

RFKELKGKEKNGEDVSQQLNELTQLKSVDHSKYGVVANMVGVIAHLLERYDYKAYISLED

LTKPYSAIDGITGQKTDAKSISGKQQDVEKYAGLGLYNFFEIQLLKKLFRIQKDSQNTLH

LVPAFRATKNYENLIAGEDKVKNRFGIVYFVDPKSTSIMCPSCGKTNNSSNKEKRVVRDK

KNGNDIIYCEFCGFDTRNDYKENPLKFIKSGDDNAAYIISTHTAKKAYELAKSIL

>ca7Cas12a2

MSLAAFTNQYQLSKTLRFGFTQKEKVRKENFDGSIYQSHAALRELTIESERLIKGKLKSN

TDTALPLEKIRACIEEIKRYTDTWSKIFTRDDQLALSKEYYRVMARKARFDAFWKNYRDV

KQPQSQIVRLSSLKSKYNGKERKAYLVDYWAGNLQTVKQRLVDFEPAIRQFESALKDNRT

DRKLNEVDFRKMFLSICKLVNETLVPLCNSSLCVPDLEKLLDNEASQELRDFVMMDIFQL

QEQIEALKIYFGENGGYVPYGRTTLNKYTALQKPHAFDEEIEAILVKLKLSDVINNLIKQ

DDVSDYFENVKDKIGQLSNSSMSVIECVQLFKYKPIPVSVRYSLVEYFHRKLGIDKDELG

TLLDTIGKPKSPAKDYADLQDKGDFNLYKYPLKVAFDFTWESLAKAQYHEGLNFPEVQCQ

KFLENIFFVNTSCEAFKTYALLLHLRGLLAKLDHEEPNDREAIIDKAISLMNEDAFPKVP

LRGTKGDSANQAILSWLQLSKEEQVYKKEKKDQSYNQYEKAKNKIGLLRGEQKNKIGKYR

EVTEQFKDLASNFGKLFGALREKFQAKNELNKITHYGTIIEDNNQDRYVLLYPLSEGIID

LDKLFVHEESGTLTSYYVKSLTSKTLNKLIKNKGGFKDFHMDGQQPDWERVKKRWSVYKD

DKAFLKYVKRCLNESEMAKAQNWGEFGWDFSSCDSFEEIEREVDKKGYSFKNDRKLSEDT

VKRLVKEEKCLLLPIINQDIIVEETKLRNQFSKDWVNIFDADCTEYRLHPEFGMSYRMPT

PNYPKPEQKRYSRFQMIGYFQCEIVPIKTEYLSKKEQIEIFNDADAQKEAVEKFNEIVNG

SVKPNDYVVIGIDRGLKQLATLCVLNKDGAIQGGFEIYTRSFNADKKQWEHRFMDNRDIL

DLSNLRVETTVDGKKVLVDLSSIKVKDQRGNYTQDNQQKVKLKQLAYIRKLQYQMQVNPE

KVKAFAAQHRTPQDIKDHMKELITPYKEGSHFADLPLDRIKYMLEAFCAFHTENDQTSLR

ELIELDAADNLKSGIVGNIVGVIAFLLKRFSYNAYISIENLTRAFYNQRDGLSEKEIPRD

HDFMDQENLVLAGLGTYHYLEVQLLRKLFRIQCDAGIINLVPAFRSNDNYETTRKLSKKQ

GVEYVCKPFGIVHFVDPMYTSKKCPACGGTTVQRGSFKDDITCQNPLCGYGTSLDISEKI

QKLISANKAGQNIHLISNGDENGAYHIALKTLKNLFGNIQNANNERRYVKSFRSK

>ca11Cas12a

MKNLSDFTNLYSLSKTLRFELIPQGKTLEHIEKNGLLEQDRQRADSYQKVKKIIDEYHKA

FIKRSLKGCKLSNELLTDYYFYYNLKLKRDETQKKKIEELQSKLRKEVASKFLNLKELES

ADLYNASKKDNRAEGKLNSWLKEKPHIAAKLIIDLKLNQHYSFEDDLKRFESFATYFTGF

HENRKNMYSADDKATAIAYRLIHENLPKFIDNINVFEKVKNSDVAQNFEKLYKDLEEYLN

VLKIEDIFQLDSFSDFLTQEQIDVYNAVIGGKAEEQGKPKIQGLNEHINLYNQKQTDKDK

KLPKLKPLFKQILSDRDTISWLPEEFKKGNDNEVLESIEKCYQELCNEIFAPDKKQPLIG

LLLNLNDYDLNKIYLRNDTSLTDISQQLFGSWAVIQKAVETKYERENPRKKNEEEEKFAK

RKDKFFKNYDNFSIGYLSDCLKCLDNYDDLPTKNIADYFTNLGKKEDSENLFSQIEKNYA

EVKNLLNTPYSGDKDLAQDKNAVEKIKIFLDSIKSVQWFVKPLLGKNESDKDEKFYGEFT

PFWKELDKITPLYNKVRNYMTQKPYSEEKIKLNFENSTLCNGWDKNKEQDNTAILLRKDG

NYYLAIMNYRAKTNFDKTEIAKESESFYEKADYKLLPGPNKMLPKVFLSTKGKAEFNPSS

EILENYENGCHKKGDTFDKDKMHKLIDFFKISIAKHKDWKNFGFNFSDTPSYESIDKFYK

EVLEQGYKISFRNISEKHINELVEAGKLYLFQIYNKDFSPYSKGTPNMHTLYWRELFSEE

NLKNVVYKLNGEAEVFFRKSSIKSENKIEHKANEVIKNKNEQNKKLESTFKYDLIKDKRY

TVDKFQFHCPITMNFKAEGTENINPKVNQFLKENANNIHIIGIDRGERHLLYLSLIDAKG

KIIKQFSLNEIVNKHNENTYTTNYHSLLDEREKERQKERESWKVIESIKELKEGYISQVI

HKIAELMVEYNAIVVLEDLNMGFMRGRQKVEKSVYQKFEKMLIDKLNYLVDKKKQASEQG

GTLQALQLTNKFESFQKLGKQSGFLFYVPAWNTSKMCPVTGFVNLFDTRYENVEKAKAFF

SKFDSITFHSAKNYFEFEVKDYSKFSGKAEGTRLNWTICTNGTRIETFRNPEQNNNWDNR

EVNITDEFVKLFGTKNADFKNLIQQQSEKAFFERLLYLFKLTVQMRNSITNSDIDYMISP

VADKNGRFYDSGNCDEALPDNADANGAYNIARKGLWIIEQLKQTEDLKKPRLAVSNKEWL

QFVQKLNK

>ca11Cas12a2

MEKNEIAISEYQTQKTIRFGLTATNQNLYSEEIMKLLDISEKRVEKQAEQAKKVNNDADK

NNQLRCCLDQIKEYLKTWSNIYPQIDFLAITKDFYKVISKKARFDFDKGNGSEIKLSYLQ

STYYNKKRYLYIIESWKENLRKTENLYRKSDDLLKVFEEAKNQNRDDKKLNKVELRKTFL

SLFNLVNESLKPLIEGNLFIVNDDKIDEQNPKHDCVSVFISKTEERRKLYDYICDLQDYF

KDNGGYVPLGRVTLNKWTALQKSNNRDAEINRIIKELKINSVSIQNIEYEYNNFANNFKE

KKDENGKIVKNNAGNIIWDLKADAKSVIEICQFFKYKQVPINARLNLAKRLEKSIDFLSE

FGVSKSPALDYKNDKNNFNLTNYPLKIAFDYAWENCAKAKHEEIPFPKEQCEKYLKDVFD

IDIECKEKCQNKECKGCEKCRGYYLNKYADLIRFKILLGRLKAEFHKTDEEKNKSNIQEL

RNIFRDLDYRGDKRLNKNEIQKAVNAWFDNKEQSIGRKKEDEIHLMENEKNKFSLSMQII

GQERGGLKSRISKYKALTEMFKVCASKFGKQFADLRDYFNEAYEVDKIKYRAWIIEDEKQ

NRFILFVNKEKEVDLTSEEGDLYFYEVKSLTSKSLVKFIKNRGAYPDFHKINNRQIDLNS

GEKDSRGNFIDDVKIHWSTYKNNQKFLDKLKDCLQNSTMATVQKWSEFEFEFDFSNCDTY

EKLEKEIDRKGHKLERKTISLTTITNLVENTACLLLPIVNQDLNKGNKQAKNQNQFTKDW

FDIFENKKRLHPEFNIFYRFKTKDYPNTKFKNGTEKTKRYSRFQMLAHFGCEVIPQGDYL

SKKEQIAIFNDDKKQTEEVKKYNKNISSDVDYVIGIDRGIKQLATLCVLDKNGVIQGGFQ

LFTRTFNSETKQWEHQELEKRNILDLSNLRVETTITGEKVLVDLASIQTKNGENRQKIKL

KELAYIRDLQYTMQTRASDLLDFASKINSADDITENNIKNFISPYKEGEKYADLPQKEMF

DLLTEWKNAEEEGKRKIAELDPADNLKSGIVANMVGVVALLCAKYKYRVRIALEDLTRAY

GIQKDALSGATIFQNDEDFKEQENRRLAGVGTMQFFEVQLLKKIFKVQIDKDLHLIPAFR

SIANYEKIVRRDKQNSGDEFVNYPFGIVCFVVPKYTSKRCPKCEKTNVNRKENIVICKEC

GFQTKEGNPYEKNNIHFITDGDQNGAYHIAKKAL

>ca14Cas12a2

MLESFKDLYEVRKTERIGLTQPNKKGEKISHLELNELVKTSLERIKSDIDKDDNSIIKNE

KNLIKNSRQFVNELNVQFGTWKNIYKRYDLISVNKDYYRILARKAKFDAFWKEGRTKYPK

ANQITLSSLQKGGRKEKIIEYWKEIIERSYYLISIFTQKLDQYENAVSNPEKLSHTKPDF

TDFRKIFLQLLKVSEDYLKPLFDKSINFKTGKKLESEEDIKEINLFISEENNKNIYSLLK

LGNDIREYFEENGAQVPYKKVSLNYYTAVQKPNNFEGEIKKAIEDLWIVEFLNKSDSQII

NYLHQGSKQKKNYYLLQNILILLSYYSYLKVKPILFSVKYNLAKFMEKNYKSEINLSYED

ILNKFNLLGRAIDIANDFKNSNNQNNFSLDEYPIKLAFDYAWENTARSLKRTIPFPKEVC

KQFLKDNFDVDIDNADFKLYANLLFIADNLATIEYNNPNNEVELINEIKQAFECISFPFD

KEVYKGHKEAILELLDKEKSQRDYSTILKAKQELGLLRGGLKNKIKKYRDLTQRLIDKKN

SHFGIASFVGKTLATIRDGLKEENELNKISDYGVIIEDSNQDKYLLTLELNGKDIRDRIR

NSLGNGEYKTYEVNSFTSKALNKFIKNPLSEDAKKFHGKYKDEYSFGYENKDGDFTYKIT

KVSKYDEQGKWTGYQESFLSHIKKCLIDSEISREQNWEAFGWNFAGCNTYEEIEKEVDSK

GYQLTENLISMGNLKSLVKDEGCLLFPIINQDISSQKQENKNIFTLDLEKVFEGKECRIH

PEFSIFYRRPIEEHKKENKSGIINRFGRLQLLANLGIEFVPRNPSFKTKKEQNRIAIDQK

KQNQLVQEFNQKKVNTYFEGLDNYYIFGIDRGIKQLATLCVTDKDGVIQDFDIYTKHFNS

ESKKWEYKFHRKDGILDLTNLKIESDRSGNKYIVDISLFQAKDEDGNPTGTNKQNIQLKQ

LAYIRKLQYQMSANEEGVLNFLGKYKNKEEREQNMEELITPYKEGKNFADLPMDIFQEMF

ENYYRLKTDQNLSESEKKNLMKITTELDASESLKKGVVANMIGVIYYLMKKYEYKVKISL

ENLSNAWLFSKDGLSGDVVLNTKNDETMDLKKQDNLALAGVGTYHFFEMQLLNKLFKIST

EEGVLHLVPSFGSVKNYIEIMKIKGKYVYKQFGIVYFVDPRNTSKKCPVCGKGGKKYISR

VDNVVTCKNCGFDTSSDNSILINNYKKQGKNIHFIKNGDDNAAYNIGEKIR

>ca17Cas12a2

MEEFKNLYQLSKTLRFGLTLKDKKRKDGYTKEIYGSHNQLKDLVRFTEKRIEEEITTVGI

NYTAELLAEKIRKCLNNMQSFISQWNKIYYRKDQVAVDKDFYKKLSKKNWIQRFLVCKEK

NKREIKRIKQPQARIIKLSALDEEDEYNRTRKDYIIDFWRENIQNAFNKYKEVERVLEEF

DFAIKANRADKSPDEVEMRKMFLSLANIVCEVLVPLCNGSICFPDIEKMSDKDENKYLRE

FAVDYKTKIDLFDSISELRKYFEENGGNVPFCRATLNPKTKIKNPSSTDNSIEAEIEKIG

MEKYLQTTDEIITAINKIKQMQDSKLPLIKRALLFKYKTIPAVVQFDLAKSIGKKLNKEE

QELRTIIRTIGQTASPAKDYADLQHKKEFNINQYPLKPAFDYAWESLAKAEYHNDIEFPN

QICKQFLKDIFNVDTETDSYFKLYAQFLYLRELLATLEHSQPTNPEKIVKIIEELLEKIN

AWKEIDDKNSEKYKTAISNWLKTFNKEDNNFKNAKQKIGLFRGGLKKKAGDKSRWEYLRK

PDEKEAPKLTAYNALTQIYKDIATNLGRNLADMRDKITNESELNKISHYAAIIEDKNGDK

YVLLKKSDKDEEFALPCEKDAEYKTYIVNSITSSAIAKMIRKKRAADLAKNIRLKSEELN

DEQKEAKNLKDWIELIKKQQYDFEFNLNLNNKNFEQIKKEIDAKCYQLQKGNISKQTLEK

LINEQKEWLLLPIINQDLAKRDKSTTNQFSKDWQKIFSDKRCGYRLTPEFRISYRKPTPN

YPQSQIGDKRYSRFQMIAHFLCDYIPQGNLEYKSTRQQIEIFKDYEKQEESVRNFTYKIY

ANNDYLIFGIDRGLKQLATLCVLNKEGKIYGGFDIYTRSFNKDKKQWEHSLSEKRNILDL

SDLRVEKTITGEKVLVDLNSIKVKGNQDNQQKIKLKELAYIRKLQFKMQTEHGTVINFIN

KYRTPEEIQKNIHELITPYKEGEHYADLPTEKICNMLQKFKEFSDKNDGKSKRELIELDS

AGELKNGIVANMVGVVAYLLEKYKYNVYISLEDLTRAYRRQTDGLDGRELFSSNDDKSVD

FKDQENTALAGLGTYHFFEIQLLRKLFRIQQEDGNILHLVPAFRSVDDYEKIIRRDKKID

GNEYVDYPFGIVRFVDPKNTSKRCPLCSSININRHKNIIKCQQKACGFKTPWDKTNDKNI

QYIQNGDENGAYHIAKKTLDNLNNKK

>ca33Cas12a

MSLTQFTNLYPIQKTLRFELKPIGKTEEHIQKNRLLENDQKRADDYDVVKDLIDSYHKWF

IEETLQAVSLDWSALQAAIANQRTEKSPASKKALEAEQALMRKKINKAFESHAKFKLLFS

EKLFKEILPAHLNVLEATDEQRKAVDTFKRFSGYFLGFHENRKNIYSADDISTGIPHRIV

QDNFPKFLNNAQIYAALPPEIKDAINAALQEDLNGRALDEVFSIGFYNTVLTQKGIDFYN

LIIGGRSPEPGQRKIQGINEIIHQYRQQYPDVKIPKMVELFKQILSDRESQSFIPKMYEN

DKEVQESIVHFHDTELRGFTQEDREVDVLTGLANLAKGIAGFDLAHIHVTAASLPELSKT

LLEEWNTLQRYIETYANERFKTKKDKEKFLKSETFSLDELNAVLEHNEQAKRIQSFWNEA

ADERLKAIDETHAKAEAILRKEYTAENKMRENPDDIALIKEFLDAVMDFMRWLKPFCIQS

EVSKDDSFYGEFEPLYKQLALVVPLFNRVRNYATQKLAETAKIKLNFENPTLANGWDKNK

EKDNTSILLMKDGKYFLGVFNVDNKPGVDGACVQTVTEACYTKMIYKLLPGPNKMLPKVF

FSKTGREAFDPPQYILDGYEAEKHKKGDAFDKTFCHNLIDYFKDAIQQHPDWKQFDFQFS

DTASYEDISGFYREVQQQGYKLTFTNIPERIISQWVDEGKLYLFQLSSKDFAAGTTGKQN

LHTLYWTQLFSPENLKDVVVKLNGEAELFYRRKSIEKPFIHRVNEKMVNRRDRDGNLIPE

KIFGELFRYFNAPDQVTLGEEATKWKDKTVVKPVKHEITKDRRYTEDKFLFHVPLTLNFK

ADGARLNEQVRGFLINNPEVNIIGIDRGERNLIYVTIINQKGEIIFTRSFNEVNGFDYHE

KLAQREHERDAARKSWQSIGKIAELKEGYLSQVVHEITKLMIQHNAIIVLEDLNFGFKRG

RFKVERQVYQKFEHALIDKLNYLVFKERKASEPGGVLKGYQFTEPLAAFKDVGKQTGMLF

YVPAAYTSKIDPVTGFANLLNLNYDNERKSKELFSTTFDFIRYNSKEDFFEFGVNYEKAK

THVTDYTSEWTVCTTNEPRYAYNPQTKKTETVMVTERLKAVFAENKIAFEDGRDLKKDIV

ERGSVGLYKTLIWLLKLVMQMRNSNAETGEDYILSPVRNKAGVFYDSRKADDTLPKDADA

NGAYHIALKGLYLLQEVFNKTQSEKDKVDLKITHPDWLAFAQKRHEVK

>ca33Cas12a2

MDADKTTKAINEYQTQKTIRFGLTATNQNLYSEEIMKLLNISEERIIKEKVKVNNDTDKT

NQLRGCLVQIKKYLKTWENIYAQIDFLAITKDYYKVISKKARFDFDKGNGSEIKLSSLQS

THNKKKRYQYIIDFWKENLRKTENLYRKSDDLLKIFEEAKNQNRDDKKLNKVELRKTFLN

LFTLVNESLKPLIEGNLFIVNDDKIDEKNSKHNYVFYFISKTEERRLLYDNICTLQDYFK

NNGGYVPFGRVTLNKWTALQKFNNRDIEINRIIKELKINNISTQKTDYKYNDFTENFKEK

KDENGKVVKNSAGNIIWDLKANAKSVIEICQFFKYKKVPINARLNLAKRLIKDNKLKKEQ

ENTFLSEFGVLKTPAFDYARDKENFNLTNYPLKVAFDYAWENCAKDKYEKIPFPKEQCER

YLQTAFEIDATKDENKKLIDTHLNKYADLLQFKILLERFKAEFHKTNEETNKNNIQKLRN

VFSGLDYHGDNRLNKNQIQKAIEAWFDNKEQNIGKKKENEKLLTENEKNNFSLSMQIIGQ

ERGGLKNGIPKYKELTEMFKVCASKFGKQFADLRDYFNEAYEVDKIKYRAWIIEDDKKNR

FVLFVNKEKAFDLTSEEGDLWFYEVKSLTSKSLVKFIKNRGAYPDFHDVKNSFHYSSIKK

DWQNYKNDPEFLDKLKECLKNSKIAKDQKWAKFCWDFKQCDTYEKLEKEVDRKGYKLEGC

KSEPKTISLTQLTDWVENKDCFLLPIVNQDINKGDKRTKNQNQFTKDWFDIFENKKRLHP

EFNIFYRFPTKDYPNTKFKNGTEKTKRYSRFQMLAYFGCEVIPSGNHLSKKEQIAIFNND

KKQKEEVEKYNKSISSDCDYVIGIDRGIKQLATLCVLDKNGVIQGDFQIFTRTFNKQTKQ

WEHKELEQRNILDLSNLRVETTITGKKVLVDLSKIKDDEGNYTNLKQTIKLKQLAYIREL

QYAMQTRPDDLLDFVKSINSANDITAENIKHFISPYKEGKNYDDLPKVEMFNLLKEWGNA

DENGKRKIAELDPADNLKSGIVANMVGVVAFLCENYNYKVRIALEDLTRAYGIQKDALNG

TAIYQNDEDFKEQENRRLAGVGTMQFFEVQLLRKLFKIQVDKNLHLIPAFRSVDNYEKIV

RRDKQNSGDEFVNYPFGIVCFVDPKYTSQQCPYCNNTHKHKKNDTETGKKAFYRNKGENK

NSLLCEKCGVSTIEGEETLSSKNDNKKQFNIHYITDGDQNGAYHIANKVVINFQKDS

>ca34Cas12a2

MEKIKNKYQVSKTLRFGLTQKEKRRKKGFVGEVYESHTELKDLTDFSVKKIRAEIKQGGN

SGIPIEKIRQCLIVIRRYMNFWENIYYRCDQLSLDKDFYKKLSKKIGFEGFWHEENRKTD

HRIKKPQARTIQLSELNKKDDHYKERKDYIVEFWESNINKAAERFKETESVFEKFEIAIN

ANRDDNRPNEVEMRKMFLSLANIIYETLVPLCNGSISFPNIEKMQNNEENNNLRKFAADD

EFRAGLLTQIEELKAYFEENGGNVHYCRATLNPKTVIKNPNSTDSSIADEVEKTGIERYL

QNKQEIKNKIEKIKDKNLPLIERALLFKYKTVPAGVQFDLAKFLSVKLGKTEQELRTIIR

CIGQIQSPAEDYAKSEDKKGFHLEQYPLKSAFDFAWESLAKAIYHKNVDFPQQQCKVFLE

KNFDIKIDSNANFKLYAQLLYLRENLATLEHSKPTDPDTFEKNIRKLLDEINWSTIDKEK

GSVYKNAISDWLKNKKAKDEKFGKVKQSIGLSRGRLKNKIKKFDDLTKDYKDIATQLGNA

FAAMRDKITNAAELNKISHYATIIEDKNGDRYILLQKVTENEKPVGENWNKNGELKTYLV

NSVTSAAISKMIRKIRTDELRKNEKMQSSITKLNEKQKEEKNINDWKNFIEEKRWDLEFK

LDLKSKNFEQIKKEIDTKCYLFNTGYASQADIKQLVKEKDALLLPIINQDLASKDKIVRN

QFSKDWQMIFSDNSQGYRLTPEFRISYRQPTSNYPQPEEKRYSRFQMIAHFLIDYIPQNN

QYISTREQVELFKDETKQREAIEEFHKQLTPKTEAEQKAESLSALAAKFNNPNNKKQKNN

ASENKPDEKFYVFGIDRGQNELATLCVINQDKKIIGDFEIYIRKFNSEKKQWEHLKLENR

HILDLSNLRVETTIVIDGKPEKKRVLVDLSEIKVKDKNGEYKNPDKMQTKMRHLAYIRKV

QFQIQNNPEGVLDFLKKFKTKNETIYNLVDKENGEKGLISFYGAGNTNEDLPKDDIWKIL

QKFQELKNKTGDDNVKKEIKELIELEPVDNLKNGVVANMVGVIAYLLEKFDYQVYIALED

LTKPFNEDIKDGTTGVKGNYKGEGKRADVEKYAGLGLYNFFEMQLLKKLFRIQQENDNVL

HLVPAFRAVKNYENLIAGKGKIKNQFGIVYFVDANSTSKTCPVCSSIPDKNNSNNKNAKG

KKIINKKGNESVIWVERDKSNGNDIIRCYACGFDTTKNYSENPLKYIKSGDDNAAFIIST

LGIKAYELAKTLVANK

>ca35Cas12a2

MKSFENSCQISKTLRFGATLKEDKKKCKSHEELQEFVDISYKTMKSSATIAESLNEIELV

KKCERCYSEIIKFHEAWGQIYCRTDQIAVYKDFYRQLARKARFDVGDQNSQLITLSSLSS

KHNVYQGLKRSQHITNYWKDNIARQKSFLKDFSQQLHQYKRALENSDKAHTKPNLINFNK

TFSILANLVNEIVIPLSNGAISFPNISKLEDGEESRHLIEFALNDYSDLVGSIGELKDAI

ATNGGYTPFAKVTLNHYTAEQKPHVFKNDIDAKIRELKLIELVEKLKNKTSKQIEEYFSK

FDKFKIYDDRNQSVIIRTQCFKYKPIPFLVKHELAKYIAEDEDAVLKVLDAIGATRSPAH

DYAHNEDVFDLKHYPIKVAFDYAWEQLANGVYTTVSFPEEKCREYLNAIYGCEVTKEPVF

KFYADLLYIRKNLAVLEHKNNLPSNPEEFICKIENTFEKIVLPYKIKEFETYKKAILTWI

NDGRGHEKYTDSKRELGLIRGGLKGRIGAQKMFRKNKKGELIPYYENPYTKLTNEFKNIS

SSYGKTFAELRDKFKEKSEITKITHFGIIIEDNNKDRYLLANGLQHDNTDQSNTQVEAIL

GKLNTSLEFTTYQVKSLTSKILIKLIKNHTTSPNAKSPYADFHTSKIHVDWTKIKKEWDT

YKSNHSLLQYVKDCLTNSTMAKNQNWAEFGWDLESCNSYEAIEHEIDQKSYILQKHTISK

ASIKSLVENGCLLLPIVNQDITSQGRKDKNQFSKDWKQIFEDSKEYRLHPEFAVSYRTPI

KDYPKDKRYGRLQFICAFNCEIIPQNGEFINLKKQIENFNDEDIQKNNVAEFNNKVNEAL

LGKEYVVIGIDRGLKQLATLCVLNKRGKILGDFEIYKKEFVRTENRSKNYWKHTLAETRH

ILDLSNLRVETTVEDNKVLVDQSLTLVKKNRDTPDEKATEENRQKIKLKQLSYIRKLQYA

MQTNEQAVLDLLKDNSDDEEFKKRVEGVISPFGEGQEYADLPIDTMRAMIQDLQGVIAKG

NKQTEKNKIIELDAADSLKQGVVANMIGVVNFILAKFNYEAYVSLEDLSRAYNVAKSGYD

GRYLPSTSQDPDMDFKEQQNQMLAGLGTYQFFEIQLLKKLQKIQSNNTVLRFVPAFRSAD

NYRNIIKLNPKYDNTEYVHKPFGIVHFIDPKDTSSKCPVCGKTGKKNVDRNEKKNNILLC

KACGFRTVWELKQPENIKSEGYCFSEDEIQKTIKNNQKQMEIKKLSDKNIHYIHNGDDNG

AYHIALKSVENLKHKE

>ca36Cas12a2

MKSIVNNYQISKTLRFGLTQKTKIQKEGYNGEIYVSHRELCDLVKISEERIKKSVSASDK

SNLELSIDIIDTCLKQIGVFLSDWQQVYYRKDQVALDKDYYKILCKKIEFDGFWKDDRGQ

RMPNSRIINISELDKRDSLGVERLHYILNYWKDNLVSASQKYSAVEEKIKRFKSAIKINR

TDNKPDEVELRKMFLSLANIVCDTLQPLCYGQISFPKINKLDDSRTDNKKLIKFATEYKS

KNDLLTSIAEQKKYFEENGGNVPFCRATLNPKTAIKDPNSTDNSIKGEIAQLGLDSILKS

FKSYLFFENSLENMSAKEKIDLMKSDGASIIKKGLMFKYKPIPVIVHREVAQELSEDLNK

TEESLSDFLRGIGQAKSPAKDYEELTDKNEFNIEAYPIKVAFDFAWESLAKAKYHSEIDL

PVDSCKKFLKTFDVTPDDANFLLYAQLQELNALISTLEYGYPSDEQSIVKKIKALSNEIQ

WEKVSDRNGQEYKRSISDWADGKRDSDGFKIAKQNIGLFRGGLRNKIDKYYSLTQIYKKT

VMDEGKIFAIMRDKIIGAAEQNKVTYYAAIIEDNTGDKYVLLQELPLNGQDRIYDKMTRD

GDGYVCCFVNSITSRTIAKQLRKKRMAELKKNKAWGAYNDNVSNTQQPVLSDEEKEERNI

REWKSFISEKRWDCEFNLNFKGKNFEEIKKEIDAKGYELENRRLSREALDELVKNNKCLL

LPIVNQDIIKENKTESNQFTKDWNSIFDDNSPWRLTPEFRVSYRKPTPDYPVSSKGDKRY

SRFQMIAHFLCDYIPNSDSYVSVREQIENYKDDKKQETAVKDFHNRLLGKTEEQKIIDRL

GAFQGWGNVIIKKTKPEKQEGSKEKFFVFGIDRGQNELATLCVIDQDKKIQGGFKIYTRS

FNSEKKQWEHKFLEERNILDLSNLRVETTIVIDGQEKKEKVLVDLSEVKVKDHFGNYVKP

NKMQVKLQRLAYIRKLQFQMQTNPERVLEWYSHNQTNDLIINNFVDKQNGEKGLVPFFGA

AVAELKDTLPIDRISEMLKQFVELKNLEKQGEEVKSKIDQLIELEPADNLKSGVVANMVG

VIAFLLEKYSYQVYISLEDLSKPFENQIVLGISGVPIGINKGMAGRSVNVEKFAGLGLYN

FFEIQLLKKLFRIQRDSCHILHLVPAFRAMKNYDNVAVGKGKIKNQFGIVFFVDAAATSK

TCPCCGANNSKKFEPDFRKFPNAKKFETPDKKSVWLERDKSDGKDIIRCHVCGFDTSKEY

DDNPRKYIKSGDDNAAYLISAEGVKAYELATTLVDNK

>ca37Cas12a2

MINLENFKNLYEVRKTVRFGLNQPNKKGDFKTHLEFKDFTEKSFQNVENELASNGKYNIG

DEEQLIKKISAFVEQLKIQLGYWKVFYQRYDLIAINKDYYKILARKAKFDAIWEISKSNK

YIKVKQPQASQISLSSLKIGNRSDSIIQYWGEIIEKTDYLLNIFKPKLEQYERAIKDANN

AHIKPDSINFRKTFLQLLKLTKEFLQPLSDRSIIFEFSKKKVSKEIEKISEFAGEKNNTK

IYNVLKNGEELRQYFEANGSQVSYGRVSLNYYTAVQKPNNFDQEIKKSIDDLGIINFLKK

SDSQIIDYLHQGSKQKIKLLLTSKHPYPIELLQLFKVKPIPFSVKYNLAKFVEKNYKSET

NLSYEDILNKFNLLGRAIDIANDFKNSNHKGNFSLDEYPVKLAFDYAWENTARSLKRDIS

FPKKVCERFLKDNFDIDVDNADFKLYANLLFIADNLATIEYNNPNNEAELINEIKQAFEC

MSFSFEKKVYEGHKKAILELLDKEKSQRDYSTISKAKQELGLLRGGLKNKIKKYRDLTQR

LIDKKNSHFGIASFIGKTLATIRDGLKEENELNKISHYGVIIEDNNQDKYILISKLEGKD

RRDKIVQKLGKGDIKVYQVNSFTSKALNKFIKNPLSEDAKKFHGKYKDEYGFGYENKNGD

FTYKVKNVSEYDEQGKWIGYQEEFLNHIKKCLIDSEISREQNWTAFGWSFDGCNNYEEIE

KEIDSKGYQLTENSISKGNLESLVKDEDCSLFPIINQDISSQKQENKNIFTLDFEKVFER

KECRIHPEFSIFYRKPIEEHKKENKSGIINRFGRLQLLANLGIEFVPRNLSFKTKKEQNR

IAIDQKKQNQLVQEFNQEKVNTYFEGLDNYYIFGIDRGIKQLATLCVTDKGGVIQDFTIF

TKHFNQASKIWEYRENRKEGILDLTNLKIESDKNGNTYIVDISLFDAKDDNGRSTGTNKQ

NIQLKQLAYIRKLQYQMSANEKGVLDFIAKYISKDEREQNIKELITPYKEGKKFADLPMD

IFQEMFENYYRLKTDQNLSEFEKKNLMKITTELDASENLKKGVVANMIGVIYYLMKYYDY

KVKITLENLNQSFGGQVDGINNHYVDIKNNFMYQENQALAGVGTYHFFEMQLLKKIFKIS

TEEGILHLVPSFGSVKNYFKDIEKYSLIVNIGRNEKYNQFGVVHFVKPDNTSKKCPVCLN

TQTTGKKEGMPIINRNYKKSNIFYCERCGFQSIHSHCQEENIKDSNDFHYSVEEVERIEM

KNKEAIEKYKKQGKNLHFIKNGDDNAAYNIGEKIRELPKKSDVKNTSISGNISFS

>ca38Cas12a2

MVEFCTGVKGMKSFENSCQISKTLRFGATLKEDKKKCKSHEELQEFVDISYKTMKSSATI

AESLNEIELVKKCERCYSEIIKFHEAWGQIYCRTDQIAVYKDFYRQLARKARFDVGDQNS

QLITLSSLSSKHNVYQGLKRSQHITNYWKDNIARQKSFLKDFSQQLHQYKRALENSDKAH

TKPNLINFNKTFSILANLVNEIVIPLSNGAISFPNISKLEDGEESRHLIEFALNDYSDLV

GSIGELKDAIATNGGYTPFAKVTLNHYTAEQKPHVFKNDIDAKIRELKLIELVEKLKNKT

SKQIEEYFSKFDKFKIYDDRNQSVIIRTQCFKYKPIPFLVKHELAKYIAEDEDAVLKVLD

AIGATRSPAHDYAHNEDVFDLKHYPIKVAFDYAWEQLANGVYTTVSFPEEKCREYLNAIY

GCEVTKEPVFKFYADLLYIRKNLAVLEHKNNLPSNPEEFICKIENTFEKIVLPYKIKEFE

TYKKAILTWINDGRGHEKYTDSKRELGLIRGGLKGRIGAQKMFRKNKKGELIPYYENPYT

KLTNEFKNISSSYGKTFAELRDKFKEKSEITKITHFGIIIEDNNKDRYLLANGLQHDNTD

QSNTQVEAILGKLNTSLEFTTYQVKSLTSKILIKLIKNHTTSPNAKSPYADFHTSKIHVD

WTKIKKEWDTYKSNHSLLQYVKDCLTNSTMAKNQNWAEFGWDLESCNSYEAIEHEIDQKS

YILQKHTISKASIKSLVENGCLLLPIVNQDITSQGRKDKNQFSKDWKQIFEDSKEYRLHP

EFAVSYRTPIKDYPKDKRYGRLQFICAFNCEIIPQNGEFINLKKQIENFNDEDIQKNNVA

EFNNKVNEALLGKEYVVIGIDRGLKQLATLCVLNKRGKILGDFEIYKKEFVRTENRSKNY

WKHTLAETRHILDLSNLRVETTVEDNKVLVDQSLTLVKKNRDTPDEKATEENRQKIKLKQ

LSYIRKLQYAMQTNEQAVLDLLKDNSDDEEFKKRVEGVISPFGEGQEYADLPIDTMRAMI

QDLQGVIAKGNKQTEKNKIIELDAADSLKQGVVANMIGVVNFILAKFNYEAYVSLEDLSR

AYNVAKSGYDGRYLPSTSQDPDMDFKEQQNQMLAGLGTYQFFEIQLLKKLQKIQSNNTVL

RFVPAFRSADNYRNIIKLNPKYDNTEYVHKPFGIVHFIDPKDTSSKCPVCGKTGKKNVDR

NEKKNNILLCKACAFRTVWEFKQPENIKNEKHCFSEDGIKKTAEDNNKKIKEKNLSDKNL

HYIHNGDDNGAYHIALKSVENLKHKK

>ca41Cas12a2

MLTKFANLYELTKTVRFGLTPKNSYKKVSDFFELTEKSLESVQAEIVEREKEKINITSTE

ESIKKIRAYVDELKKQSAQWKTIYQREDMIAVTKEYYKKLEKEAGFDGFWEDRGKKQPQT

SEIMLSALSKEYNNKPRREYIITYWALLLQKTEQLNSYFEPLLEKYEESLSRKEQAHLKP

NLVDFRKQFLSLLNVSNKWLMPVINQSIVFPKIHNASKGEQNRKVKDFISEEGRQKRIQL

QSIGRSLRDFFEANGSRVPFGKATLNYFTARQKPNRFDSEIDALMLDLEIEEIVRKVKDL

NGEELKNYFKYQTYATGDKLDAFLGHSYMTLIEKIQLFKPKPIPASVRFSLAERLSKKMN

LPMEKVTAIFDEIGNPIDIAREYEQALDQKNFDLNKYPINIAFDYAWESCASLIKGRISE

GDFPKAQCLAILKKFNADGDDFKLYANLRYIRDELAPIEHNNPSADIEKELVYNIQKTIG

IIYSSMPSEYQKYLDIILKWVEMPKIERNSSDSNFIFAKQKIGLLRGGLKNKINKHAEIT

NKFKNLSMAFGRSCSDLRDKLREEHELSKIKYYSVLIEDSKKNRHLLMLPLETGDQKMSA

VEIENRLDILENKFVEDAPSANAYIASKVSSFTSKALVKMLKNPKGSEDFHIGSEKYKLH

DYNNKIKKEWKIYQDDSKFLSYAKKCLSESKMAMNQHWEKFGWNFEGCKTYADLEKEVDI

KGYSLEKKYISFENTEMLVYEQGCLLFPIVNQDYASEVCQNIFTGNKNAFTIEIEKGLQE

QDGYLIHPEFTIFYQKPTEDYKKSNRYGRFQLSANFSLEVKKISDNFKTKKEKLKNIRDK

DSFQKDVQIFNESINAQLRINEDVYFYGIDRGINELATLCLLNNKNKIQDFIVFKKEKKK

RNSENHLREAGDFYTYVDFEKQNILDLTNIKVETWNMRDEIHPNSNFLKQAFKKIEADPE

MKMIATQLGYGINEKKQCTILVEYPSDRSNTLKLRMNHFQRALQFATSHSEKRDIVGGIA

ERNFEGTDLENPSDEVKKQIIQDLFGLGSDFLIDEKKAGVINELQTKKFRFEKIYSRNQQ

LFLEMFEPLIQQLIEWYQHKNDPDYEKTGKAKYEEMEDLDTLKRGVTANMVGVFSFLMQK

FPGVIVLENIAKEKLEKPVQSKLDEHTREVESHQEAEHHRWAGVELYRYMEKKLVQKFSN

WRQPTGEIFNLTPPFANLESLNKQTMTINSERIENMFQFGIFCYTPAEYTTKTCPCCNKY

HIRNRKKGNTFDSIICKNEHCGFDTRYNEPLHESFNPARNVEIALRLNPEFSEALPKIRS

GDQSAAFNIAKKGLIIMENKLC

>ca125Cas12a2

MGKKQAVYNKFDEYLNQKNNSGKSNSNQIFNNNRQRRDKQAKNPKNNHAGKGDDAKPQII

KIPKSAIFEQYKTIRFTLSPNETKKNKVVASELESLVGVSMEMVQSQIQEKSKSLEIKDQ

ETAIKKINEILGSLKNFTSSWLTFSERTDTIKLSEEYYRFVARKARFNSFEQYKDKYGKE

KTTPQIKSGFIPLNQKSEYQGIERSEYVIKYWQNLANNIVNLHSQLEAPVDKYITALEKQ

DKAHTKVNLIQLKKTFLSFSNLILEYLEPLTNNKILIEKIDKLPDSEENTKLKEFFSNQN

INLIRQDLQSLKQVIDYFNQNGALVSLGKVSLNYHTAVKKPDIVKSEIETIINELELETF

IKKYIQCDDQELFKKFQFDIADKKQVFLCNKNDLTKIELAQLFKPRPIPFGVIHEISEYF

EKKDFDYNNVQNILMNTGQSVNIAGDWTNTKLEDRENFDLNKYPLKSAFDYAWEYTARNK

VGLIGNNDFAKKQSEKLLTDFDVKTSDNNFESYANLLFIGDKLAVLEHSAHEIKDKKELE

NIMILISQKIENVKNPFHQNSEKDRYEKFENNCETISSSLNSSHTLKSNKNFQKAKQNLG

LVRGGLKNKGIYYKLTQQFGINKTIDNKLSLASFLGLKFADLNKKFKEKYENNKIGYYGV

IVEENINKYLLLKALENEDTREIIENDKILKSEKFENALTVKKVTSLTSNSLKKIRINKK

AYPDFHLEKIDEEKTSKEDNKALKESKRANYIKRCLLKSKMSQEQNWNQKFNWENDLNEC

QTYEQIEKVLDLKGYKLETKHISKNDLENLVTEQDCLLLPIINQDYQSEIKQGEFVGNKN

QFTVDFENVFVNKPIENKQDKKTYNYRIHPEFSLFYQLPTIKPETGQTEQDLQKEHHAKT

LNRNSRFQIIGNFGIEIKPEITEFSQFGNFINRKGKNDLTRDKQSYAEFVQGFNQKISSD

FEQKDSNRQWIYGFDRGINELATLCVLHKNSKQIDDFVVFKKEKEKEKKLGWIGEIIEEK

KKSGKELEKKDKFVFRFVKHDESKNQKGFELQHILDLTNIKVETWHFGFKDKDGNDHPND

KYLKTALSKIKENSELFEIAKKLGYFDNFSEGKRGENIKILVQYPENESNVIKLRMNHFQ

RVLSFAISQNRDGVANSIEEVFESDVEAEQVKKLFDNGILDNTKIKIDIDKHINLFKPLL

DQIKEWNENKDKPNYLDEFKKKYEEIEDLDTLKKGIVANMVGVTGFLMEKLPGVLVLEDT

YKYDPQGFRINSLTKRDENADTFGHSEHLTWAGTETYRYFEKMLVKKFVKQRLVPPFADL

ESLNTQKTKLGGNEDENNKQFGIMFYVDAGFTSKTCPCCGFKPDFKNENLKNLLEKTTSI

QKINHNYCLSDCKDNILFELKN

>EcCas12a

MIKDNFVNVYSLSKTIRMALIPWGKTEDNFYKKFLLEEDEERAKNYIKVKGYMDEYHKNF

IESALNSVVLNGVDEYCELYFKQNKSDSEVKKIESLEASMRKQISKAMKEYTVDGVKIYP

LLSKKEFIRELLPEFLTQDEEIETLEQFNDFSTYFQGFWENRKNIYTDEEKSTGVPYRCI

NDNLPKFLDNVKSFEKVILALPQKAVDELNANFNGVYNVDVQDVFSVDYFNFVLSQSGIE

KYNNIIGGYSNSDASKVQGLNEKINLYNQQIAKSDKSKKLPLLKPLYKQILSDRSSLSFI

PEKFKDDNEVLNSINVLYDNIAESLEKANDLMSDIANYNTDNIFISSGVAVTDISKKVFG

DWSLIRNNWNDEYESTHKKGKNEEKFYEKEDKEFKKIKSFSVSELQRLANSDLSIVDYLV

DESASLYADIKTAYNNAKDLLSNEYSHSKRLSKNDDAIELIKSFLDSIKNYEAFLKPLCG

TGKEESKDNAFYGAFLECFEEIRQVDAVYNKVRNHITQKPYSNDKIKLNFQNPQFLAGWD

KNKERAYRSVLLRNGEKYYLAIMEKGKSKLFEDFPEDESSPFEKIDYKLLPEPSKMLPKV

FFATSNKDLFNPSDEILNIRATGSFKKGDSFNLDDCHKFIDFYKASIENHPDWSKFDFDF

SETNDYEDISKFFKEVSDQGYSIGYRKISESYLEEMVDNGSLYMFQLYNKDFSENRKSKG

TPNLHTLYFKMLFDERNLEDVVYKLSGGAEMFYRKPSIDKNEMIVHPKNQPIDNKNPNNV

KKTSTFEYDIVKDMRYTKPQFQLHLPIVLNFKANSKGYINDDVRNVLKNSEDTYVIGIDR

GERNLVYACVVDGNGKLVEQVPLNVIEADNGYKTDYHKLLNDREEKRNEARKSWKTIGNI

KELKEGYISQVVHKICQLVVKYDAVIAMEDLNSGFVNSRKKVEKQVYQKFERMLTQKLNY

LVDKKLDPNEMGGLLNAYQLTNEATKVRNGRQDGIIFYIPAWLTSKIDPTTGFVNLLKPK

YNSVSASKEFFSKFDEIRYNEKENYFEFSFNYDNFPKCNADFKREWTVCTYGDRIRTFRD

PENNNKFNSEVVVLNDEFKNLFVEFDIDYTDNLKEQILAMDEKSFYKKLMGLLSLTLQMR

NSISKNVDVDYLISPVKNSNGEFYDSRNYDITSSLPCDADSNGAYNIARKGLWAINQIKQ

ADDETKANISIKNSEWLQYAQNCDEV

>FnCas12a

MSIYQEFVNKYSLSKTLRFELIPQGKTLENIKARGLILDDEKRAKDYKKAKQIIDKYHQF

FIEEILSSVCISEDLLQNYSDVYFKLKKSDDDNLQKDFKSAKDTIKKQISEYIKDSEKFK

NLFNQNLIDAKKGQESDLILWLKQSKDNGIELFKANSDITDIDEALEIIKSFKGWTTYFK

GFHENRKNVYSSNDIPTSIIYRIVDDNLPKFLENKAKYESLKDKAPEAINYEQIKKDLAE

ELTFDIDYKTSEVNQRVFSLDEVFEIANFNNYLNQSGITKFNTIIGGKFVNGENTKRKGI

NEYINLYSQQINDKTLKKYKMSVLFKQILSDTESKSFVIDKLEDDSDVVTTMQSFYEQIA

AFKTVEEKSIKETLSLLFDDLKAQKLDLSKIYFKNDKSLTDLSQQVFDDYSVIGTAVLEY

ITQQIAPKNLDNPSKKEQELIAKKTEKAKYLSLETIKLALEEFNKHRDIDKQCRFEEILA

NFAAIPMIFDEIAQNKDNLAQISIKYQNQGKKDLLQASAEDDVKAIKDLLDQTNNLLHKL

KIFHISQSEDKANILDKDEHFYLVFEECYFELANIVPLYNKIRNYITQKPYSDEKFKLNF

ENSTLANGWDKNKEPDNTAILFIKDDKYYLGVMNKKNNKIFDDKAIKENKGEGYKKIVYK

LLPGANKMLPKVFFSAKSIKFYNPSEDILRIRNHSTHTKNGSPQKGYEKFEFNIEDCRKF

IDFYKQSISKHPEWKDFGFRFSDTQRYNSIDEFYREVENQGYKLTFENISESYIDSVVNQ

GKLYLFQIYNKDFSAYSKGRPNLHTLYWKALFDERNLQDVVYKLNGEAELFYRKQSIPKK

ITHPAKEAIANKNKDNPKKESVFEYDLIKDKRFTEDKFFFHCPITINFKSSGANKFNDEI

NLLLKEKANDVHILSIDRGERHLAYYTLVDGKGNIIKQDTFNIIGNDRMKTNYHDKLAAI

EKDRDSARKDWKKINNIKEMKEGYLSQVVHEIAKLVIEYNAIVVFEDLNFGFKRGRFKVE

KQVYQKLEKMLIEKLNYLVFKDNEFDKTGGVLRAYQLTAPFETFKKMGKQTGIIYYVPAG

FTSKICPVTGFVNQLYPKYESVSKSQEFFSKFDKICYNLDKGYFEFSFDYKNFGDKAAKG

KWTIASFGSRLINFRNSDKNHNWDTREVYPTKELEKLLKDYSIEYGHGECIKAAICGESD

KKFFAKLTSVLNTILQMRNSKTGTELDYLISPVADVNGNFFDSRQAPKNMPQDADANGAY

HIGLKGLMLLGRIKNNQEGKKLNLVIKNEEYFEFVQNRNN

>KaCas12a

MNVKKSPTTFAEFTNQYSLSKTLRFELKPVGNTQKMLEDFGIIEKDEIIAQKYKITKKYF

DKLHRTFVEEALFGKKIEGLDNYFELYRNWKKDKKANQKDLQNKETELRKNIVTLFNKMA

KTWSQKETFKGLKNNNIEILFEEAVFKSVLKVLYGDEDESFLRDKVSGEFLLDKDGNKIS

VFDSWKGFTGYFTKFNETRKNFYKDDGTATAHATRIIDQNLKRFCENIIIFESIKDKLDL

SEVENVFDISLVDIFSVTNYGDCFLQAGIDSYNTVLGGETLPNGEKKKGLNELINLYKQQ

TREKLPFFKSLDKQILSEKDKFTDEIEDAGALKEVLTVFLKNVDVKIQKFSLLITDVLNN

PDAYDLSQIYLAKEALNTISRRWLVSFSTFEDALYTVLKNAKIISSSAKKTEGGYSFPEF

IPLSYLKEALEKMVEDKIWKEQFYVTVEGLKNNGVIIGNKPVWQQFWEIYRFEFEQLKSR

IEIDSQIGTTHPAGYSVSKTALETILKDFKEDDVAPKLAIKNLADDVLHIYQLAKYFAIE

KKIEKKRRWEGDNFELDVFYTNPDFGYFETFYKNAYEDIVQVYNKLRNYLTKKPYSQEKW

KLNFDNPTLAAGWDKNKEADNSAVLLKKDGKYYLGLMEKSHNRIFDKKHEALFLDNNQSE

NYEKVVYKFFPDQAKMFPKVCFSVKGLDFFKPSEEIYKIYKNAEFKKGDTFSIKSMQSLI

KFYIDCLQKYDGWKGYNFKHIKSSNQYTNNIGDFFRDVAEDGYKITYQDVSDSYIKEKNN

NGELYLFQIKNQDWNLDKAINGIKKTTTKNLHTLYFEALFSKENAEQNFPIKLNGQAEIF

YRPKTSQGKLGTKLVNNKEVVNHKRYSEDKIFFHVPLTLNRTKNRTKNEAQKFNSKVNEF

LVNNPNINIIGIDRGEKHLAYYSVINQQGEILSDAKGIPAIGSLNFVGQGADGKEIDYAE

KLAIRASERESSRRDWLDIEKIKDLKKGYISQVVRKLADLAIEHNAIIVLEDLNMRFKQI

RGGIEKSIYQQLEKALIEKLSFLVDKGETDPEKAGHLFRAYQLTAPFESFQNIGKQTGII

FYTQANYTSKTCPECGFRPNVRFDESIDWNLINIKYYDNDFTISYSLKTFTKAKETSKRS

NQLFTGINKKDSFILSTKNAIRYKWFRRNISAVELSAGESKLINETGSGVTIQYQLTDCM

KGLFEGNGIDIKGNINSQLRTGGHLMKFYKQLANYLYLLSNTRSSVSGTDIDIINCPCCG

FDSTQGFKGVAYNGDANGAYNIGRKGLIVLDKIKNYQKNNGPLEKMNWGDIFIDIDEWDK

FTQK

>Lb2Cas12a

MHENNGKIADNFIGIYPVSKTLRFELKPVGKTQEYIEKHGILDEDLKRAGDYKSVKKIID

AYHKYFIDEALNGIQLDGLKNYYELYEKKRDNNEEKEFQKIQMSLRKQIVKRFSEHPQYK

YLFKKELIKNVLPEFTKDNAEEQTLVKSFQEFTTYFEGFHQNRKNMYSDEEKSTAIAYRV

VHQNLPKYIDNMRIFSMILNTDIRSDLTELFNNLKTKMDITIVEEYFAIDGFNKVVNQKG

IDVYNTILGAFSTDDNTKIKGLNEYINLYNQKNKAKLPKLKPLFKQILSDRDKISFIPEQ

FDSDTEVLEAVDMFYNRLLQFVIENEGQITISKLLTNFSAYDLNKIYVKNDTTISAISND

LFDDWSYISKAVRENYDSENVDKNKRAAAYEEKKEKALSKIKMYSIEELNFFVKKYSCNE

CHIEGYFERRILEILDKMRYAYESCKILHDKGLINNISLCQDRQAISELKDFLDSIKEVQ

WLLKPLMIGQEQADKEEAFYTELLRIWEELEPITLLYNKVRNYVTKKPYTLEKVKLNFYK

STLLDGWDKNKEKDNLGIILLKDGQYYLGIMNRRNNKIADDAPLAKTDNVYRKMEYKLLT

KVSANLPRIFLKDKYNPSEEMLEKYEKGTHLKGENFCIDDCRELIDFFKKGIKQYEDWGQ

FDFKFSDTESYDDISAFYKEVEHQGYKITFRDIDETYIDSLVNEGKLYLFQIYNKDFSPY

SKGTKNLHTLYWEMLFSQQNLQNIVYKLNGNAEIFYRKASINQKDVVVHKADLPIKNKDP

QNSKKESMFDYDIIKDKRFTCDKYQFHVPITMNFKALGENHFNRKVNRLIHDAENMHIIG

IDRGERNLIYLCMIDMKGNIVKQISLNEIISYDKNKLEHKRNYHQLLKTREDENKSARQS

WQTIHTIKELKEGYLSQVIHVITDLMVEYNAIVVLEDLNFGFKQGRQKFERQVYQKFEKM

LIDKLNYLVDKSKGMDEDGGLLHAYQLTDEFKSFKQLGKQSGFLYYIPAWNTSKLDPTTG

FVNLFYTKYESVEKSKEFINNFTSILYNQEREYFEFLFDYSAFTSKAEGSRLKWTVCSKG

ERVETYRNPKKNNEWDTQKIDLTFELKKLFNDYSISLLDGDLREQMGKIDKADFYKKFMK

LFALIVQMRNSDEREDKLISPVLNKYGAFFETGKNERMPLDADANGAYNIARKGLWIIEK

IKNTDVEQLDKVKLTISNKEWLQYAQEHIL

>LbCas12a

AASKLEKFTNCYSLSKTLRFKAIPVGKTQENIDNKRLLVEDEKRAEDYKGVKKLLDRYYL

SFINDVLHSIKLKNLNNYISLFRKKTRTEKENKELENLEINLRKEIAKAFKGAAGYKSLF

KKDIIETILPEAADDKDEIALVNSFNGFTTAFTGFFDNRENMFSEEAKSTSIAFRCINEN

LTRYISNMDIFEKVDAIFDKHEVQEIKEKILNSDYDVEDFFEGEFFNFVLTQEGIDVYNA

IIGGFVTESGEKIKGLNEYINLYNAKTKQALPKFKPLYKQVLSDRESLSFYGEGYTSDEE

VLEVFRNTLNKNSEIFSSIKKLEKLFKNFDEYSSAGIFVKNGPAISTISKDIFGEWNLIR

DKWNAEYDDIHLKKKAVVTEKYEDDRRKSFKKIGSFSLEQLQEYADADLSVVEKLKEIII

QKVDEIYKVYGSSEKLFDADFVLEKSLKKNDAVVAIMKDLLDSVKSFENYIKAFFGEGKE

TNRDESFYGDFVLAYDILLKVDHIYDAIRNYVTQKPYSKDKFKLYFQNPQFMGGWDKDKE

TDYRATILRYGSKYYLAIMDKKYAKCLQKIDKDDVNGNYEKINYKLLPGPNKMLPKVFFS

KKWMAYYNPSEDIQKIYKNGTFKKGDMFNLNDCHKLIDFFKDSISRYPKWSNAYDFNFSE

TEKYKDIAGFYREVEEQGYKVSFESASKKEVDKLVEEGKLYMFQIYNKDFSDKSHGTPNL

HTMYFKLLFDENNHGQIRLSGGAELFMRRASLKKEELVVHPANSPIANKNPDNPKKTTTL

SYDVYKDKRFSEDQYELHIPIAINKCPKNIFKINTEVRVLLKHDDNPYVIGIDRGERNLL

YIVVVDGKGNIVEQYSLNEIINNFNGIRIKTDYHSLLDKKEKERFEARQNWTSIENIKEL

KAGYISQVVHKICELVEKYDAVIALEDLNSGFKNSRVKVEKQVYQKFEKMLIDKLNYMVD

KKSNPCATGGALKGYQITNKFESFKSMSTQNGFIFYIPAWLTSKIDPSTGFVNLLKTKYT

SIADSKKFISSFDRIMYVPEEDLFEFALDYKNFSRTDADYIKKWKLYSYGNRIRIFAAAK

KNNVFAWEEVCLTSAYKELFNKYGINYQQGDIRALLCEQSDKAFYSSFMALMSLMLQMRN

SITGRTDVDFLISPVKNSDGIFYDSRNYEAQENAILPKNADANGAYNIARKVLWAIGQFK

KAEDEKLDKVKIAISNKEWLEYAQTSVK

>LsaCas12b1

MSIRSFKLKLKTKSGVNAEQLRRGLWRTHQLINDGIAYYMNWLVLLRQEDLFIRNKETNE

IEKRSKEEIQAVLLERVHKQQQRNQWSGEVDEQTLLQALRQLYEEIVPSVIGKSGNASLK

ARFFLGPLVDPNNKTTKDVSKSGPTPKWKKMKDAGDPNWVQEYEKYMAERQTLVRLEEMG

LIPLFPMYTDEVGDIHWLPQASGYTRTWDRDMFQQAIERLLSWESWNRRVRERRAQFEKK

THDFASRFSESDVQWMNKLREYEAQQEKSLEENAFAPNEPYALTKKALRGWERVYHSWMR

LDSAASEEAYWQEVATCQTAMRGEFGDPAIYQFLAQKENHDIWRGYPERVIDFAELNHLQ

RELRRAKEDATFTLPDSVDHPLWVRYEAPGGTNIHGYDLVQDTKRNLTLILDKFILPDEN

GSWHEVKKVPFSLAKSKQFHRQVWLQEEQKQKKREVVFYDYSTNLPHLGTLAGAKLQWDR

NFLNKRTQQQIEETGEIGKVFFNISVDVRPAVEVKNGRLQNGLGKALTVLTHPDGTKIVT

GWKAEQLEKWVGESGRVSSLGLDSLSEGLRVMSIDLGQRTSATVSVFEITKEAPDNPYKF

FYQLEGTEMFAVHQRSFLLALPGENPPQKIKQMREIRWKERNRIKQQVDQLSAILRLHKK

VNEDERIQAIDKLLQKVASWQLNEEIATAWNQALSQLYSKAKENDLQWNQAIKNAHHQLE

PVVGKQISLWRKDLSTGRQGIAGLSLWSIEELEATKKLLTRWSKRSREPGVVKRIERFET

FAKQIQHHINQVKENRLKQLANLIVMTALGYKYDQEQKKWIEVYPACQVVLFENLRSYRF

SFERSRRENKKLMEWSHRSIPKLVQMQGELFGLQVADVYAAYSSRYHGRTGAPGIRCHAL

TEADLRNETNIIHELIEAGFIKEEHRPYLQQGDLVPWSGGELFATLQKPYDNPRILTLHA

DINAAQNIQKRFWHPSMWFRVNCESVMEGEIVTYVPKNKTVHKKQGKTFRFVKVEGSDVY

EWAKWSKNRNKNTFSSITERKPPSSMILFRDPSGTFFKEQEWVEQKTFWGKVQSMIQAYM

KKTIVQRMEE

>LseCas12b1

MSIRSFKLKIKTKSGVNAEELRRGLWRTHQLINDGIAYYMNWLVLLRQEDLFIRNEETNE

IEKRSKEEIQGELLERVHKQQQRNQWSGEVDDQTLLQTLRHLYEEIVPSVIGKSGNASLK

ARFFLGPLVDPNNKTTKDVSKSGPTPKWKKMKDAGDPNWVQEYEKYMAERQTLVRLEEMG

LIPLFPMYTDEVGDIHWLPQASGYTRTWDRDMFQQAIERLLSWESWNRRVRERRAQFEKK

THDFASRFSESDVQWMNKLREYEAQQEKSLEENAFAPNEPYALTKKALRGWERVYHSWMR

LDSAASEEAYWQEVATCQTAMRGEFGDPAIYQFLAQKENHDIWRGYPERVIDFAELNHLQ

RELRRAKEDATFTLPDSVDHPLWVRYEAPGGTNIHGYDLVQDTKRNLTLILDKFILPDEN

GSWHEVKKVPFSLAKSKQFHRQVWLQEEQKQKKREVVFYDYSTNLPHLGTLAGAKLQWDR

NFLNKRTQQQIEETGEIGKVFFNISVDVRPAVEVKNGRLQNGLGKALTVLTHPDGTKIVT

GWKAEQLEKWVGESGRVSSLGLDSLSEGLRVMSIDLGQRTSATVSVFEITKEAPDNPYKF

FYQLEGTELFAVHQRSFLLALPGENPPQKIKQMREIRWKERNRIKQQVDQLSAILRLHKK

VNEDERIQAIDKLLQKVASWQLNEEIATAWNQALSQLYSKAKENDLQWNQAIKNAHHQLE

PVVGKQISLWRKDLSTGRQGIAGLSLWSIEELEATKKLLTRWSKRSREPGVVKRIERFET

FAKQIQHHINQVKENRLKQLANLIVMTALGYKYDQEQKKWIEVYPACQVVLFENLRSYRF

SYERSRRENKKLMEWSHRSIPKLVQMQGELFGLQVADVYAAYSSRYHGRTGAPGIRCHAL

TEADLRNETNIIHELIEAGFIKEEHRPYLQQGDLVPWSGGELFATLQKPYDNPRILTLHA

DINAAQNIQKRFWHPSMWFRVNCESVMEGEIVTYVPKNKTVHKKQGKTFRFVKVEGSDVY

EWAKWSKNRNKNTFSSITERKPPSSMILFRDPSGTFFKEQEWVEQKTFWGKVQSMIQAYM

KKTIVQRMEE

>MaCas12a

MDAKEFTGQYPLSKTLRFELRPIGRTWDNLEASGYLAEDRHRAECYPRAKELLDDNHRAF

LNRVLPQIDMDWHPIAEAFCKVHKNPGNKELAQDYNLQLSKRRKEISAYLQDADGYKGLF

AKPALDEAMKIAKENGNESDIEVLEAFNGFSVYFTGYHESRENIYSDEDMVSVAYRITED

NFPRFVSNALIFDKLNESHPDIISEVSGNLGVDDIGKYFDVSNYNNFLSQAGIDDYNHII

GGHTTEDGLIQAFNVVLNLRHQKDPGFEKIQFKQLYKQILSVRTSKSYIPKQFDNSKEMV

DCICDYVSKIEKSETVERALKLVRNISSFDLRGIFVNKKNLRILSNKLIGDWDAIETALM

HSSSSENDKKSVYDSAEAFTLDDIFSSVKKFSDASAEDIGNRAEDICRVISETAPFINDL

RAVDLDSLNDDGYEAAVSKIRESLEPYMDLFHELEIFSVGDEFPKCAAFYSELEEVSEQL

IEIIPLFNKARSFCTRKRYSTDKIKVNLKFPTLADGWDLNKERDNKAAILRKDGKYYLAI

LDMKKDLSSIRTSDEDESSFEKMEYKLLPSPVKMLPKIFVKSKAAKEKYGLTDRMLECYD

KGMHKSGSAFDLGFCHELIDYYKRCIAEYPGWDVFDFKFRETSDYGSMKEFNEDVAGAGY

YMSLRKIPCSEVYRLLDEKSIYLFQIYNKDYSENAHGNKNMHTMYWEGLFSPQNLESPVF

KLSGGAELFFRKSSIPNDAKTVHPKGSVLVPRNDVNGRRIPDSIYRELTRYFNRGDCRIS

DEAKSYLDKVKTKKADHDIVKDRRFTVDKMMFHVPIAMNFKAISKPNLNKKVIDGIIDDQ

DLKIIGIDRGERNLIYVTMVDRKGNILYQDSLNILNGYDYRKALDVREYDNKEARRNWTK

VEGIRKMKEGYLSLAVSKLADMIIENNAIIVMEDLNHGFKAGRSKIEKQVYQKFESMLIN

KLGYMVLKDKSIDQSGGALHGYQLANHVTTLASVGKQCGVIFYIPAAFTSKIDPTTGFAD

LFALSNVKNVASMREFFSKMKSVIYDKAEGKFAFTFDYLDYNVKSECGRTLWTVYTVGER

FTYSRVNREYVRKVPTDIIYDALQKAGISVEGDLRDRIAESDGDTLKSIFYAFKYALDMR

VENREEDYIQSPVKNASGEFFCSKNAGKSLPQDSDANGAYNIALKGILQLRMLSEQYDPN

AESIRLPLITNKAWLTFMQSGMKTWKN

>Mb1Cas12a

MLFQDFTHLYPLSKTVRFELKPIDRTLEHIHAKNFLSQDETMADMHQKVKVILDDYHRDF

IADMMGEVKLTKLAEFYDVYLKFRKNPKDDELQKQLKDLQAVLRKEIVKPIGNGGKYKAG

YDRLFGAKLFKDGKELGDLAKFVIAQEGESSPKLAHLAHFEKFSTYFTGFHDNRKNMYSD

EDKHTAIAYRLIHENLPRFIDNLQILTTIKQKHSALYDQIINELTASGLDVSLASHLDGY

HKLLTQEGITAYNTLLGGISGEAGSPKIQGINELINSHHNQHCHKSERIAKLRPLHKQIL

SDGMSVSFLPSKFADDSEMCQAVNEFYRHYADVFAKVQSLFDGFDDHQKDGIYVEHKNLN

ELSKQAFGDFALLGRVLDGYYVDVVNPEFNERFAKAKTDNAKAKLTKEKDKFIKGVHSLA

SLEQAIEHYTARHDDESVQAGKLGQYFKHGLAGVDNPIQKIHNNHSTIKGFLERERPAGE

RALPKIKSGKNPEMTQLRQLKELLDNALNVAHFAKLLTTKTTLDNQDGNFYGEFGVLYDE

LAKIPTLYNKVRDYLSQKPFSTEKYKLNFGNPTLLNGWDLNKEKDNFGVILQKDGCYYLA

LLDKAHKKVFDNAPNTGKSIYQKMIYKYLEVRKQFPKVFFSKEAIAINYHPSKELVEIKD

KGRQRSDDERLKLYRFILECLKIHPKYDKKFEGAIGDIQLFKKDKKGREVPISEKDLFDK

INGIFSSKPKLEMEDFFIGEFKRYNPSQDLVDQYNIYKKIDSNDNRKKENFYNNHPKFKK

DLVRYYYESMCKHEEWEESFEFSKKLQDIGCYVDVNELFTEIETRRLNYKISFCNINADY

IDELVEQGQLYLFQIYNKDFSPKAHGKPNLHTLYFKALFSEDNLADPIYKLNGEAQIFYR

KASLDMNETTIHRAGEVLENKNPDNPKKRQFVYDIIKDKRYTQDKFMLHVPITMNFGVQG

MTIKEFNKKVNQSIQQYDEVNVIGIDRGERHLLYLTVINSKGEILEQCSLNDITTASANG

TQMTTPYHKILDKREIERLNARVGWGEIETIKELKSGYLSHVVHQISQLMLKYNAIVVLE

DLNFGFKRGRFKVEKQIYQNFENALIKKLNHLVLKDKADDEIGSYKNALQLTNNFTDLKS

IGKQTGFLFYVPAWNTSKIDPETGFVDLLKPRYENIAQSQAFFGKFDKICYNADKDYFEF

HIDYAKFTDKAKNSRQIWTICSHGDKRYVYDKTANQNKGAAKGINVNDELKSLFARHHIN

EKQPNLVMDICQNNDKEFHKSLMYLLKTLLALRYSNASSDEDFILSPVANDEGVFFNSAL

ADDTQPQNADANGAYHIALKGLWLLNELKNSDDLNKVKLAIDNQTWLNFAQNR

>Mb2Cas12a

MLFQDFTHLYPLSKTVRFELKPIGKTLEHIHAKNFLNQDETMADMYQKVKAILDDYHRDF

IADMMGEVKLTKLAEFYDVYLKFRKNPKDDGLQKQLKDLQAVLRKEIVKPIGNGGKYKAG

YDRLFGAKLFKDGKELGDLAKFVIAQEGESSPKLAHLAHFEKFSTYFTGFHDNRKNMYSD

EDKHTAIAYRLIHENLPRFIDNLQILATIKQKHSALYDQIINELTASGLDVSLASHLDGY

HKLLTQEGITAYNTLLGGISGEAGSRKIQGINELINSHHNQHCHKSERIAKLRPLHKQIL

SDGMGVSFLPSKFADDSEVCQAVNEFYRHYADVFAKVQSLFDGFDDYQKDGIYVEYKNLN

ELSKQAFGDFALLGRVLDGYYVDVVNPEFNERFAKAKTDNAKAKLTKEKDKFIKGVHSLA

SLEQAIEHYTARHDDESVQAGKLGQYFKHGLAGVDNPIQKIHNNHSTIKGFLERERPAGE

RALPKIKSDKSPEIRQLKELLDNALNVAHFAKLLTTKTTLHNQDGNFYGEFGALYDELAK

IATLYNKVRDYLSQKPFSTEKYKLNFGNPTLLNGWDLNKEKDNFGVILQKDGCYYLALLD

KAHKKVFDNAPNTGKSVYQKMIYKLLPGPNKMLPKVFFAKSNLDYYNPSAELLDKYAQGT

HKKGDNFNLKDCHALIDFFKAGINKHPEWQHFGFKFSPTSSYQDLSDFYREVEPQGYQVK

FVDINADYINELVEQGQLYLFQIYNKDFSPKAHGKPNLHTLYFKALFSEDNLVNPIYKLN

GEAEIFYRKASLDMNETTIHRAGEVLENKNPDNPKKRQFVYDIIKDKRYTQDKFMLHVPI

TMNFGVQGMTIKEFNKKVNQSIQQYDEVNVIGIDRGERHLLYLTVINSKGEILEQRSLND

ITTASANGTQMTTPYHKILDKREIERLNARVGWGEIETIKELKSGYLSHVVHQISQLMLK

YNAIVVLEDLNFGFKRGCFKVEKQIYQNFENALIKKLNHLVLKDKADDEIGSYKNALQLT

NNFTDLKSIGKQTGFLFYVPAWNTSKIDPETGFVDLLKPRYENIAQSQAFFGKFDKICYN

ADRGYFEFHIDYAKFNDKAKNSRQIWKICSHGDKRYVYDKTANQNKGATIGVNVNDELKS

LFTRYHINDKQPNLVMDICQNNDKEFHKSLMYLLKTLLALRYSNASSDEDFILSPVANDE

GVFFNSALADDTQPQNADANGAYHIALKGLWLLNELKNSDDLNKVKLAIDNQTWLNFAQN

R

>Mb3Cas12a

MLFQDFTHLYPLSKTVRFELKPIGRTLEHIHAKNFLSQDETMADMYQKVKVILDDYHRDF

IADMMGEVKLTKLAEFYDVYLKFRKNPKDDELQKQLKDLQAVLRKESVKPIGNGGKYKAG

HDRLFGAKLFKDGKELGDLAKFVIAQEGKSSPKLAHLAHFEKFSTYFTGFHDNRKNMYSD

EDKHTAIAYRLIHENLPRFIDNLQILTTIKQKHSALYDQIINELTASGLDVSLASHLDGY

HKLLTQEGITAYNRIIGEVNGYTNKHNQICHKSERIAKLRPLHKQILSDGMGVSFLPSKF

ADDSEMCQAVNEFYRHYTDVFAKVQSLFDGFDDHQKDGIYVEHKNLNELSKQAFGDFALL

GRVLDGYYVDVVNPEFNERFAKAKTDNAKAKLTKEKDKFIKGVHSLASLEQAIEHHTARH

DDESVQAGKLGQYFKHGLAGVDNPIQKIHNNHSTIKGFLERERPAGERALPKIKSGKNPE

MTQLRQLKELLDNALNVAHFAKLLTTKTTLDNQDGNFYGEFGVLYDELAKIPTLYNKVRD

YLSQKPFSTEKYKLNFGNPTLLNGWDLNKEKDNFGVILQKDGCYYLALLDKAHKKVFDNA

PNTGKSIYQKMVYKYLEVRKQFPKVFFSKEAIAINYHPSKELVEIKDKGRQRSDDERLKL

YRFILECLKIHPKYDKKFEGAIGDIQLFKKDKKGREVPISEKDLFDKINGIFSSKPKLEM

EDFFIGEFKRYNPSQDLVDQYNIYKKIDSNDNRKKENFYNNHPKFKKDLVRYYYESMCKH

EEWEESFEFSKKLQDIGCYVDVNELFTEIETRRLNYKISFCNINADYIDELVEQGKLYLF

QIYNKDFSPKAHGKPNLHTLYFKALFSEDNLADPIYKLNGEAQIFYRKASLDVNETTIHR

AGEVLENKNPDNPKKRQFVYDIIKDKRYTQDKFMLHVPITMNFGVQGMTIKEFNKKVNQS

IQQYDEVNVIGIDRGERHLLYLTVINSKGEILEQRSLNDITTASANGTQMTTPYHKILDK

REIERLNARVGWGEIETIKELKSGYLSHVVHQISQLMLKYNAIVVLEDLNFGFKRGRFKV

EKQIYQNFENALIKKLNHLVLKDKADDEIGSYKNALQLTNNFTDLKNIGKQTGFLFYVPA

WNTSKIDPETGFVDLLKPRYENIAQSQAFFGKFDKICYNADKDYFEFHIDYAKFTDKAKN

SRQTWTICSHGDKRYVYDKTANQNKGATKGINVNDELKSLFARHHINDKQPNLVMDICQN

NDKEFHKSLMYLLKTLLALRYSNASSDEDFILSPVANDGGVFFNSALADDTQPQNADANG

AYHIALKGLWLLNELKDSDDLNKVKLAIDNQTWLNFAQNR

>McCas12a

MLFQDFTHLYPLSKTMRFELKPIGKTLEHIHAKNFLSQDETMADMYQKVKAILDDYHRDF

IADMMGEVKLTKLAEFYDVYLKFRKNPKDDGLQKQLKDLQAVLRKEIVKPIGNGGKYKAG

YDRLFGAKLFKDGKELGDLAKFVIAQEGESSPKLAHLAHFEKFSTYFTGFHDNRKNMYSD

EDKHTAITYRLIHENLPRFIDNLQILATIKQKHSALYDQIINELTASGLDVSLASHLDGY

HKLLTQEGITAYNTLLGGISGEAGSRKIQGINELINSHHNQHCHKSERIAKLRPLHKQIL

SDGMGVSFLPSKFADDSEMCQAVNEFYRHYADVFAKVQSLFDGFDDHQKDGIYVEHKNLN

ELSKQAFGDFALLGRVLDGYYVDVVNPEFNERFAKAKTDNAKAKLTKEKDKFIKGVHSLA

SLEQAIEHYTARHDDESVQAGKLGQYFKHGLAGVDNPIQKIHNNHSTIKGFLERERPAGE

RALPKIKSGKNPEMTQLRQLKELLDNALNVAHFAKLLTTKTTLDNQDGNFYGEFGALYDE

LAKIPTLYNKVRDYLSQKPFSTEKYKLNFGNPTLLNGWDLNKEKDNFGIILQKDGCYYLA

LLDKAHKKVFDNAPNTGKNVYQKMIYKLLPGPNKMLPKVFFAKSNLDYYNPSAELLDKYA

QGTHKKGNNFNLKDCHALIDFFKAGINKHPEWQHFGFKFSPTSSYQDLSDFYREVEPQGY

QVKFVDINADYINELVEQGQLYLFQIYNKDFSPKAHGKPNLHTLYFKALFSKDNLANPIY

KLNGEAQIFYRKASLDMNETTIHRAGEVLENKNPDNPKKRQFVYDIIKDKRYTQDKFMLH

VPITMNFGVQGMTIKEFNKKVNQSIQQYDEVNVIGIDRGERHLLYLTVINSKGEILEQRS

LNDITTASANGTQMTTPYHKILDKREIERLNARVGWGEIETIKELKSGYLSHVVHQISQL

MLKYNAIVVLEDLNFGFKRGRFKVEKQIYQNFENALIKKLNHLVLKDEADDEIGSYKNAL

QLTNNFTDLKSIGKQTGFLFYVPAWNTSKIDPETGFVDLLKPRYENIAQSQAFFGKFDKI

CYNADKDYFEFHIDYAKFTDKAKNSRQIWKICSHGDKRYVYDKTANQNKGATKGINVNDE

LKSLFARHHINDKQPNLVMDICQNNDKEFHKSLIYLLKTLLALRYSNASSDEDFILSPVA

NDEGMFFNSALADDTQPQNADANGAYHIALKGLWVLEQIKNSDDLNKVKLAIDNQTWLNF

AQNR

>MlCas12a

MLFQDFTHLYPLSKTVRFELKPIGKTLEHIHAKNFLSQDETMADMYQKVKAILDDYHRDF

ITKMMSEVTLTKLPEFYEVYLALRKNPKDDTLQKQLTEIQTALREEVVKPIDSGGKYKAG

YERLFGAKLFKDGKELGDLAKFVIAQEGESSPKLPQIAHFEKFSTYFTGFHDNRKNMYSS

DDKHTAIAYRLIHENLPRFIDNLQILVTIKQKHSVLYDQIVNELNANGLDVSLASHLDGY

HKLLTQEGITAYNRIIGEVNSYTNKHNQICHKSERIAKLRPLHKQILSDGMGVSFLPSKF

ADDSEMCQAVNEFYRHYAHVFAKVQSLFDRFDDYQKDGIYVEHKNLNELSKQAFGDFALL

GRVLDGYYVDVVNPEFNDKFAKAKTDNAKEKLTKEKDKFIKGVHSLASLEQAIEHYIAGH

DDESVQAGKLGQYFKHGLAGVDNPIQKIHNSHSTIKGFLERERPAGERTLPKIKSDKSLE

MTQLRQLKELLDNALNVVHFAKLLTTKTTLDNQDGNFYGEFGALYDELAKIATLYNKVRD

YLSQKPFSTEKYKLNFGNPTLLNGWDLNKEKDNFGVILQKDGCYYLALLDKAHKKVFDNA

PNTGKSVYQKMVYKLLPGPNKMLPKVFFAKSNLDYYNPSAELLDKYAQGTHKKGDNFNLK

DCHALIDFFKASINKHPEWQHFGFEFSLTSSYQDLSDFYREVEPQGYQVKFVDIDADYID

ELVEQGQLYLFQIYNKDFSPKAHGKPNLHTLYFKALFSEDNLANPIYKLNGEAEIFYRKA

SLDMNETTIHRAGEVLENKNPDNPKERQFVYDIIKDKRYTQDKFMLHVPITMNFGVQGMT

IKEFNKKVNQSIQQYDEVNVIGIDRGERHLLYLTVINSKGEILEQRSLNDIITTSANGTQ

MTTPYHKILDKREIERLNARVGWGEIETIKELKSGYLSHVVHQISQLMLKYNAIVVLEDL

NFGFKRGRFKVEKQIYQNFENALIKKLNHLVLKDKADNEIGSYKNALQLTNNFTDLKSIG

KQTGFLFYVPAWNTSKIDPVTGFVDLLKPRYENIAQSQAFFDKFDKICYNADKGYFEFHI

DYAKFTDKAKNSRQIWTICSHGDKRYVYDKTANQNKGATIGINVNDELKSLFARYRINDK

QPNLVMDICQNNDKEFHKSLTYLLKALLALRYSNASSDEDFILSPVANDKGVFFNSALAD

DTQPQNADANGAYHIALKGLWLLNELKNSDDLDKVKLAIDNQTWLNFAQNR

>Pb2Cas12a

MKFTDFTGLYSLSKTLRFELKPIGKTLENIKKAGLLEQDQHRADSYKKVKKIIDEYHKAF

IEKSLSNFELKYQSEDKLDSLEEYLMYYSMKRIEKTEKDKFAKIQDNLRKQIADHLKGDE

SYKTIFSKDLIRKNLPDFVKSDEERTLIKEFKDFTTYFKGFYENRENMYSAEDKSTAISH

RIIHENLPKFVDNINAFSKIILIPELREKLNQIYQDFEEYLNVESIDEIFHLDYFSMVMT

QKQIEVYNAIIGGKSTNDKKIQGLNEYINLYNQKHKDCKLPKLKLLFKQILSDRIAISWL

PDNFKDDQEALDSIDTCYKNLLNDGNVLGEGNLKLLLENIDTYNLKGIFIRNDLQLTDIS

QKMYASWNVIQDAVILDLKKQVSRKKKESAEDYNDRLKKLYTSQESFSIQYLNDCLRAYG

KTENIQDYFAKLGAVNNEHEQTINLFAQVRNAYTSVQAILTTPYPENANLAQDKETVALI

KNLLDSLKRLQRFIKPLLGKGDESDKDERFYGDFTPLWETLNQITPLYNMVRNYMTRKPY

SQEKIKLNFENSTLLGGWDLNKEHDNTAIILRKNGLYYLAIMKKSANKIFDKDKLDNSGD

CYEKMVYKLLPGANKMLPKVFFSKSRIDEFKPSENIIENYKKGTHKKGANFNLADCHNLI

DFFKSSISKHEDWSKFNFHFSDTSSYEDLSDFYREVEQQGYSISFCDVSVEYINKMVEKG

DLYLFQIYNKDFSEFSKGTPNMHTLYWNSLFSKENLNNIIYKLNGQAEIFFRKKSLNYKR

PTHPAHQAIKNKNKCNEKKESIFDYDLVKDKRYTVDKFQFHVPITMNFKSTGNTNINQQV

IDYLRTEDDTHIIGIDRGERHLLYLVVIDSHGKIVEQFTLNEIVNEYGGNIYRTNYHDLL

DTREQNREKARESWQTIENIKELKEGYISQVIHKITDLMQKYHAVVVLEDLNMGFMRGRQ

KVEKQVYQKFEEMLINKLNYLVNKKADQNSAGGLLHAYQLTSKFESFQKLGKQSGFLFYI

PAWNTSKIDPVTGFVNLFDTRYESIDKAKAFFGKFDSIRYNADKDWFEFAFDYNNFTTKA

EGTRTNWTICTYGSRIRTFRNQAKNSQWDNEEIDLTKAYKAFFAKHGINIYDNIKEAIAM

ETEKSFFEDLLHLLKLTLQMRNSITGTTTDYLISPVHDSKGNFYDSRICDNSLPANADAN

GAYNIARKGLMLIQQIKDSTSSNRFKFSPITNKDWLIFAQEKPYLND

>PsCas12a

MENFKNLYPINKTLRFELRPYGKTLENFKKSGLLEKDAFKANSRRSMQAIIDEKFKETIE

ERLKYTEFSECDLGNMTSKDKKITDKAATNLKKQVILSFDDEIFNNYLKPDKNIDALFKN

DPSNPVISTFKGFTTYFVNFFEIRKHIFKGESSGSMAYRIIDENLTTYLNNIEKIKKLPE

ELKSQLEGIDQIDKLNNYNEFITQSGITHYNEIIGGISKSENVKIQGINEGINLYCQKNK

VKLPRLTPLYKMILSDRVSNSFVLDTIENDTELIEMISDLINKTEISQDVIMSDIQNIFI

KYKQLGNLPGISYSSIVNAICSDYDNNFGDGKRKKSYENDRKKHLETNVYSINYISELLT

DTDVSSNIKMRYKELEQNYQVCKENFNATNWMNIKNIKQSEKTNLIKDLLDILKSIQRFY

DLFDIVDEDKNPSAEFYTWLSKNAEKLDFEFNSVYNKSRNYLTRKQYSDKKIKLNFDSPT

LAKGWDANKEIDNSTIIMRKFNNDRGDYDYFLGIWNKSTPANEKIIPLEDNGLFEKMQYK

LYPDPSKMLPKQFLSKIWKAKHPTTPEFDKKYKEGRHKKGPDFEKEFLHELIDCFKHGLV

NHDEKYQDVFGFNLRNTEDYNSYTEFLEDVERCNYNLSFNKIADTSNLINDGKLYVFQIW

SKDFSIDSKGTKNLNTIYFESLFSEENMIEKMFKLSGEAEIFYRPASLNYCEDIIKKGHH

HAELKDKFDYPIIKDKRYSQDKFFFHVPMVINYKSEKLNSKSLNNRTNENLGQFTHIIGI

DRGERHLIYLTVVDVSTGEIVEQKHLDEIINTDTKGVEHKTHYLNKLEEKSKTRDNERKS

WEAIETIKELKEGYISHVINEIQKLQEKYNALIVMENLNYGFKNSRIKVEKQVYQKFETA

LIKKFNYIIDKKDPETYIHGYQLTNPITTLDKIGNQSGIVLYIPAWNTSKIDPVTGFVNL

LYADDLKYKNQEQAKSFIQKIDNIYFENGEFKFDIDFSKWNNRYSISKTKWTLTSYGTRI

QTFRNPQKNNKWDSAEYDLTEEFKLILNIDGTLKSQDVETYKKFMSLFKLMLQLRNSVTG

TDIDYMISPVTDKTGTHFDSRENIKNLPADADANGAYNIARKGIMAIENIMNGISDPLKI

SNEDYLKYIQNQQE

>Rb1Cas12a

MKSFDSFTNLYSLSKTLKFEMRPVGNTQKMLDNAGVFEKDKLIQKKYGKTKPYFDRLHRE

FIEEALTGVELIGLDENFRTLVDWQKDKKNNVAMKAYENSLQRLRTEIGKIFNLKAEDWV

KNKYPILGLKNKNTDILFEEAVFGILKARYGEEKDTFIEVEEIDKTGKSKINQISIFDSW

KGFTGYFKKFFETRKNFYKNDGTSTAIATRIIDQNLKRFIDNLSIVESVRQKVDLAETEK

SFSISLSQFFSIDFYNKCLLQDGIDYYNKIIGGETLKNGEKLIGLNELINQYRQNNKDQK

IPFFKLLDKQILSEKILFLDEIKNDTELIEALSQFAKTAEEKTKIVKKLFADFVENNSKY

DLAQIYISQEAFNTISNKWTSETETFAKYLFEAMKSGKLAKYEKKDNSYKFPDFIALSQM

KSALLSISLEGHFWKEKYYKISKFQEKTNWEQFLAIFLYEFNSLFSDKINTKDGETKQVG

YYLFAKDLHNLILSEQIDIPKDSKVTIKDFADSVLTIYQMAKYFAVEKKRAWLAEYELDS

FYTQPDTGYLQFYDNAYEDIVQVYNKLRNYLTKKPYSEEKWKLNFENSTLANGWDKNKES

DNSAVILQKGGKYYLGLITKGHNKIFDDRFQEKFIVGIEGGKYEKIVYKFFPDQAKMFPK

VCFSAKGLEFFRPSEEILRIYNNAEFKKGETYSIDSMQKLIDFYKDCLTKYEGWACYTFR

HLKPTEEYQNNIGEFFRDVAEDGYRIDFQGISDQYIHEKNEKGELHLFEIHNKDWNLDKA

RDGKSKTTQKNLHTLYFESLFSNDNVVQNFPIKLNGQAEIFYRPKTEKDKLESKKDKKGN

KVIDHKRYSENKIFFHVPLTLNRTKNDSYRFNAQINNFLANNKDINIIGVDRGEKHLVYY

SVITQASDILESGSLNELNGVNYAEKLGKKAENREQARRDWQDVQGIKDLKKGYISQVVR

KLADLAIKHNAIIILEDLNMRFKQVRGGIEKSIYQQLEKALIDKLSFLVDKGEKNPEQAG

HLLKAYQLSAPFETFQKMGKQTGIIFYTQASYTSKSDPVTGWRPHLYLKYFSAKKAKDDI

AKFTKIEFVNDRFELTYDIKDFQQAKEYPNKTVWKVCSNVERFRWDKNLNQNKGGYTHYT

NITENIQELFTKYGIDITKDLLTQISTIDEKQNTSFFRDFIFYFNLICQIRNTDDSEIAK

KNGKDDFILSPVEPFFDSRKDNGNKLPENGDDNGAYNIARKGIVILNKISQYSEKNENCE

KMKWGDLYVSNIDWDNFVTQANARH

>Rb2Cas12a

MSVFKSFTNCYALSKTLRFELKPVGKTFENMRTQLAYNKDLQTFLKDQAIEDAYQKLKPL

FDKLHEEFITDSLDSDQAKKIDFSEYLVLYEAKKELQAIEKKLREEIGKTFIAAGEKWKQ

EKYAQYTWKKGSKVANGSDILLTQDVLELIRDLNDKNEELKKMIEETFKGFFTYLSGFNQ

NRKNYYTIKEEKATAVATRIVHENLPKFCDNILFFIDRQTEYLIAHSFLKEKGRDLVNKD

GKALLPITDSIFSIEHFNHCFSQKQIEAYNAQIGNANVLINLYNQAHNDEQGFKRLPAFK

TLYKQIGCGKRKSLFFTLTCDTEAEASKMRNENKEAFSVEEVLNLAYKAGEKYFQNSIEN

DSNLTIPKFCSYIEAQQDYDGIYWSKAALNTISNKYFANYHVLKDRLKEVKVFQKAAKGS

EEDVKIPEAIELEGLFAVIDEVDGWRDEDIFFKKSLIEERKDEKENKKNKKRLEVIKKAE

KPSQALINLIFFDINEHIEQFFDTSKAILSLQEYKSKESKEAIKAWMDHALAVNQILKYF

LVKENKTKGNPLDSEISNALKNILFEGKIIFDGKEIDVDWFRWYDALRNYLTKKPQDDAK

ENKLKLNFKNSTLAGGWDINKEPDNHCVILQDQNDKQYLGVIAKKEKQRGYNKIFEKTPE

NPLYKIDSGEVWQKMEYKQIAAPTGIGGFVRKCFKTAQQYGWKCPDNCLNSEGKIIIKND

EAKENLEAIIDCYKDFFIKYEKDGFSYKKFSFNFKKSSEYEELNNFFSDVERQGYKLDFT

TINKAIIDQWVEDGTIYLFEIKNQDANDGKKEGHKNNLHTIYWKALFENNEDKPKLNGGA

ELFYRKALPKSKQEKIKDNHGKEIIKNFRFSKEKFLFHCPIKMNYKAKSYSDPKYALPEI

NNQINEALTTFGDIHFLGIDRGEKHLAYYSLVDKNGEMIDKGTLNLPFIDQEGKPRSIKK

PKYFYNKKKDKWESEEINCWDYNDLLDAMASNRDMARKNWQTIGTIKELKEGYISQVVRK

IADIVVEHGAFIVLEDLNTGFKRGRQKIEKSVYQKFELALAKKLNFLVDKSAKSGEIGSV

TRALQLTPPVNNYGDIEKRKQVGIMLYTRANYTSQTDPETGWRKIIYLKKGNEEAIKEQI

LQNFTDIWFDGLDYYFEYPNKNKSDKPWKLYSGKGGKSLDRFRRSRGKDKNEWTIEPVNV

VNILKQVFVNFDEKRSLRSQIIEGKALARTKEKTDFTAWEALRFAIDLIQQIRNTGNNEK

DADFLHSPVRDTNGNHFDSRSVSHDRPTSGDANGAYNIARKGLMMNEHIRTWAKKGKPKY

DKNTNDLNLFISEEEWDLYLADKKAWQEKLLMFSSRKAMDEEKKKHI

>Sm1Cas12a

MQTLFENFTNQYPVSKTLRFELIPQGKTKDFIEQKGLLKKDEDRAEKYKKVKNIIDEYHK

DFIEKSLNGLKLDGLEEYKTLYLKQEKDDKDKKAFDKEKENLRKQIANAFRNNEKFKTLF

AKELIKNDLMSFACEEDKKNVKEFEAFTTYFTGFHQNRANMYVADEKRTAIASRLIHENL

PKFIDNIKIFEKMKKEAPELLSPFNQTLKDMKDVIKGTTLEEIFSLDYFNKTLTQSGIDI

YNSVIGGRTPEEGKTKIKGLNEYINTDFNQKQTDKKKRQPKFKQLYKQILSDRQSLSFIA

EAFKNDTEILEAIEKFYVNELLHFSNEGKSTNVLDAIKNAVSNLESFNLTKIYFRSGTSL

TDVSRKVFGEWSIINRALDNYYATTYPIKPREKSEKYEERKEKWLKQDFNVSLIQTAIDE

YDNETVKGKNSGKVIVDYFAKFCDDKETDLIQKVNEGYIAVKDLLNTPYPENEKLGSNKD

QVKQIKAFMDSIMDIMHFVRPLSLKDTDKEKDETFYSLFTPLYDHLTQTIALYNKVRNYL

TQKPYSTEKIKLNFENSTLLGGWDLNKETDNTAIILRKENLYYLGIMDKRHNRIFRNVPK

ADKKDSCYEKMVYKLLPGANKMLPKVFFSQSRIQEFTPSAKLLENYENETHKKGDNFNLN

HCHQLIDFFKDSINKHEDWKNFDFRFSATSTYADLSGFYHEVEHQGYKISFQSIADSFID

DLVNEGKLYLFQIYNKDFSPFSKGKPNLHTLYWKMLFDENNLKDVVYKLNGEAEVFYRKK

SIAEKNTTIHKANESIINKNPDNPKATSTFNYDIVKDKRYTIDKFQFHVPITMNFKAEGI

FNMNQRVNQFLKANPDINIIGIDRGERHLLYYTLINQKGKILKQDTLNVIANEKQKVDYH

NLLDKKEGDRATARQEWGVIETIKELKEGYLSQVIHKLTDLMIENNAIIVMEDLNFGFKR

GRQKVEKQVYQKFEKMLIDKLNYLVDKNKKANELGGLLNAFQLANKFESFQKMGKQNGFI

FYVPAWNTSKTDPATGFIDFLKPRYENLKQAKDFFEKFDSIRLNSKADYFEFAFDFKNFT

GKADGGRTKWTVCTTNEDRYAWNRALNNNRGSQEKYDITAELKSLFDGKVDYKSGKDLKQ

QIASQELADFFRTLMKYLSVTLSLRHNNGEKGETEQDYILSPVADSMGKFFDSRKAGDDM

PKNADANGAYHIALKGLWCLEQISKTDDLKKVKLAISNKEWLEFMQTLKG

>TsCas12a

MTKTFDSEFFNLYSLQKTVRFELKPVGETASFVEDFKNEGLKRVVSEDERRAVDYQKVKE

IIDDYHRDFIEESLNYFPEQVSKDALEQAFHLYQKLKAAKVEEREKALKEWEALQKKLRE

KVVKCFSDSNKARFSRIDKKELIKEDLINWLVAQNREDDIPTVETFNNFTTYFTGFHENR

KNIYSKDDHATAISFRLIHENLPKFFDNVISFNKLKEGFPELKFDKVKEDLEVDYDLKHA

FEIEYFVNFVTQAGIDQYNYLLGGKTLEDGTKKQGMNEQINLFKQQQTRDKARQIPKLIP

LFKQILSERTESQSFIPKQFESDQELFDSLQKLHNNCQDKFTVLQQAILGLAEADLKKVF

IKTSDLNALSNTIFGNYSVFSDALNLYKESLKTKKAQEAFEKLPAHSIHDLIQYLEQFNS

SLDAEKQQSTDTVLNYFIKTDELYSRFIKSTSEAFTQVQPLFELEALSSKRRPPESEDEG

AKGQEGFEQIKRIKAYLDTLMEAVHFAKPLYLVKGRKMIEGLDKDQSFYEAFEMAYQELE

SLIIPIYNKARSYLSRKPFKADKFKINFDNNTLLSGWDANKETANASILFKKDGLYYLGI

MPKGKTFLFDYFVSSEDSEKLKQRRQKTAEEALAQDGESYFEKIRYKLLPGASKMLPKVF

FSNKNIGFYNPSDDILRIRNTASHTKNGTPQKGHSKVEFNLNDCHKMIDFFKSSIQKHPE

WGSFGFTFSDTSDFEDMSAFYREVENQGYVISFDKIKETYIQSQVEQGNLYLFQIYNKDF

SPYSKGKPNLHTLYWKALFEEANLNNVVAKLNGEAEIFFRRHSIKASDKVVHPANQAIDN

KNPHTEKTQSTFEYDLVKDKRYTQDKFFFHVPISLNFKAQGVSKFNDKVNGFLKGNPDVN

IIGIDRGERHLLYFTVVNQKGEILVQESLNTLMSDKGHVNDYQQKLDKKEQERDAARKSW

TTVENIKELKEGYLSHVVHKLAHLIIKYNAIVCLEDLNFGFKRGRFKVEKQVYQKFEKAL

IDKLNYLVFKEKELGEVGHYLTAYQLTAPFESFKKLGKQSGILFYVPADYTSKIDPTTGF

VNFLDLRYQSVEKAKQLLSDFNAIRFNSVQNYFEFEIDYKKLTPKRKVGTQSKWVICTYG

DVRYQNRRNQKGHWETEEVNVTEKLKALFASDSKTTTVIDYANDDNLIDVILEQDKASFF

KELLWLLKLTMTLRHSKIKSEDDFILSPVKNEQGEFYDSRKAGEVWPKDADANGAYHIAL

KGLWNLQQINQWEKGKTLNLAIKNQDWFSFIQEKPYQE

>UbCas12a

MKDFYQFTNLYALSKTLRFSLIPTPATKQMLEDAKVFEKDETIQKKYEATKPYFDRLHRE

FALEALQDQKLDFKNYLELYRKYKADKKASGKLLINIEKDLRKEVVKLFDKQGEKWAKQY

PGLKNKNIGVLFKEAVFTVILKERYGNEKETQILDESSGQLVSIFDSWKGFIGYFKKFHE

TRKNFYKDDGTSTALATRIIDQNLKRFCDNILIFESTKEKVDFSEVEISFGKPLSEVFTL

EFYNTCFLQNGIDFYTKILGGETLQNGEKVKGLNECINLHKQKTGEKLPFFKSLDKQILS

EKDKFFIDEISNETQLLEVLKSFVASAESKTDTIKTLVDDFVKDQDKYDLNYIYFSNDGL

NTITRKWTTETQVFEEALYTALKAAKVVSSSAKKNEGGYSFPDFIPFAHLKTALESIKID

GTIWRDNFNAIENFEEKSIWAQFLAIYNFELSNLFETEIKNPEIGNCPTIGYNVYKQDFE

ELLKSFVYDPNAKVTIKNFADNVLSIYQMAKYFAVEKKRGWNTDYELDVFYTDPQNGYLQ

YYENAYEEIVQVYNKLRNYLTKKPYSEEKWKLNFDSGTPIKYTTRAIIFNNTTNERYYLG

LLKKGVAKPREFEPINNNIISSGEFRRMIIQQLKFQTLAGKGYVRDFGVKYSEDKDGVKH

LQQLIKKQYLSKYPCLKKIADGVYNDKKAFDADIKDVLLETYNLDFQPISEEFILNKNRL

GEIYLFEIHNKDWNLKDGKNKSGSKNLHTMYFESLFVDKTTFKLNNEGAEVFYRPATNEG

KLGTKKDRNGKIIINHKRYATDKILFHCPIGLNKDAGKSYTFNAKINNMLANNPDINIIG

VDRGEKHLAYYSVITQKGKILDRGSLNKVEGGDKQEIDYAKKLEETAKNREQARKDWQAV

EGIKDLKRGYISQVVRKLADLAIEHNAIIVFEDLNMRFKQIRGGIEKSVYQQLEKALIDK

LSFLVMKGEADPEKAGHLLKAYQLVAPFESFQSMGKQTGIIFYTQANYTSKIDPITGWRP

NLYLKYTSAEKAKADILKFSKIEFVNNRFELTYDIKNFVLDKKVVLSNKTKWTVCSSVER

FRWNRRLESNQGNYEHYENLTENLSSLFKDFGFEIEQNIIRQVEQLATKGNEQFFRSFIF

YVNLIFQIRNTDAKAKDQNKEDFILSPVEPFFDSRTPEKFGENLPENGDDNGAFNIARKG

IIMLNKISAYKQEVGNVDKIIWKDLFISAAEWDNFTQE

>MCK9303417.1 hypothetical protein [Bacteroidales bacterium]

MEAIKNNYQLSKTLRFGLTQKSKTRKDGFTGEIYQSHNELKDLVKSSEDRIKKSVSTDEK

SEMSLSVDKIRCCLVMISDFLSSWQQVYSRADQIALDKDYYKILCKKIGFDGFWVDERYD

RKNDKTVRTKKPQSRTINLSELDKKDDKGIERRQYLLTYWRDNLINAADKFEVVTEKLKQ

FEDALNINRTYNKPNEVELRKLFLSLTNIVQETLQPLCLGQICFPKLEKIDDSRTENKHL

IDFATDYQSKSDLLSEISELKKYFEENGGNVPYCRATLNQKTAVKNPNSTDNSIDSEIKK

LGLDKILKENKDALYFANKIYSLSAKEKLSKLDDKTTGLIERSLLFKYKPVPAIVQYEIA

KTLSETINKSEEDLLEFLRSIGQTKSPTKDYADLQDKNDFDLDAYPLKVAFDFAWENLAR

SIYHSDADIPVAVCEGFLKKNFGIDKSNADFKLYAQLQELKAVLATLEYGNPTNRQTFIN

EATKLLSPISWDKIGRNGNQNKYSIEKWLKTLTKDDKDYKDAKQQIALFRGRLKNNIKTF

DDITKYFKSVAMEMGRTFAQMRDKITGAAELNKVTHYAIIIEDQNFDKYVLLQEFVDKKE

NRIYAKTDRHHSDFTTYSVNSVTSGSIAKMLRKKRMDELNRNNRNSFEQKPELSEEQKEQ

RNIREWKEFIEDKRWDLEFQLNLSNKTFEQIKKEVDAKCYELDINYISQETLSDLVNKKG

CLLLPIVSQDIAKENKTEGNQFTKDWNAIFTQETPWRLTPEFRVSYRKPTPNYPVSDKGD

KRYSRFQMIAHFLCDYLPKSDSYISNWEQIANYKDDKLQEKAVKEFNADLRGRTEEEKQS

ESVNALLAHFGNQNKKQKPVERPKEKFYVFGIDRGQKELATLCVIDQDKKIVGDFDIYTR

SFNSEAKQWEHKLLEKRVILDLSNLRVETTIVIDGKPEKKKVLVDLSEVKVKDKDGKYSK

PNKMQVKMQQLAYIRKLQFQMQTNPDAVLEWYANNKTKEQIMSNFVDNENGDKGLVSFYG

TAVEELNETLPIDKIEEILKKFQELKDKEKQGETVKLEIDKLVQLEPVDNLKNGVVANMV

GVIAYLLQNLDYQVYISLEDLSKPFSGQIIGGIAGVPTKTNKEEGRRADVEKYAGLGLYN

FFEMQLLKKLFRIQQDSQNILHLVPPFRAMKNYDHVAVGKGKVKNQFGIVFFVDADATSK

TCPCCGSSNNKPNLKMYPNAKKGLSKEGKEVWVERDKSEGNDIIRCFVCGFDTTKDYSEN

PLRYIKSGDDNAAYLISAEGIKAYELATTLVNNK

>MCI1720368.1 transposase [Bacteroidales bacterium]

MKNIVNNYQISKTLRFGLTQKTKIQKEGYNGEIYVSHRELGDLVKISEERIKKSVSSSDK

SNLELSLDKIDICLKQIGAFLLDWQQVYYRKDQVALDKDYYKILCKKIEFDGFWKDGRGQ

RMPNSRIINISELDTKDILGVERLQYILNYWKDNLVSASQKYSVVEEKIKRFKLAIKINR

TDNKPDEVELRKMFLSLVNIVCDTLQPLCYGQISFPKINKLDDSRADNKKLIKFATDYKS

KNDLLTSIAEQKKYFEENGGNVPFCRATLNPKTAIKDPNSTDNSIKGEIVQLGLDSILKS

FKSYLFFENSLENMPAKEKIELMKSSGANVVKKGLMFKYKPIPVIVHREVAQELSKDLNK

TEESLSDFLRGIGQAKSPAKDYEELTDKNKFNIEAYPIKVAFDFAWESLAKAKYHSEIDL

PVAACEKFLNTFFDVKPNNVHFLLYAKLQELNALISTLEYGNPSDEQSIVKKIKALSNEI

KWDDFGGSGQGYKKSISDWADSKKDSDGFKIAKQKIGLFRGGLRNEIAEYYNLTQIYKKT

VMQEGKLFATMRDKITGAAEQNKVTHYAAIIEDKTGDKYVLLQEVPLDKQDRIYDKMDRN

GDGYVSYFVNSITSRTIAKQLRKKRMAELMKNNARGIYNNVSNIQQPALSDKEKEERNIK

EWISFISEKRWNYEFNLNFKGKNFEEIKKEVDANGYELENRILSRDALEELVKNKKCLLL

PIVNQDIIKESKTESNQFTKDWNSIFDGNSPWRLTPEFRVSYRKPTPDYPMSDKGDKRYS

RFQMIAHFLCDYIPQGGSYVSVREQIDNYKDDKKQEIAVKDFHDRLLGKTDEQKFIEGLG

GLSSLGNVTIKTKLKKQDISKEKFYVFGIDRGQNELATLCVIDQDKKIQGGFRIYTRLFN

NEKKQWEHKFLEERNILDLSNLRVETTIVIDGQEKREKVLVDLSEVKVKDNSGNYVKPNK

TQIKLQQLAYIRKLQFQMQTNPGRVLEWYSHNQTSDLIIDNFVDKQNGEEGLVPFFGAAV

AELKDTLPIDRISDMLKQFVELKNLEKQGEDVKSKIDQLIELEPADNLKSGVVANMVGVI

AFLLKKYNYKVYISLEDLSKPFKDQIVLGISGVPIGIKKGMAGRSINVEQYAGLGLYNFF

EMQLLKKLFRIQQDSSHILHLVPAFRAMKNYDNVAVGKGKIKNQFGIVFFVDAAATSKTC

PCCGAINDRQFAPDLRKFPNAKKIETPDGKSVWLERDKTDGKDIIRCHVCGFDTSKEYDD

NPRKYIKSGDDNAAYLISAECIKAYELATTLVDNK

>NUJ97582.1 transposase [Candidatus Gracilibacteria bacterium]

MRFGLTQPNKKGELKTHIEFSDLVNKSFENIKKEVNSKDKSKFDTRKELIDKINQFISGL

ENQLGDWKNMYERYDLISVNKDYYKILARKAKFDAFKKDKKGVKQPQANQIKLSSLRYNK

ELIINYWGNIISRSDYLINVFKPKLEQYLNAVNNPNNSSHTKPDLIDFRKVFLQLLKISE

EYLQPLFNKSIQFETGKKENSGDIKRVNDFSGNENNKEINDLLDLGKEIREYFEANGSQV

PYGKVSLNYYTAVQKPNNFDKEIKEGIKDLGIIEFLKKSEEDIKNYLKQDSKEKIYLLNN

SKNPYSIELIQLFKPKTIPFSVKYNLSKYLEKNYNLKYEDILNKFDLLGKSVDIGKDYLE

CKDKEKFSLEKYPIKSAFDYSWENLARSLKRDVDFPKNVCEKYLNDNFNINVGNSSFNLY

ANLLFIAENLATIEYGKPNNEKEIIDSIKETFLELSDEIEKNNKKNEVENIIKYLNLNTD

ERKNIKDLQKKYFKNLDTKEQNILNIFDSFTKSKQSLGLLRGQQKNKIDKYRNLTQKLVD

KKDSHIGIASFIGRTLASIREGLKEENELNKITDYGIIIEDKNQDKYILTLKLNGKDTRE

KIKNNLGNGEYKVFEINSFTSKALNKFIKNPLGEDSKKFHGYFQYKHREVSIYDENEKWV

GYKEEFLKHLKHSLINSQIAVEQNWKDFGWNFDNCDTYEKIEKEVDKKGYKLIETSISKE

NLENLIHKEDCLLFPLINQDISSKKEENKNDFTKNFEKVFLGDGYRIHPEFSIFYRQPNE

ENLKPNKSGIINRFGRLQLLANIGVEYIPQNNDYTTRKEQNKISIDQTKQNESVQKFNKE

KVNPYFDSLEDYYIFGIDRGIKQLATLCITNKKGVIQNFDIYTKHFNDNSKNWEYKNNRT

EGILDLTNLKVESDKEGNKYLVDLSLFEAKDENGNLTGTNKQNVKLKQLAYIRKLQYQMS

SNEEGVLSFLNKYKTKEERQNNIKELITPYKEGHHFEDLPMNIFEEMFENYEKLKNNKTL

SEGEKQNLMKLTTELDASEDLKKGVVANIIGVIVHLMKEYDYKVKIAIEDLSNAWYFSKD

GLSGDSILNSKIDEEMDLKKQDNLALAGVGTYHFFEMQLFKKLFKISVEKGILHLVPSFG

NVRNYTDLLKEKYKYQYQQFGVIYFISPKFTSSKCPICGKGGKKHIKRENNVITCKECGF

VSGKDNSINIKNNKKEGLNLDLIKNGDDNGSYNIGGKIK

>MBR4267799.1 hypothetical protein [Bacteroidales bacterium]

MTKYQLTKTLRFGLTKVRKKTKLVAGKEVDAKYLSHEELDDLVMRSEYNLIKRNVLEWSK

KKESDKSNYIFDELDKRFEEIIEIEDRDQRNAEYVEFLNEFFKEVNHQTINQLDESTFIN

KIGDCSKAIKEYLLSWGKVCRRIDKITVRKDYFKILARKTFFKYESKVGKKRTPLPSEVK

LSGQKGNNYFDEPINEGISQFWQNRVAKALNLHSQLESMLFDYKKAIETEKHNQENPKGD

NGSFDKLHLVDFRKMFLSVCSLVMDSLRPIVNELIIVSDNVSKDEDKYILDFVNDKKTQW

DLFQQIENLQTICKDNGENIFFGKATFNKYTSEQAPNHRNNDIAKVLRELKIEKFVSDYI

DLDQEAINRKIYQSTQSRLENLNNPQISPIIRAQYFKYKPIPTLVRFGLAKELAKQQGKK

YSDRLKGIQELFRIFGSSKSPALDYKNNRTDFSLDNYPIKVAFDYAWEMCARSEYAQKPV

DFPKSICEKFLEKFFECKSNEKYQQSFVTYARLLKINEDLATLEHFENEPPKDIESIYQD

AQRYLDEVGNLCSNEDRAAIAKWYEEYNKLWTKGDHKKLKEWIVSKSSIITNFTQAKMHL

GQKRGSQKTFVLKSYFHSSYGKIRDNNRFVNSNVTEVFKTIASTFGKSFATIREYFNEES

EVNKIEYGAVIIKDKNGDKYLLLQKKNEGGIDMPIFNKSDENGDCDLYQVKSLTSKTVRK

IIASPNKYNDFFVNNDGKKIIYPDKTDFKYKINPYDKEEVKKRKRELYNNDLVRPIIYSL

TQSKFANKQNFEKYFDWTKALKQCSNIEQLYKTIDQKGYSLNPSKISKEQIADLVNNMNC

YLLPIVNQNITAKTKNDTNQFTKDWNKIFNEVDKDYRIHPEFTMFYRYPTPDYPKFGEKR

YSRFQMNVNFLMEVIPADGEYCSRKEQIEIYNAPKDNENCQKNVVERFNNKIKALKPSYF

IGIDRGINELATLCVIDKEGKIVGDFEIYKREFDSNLKRQKYTSIETRDILDLSYLRVEK

DENGESRLVDLSESEVWIDSLDHDKGKRANKQIVHLKHLYYLRCIAHLLQSSDYKSIVLE

KLKDCNNLKDEEIKKVFEKDKFVDSYKGGEAYTDLPYDEIRKLISDYQEIEQSNQTESEK

SKALNTLCQLDASEYLKKGVVANMIGVVVYVLKKYNYDAYISLENLCYAYGYSKDTLSGY

SITSTKEDPYLDFKDQENAKLAGLGTYSFFEVQLLKKLFKLQIEENTELIPAFRSVDNYE

KIFLLKNIDNKIYQFGIVYFVDPKYTSLCCPICGEHGKKNVDRKKHTKKYDEDELVCKQC

GFHTNLSHIETRVMEDKTIKNSYDECNLKAIVSGDANAAYNIAIRLGKNIYSTIADKVKD

LHHEGKKYIIVKG

>KKQ38176.1 hypothetical protein US54_C0016G0017 [Candidatus Roizmanbacteria bacterium GW2011_GWA2_37_7]

MEIQELKNLYEVKKTVRFELKPSKKKIFEGGDVIKLQKDFEKVQKFFLDIFVYKNEHTKL

EFKKKREIKYTWLRTNTKNEFYNWRGKSDTGKNYALNKIGFLAEEILRWLNEWQELTKSL

KDLTQREEHKQERKSDIAFVLRNFLKRQNLPFIKDFFNAVIDIQGKQGKESDDKIRKFRE

EIKEIEKNLNACSREYLPTQSNGVLLYKASFSYYTLNKTPKEYEDLKKEKESELSSVLLK

EIYRRKRFNRTTNQKDTLFECTSDWLVKIKLGKDIYEWTLDEAYQKMKIWKANQKSNFIE

AVAGDKLTHQNFRKQFPLFDASDEDFETFYRLTKALDKNPENAKKIAQKRGKFFNAPNET

VQTKNYHELCELYKRIAVKRGKIIAEIKGIENEEVQSQLLTHWAVIAEERDKKFIVLIPR

KNGGKLENHKNAHAFLQEKDRKEPNDIKVYHFKSLTLRSLEKLCFKEAKNTFAPEIKKET

NPKIWFPTYKQEWNSTPERLIKFYKQVLQSNYAQTYLDLVDFGNLNTFLETHFTTLEEFE

SDLEKTCYTKVPVYFAKKELETFADEFEAEVFEITTRSISTESKRKENAHAEIWRDFWSR

ENEEENHITRLNPEVSVLYRDEIKEKSNTSRKNRKSNANNRFSDPRFTLATTITLNADKK

KSNLAFKTVEDINIHIDNFNKKFSKNFSGEWVYGIDRGLKELATLNVVKFSDVKNVFGVS

QPKEFAKIPIYKLRDEKAILKDENGLSLKNAKGEARKVIDNISDVLEEGKEPDSTLFEKR

EVSSIDLTRAKLIKGHIISNGDQKTYLKLKETSAKRRIFELFSTAKIDKSSQFHVRKTIE

LSGTKIYWLCEWQRQDSWRTEKVSLRNTLKGYLQNLDLKNRFENIETIEKINHLRDAITA

NMVGILSHLQNKLEMQGVIALENLDTVREQSNKKMIDEHFEQSNEHVSRRLEWALYCKFA

NTGEVPPQIKESIFLRDEFKVCQIGILNFIDVKGTSSNCPNCDQESRKTGSHFICNFQNN

CIFSSKENRNLLEQNLHNSDDVAAFNIAKRGLEIVKV

>MBO5704032.1 hypothetical protein [Bacteroidaceae bacterium]

MEYIKTTKTIRFKLQHGHENKRIVKTISNLQNEEFDLAEFIVNLKQFIDLFYKFIFYKKD

GDDYLQEHMTLKTEWLKKYARHKIKVSTTAKRQQHSIKKYKVDDIIKETLYNMEILYGEL

AKDASAELNERAKRARTGLLLKSLYAKHNLPCLVDLADNIVDKKETTNRSLRLKKMGRDL

LQQLEAGMKIYLPEQSKGAVIAKASFNYYTLNKKPIDFQQKINELEHSLYTCLGDCRRDI

HCNMNLWNAITNDINERTDGKQLCWGDSPFLEEGTYVSLRQTLKNILAEQKAAFSEMMQD

DASYKELRESDLYLFRGISRKEFDDYKDLTNEIEDIATKKNQTSNDNRKKYLQTQLNNLK

KQRGSLISAADYCTKNLFTDYKAFAAWYRDVAKKHGKIFAQLKGIEKERVDSQLLQYWAF

MAEQNGQHQLYLVPREEAKGCYDFLKNATSSEASAACYWLESFTYRSLQKLCFGNLESGS

NDFYREIGRELLQYTTLDCRGYPQFISGEYKFHGDEQKKIKFYQDVLASKSASKIIGLPQ

QELKKKVTDRAFSCLDDFKIALEQVCYRRFMRMSPNILTTIKERFNAQVFEITSLDLRNA

ENIKEYEKCFEHHDKLHTTIWKQFWSNDNANNHFDIRINPELSIIYRTPKDSRVEKYGVD

SERYDERKKNRYLHEQLTLALTISEHCNAPFVNQAFSKVEEVVNNIKNFNQKLNDHHYKF

ALGIDNGEVELSTLGVYLPEFAKSSHEEVLEALRQVDKYGFEVLTITDLSYEESDTNGKS

RRILQNPSYFLNPDLYCRTFGKSATEYEEMFERIFEKRHTLSLDLSTAKVICGHIVTNGD

VTSLFNLWLRHAQRNVYEMNDHAHKEGVKNIALKKSDNLNNGERRKFIDYLAEKSEDYKK

MTEAQKNEYVAWVYKRWNFEDVTPDSVFVRILNKHRVRGNYLHDVLMAVAFEGEELVSVS

EIFNIRNVFKLRKDFYRVKSQDDILAEINAYNKRIISNEELDLKLNQLKSSLVANVVGVI

DFLYKDYQSRYGGEGLVAMEDFNVKNVDAAMEKFAGNIYRMLERKLYLKLQNYGLVPPVK

NLLQVRDNNKLKQIGNVCFVDEAGTSQMCPVCEAGRLGHTETCSRNCGFTSVGILHSNDG

IAGYNIAKKGWNMNKY

>MBL6666846.1 type V CRISPRassociated protein Cas12a/Cpf1 [Flavobacteriaceae bacterium]

MIKEQIRKVRSCKLEELYKQILSDRSIISFRLNEIKNDGELCQLIQNLFRINFNNEIVCK

KEIIDKETAEIIEEVNISEHLENSFRHISEADPDQLYIRNDRALTDISQAIFGNWSLIGN

SLKFYAEKEELGKNKNGIPKKLTSKEQEMWLKHNYFSFGTIHKALQSYFNQFDANELDKE

PSDAQNSANAITVETKELAENKPLFEYFKSFETSKKNKETNAFEKVSLLKSVKSAFQPAL

EMLKKYENEGGEKLKNEKTDVEKIKIYLDALMDLLHFLKPLYVELSRKEEMKAEVFEKDT

GFYDDFDKAYNTLKQIIPLYNQTRNHLTKKPFSVEKYKLNFDNSTLAAGWDKNKETANTA

VILIKDGKYFLAIMTKEYNKLFQTEPHINTAPAYKKMDYKLLPGASKMLPKVFFSNKNID

YYNPSKEVLHIRNHGTHTKNGRPQKGFTKQDFTKKDCHTMVNFYKNSISRHAEWKEFNFE

FSQTNSYNTIDEFYREVEQKGYKITFSDISESYINQCVKEGKLYLFEIYSKDFSKHSKGK

PNLQTLYWKEIFSERNLKNVVYKLNGEAELFFRKASIQYPDIVWKEGHHKNDPLKKQKYP

IIKDRRYAKDTFLFHCPVTCNFKAQGIPRFNDKVNTFLKNNPDVNILGIDRGERHLAYWT

LINQKREIIKQGSFNNPQGKKDYHDLLDIREKERDKARKSWSTIEKIKDLKAGYLSLLVH

KIAKMVEEHNAIVVFEDLNFGFKRGRFKVEKQIYQKLEKALIDKLNYLVFKDKNTAEAGG

VLNGLQLTAPFESFQKMGKQTGVVFYVPAYHTSKVCPATGFVNLLYPRYETIKKSQEFFK

NFCEIKFNSQEDYFEFSFNYEKFTKKAEGSRQNWTLYTYGERLENYRNSLQNNQWDTREV

KLTEAFKKLFNQYVIDFNDGTDIKDQIVRQTEGKFFRQLTHLLKLTLQLRNSRINSDEDW

MISPVRGNNGTFFDSRKEDSKMPENADANGAYHIGLKGLWVLEQINNQEEGKKLSIAITN

KEWYQFIQDKKYQE

>MBK8394035.1 hypothetical protein [Leptospiraceae bacterium]

MEKYQITKTIRFKLEAVNGKANSLKANQTDSKIDLQILIYSLEKVKSDFKTIFFSKDENT

NEDKFKSVFKIKYTWMRTFLKNQFYDSRNMSASSKSYSLIDLDYVLVEFKERWLKEWESI

LNRLKVFHAAPDESYFRKSETALTINQLAKRTNFEFIKEFSYSIEATQKPEIDDSIRIFI

SDLDKLQNTIKSFQKIFLPSQSGGLLVAGGSLNYYTINKTTKFEKQLSDIQENLKKKINS

FRDRDGRQQFYLDQSIVDKIGLPSDIFDKTLDEVYPFLKKWKSDKKNGIIETAQKVELSK

NDILEKGEYNLFHTSDSNLEKFIERTQKIGELAEQKNSDSTPAHLKKELKEKIRELKVKR

GEFFNTPERPVQTQNYKKFCDFYKKIALKRGQELAKKRGIEKEKITAELLKYWCVILEKE

NSRFLYMIPIGNDDNIKKAKEYIKSGKANGSANEPKLYYFQSLTLRALRKLCFKSTDNTF

KHATQKKDFPQYEQAMTSDREKIKFYQEVLKIWKEKGDDNFGLNLSNFNIEDLCNKNYPS

LEEFEIALNKVCYIKKMYISSTIENDLENKFHAISFKIISYDIYHNNLKYNHSNYKNKHH

TDIWKKFWESSDTSHYPLRLNPELKITWREPNDRTVKKYGKDSSLYDPKKKNRYLKEQFT

LGLTFTENAHGKKFDFSFTDHKSIKNAIDGFNAKLAQENKGEWIYGIDRGTKELATLCLV

KFPSPDFKNPEFPKLNLYKLKPNKYFFQDEAKTRYGQVDQISYSYNLMVVSNEEEKKSKI

HVFQEEFQNQKNKYKTLVLKIENKWNLVYFNDQNILREVSLDKQVELIAEFNKSSKKDVK

KINRKLRLICNHKGFSGPIKNISYYLNKLNPDKTNDWFEKIPEQESACFDLTTAKLIKGR

IILNGDLMTFFKLKRLVAQRIIFDLFREGKITQSSEIRTRENKNYLLSVYNGIEVIKPNR

IEEDDDAKNGLVIYRLSLEEKSNFETAGELIIEELNAYLKSLFEKNGTIYEPSIDKINHL

RDSITANMIGIIKFLQTSYPIKSIALENLDEFKETGNKFITDHFIQSEVSIERRLEWSLY

RSLQDLSLVPPNIKETILLKDEFQHLKFGVMKFIETKGTSSNCPACGNKYRKTGNHYICK

DISNCGFSSKDNRKGLDPLINSDKVAAYNIAKRGLEILN

>OGW03971.1 type V CRISPRassociated protein Cpf1 [Nitrospinae bacterium RIFCSPLOWO2_02_FULL_39_110]

MENKELFNSFTKKCQLSKTLRFELVPQGETEKFIKKKGLLKQDKDRADDYRKAKKLIDEY

HKDFIEHALSGKQLKKLQAYYDEFTALLGKPAKERDTRVLSNISEALRKEIAGWLKDNPI

KDEKKLIKEIVPNFLKSKDRNDDAALVLKFKGFTTYFGGFNENRRNMYSAEEKSTAIAYR

IVHENLPKFISNMKTFQRLTENHKIDFLEVETQMNDELDGNKLKDVFSIDYFNQCLTQSG

IDRYNTILGGKTDSRGAQIQGVNIKINLFGQKNNLKSKDAPVLARLYKQILSDRTSASFL

PEKFETAKELIASIREFYEKAINGFIKNNERKNVIDEIESLFNKKLNPTDCDLSQIYIRN

IFISAVSQQIFGSYSVITSALNAFIDKTYKTKKEKEKQQKKSYFGIAEIEAALKAYLQET

EIENKEKREEVLKHANPVLQYLNSLTTNIEINNKKEKANVIVRFRQQHEAVQSVLNKEYG

NNSELIQDKNAVAAIKDYLDSCQHILHFVKPLFVVPPKKENAELPDKDAFYSEFDELFEQ

LKSVTPLYNMVRNFLTQKPYSTEKIKLNFENVSLLSGWDVNKETDNTSVIFRKDSLYYLG

IMDKKHNKVFHDIPASSDNDFYEKMNYKLLPGANKMLPKVIFSDRWIEYFNPSKELIEKY

KKGTHKKGEIFSLSDCHALIDFFKTSIAKHEDWKQFNFQFSPTKTYQDINEFYREVEHQG

YKISFQNVSAKYIHQLVSEGKLFLFKIYNKDFSPYSKGKPNLHTLYWKAVFDKENLKDVV

VKLNGEAEVFYRPQSIPKKSKIVHKKNKPIVNKNPNNPKKQSLFKEYDIVKDKRYTEDKY

HFHVPITLNFKEAKVPSENDRKKAFAYTFNKGANETLKENFPNIIGIDRGERHLAYYSLI

NSKGEILEQGSFNVIKNEFNGTKHETDYYASLVEKEKARDAARKSWDTIGKIKELKEGYL

SQVVHKIARMMIEHNAIIVLEDLTMKFKDIRKKVERQVYQKLEKMLIDKLNYLVFKDREA

YEPGGVLNAYQLAAPFTSFKDMGKQTGFIFYVPAANTSKIDYATGFMNFIYPKIEDAVEF

FQKFEYIRFNPDKNYFEFAARYNNFVKDDEAKLPDEYNEPWIICTHGEQRFQYQPSYKSF

KEVNVTQELQDLFSKHAIMPYAHGNDLSPQIKDKGNTSFYKGLIALLRLVLQMRYTDGKG

RDFILSPVANDEGVFFNSEKAKETEPNDADANGAYHIALKGLLLLKRIKGEFKKEEKTGK

KKTQKENWIDLSISNKEWYEFAQKKEYRK

>BaCas12a3

MDFIKTTKAVRFRLESNNENTLIQESINNLNSRKEFDLNTFVDDLDAFINDCNAFLFCSK

NKGRREIFYVNPSLIVKNEWLKKYAKQDLAELKQNHTAQRVQYKIGDIDGLCYRIQDLID

DLDDIYVKLCDDASAELHERAKRAQTALLLKRLFANNALPCLVSLIDNTVDKNEKDNLSL

KLKSLGKKLLAQLELGIQEYLPEQSSGVNIAKASFNYYTINKKPIDYDRKIEELSDKLVT

TLDFWKRDGSCNFNKSLWKLIEVKSEGKTLYLGDSPLSDTDEYASLRQILKNILAEQKAE

FSEKMQEKISYEDLTKSDLFLFNNISKEEYNGYLELTNQIEELATDINQEDNEYKLKKLR

SDLMKLKKNRGSLINAADRRTKEKFKTYKSFADFYRKVSQRHGKILAQLKGIEKERSESQ

LLQYWALMLEVNNQHKLVLIPKDKAQECKSRLESSNEQAQGTKLYWFESFTFRSLQKLCF

GNLENGSNSFYPGIRKELQYKYSTEDRNGYPQFISGEFEFKGDEQKKIQFYKDVLNTKYA

QSALSFPKEEVKRNIIEKDFESLDDFVIALEQICYQRYVCVNSHMINALGSYFNAQILDI

TSLDLRNPLNSQEKETVYAHADKKHTEIWKKFWTADNEKDNFDIRLNPEITITYRKPKES

RIAKYGVESDKYDANKKNRYLHDQLTLVTTISEHSNSPAKNLAFTTDAELKDMIERFNAE

IKKEKIKFALGIDNGEVELSTLGVYLPGFKKDTKEEVFAELKKVDEYGFKVLEIRNLRYS

ENDYNGKERRIIQNPSYFMNKELYCRTFNKTAAEYDAMFAEVFEEKQLLTLDLTTAKVIN

GHIVTNGDVISLFNLWMRHAQRSIYEMNDHAIKETANDIVLKRSETLNDAEKRKFIDYLN

GKNKKYEDLSEREKSEYVKWVYRIWGGDYSEYGKNKAFAEISKGQRVGDYLNNVLVAVTF

TGKELTNVVDIFDIRNVFKFKEDFYSLKSETEIMEEVNKYNVKNTKSISNEELDLKLNQL

KSSLVANVVGVIDFLYKQYKERFGGDGIIVKEGFDSAKVESDREKFSGNIYRLLERKLYQ

KFQNYGLVPPVKNLMLMRDVDLNDTNEFMQLGNICFVGYEGTSQNCPVCEKGRLGHTEKC

SDNCGFESKGIMHSNDGIAGYNIAKRGFNNFMRK

>SuCas12a2

MLHAFTNQYQLSKTLRFGATLKEDEKKCKSHEELKGFVDISYENMKSSATIAESLNENEL

VKKCERCYSEIVKFHNAWEKIYYRTDQIAVYKDFYRQLSRKARFDAGKQNSQLITLASLC

GMYQGAKLSRYITNYWKDNITRQKSFLKDFSQQLHQYTRALEKSDKAHTKPNLINFNKTF

MVLANLVNEIVIPLSNGAISFPNISKLEDGEESHLIEFALNDYSQLSELIGELKDAIATN

GGYTPFAKVTLNHYTAEQKPHVFKNDIDAKIRELKLIGLVETLKGKSSEQIEEYFSNLDK

FSTYNDRNQSVIVRTQCFKYKPIPFLVKHQLAKYISEPNGWDEDAVAKVLDAVGAIRSPA

HDYANNQEGFDLNHYPIKVAFDYAWEQLANSLYTTVTFPQEMCEKYLNSIYGCEVSKEPV

FKFYADLLYIRKNLAVLEHKNNLPSNQEEFICKINNTFENIVLPYKISQFETYKKDILAW

INDGHDHKKYTDAKQQLGFIRGGLKGRIKAEEVSQKDKYGKIKSYYENPYTKLTNEFKQI

SSTYGKTFAELRDKFKEKNEITKITHFGIIIEDKNRDRYLLASELKHEQINHVSTILNKL

DKSSEFITYQVKSLTSKTLIKLIKNHTTKKGAISPYADFHTSKTGFNKNEIEKNWDNYKR

EQVLVEYVKDCLTDSTMAKNQNWAEFGWNFEKCNSYEDIEHEIDQKSYLLQSDTISKQSI

ASLVEGGCLLLPIINQDITSKERKDKNQFSKDWNHIFEGSKEFRLHPEFAVSYRTPIEGY

PVQKRYGRLQFVCAFNAHIVPQNGEFINLKKQIENFNDEDVQKRNVTEFNKKVNHALSDK

EYVVIGIDRGLKQLATLCVLDKRGKILGDFEIYKKEFVRAEKRSESHWEHTQAETRHILD

LSNLRVETTIEGKKVLVDQSLTLVKKNRDTPDEEATEENKQKIKLKQLSYIRKLQHKMQT

NEQDVLDLINNEPSDEEFKKRIEGLISSFGEGQKYADLPINTMREMISDLQGVIARGNNQ

TEKNKIIELDAADNLKQGIVANMIGIVNYIFAKYSYKAYISLEDLSRAYGGAKSGYDGRY

LPSTSQDEDVDFKEQQNQMLAGLGTYQFFEMQLLKKLQKIQSDNTVLRFVPAFRSADNYR

NILRLEETKYKSKPFGVVHFIDPKFTSKKCPVCSKTNVYRDKDDILVCKECGFRSDSQLK

ERENNIHYIHNGDDNGAYHIALKSVENLIQMK

>MBX2887915.1 MAG: hypothetical protein KF829_04615 [Ferruginibacter sp.]

MENYKLTKAIRFRLEPNEGSNLIQQQVEDMKNISFDLVSFISGLGNFNDLLKRYLLRARK

DGLIKMNSSLSIKSGWLKRYAKQIFHDSEALRDKTRRKQTKTIGDFEGLSAEIEGRLETY

LAIYQSLVNDGVAALNERARKANTGLLLKQLGNQNGLSYFVGLIEASVDKREVDDLSTQL

KKQGKALQEELLAGIQKYLPEQSTGFPLAKASFNYYTINKKPVDYAQKIREQEGKLRITD

WDAQVWSNSGLSNAIKAAVQWGITDLGSLRQKLKNIKAQQKASFNELMTQDLNFQELREN

ADLYLFSNITEHEFNQYKNLTEKIEDKATRINQSSNDNDKRQLRSDLQKLKKDRGALINA

ADKRNQNSFKTYKYFANYYRNIALKHGRILSTIKGIEKEAVESQLLQYWSLVLEQKGKHQ

LLLIPKQKAAECYQFVSAASAIQQSSTSKIYWFESFTLRSLRKLCFGYLDAGTNTFNQNI

QEILRRHQYSEMDNRGRQNQVRIEGEFSFKGNTQKIIACYKDVLASRYTQSVLRLPVRQL

RDEVIEPNFETLDDFQIALERICYKRYALLSEQNLDTITRRYQAQVFAITSADLEKGNDG

NDKAHTQIWKKFWAQENEQTYFDVRINPELSIQWRPAKESRVHKYGRDSALYDEQKNNRY

LHPQFTLVTTISEHSNTPTKVLSFMTDEEFKISVDAFNSKFNNADFKFALGLDNGETELA

TLGVYLPAFKQESNEASIAEIRKIEAYGFKVLSITDLTHQEKDTNGKERKIIQNPSYFLV

KEQYMKTFGKTEAEYAAMFSKLFKESQLLTLDLTTAKVIKGHIITNGDVPSFFNLWMRHA

QRLLWDMNDHAQKQTAKHLRLKRSEHLNDKEKEKFIEYLNSVRVNKEFQKLNAEEKSQYT

QWIYALWNREAHSFSKEQVAKFAKVKKDQRVGYYSKYIILAVCYIGEDLQSATEIFDVRN

IFKLRKDFYALKSEQEIIQELNTYNTTESRQQIGNEELDLKINHLREAVVSNAVGVVDSL

YKQYKEKTGGEGLIIKEGFDTNKTKEDLEKFSGNIYRILERKLYQKFQNYGLVPPIKSLM

AVRSEGIKNNKGAILRLGNIGFIDPEETSQNCPVCATKFGHHQDVFVCQTCSFTPEKIMH

SNDGVAGYNIAKGGFNNFLNPAPLIKKEEQKAANSNTKQYSNKNNTPRQKTAMEILLEKA

GLKNKNNE

>MBX2887915.1 hypothetical protein [Ferruginibacter sp.]

MENYKLTKAIRFRLEPNEGSNLIQQQVEDMKNISFDLVSFISGLGNFNDLLKRYLLRARK

DGLIKMNSSLSIKSGWLKRYAKQIFHDSEALRDKTRRKQTKTIGDFEGLSAEIEGRLETY

LAIYQSLVNDGVAALNERARKANTGLLLKQLGNQNGLSYFVGLIEASVDKREVDDLSTQL

KKQGKALQEELLAGIQKYLPEQSTGFPLAKASFNYYTINKKPVDYAQKIREQEGKLRITD

WDAQVWSNSGLSNAIKAAVQWGITDLGSLRQKLKNIKAQQKASFNELMTQDLNFQELREN

ADLYLFSNITEHEFNQYKNLTEKIEDKATRINQSSNDNDKRQLRSDLQKLKKDRGALINA

ADKRNQNSFKTYKYFANYYRNIALKHGRILSTIKGIEKEAVESQLLQYWSLVLEQKGKHQ

LLLIPKQKAAECYQFVSAASAIQQSSTSKIYWFESFTLRSLRKLCFGYLDAGTNTFNQNI

QEILRRHQYSEMDNRGRQNQVRIEGEFSFKGNTQKIIACYKDVLASRYTQSVLRLPVRQL

RDEVIEPNFETLDDFQIALERICYKRYALLSEQNLDTITRRYQAQVFAITSADLEKGNDG

NDKAHTQIWKKFWAQENEQTYFDVRINPELSIQWRPAKESRVHKYGRDSALYDEQKNNRY

LHPQFTLVTTISEHSNTPTKVLSFMTDEEFKISVDAFNSKFNNADFKFALGLDNGETELA

TLGVYLPAFKQESNEASIAEIRKIEAYGFKVLSITDLTHQEKDTNGKERKIIQNPSYFLV

KEQYMKTFGKTEAEYAAMFSKLFKESQLLTLDLTTAKVIKGHIITNGDVPSFFNLWMRHA

QRLLWDMNDHAQKQTAKHLRLKRSEHLNDKEKEKFIEYLNSVRVNKEFQKLNAEEKSQYT

QWIYALWNREAHSFSKEQVAKFAKVKKDQRVGYYSKYIILAVCYIGEDLQSATEIFDVRN

IFKLRKDFYALKSEQEIIQELNTYNTTESRQQIGNEELDLKINHLREAVVSNAVGVVDSL

YKQYKEKTGGEGLIIKEGFDTNKTKEDLEKFSGNIYRILERKLYQKFQNYGLVPPIKSLM

AVRSEGIKNNKGAILRLGNIGFIDPEETSQNCPVCATKFGHHQDVFVCQTCSFTPEKIMH

SNDGVAGYNIAKGGFNNFLNPAPLIKKEEQKAANSNTKQYSNKNNTPRQKTAMEILLEKA

GLKNKNNE

>MBS1614876.1 hypothetical protein [Bacteroidetes bacterium]

MENYKLTKAIRFRLEPTEENNLIQQQVDELKNISFDLVSFISKLGNFKDSLKQYLLHDHR

NGTKQVKSGLSIKSEWLKRYAKQIFHDSEAERDKTKRKQTKTIGDFGSLAKDIEGRLEEL

ENTYQELTDDAIAQLNERARNAHSGLLLKRLGSRNGLSYFVDLVENSVDKHEEGDLSIQL

KSQGKALHLELLAGIKKFLPEQSTGVPIAKASFNYYTINKMPVDYAQKVREQQSKLLITD

WNGQIWSGKSYSREIKDDVQAGGSDLEALRQKLKNIKAQQKARFNELMTQNLSFQELKNN

AELYLFSSISEQEFNSYKDLTGIIEKEATKLSQASDDSQKLQLRSNLQRLKKERGALINA

ADKRNQNSFKTYKGFADLYRNIALKHGRILSTIKGIEKEAVESQLLQYWSLVLEREGRHQ

LLLIPKQFAAKCYAFVCNAPVCQQTSASKIYWLESFTLRSLRKLCFGHLDAGTNTFNKNI

QSVLRYHSYSETDNRGQNKQVNIEGEFSFKGDARKIIAFYKDVLASRYTQPVLKLPIRAL

REEVTEKNFETLDEFQIALEKICYKRYALLPEQILDTIIRRYQPQVFDITSTDLTKGESD

NEKRHTQIWRRFWTNENEQQNFDIRINPELAILWRPAKESRVHKYGKESALFDENKHNRY

LHPQFTLVTTISEHSNTPTKELSFMTDEEFTHSVEGFNNKFKKEDFKFALGIDNGETELA

TLGVYLPVFKQNTHEESIQEIDKINEYGFKVLSITNLSHSEADKNGKDRKIIQNPSYFLS

KEQYTRTFEKTETEYQTMFSKLFKEDSLLTLDLTTAKVIGGHIITNGDVPTYFNLWMRHA

QRMLWDMNDHAEKQTAKQITLKKSEDLNQAEKEKFIEYLNLTKNNKKYFSLSETEKTEYT

NWIYAIWNRQTTNLPAEKNQKYETVKKDQRVGYYSKHIILAVYYIGEDLQSVTEIFDVRN

IFKLRKDFFAINTEQEIIQELNNYNTNENRQQIGNEELELKINHLRAAVVANAIGVIDSL

YHYYKSKTQGEGLVIKEGFDTNKVAEDLGKFSGNIYRVLERKLYQKFQNYGLVPPIKSLM

TVRSEGLAGENNKDAIIRLGNICFISDFQTSNLCPSCGRNNPGHIDPYICNNCNFNSTGL

MHSNDGVAGYNIAKRGFYNFLNPAPHIIKEVKKSSEQPHKNTNNNRPQNNNKLSQHKTDL

EIALEKAGFKNKR

>MBS1630098.1 hypothetical protein [Bacteroidetes bacterium]

MENYKLTKAIRFRLEPTEENNLIQQQVDELKNISFDLVSFISKLGNFKDSLKQYLLHDHR

NGTKQVKSGLSIKSEWLKRYAKQIFHDSEAERDKTKRKQTKTIGEFGSLAKDIEDRLDEL

ENTYQELTDDAIAQLNERARNAHSGLLLKRLGSRNGLSYFVDLVENSVDKHEEGDLSIQL

KSQGKALHLELLAGIKKFLPEQSTGVPIAKASFNYYTINKMPVDYAQKVREQQSKLLITD

WNGQIWSGKSYSREIKDDVQAGGSDLEALRQKLKNIKAQQKARFNELMTQNLSFQELKNN

AELYLFSSISEQEFNSYKDLTGIIEKEATKLSQASDDSQKLQLRSNLQRLKKERGALINA

ADKRNQNSFKTYKGFADLYRNIALKHGRILSTIKGIEKEAVESQLLQYWSLVLEREGRHQ

LLLIPKQFAAKCYAFVCNAPVCQQTSASKIYWLESFTLRSLRKLCFGHLDAGTNTFNNDI

QSVLRYHSYSETDNRGQNKQVNIEGEFSFKGDARKIIAFYKDVLASRYTQPVLKLPIRAL

RKEVTEKNFETLDEFQIALEKICYKRYALLPEQILDTIIRRYQPQVFDITSTDLTKGESD

NEKRHTQIWRRFWTNENEQQNFDIRINPELAILWRPAKESRVHKYGKESALFDENKHNRY

LHPQFTLVTTISEHSNTPTKELSFMTDEEFTHSVEEFNNKFKKEDFKFALGIDNGETELA

TFGVYFPVFKQNTHEESIQEIDKINEYGFKVLSITNLSHSEADKNGKDRKIIQNPSYFLS

KEQYTRTFEKTETEYQTMFSKLFKEDSLLTLDLTTAKVIGGHIITNGDVPTYFNLWMRHA

QRMLWDMNDHAEKQTAKQITLKKSEDLNQAEKEKFIEYLNLTKNNKKYFSLSETEKTEYT

NWIYAIWNRQTTNLPAEKNQKYETVKKDQRVGYYSKHIILAVCYIGEDLQSVTEIFDVRN

IFKLRKDFFAIKTEQEIIQELNNYNTNENRQQIGNEELELKINHLRAAVVANAIGVIDSL

YHYYKSKTQGEGLVIKEGFDTNKVAEDLGKFSGNIYRVLERKLYQKFQNYGLVPPIKSLM

TVRSEGLAGENNKDAIIRLGNICFISDFQTSNLCPSCGRNNPGHIDPYICNNCNFNSSGL

MHSNDGVAGYNIAKRGFYNFLNPAPHIIKEVKKSSEQPHKNTNNNRPQNNNKLSQHKTDL

EIALEKAGFKNKR

>MBS1747704.1 transposase [Bacteroidetes bacterium]

MNIKLLNHSIMENYKLTKAIRFKLENSSENNLILQSVEQIKNTSFDLVAFVSDLHNFIDG

LRGYLFFKKNNDTLSTKQGLSIKTAWLKLYAKQAFHDSEAVRDKSRRKQTKTISDYENLT

FEIQKRFDEFEQIYKSLADIANTPLNARSRHAEIGLLLKKLANKHNLPYFVDLVENTVDK

KEEGDLSIQLKSLGKQLINNSLLGVQKYLPEQSSGLLIAKASFNYYTLNKKPIDYAKKKE

EQLNKLKITDWNCQDFVRSSLSNVIKEEIKKGVTDLDELRQKLKNVKAQQKATFNELMTQ

DISYENLKENTELYLFNNISKQEFDSYKKQTEDIEAKATKLNQCTNDNEKRRLRSDLQKL

KKDRGAMMNAADRRFSSNFTVYKVYANLYRDIALKQGRINATLKGIEREQVESQLLQYWA

LVLADAGGHKLVLIPKKHAAECYQLLTTANQNAVTEPNSIFWFESFTLRSLRKLCFGNLE

AGTNSFNENIQSVLREHSYSEIDNRGQNKQVRIEGEFSFKGDTQKIINFYKDVLGSRYAQ

SVLNLPSTQLRVEVLDVNFENLDDFQIALEKVCYKRFALVAHSVLGIIEKQYNPQIFDIT

TYDLRKAVESNDKAHTQLWKEFWTTANEQNNFDIRINPEIAIQWRNAKESRVHKYGKDTA

LYDEKKNNRYLHEQLTLVTTISEHSNTPTKVLAFMTDDDFETSVNAFNEKFNKDDFKFAL

GLDNGETELATLGVYLPAFKQESNEASIAEIKKIEEHGFKVLTITNLSHQEIDKNGKERK

IIQNPSYFLLKEQYTRTFGKTDEEYAAMFSKLFKQTQLLTLDLTTAKVIQGHIVTNGDVP

TYFNLWMRHAQRMLWDMNDHAEKQTAKQISLKKSEELNQIEKEKFIEYLNVTKNNKKYFA

LSKAEKTEYTNWIYAIWNRQTTNLPAEKNKKYESVKKDQRVGYYSKHVILAVCYIGEDLQ

SVTEIFDVRNIFKLRKDFYAIKTEQEIIQELNNYNTNESRQQIGNEELDLKINHLRAAVV

ANAIGVIDSLYHYYKDKTQGEGLVVKEGFDTNKVAEDLGKFSGNIYRVLERKLYQKFQNY

GLVPPIKSLMAVRSKGLKDNKGVILRLGNIGFIDPAGTSQNCPVCNTKFGNHQNNVVCEN

CGFSTQNIMHSNDGIAGYNIAKRGFENFSNPSLSTTNSNRAENNRNRSNNLPNNRNESQP

LTALGAALAAAEAKRNKKRK

>MBS1644463.1 transposase [Bacteroidetes bacterium]

MENYKLTKAIRFKLENSSENNLILQSVEQIKNTSFDLVAFVSDLHNFIDGLRGYLFFKKN

NDTLSTKQGLSIKTAWLKLYAKQAFHDSEAVRDKSRRKQTKTISDYENLTFEIQKRFDEF

EQIYKSLADIANTPLNARSRHAEIGLLLKKLANKHNLPYFVDLVENTVDKKEEGDLSIQL

KSLGKQLINNSLLGVQKYLPEQSSGLLIAKASFNYYTLNKKPIDYAKKKEEQLNKLKITD

WNCQDFVRSSLSNVIKEEIKKGVTDLDELRQKLKNVKAQQKATFNELMTQDISYENLKEN

TELYLFNNISKQEFDSYKKQTEDIEAKATKLNQCTNDNEKRRLRSDLQKLKKDRGAMMNA

ADRRFSSNFTVYKVYANLYRDIALKQGRINATLKGIEREQVESQLLQYWALVLADAGGHK

LVLIPKKHAAECYQLLTTANQNAVTEPNSIFWFESFTLRSLRKLCFGNLEAGTNSFNENI

QSVLREHSYSEIDNRGQNKQVRIEGEFSFKGDTQKIINFYKDVLGSRYAQSVLNLPSTQL

RVEVLDVNFENLDDFQIALEKVCYKRFALVAHSVLGIIEKQYNPQIFDITTYDLRKAVES

NDKAHTQLWKEFWTTANEQNNFDIRINPEIAIQWRNAKESRVHKYGKDTALYDEKKNNRY

LHEQLTLVTTISEHSNTPTKVLAFMTDDDFETSVNAFNEKFNKDDFKFALGLDNGETELA

TLGVYLPAFKQESNEASIAEIKKIEEHGFKVLTITNLSHQEIDKNGKERKIIQNPSYFLL

KEQYTRTFGKTDEEYAAMFSKLFKQTQLLTLDLTTAKVIQGHIVTNGDVPTYFNLWMRHA

QRMLWDMNDHAEKQTAKQISLKKSEELNQIEKEKFIEYLNVTKNNKKYFALSKAEKTEYT

NWIYAIWNRQTTNLPAEKNKKYESVKKDQRVGYYSKHVILAVCYIGEDLQSVTEIFDVRN

IFKLRKDFYAIKTEQEIIQELNNYNTNESRQQIGNEELDLKINHLRAAVVANAIGVIDSL

YHYYKDKTQGEGLVVKEGFDTNKVAEDLGKFSGNIYRVLERKLYQKFQNYGLVPPIKSLM

AVRSKGLKDNKGVILRLGNIGFIDPAGTSQNCPVCNTKFGNHQNNVVCENCGFSTQNIMH

SNDGIAGYNIAKRGFENFSNPSLSTTNSNRAENNRNRSNNLPNNRNESQPLTALGAALAA

AEAKRNKKRK

>MBR4156861.1 hypothetical protein [Bacteroidales bacterium]

MDFIKTTKAVRFRLESNNENTLIQESINNLNSRKEFDLNTFVDDLDAFINDCNAFLFCSK

NKGRREIFYVNPSLIVKNEWLKKYAKQDLAELKQNHTAQRVQYKIGDIDGLCYRIQDLID

DLDDIYVKLCDDASAELHERAKRAQTALLLKRLFANNALPCLVSLIDNTVDKNEKDNLSL

KLKSLGKKLLAQLELGIQEYLPEQSSGVNIAKASFNYYTINKKPIDYDRKIEELSDKLVT

TLDFWKRDGSCNFNKSLWKLIEVKSEGKTLYLGDSPLSDTDEYASLRQILKNILAEQKAE

FSEKMQEKISYEDLTKSDLFLFNNISKEEYNGYLELTNQIEELATDINQEDNEYKLKKLR

SDLMKLKKNRGSLINAADRRTKEKFKTYKSFADFYRKVSQRHGKILAQLKGIEKERSESQ

LLQYWALMLEVNNQHKLVLIPKDKAQECKSRLESSNEQAQGTKLYWFESFTFRSLQKLCF

GNLENGSNSFYPGIRKELQYKYSTEDRNGYPQFISGEFEFKGDEQKKIQFYKDVLNTKYA

QSALSFPKEEVKRNIIEKDFESLDDFVIALEQICYQRYVCVNSHMINALGSYFNAQILDI

TSLDLRNPLNSQEKETVYAHADKKHTEIWKKFWTADNEKDNFDIRLNPEITITYRKPKES

RIAKYGVESDKYDANKKNRYLHDQLTLVTTISEHSNSPAKNLAFTTDAELKDMIERFNAE

IKKEKIKFALGIDNGEVELSTLGVYLPGFKKDTKEEVFAELKKVDEYGFKVLEIRNLRYS

ENDYNGKERRIIQNPSYFMNKELYCRTFNKTAAEYDAMFAEVFEEKQLLTLDLTTAKVIN

GHIVTNGDVISLFNLWMRHAQRSIYEMNDHAIKETANDIVLKRSETLNDAEKRKFIDYLN

GKNKKYEDLSEREKSEYVKWVYRIWGGDYSEYGKNKAFAEISKGQRVGDYLNNVLVAVTF

TGKELTNVVDIFDIRNVFKFKEDFYSLKSETEIMEEVNKYNVKNTKSISNEELDLKLNQL

KSSLVANVVGVIDFLYKQYKERFGGDGIIVKEGFDSAKVESDREKFSGNIYRLLERKLYQ

KFQNYGLVPPVKNLMLMRDVDLNDTNEFMQLGNICFVGYEGTSQNCPVCEKGRLGHTEKC

SDNCGFESKGIMHSNDGIAGYNIAKRGFNNFMRK

>MBR2051365.1 hypothetical protein [Bacteroidales bacterium]

MKQIKTTKAIRFKLENDDKNTIINNSITNLNSRRDFDLNLFVDDLDTFITDCNDYLFCFK

KKGCREIFYVNPNLIVKNEWLKKYAKQDLAKLKQNHTAQRIQYKIGDIDGLNLRIQDLID

DIDDIYTKLCDDASAELHERAKRAQTALLLKRLSANNSLPCLVSLINNIVDKNEKDNISL

KLKSSGKKLLSQLEIGLQEYLPEQSSGVAIAKASFNYYTINKKPIDYDRKIEELSQNMST

SLDEWKNKWQFNEGLWELIEIKSEDKTLYLGDSPFSDDGEYASLRQILKNILAEQKARFS

EMMQDNVSYEELKDSDLFLFNNISEDDYNKYKDLTDRIENVATDINQEDDESKKKDLRSD

LMKLKKERGSLINAADKKTKDRFKTYKTFADFYRKVSQRHGKILAQLKGIEKERTESQLL

QYWALILEVNNQHKLVLIPKEKAQKFKSRLTKSNDKKSTIKLYWFESFTFRSLQKLCFGN

IENGSNTFYPRIKKELQYKYSTTDRNGYPQFIAGEFEFKGDEQKKIQFYKDVLNTKYAQS

ALSFPVNEVKRNIIDKDFETLDDFIIALEQICYQRYVSVNPHLINALSDYFNAQILDITS

LDLRNESNVKDKDVAYSHDDKIHTKIWRQFWSVDNEKNGFDVRLNPEITITYRKPKESRV

IKYGNDSELYDANKKNRYLHDQLTLVTTISEHSNSPAKNLSFTTESELKDMIERFNSEIK

KEKIKFAFGIDNGEVELSTLGVYLPDFKQDSNEATFAELKKVDKYGFKVLEIKNLLYSEK

DYNGKERKIIQNPSYFLNKELYCRTFKKTVAEYDMMFAEVFEEKQLLTLDLTTAKVISGH

IVTNGDVVSLFNLWMRHAQRNIYEMNDHANKETAKNIILKRSETLNETEKRKFIDHLNEN

NKKYERLSEAEKVRYVKWVYDIWNGDFSEDGNNQTFADISKGQRVGDYLNNVLMAVVHEK

GQIESVIDIFDIRHVFKFRKDFYSLKSETEILEEINKYNVKSISNEELDLKLAQLRSSKS

ETEILEEINKYNVKSISNEELDLKLAQLRSSLVANVVGVIDFLYKQYKERFGGEGIIVKE

GFNSNKVESDREKFSGNIYRSLERKLYQKFQNYGLVPPIKNLMLVRNDDLKTANEFMHLG

NICFVGYEGTSQNCPVCEEGRLRHTERCSNNCGFESKGIMHSNDGIAGYNIAKRGFINLN

DNHGCSQQ

>PKP47251.1 hypothetical protein CVT95_05895 [Bacteroidetes bacterium HGWBacteroidetes12]

METYKVTKTIRFKLEAQNVPEIQKDIEGLQSEFNLANFVSDLKNYLDQVRNYLFSEGKEH

VFVNNKITIKREWLQNHAKQEWVDFLEKNKRNNSLNNHTRRIQMSIGDFTGLASKIEGTF

DEINSICKGLADAAGAQANKRTKRERTGLLLKRLQARKALPSLFSLIENSADKNETGNLS

LQLKNKSILIEQQLAAGVQTFLPAQSGGLPVAKASFNYYTINKKPVDFGNEKSELESRLK

ISIDTIFKLTRENFSKKIEEAITADIQKELNNGKTLLLGDVPMLGIENYVSLRQILKNIK

SNQKKAFSDLMQSGKNYNELKATNLYLLNTIEQRQFDNYKVKTNELEKLAVKINQATNDN

QKKELISNKQRVAKQRGIIMRDNFATWKSFSNFYRTISQEHGKILALLKGIEKERTESQL

LKYWALILENNGQHKLILIPREKAASCKQWIASLNPSGDKLTKLFWFESLTYRSLQKLCF

GFTENGNNKFNKNIQNLLPKDNSRKIINGEFAFQGDEQKKIKFYQSVLESKYAQSVLNIP

IQQVQADIINQSFASLDDFQIALEKICYRLFAVVEANIEAELLKNDKAQIFNITSSDLRK

EAKDKIKSHTQIWKAFWTSENKQNNFETRLNPEITITYRQPKQSKIDKYGERSQKNNRYL

HAQYTLITTISEHSNSPTKILSFMSDDEFKSSVDTFNKKFKKDEIKFAFGIDNGEVELST

LGVYFPAFDKTTYKEKVAELEKVNDYGFEVLTIRNLNYKETDYNGKERKIIQNPSYFLKK

ENYLRTFNKSETAYQKMFTEQFEKKKLLTLDLTTAKVICGHIVTNGDVPALFNLWLKHAQ

RNIFEMNDHIQKETAKKIVLKNQLDTDNEKLKFAEYISKEKEFGKLNDDEKMKYTKWIFE

DRDQNNFTEVENKKFKRCQKIYGNYSTKAKAPVLFASCFIDEELQSVTDIFDVRHIFKKR

EDFYALKTEEEIKQLIDSYNTNRASHDISNEELDLKILNTKKALVANAVGVIDFLYKHYE

RRLGGEGLIIKEGFGTGKVEDGIEKFSGNIYRILERKLYQKFQNYGLVPPIKSLMAVRAN

GIENNKNAILQLGNVGFIDPAGTSQECPVCIEGRLEHTTTCPNKCGFNSERIMHSNDGIA

SFNIAKRGFNNFVKSKTDKQ

>HAG50354.1 hypothetical protein [Deltaproteobacteria bacterium]

LGSFISQFNEYIFYESRNNEFIIKGNLTVKNIWLKQYAKQEIVGLKLKRGQTIGDIKGLS

DKIRKARDEVNDIYLKLCDETSKLNERANRAKKGLLIKRLNARNSLPLLLSLIENTTDKN

ETGNLSIQLKQLGAKLQSQLDSGVNMYLPAQSNGLPIAKASFNYYTINKKPIDYDYGQKK

QEQIDKLEINLDSNFDSLIPQHINISNNLKQAIKSDISDIAKKGKDNTLLLGDAPFVNHD

YVSIRQILKNIKAEQKKIFSEFIPRDDSTFKNLKTNSRLYLFNDISGEEFNDYKHKTRVI

EEKAEKKNQCNNDELKRELNSELQNLKKERGSLINAADRNTKGKFKTYKAFANIYGKVAR

DHGRILSTLKGIEKEQVESQLLKYWAVVIEENGQHSLALIPKKNAEEFKKWLENNASTSS

VSPIKTYWFESLTLRSLRKLCFGYIENQTHSNTFYPELKKSPELENYKDDRGDFIRGEFF

FEGDEQRLIKFYKDVLGSQHAQKVLSFPKQQVKDDIIDKQFNTLDEFQIALEKVCYRRFV

LCSSDIKLKLKEYEAEIFGITSLDLRNATKTNIKNHTLIWRMFWNTKNESNNFDIRLNPE

ITLTYRSPNSGRIEKYGKDSELYDERKNNRYLHPQYTLVTTISEHSNAPTKIMSFMTDDE

FKTSVDAFNQKLKKEDVKFALGIDNGEAELSTLGIYLPVFNKDSNSEIWKLFKNVSEYGF

KFLTIRNLSYSEIDANERKRKIIQNPSYFLNKNLYMGIFGRTEQDYDQMLEKQFEEKSLL

SLDLTTAKVINGHIITNGDVPTFFNLWMRHAQRNVWDMNDHSKYKTAKKITIKNNDELTE

AEKAKFAEYISDGKKYEPLTAKEKSKYLKWIFENRNLLQFTDEEKKKFDNCQRRKDNFSK

NILFAVCSIGTDIYSVTDVFDVRHIFKKRKNFDVLKSEAEIIKEIESYNTNKSIRKISNE

ELDLKINHLKQSVVANAVGVIHFLYNYYKEKTGGEGLIVKEGFDTKTVADGLEKFSGNIY

RILERKLYQKFQNYGLVPPIKSLMAVRGGGIKDNGLSKTIRLGNVCFVSQSKTSGICPLC

NSKKLDHNTTCPDCQSDLIEIFHSNNGVAGYTIAKRGFENFNNNSNKGVSEK

>MBQ8223513.1 hypothetical protein [Bacteroidales bacterium]

MEQIKTTKAIRFKLESNDNNNLVIESISNINSNREFDLDIFVDDIDNFINYCNDFLFVSK

KIKGKDVFFVNRMLIVKNEWLKVYAKNEFAELKHNNNAQRVQYKIGDIDGLDARIEDLIN

EIEGIYNALVDDASAELHERAKRTQTALLLKRLYSNNSLPCLVSLIENTVDKNERGNLSL

KLKSMGKKLLKQLELGIQEYLPEQSAGVTLAKASFNYYTINKKPIDYEKKIDELSNQLKT

SVEDWKNLKNREGRPMFRFDDALWKLILVKAQNKQLYLGDSPFAEVNEYASLRQILKNIM

AKQKAEFSEMMQDGASYRELKNSDLYLFNSISEDEYIKYFDLTEKIEEFATKVNQCSDDN

KKKNLRSELNKLKKKRGSLINAAERNTSNYFKTYKSFAELYRKVSQKHGKILAQLKGIEK

EKSESQLLKYWAMILQDNNGHKLVLIPKDKIKDCKSRLTESIANNGRIKLYWFESFTFRS

LQKLCFGNLDTNTFYPEIKKELLGKYSEIDGRGYPKFISGEHEFKGDEQKKIQFYKDVLS

TKYVRQNLNLPFNEVKNKVIDKSFACLDDFKIALEQICYKRMLTVDVHLINALQTYFNAQ

ILDITSLDLRNNNETHKDKIHTEIWKQFWSSENERNNFDIRINPEITIIYRKPKESRLAK

YGVGSDKYDPNKNNRYLHEQLTLVTTISEHSNAVSKNLSFTTDEELIDMINDFNAKIRDE

KIKFAFGIDNGEVELSTLGVYLPEFDKPTNDEKIAELKNVEKYGFKVLEITNLLYSEEDY

NDKDRRIIQNPSYFMNKELYCRTFKKTETEFEEMFKNVFEEKHLLTLDLSTAKVINGQIV

SNGDVVSLFNLWMRHAQRNIYELNDHANKKTAKEIVLKKSEELNAGERRKLIDGLNENNK

KYNDLSDYQKEQYVEWNYKRWRFEDVENEDFEKIYKNGQRVGNYLHNVLMAVVHDKGNIE

SVIDIFDIKNVFKFRKDFYSLKSENEILEEINKYNVKSISNEELDLKLNQLKSSLVANVV

GVIDFLYKQYKSRFGGEGIIVKEGFNSNKVASDREKFSGNIYRLLERKLYQKFQNYGLVP

PVKNLMLVRNDDLRDENAFFNLGNICFVGYVGTSQRCPVCETGKLGHTETCSNDCGFESK

GIMHSNDGIAGFNIAKRGLNNFNK

>MBQ7612354.1 hypothetical protein [Spirochaetaceae bacterium]

MQKYKITKTIRFKLEAETIGEQLQVDIQALNKKSEFNLAHFITQLKCFCDDMKKYLFFEK

GNELEVDGKLIIKKEWLRIYAKQALADVTEKEKANLYNREGNRKLRREQYTIDKYKDLKF

TIKEIFDNVDKIYCDIATDASAELNERSRRAHTGLLLQRLSAKRALPCLVSLVENTSKKN

EIDDLSVRLKYFGAKLLQQLTFGTCEYLPAQSNGIPIAKASFNYYTINKKTTDYKSIHKN

IEEKLLVADLKTLIPQMFNLSKAVKESITEDIKAKTDNKQLLFGDSPFADTSTASLRQIL

KNIKAEQKAKFQEFMNNNPSLEKLKSKTNLYLFNNITQAEFDEYAELTKQIEEKGISINQ

CSSESQKRKLRSDMQHLKKLRGKLLKDNFNTYKTFANFYGDVARNHGKLLAQLKGIEKER

TESQLSNYWALILEQDGSHKLILVPKEVASEFRKQLNPIIKNNSEDKITWIESLTYRSLR

KLCFGNLENGTHSNTFNPEIKKEIHMPNGEFEFQGDEQQKIKFYQEVLNTNYAKQVLDLP

QEVNTRIIGKNFDSLYDFIIALEKICYRRFSTVDENTIKELKRIGAQVFNITSLDLRNEK

NSKDKVARYAHTDKMHTQIWKNFWSVDNENNNFDIRLNPEITISYRKPKESRIVKYGKES

KLFDQNKKNRYLHEQFALITTISEHCNTPAKDLSFVSDEDFKKSIDEFNKTLTEDKVKFA

LGLDNGEVELSTLGVYLPQFDKSTNEEKIAELKKTKDCGFQVFKIKNLSYEENDTNGKPR

KIIQNPSYFVKEDIYCRTFGKTHEEYETMFANVFEEKYLLTLDLTTAKVIDGKIIENGDV

ISLFNLWMRHAERNIYEMNDHAKKETAKTIILKKSDELNDEEKAKFIDYLNEGNKKYASL

SSEEKGEYVKWIYDVWNKKIKEDDNDEFKKVKKECKRKGKFIPDKKQDNKDNKVIDIVFA

VCKIGEDIQSVQVVFDIRNVFKLRKDFHSIKSEKEIFDELGKYNKRTISNEELDLQINQR

KESLVANVVGVVDFLYKQYQKRFGGEGIIVKEGFDHGKVAQDREKFSGNIYRLLERKLYQ

KFQNYGLVPPIKNLMLLRSEGVKKDKEKEKDKDETKIIRRLGNICFVHEKGTSQECPVCE

KGRLNHTEVCPESCGFDVKGIMHSNDGIAGFNIAKRGFNNFY

>MBO7227809.1 hypothetical protein [Bacteroidales bacterium]

MNNIKTTKAIRFKLENSASNREIQERIDNLSSNNGFNLVSFVTELNNYIDLLNGYLFCSR

GDRFYLNDKFILKKEWMKNYAKQELAEFKTKIDNTANGVRVQYTVGDYRVESKIQAAFDN

IDEIYGELCDDASRELNERSKRTRTALLLKRLYAKNNLPLLIDFVENSTYKKEVGNDSLV

LKSIGKRLMEHLELGIQEYLPEQSAGVSIAKASFNYYTINKKPIDYDAKIEEIEKKLVVS

NIDRTWNDRDNKIKNNGRLWNIVKADIASRQNNKPLCIGDSPFMDVDEYASLRQIMKNIL

SEQKAAFNELMQQGGSYKELQNSELYLFKEVTKDEFDEYSKLTEDIEELATKRNQTKNER

EKKELKSQIERLSKQRGILISEADGRTANKFRTYKDFSILYRKLAQAHGRQLAKLKGIEK

EKTESQQLTYWAMILEKDNRHKLVLIPKAKSSECRSRLVEADNANGNIKLYWFESFTYRS

LQKLCFGNIENGSNTFYPQIRRELNFRYSIPDNRGNYKFISGEHEFKGEEQRKIQFYQDV

LKTDYARKNLNLPYREVEDRIINSTFESLDDFKIALESICYRRAVCTNIHLIDALESEYG

AQIFDITSLDLRREDNIKDKEEKYSYSYKSHTAVWKEFWTAENEKRNFDIRLNPEISIIY

RKAKESRIEKYGVDSKKNNRYLHDQLTLVTTLSEHCNSPERNIAFATIDEEERIIAEFNS

KLYKENIRFAFGIDNGEVELSTLGVYLSEFKQDSIKSSLKEVKNVDKYGFDTLTIRNLMH

SEKDVNGRDRRIIDNPSYFLKEDLYCRTFGKNGEEYKAMFDKVFEQRRLLTLDLSTAKVI

CGRIVTNGDVISLYNLWLRHAQRNIYDMNEHIEGGGAKVYLKKSEELNDNEKRKFLDYLN

DENERYEKLSDTDKKDYIDWIYQIWDGKDVENKRFAEVRKKQKRPGFYFHNVLLAASYIG

EEVQDVKDIANIDDVRHVFKFREDFKQFKPEKEILDEINKYNIKVISNEELDLRLNQLKS

SLVANVIGVIDYLYKQYKERFGGEGIIAKEGFGIEKVESDRMKFSGNIYRMLERKLYQKF

QNYGMVPPIKNLTAFRSKDKSYTQIGYICFINYDGTSQRCPICNTKLAYNHGLECSAKCG

FDSEGIMHSNDGIAGYNIAKKGFESIVNK

>MBQ8720783.1 hypothetical protein [Paludibacteraceae bacterium]

MENNNVYQVKKAIRFKLQPAESNSVNLSEAIEGNVEFNLPNFVVHLVDFYDDLNEFFFYE

DEDRWLVKEKMVVKKEWLQENEKEAYLKWKREKKDKENKEKVDVSKNKRVQLTIENIDGL

ALRIERAIDNLGMLIDELEDDASRNLEERAKRTRTALLLKRLKTRRMLPYLASIVENVVS

KSERDDLSVRLKRLGRSVMRELELGIQAYLPEQSGGMPVAKASFNYYILNKKPIDFDKEK

QELMSELEIDVDSFVFPQGPKSKKDPILDNAIRNSVRKEAKGKCVLLGDVPMVDFEDYIS

LRQVLKNIKSRQKAKFTEYMNNDPIYEDLREYDSLYLFNRISKDEFDEYLDLTEKIQEVA

TKYGQYKSKEFKSKVEDLRKKRGAYINAADKTKEKYFGDYKKFADIYRRVAQQHGKILSR

LKGIEIEQYESQLLNYWALMMEEGGRHRLVLVPRESASKAKSKVEQKRGVSGGDVKLYWF

ESFTFRSLRKLCFGNLDNGTNTFYPELRNDRDFFRKYSTLDRNGRPQFVQGEFEFKGDEQ

RIIQFYKDVLMTSHAQENLNLPIKLVREQIVNTEFESLDDFVIALERVCYTRFVTTDKDT

INYLKNECNALVFDIKSQDLDNAENTKDKEVKYTHHDKRHTQIWRRFWTTENENDNFDLR

LNPEITILYRKPKPSRVAKYGEGTENYDPTRKNRYLCDQYTLVTTFTDHSNTPQKNLAFA

TDEEVKCVVDEFNAKLAMPSFAFGLDNGETELSTFGVYVPEFDKAINEEVIAELNNVDKY

GFDTLKIKDLLYSEIDKNNNVKRIVLNPSYFVDEELYCRTFGKTHEEYVAMFEAQFEKRR

LLTLDLTTAKVINGHIVENGDVASYFSLCMRNAQREIYEMNDKAKEETARKIMLKRSETL

DAKERREFVNYLNKKNDEYKKLTEEEKNKYIQVLYTHWGGENIEDKWFKKIKKDQRVGYF

ASDVVCGVCYEGNELQTVIDVFDVHNIFKLRKDFYSIRSEQEILDEINEYNQKRVISNEE

LDLKIVNVKRAVVANVVGVVDKLYHEYAERFGGDGVIVKEGFDSKKEEADRNKFEGNIYR

MLERKLYQKFQNYGLVPPLKNLMMLRDGGIKDNRDAIMQVGIVAFVDPAGTSQECPVCVE

GKLRHTTICPNKCGFTSDGIMHSNDGIAAYNIAKRGFNNLKNNR

>MCI6417139.1 transposase [Bacteroidales bacterium]

METIRSNRFIRFKLQADGENASIQNKIDALNVDGEFDLVNFFSKLKNYLYDINDYLFFER

EDGSIFKGGIVVKSEWMKTHAKQQLAEFRYKERSEDNQRNDSRRVKRPRMQHTLSEYKVS

DRIIDVHDDIVDVYEKLQDVVSRERHERADRAKIALLLGRLKTKGGLPLLVSLVENTTDK

KEQGELSLRLKKSGKELLNLLEMGVQSYLPEQSKGYPIAKASFNFYTINKRPIDYTRKIK

EINDRLRVDWKKCEEWCFGKNKSANDIRWWKTIKADVERRAKGKELLLGEMPMRDIEDYA

NLRQILKNILAQQRAEFSEMIQSAESYSSLQESGLYLFDRISDEDFDEYFDFSEAIEQTA

TKRNQTRDQYKQRELRTKINRLKKQRGDLLNDACRETRDRFVDYKRFAAFYRKVAQQHGR

LYAQLKGIEKERNESQMISYWAMILQSGSKHKLVLVPKEKASELKNGLNVSEDKTSNTKL

FWFESFSFRSLQKLCFSNIESGSNKFYSDLRNEAEFSRKYSFGGKFISGEFDLQGDEQKK

IEFYKDVLSSRTAGKMLSISKEELKQEVLDVDFDCLDDFKVALERVSYKRMFTVNPNIID

ALKSDFAAQVFDITSLDLRHEESCKDKETVFEYSDKMHTRIWKEFWSEDNEDSNFDVRLC

PEITLLYRKPKSSRIAKYGADSTQRNRYLHEQLTLVTRFTERSNSPGRELSFLAEEDEKK

IVENFNREVLKEDFRFALGIDNGEVELSTLGVYLPQFAQESNDATFEMLKKVDEFGFPTL

TIRDLKYKEKDIKGIDRRVVQNPSYFIKEDLYCRTFGKSHQEYEAMFETLFERRNLLTLD

LSTAKVISGHIVTNGDVVSLFNLWKRHAQRNIYAMMEHQLAEGSVEIELKRSYELSDNER

QTFIDYLNADNKKYEKLSKTDKSKYVTWIYDCWNGKEVENKDFENVYNECKRKGNFADIV

LYAVCTNAGKVETVVDVFDVRNVFKLRKDFDCLMSQEEIKAELNRYNVRVISDEELDLNL

RQTKSSLVANVIGVIDFLYARYKEKFGGEGLVVSEGFGVSKVEQDLEKFSGNIYRLLERK

LYQKFQAKGLVPPIKNILMFREEDNNRESKKSVPFLHIGNICFVDPRETSQNCPVCENGK

LGHTEICPNNCGFKSEGIMHSNDGIAGYNIAKRGFLEFKKLK

>MCI7571710.1 transposase [Bacteroidales bacterium]

METIRSNRFIRFKLQADGENASIQSKIDALNVDGEFDLVNFFSKLEDYLSGINDYLFFER

EDGSIFKGGIVVKSEWMKTHAKQQLAEFRYKERSEDNQRNDSRRVKRPRMQHTLSDYKLS

DRIIGVYDDIVDVYEKLQDVVSRERHERADRAKIALLLGRLKTKGGLPLLVSLVENTTDK

KEQGELSLRLKKSGKELLNLLEMGVQNYLPEQSKGYPIAKASFNFYTINKRPIDYTRKIK

EINDRLRVDWKKCEEWCFGKNKSANDIRWWKTIKADVERRAKGKELLLGEMPMRDIEDYA

NLRQILKNILAQQRAEFSEMIQSAESYSSLQESGLYLFDRISDEDFDEYFDFSEEIEQTA

TKRNQTRDQYKQRELRTKINRLKKQRGDLLNDACRETRDRFVDYKRFAAFYRKVAQQHGR

LYAQLKGIEKERNESQMISYWAMILQNGSKHKLVLVPKEKASELKNGLNVSEDKTSNTKL

FWFESFSFRSLQKLCFSNIESGSNKFYSDLRNEAEFSRKYSFGGKFISGEFDLQGDEKKK

IEFYKDVLSSRTAGKVLSISKEELKQEVLDVDFDCLDDFKVALERVSYKRMFTVNPHIID

ALRGNFAAQVFDITSLDLRHEESCKDKETVFEYSDKMHTRIWKEFWSEDNEDSNFDIRLC

PEITLLYRKPKSSRIAKYGADSTQRNRYLHEQLTLVTRFTERSNSPGRELSFLAEEDEKK

IVENFNREVLKEDFRFALGIDNGEVELSTLGVYLPQFAQESNDATFEMLKKVDEFGFPTL

TIRDLNYKEKDIKGIDRRVVQNPSYFIKEDLYCRTFGKSHQEYEAMFETLFERRNLLTLD

LSTAKVISGHIVTNGDVVSLFNLWKRHAQRNIYAMMEHQLAEGSVEIELKRSYELSDNER

QTFIDYLNADNKRYEKLSKTDKSKYVTWIYDRWNGKEVANKDFENVYNECKRKGNFADIV

LYAVCTNAGKVETVVDVFDIRNVFKLRKDFDCLMSQEEIKAELNRYNVRVISDEELDLNL

RQTKSSLVANVIGVIDFLYARYKEKFGGEGLVVSEGFGVNKVEQDLEKFSGNIYRLLERK

LYQKFQAKGLVPPIKNILMFREEDNNRESKKSVPFLHIGNICFVDPSGTSQNCPVCENGK

LGHTEICPNNCGFESKGIMHSNDGIAGYNIAKRGFLEFKKLK

>MCI7701238.1 transposase [Bacteroidales bacterium]

METIRSNRFIRFKLQADGENASIQNKIDALNVDGEFDLVNFFSKLEDYLYDINDYLFFER

EDGSIFKGGIVVKSEWMKTHAKQQLAEFRYKERSEDNQRNDSRRVKRPRMQHTLSDYKLS

DRIIGVYDDIVDVYEKLQDVVSRERHERADRAKIALLLGRLKTKGGLPLLVSLVENTTDK

KEQGELSLRLKKSGKELLNLLEMGVQNYLPEQSKGYPIAKASFNFYTINKRPIDYTRKIK

EINDRLRVDWKKCEEWCFGKNKSANDIIWWKTIKADVERRAKGKELLLGEMPMRDIEDYA

NLRQILKNILAQQQAEFSEMIQSAESYSSLQESGLYLFDRISDEDFDEYFDFSEEIEQTA

TKRNQTRDQYKQRELRTKINRLRKQRRDLLNDACRETRDRFMDYKRFVAFYRKVAQQHGR

LYAQLKGIEKERNESQMISYWAMILQNGSKHKLVLVPKEKARELKNGLNVSEDKTSNTKL

FWFESFSFRSLQKLCFSNIESGSNKFYSDLRNEAEFSRKYSFGGKFISGEFDLQGDEQKK

IEFYKDVLSSRTAGKMLSISKEELKQEVLDVDFDCLDDFKVALERVSYKRMFTVNPHIID

ALRGNFAAQVFDITSLDLRHEESCKDKETVFEYSDKMHTRIWKEFWSEDNEDSNFDVRLC

PEITLLYRKPKSSRIAKYGADSTQRNRYLHEQLTLVTRFTERSNSPGRELSFLAEEDEKK

IVENFNREVLKEDFRFALGIDNGEVELSTLGVYLPQFAQESNDATFEMLKKVDEFGFPTL

TIRDLKYKEKDIKGIDRRVVQNPSYFIKEDLYCRTFGKSHQEYEAMFETLFERRNLLTLD

LSTAKVISGHIVTNGDVVSLFNLWKRHAQRNIYAMMEHQLAEGSVEIELKRSYELSDNER

QTFIDYLNADNKKYEKLSKTDKSKYVTWIYDCWNGKEVENKDFENVYNECKRKGNFADIV

LYAVCTNAGKVETVVDVFDIRNVFKLRKDFDCLMSQEEIKAELNRYNVRVISDEELDLNL

RQTKSSLVANVIGVIDFLYARYKEKFGGEGLVVSEGFGVSKVEQDLEKFSGNIYRLLERK

LYQKFQAKGLVPPIKNILMFRENDNNRESKKSVPFLHIGNICFVDPSGTSQNCPVCENGK

LGHTEICPNNCGFESKGIMHSNDGIAGYNIAKRGFLEFKKLK

>MCO5290432.1 transposase [Chitinophagaceae bacterium]

MEKFKITKTIRFKLANSEGSILKDEITKLNNNIESFEVKNFIAELFNLFEKLEAYLFFTE

DNELQVSEHINFKKSWLRLNVKNEYYKFLESKEPNKIISIKVSDIPGLEEIIKQRFDEVR

KQIKKLNECENLDLNQRARDEHISLLIKQIAGRNSIPFFLSLVEHTDKMKEEDDASMYLK

NKAEVIKNTLLVAIQKYLPNQSQGLSILKASFNYYTINKTPIDFQKKKEELEKQLVIKID

NGRIHVQEHKKGKLEYVSLFDKKMNEEVVKIFKKELAKNENAILTIGHNPLSDEKTFSLR

QFLKQIKAEQRKKFEEAICGPDSTYERLKQNPELFLFNNILIGEYNNYKKLTEEIQELAN

QKNNTQDDAKKEQLTSTIKSRKQKRGELLDFQNKDNKQAFKTYKGFIRKYTDVARIHGKT

LALLKSIEKESIESALLSHWALIIENNQQHQLALIPKEKAKLCKEWLETSQTNNSEQDAI

IWFESLQLRSLRKLCFGFVENRTNSFYKAISQSPELKKYTSKNGRGFIKGEYEFEGDEQK

IIQFYKHVLRTKYTESVLKLPFKEMQDTIFVQDFENLDEFKIALERITYKRYVCKPVNYL

EELKKFSPKIFNITSQDLEKAEEKNLKHHTILWKKFWTKENEVNHFAFRLNPEIQITYRL

SKTSRMLKYGHTSSHYDPSKKNRYLHPQFTLVITISENCNSNEPEMNFKTDEELAEQITG

YNQRLSTDFKFALGIDNGVAELATLGVCHPDFLKKSAQERLETFQNINDYGFKTLEIKNP

SYYELYKGKDKRIIAQNPSYFLNKETYKAVFDKTEEEFKAMFANQFKEKTSLSLNLSIAK

VINNQLVSNGDIYTFFNLCLKEAQRLIYEMNDHSKITSSMQVVLKTNEQLNLNEKWAFIE

KNHSAKRLTSLETNEQRNKYIEFVFEGKHNNDEELKDIFKGIQIRRNYTQRAIYAINYME

EDGAISDIKIVFQLKPLFNPIMCEEEIMDKLKEFNVKQISNEELDLKLINLRKAVVSNAI

GVIDFLYKAFQARYGGEGIIVKEHFKPEDVDNKVADFRGNIYRLLEMKLYQKFQNYGLVP

PIKRILSLREEKIGIENIQNFGIIGFVNPQHTSNLCPVCEKNSMHHTTICPENCGFDSMD

KFHSNDGIAGFNIAKRGLEKIRKNQLINKLNKTQ

>MBK7097284.1 hypothetical protein [Sphingobacteriales bacterium]

MENQLTIKITNEQIQIQKFKKGSLEFESPFDKKTNAEIVKIVNEELAKNKNAKIYIGHNP

LSEEMILSLRQILKQIKAQQRKIFEETICNPNSTFEQLKQDPKLFLFNNISIDEYNNYKK

LTKEIQELASQKNNTQEDSKKEQLTSEIKSKTGERGGFLDYQNKDKDKSKLFRTYKDFIR

KYADVARIHGSTLALLKSIEKESVESALLSHWALIVENNKQHQLVLIPKEKAKECKEWLE

NKRENPENDFQSTIFWFESLQLRSLRKLCFGFVENKTNSFYKEISKCGDLQKYISNNGRG

FIKGEYEFKGDEQKIIQFYKDVLSAKYTQSVLKLPFKEINDTIIKKDFENLDEFKIALKR

ITYKRFVCKPTNYLEEIKKFSPKILVITSQDLEKEEVKNFKNHTILWKKFWTKENEVKYF

DFRLNPEIQITYRLPKTSRVLKYGSASSNYDPTKKNRYLNPQFTLVTTISENCNSNEPQL

AFKTDEELNGQIRHYNQRLCTDFKFAFGIDNGVAELAAFGVCHTDFLKSSTQDRLEAFHN

VNDYGFKTIKIKNPSYYELYKGKDKRTIAQNPSFFLNMETYRKVFNKTKDDFIAMYAEQF

EEGISLSLNLSIAKVINNQLVLNGDIYTFFNLSLREAQRLIYEMNDHSKLVSSMQVILKT

NEQLNQNEKWAFIEKNHSAKSLKPLETNEQRNKYIEYVFEGKHNNDEELIKIFNGIQIIR

KYTQKAIYAISYMEGNGAILDIKMVFQSKPLFDCIMSENEIINKLEGFNRKEISNEELDL

KLVNLKKAVVSNAVGVIDYLYKSYKARFDGEGLIVKEHFKPEDVDNKVADFRGNIYRLLE

MKLYQKFQNYGLVPPLKRILSIREKKIDLENIQNIGIVGFVNPQDTSNQCPVCNMNKMRH

TTICPENCGFDSKDIFHSNDGIAGFNIAKRGLGNIRNKI

>MBP6739571.1 hypothetical protein [Leptospiraceae bacterium]

MEKYQITKSISFRLNKVKAPILEEQVEKIGDVKEAEASLLRLFSVGQDLAGLLKQYIYKD

NQKLKSSVTVHFRWLRDYTKDKFYNWKTESRRDFSREKRFKLSEVDYLKEVFEHFISDWD

AIIQNLGSEISRKQEALSRKAEIGKIIKRIGTRNVFLFFENLIIESKDKNEIHLEAQLNE

KIKVFKENLIKAEQWALPAQSAGIELAKASFNYYTINKKPKDFELEKRNLEKQLHNYDER

FMKDIAQRENKLFGELRIIGKDDATNMYRFLYELIQNSEGKQVLEKTDKGLTVDLDSLYK

NLKVFRAEQKKSFQEAVSKGSSYEDTAKINILFSSVDQKNNEKNLKAFNKYKEFTDTIQK

LADQKNKVERDSLNSRKLQEEISRIRQYRGKLLFQGPSKRGEPEERFDKYLKFCKVYKDI

ALKRGRLVARLKGIEKEKIDSQRLKYWTFIVEKDKQHQLVLIPKSQSQKVYQKISSYESL

GAGSSNIYYFESLTYRALKKLCFGVNGNTFIPEISEELRKAGKWNYEERDFGEHIFKIRS

GVEEIRDEKRLIAFYKDVLGTDYIRKNREGGKIVLPSSFDEEVLIPEFSTEQEFHAALEK

CCYLRKKKIPKDDIEKIFSECSAQIFTITSYDLEKQDKTNLKAHTKLWFDFWTGENEAQR

FPLRLNPEIRILWREPKATRVEKYGKDSKLFDPKKRNRYLHPQFTLTTSFIENALHPEIN

YRFLDAKQKKDNVKKFNMDINAKLKPLPKAWYYGIDTGIVELASLCLIQKNGIPQTFEVL

NLKEDKMNYDKTGYLKDGTKKKYKAIQNLSYFLNEDLYKRTFQDESFLDTFAEIFEKKEV

SALDLSVSKIISGHIVMNGDITARLNVALANARRKIVDALIKNPSVQLSEQDYNIKVGDE

IIFKGRIEFNAIKSWEDIQAIIHSSFEKDKENLARVQEDINKYRQVIAANVTGVIHFLYK

KFPGLITIEYLDQSEVESHRKDFEGIIDNPVERALYRKFQTEGLTPPVSELWSIKNKNEQ

IGIIYFADPKDGKKGAHGIRCPKCGEKAYPRSEDDGYKKDKNNKIFQCKQCGFHNLNNSM

GLLGLDSNDKVAAYNIAKRGLEILK

>KFO67988.1 hypothetical protein ER57_07110 [Smithella sp. SCADC]

MEKYKITKTIRFKLLPDKIQDISRQVAVLQNSTNAEKKNNLLRLVQRGQELPKLLNEYIR

YSDNHKLKSNVTVHFRWLRLFTKDLFYNWKKDNTEKKIKISDVVYLSHVFEAFLKEWEST

IERVNADCNKPEESKTRDAEIALSIRKLGIKHQLPFIKGFVDNSNDKNSEDTKSKLTALL

SEFEAVLKICEQNYLPSQSSGIAIAKASFNYYTINKKQKDFEAEIVALKKQLHARYGNKK

YDQLLRELNLIPLKELPLKELPLIEFYSEIKKRKSTKKSEFLEAVSNGLVFDDLKSKFPL

FQTESNKYDEYLKLSNKITQKSTAKSLLSKDSPEAQKLQTEITKLKKNRGEYFKKAFGKY

VQLCELYKEIAGKRGKLKGQIKGIENERIDSQRLQYWALVLEDNLKHSLILIPKEKTNEL

YRKVWGAKDDGASSSSSSTLYYFESMTYRALRKLCFGINGNTFLPEIQKELPQYNQKEFG

EFCFHKSNDDKEIDEPKLISFYQSVLKTDFVKNTLALPQSVFNEVAIQSFETRQDFQIAL

EKCCYAKKQIISESLKKEILENYNTQIFKITSLDLQRSEQKNLKGHTRIWNRFWTKQNEE

INYNLRLNPEIAIVWRKAKKTRIEKYGERSVLYEPEKRNRYLHEQYTLCTTVTDNALNNE

ITFAFEDTKKKGTEIVKYNEKINQTLKKEFNKNQLWFYGIDAGEIELATLALMNKDKEPQ

LFTVYELKKLDFFKHGYIYNKERELVIREKPYKAIQNLSYFLNEELYEKTFRDGKFNETY

NELFKEKHVSAIDLTTAKVINGKIILNGDMITFLNLRILHAQRKIYEELIENPHAELKEK

DYKLYFEIEGKDKDIYISRLDFEYIKPYQEISNYLFAYFASQQINEAREEEQINQTKRAL

AGNMIGVIYYLYQKYRGIISIEDLKQTKVESDRNKFEGNIERPLEWALYRKFQQEGYVPP

ISELIKLRELEKFPLKDVKQPKYENIQQFGIIKFVSPEETSTTCPKCLRRFKDYDKNKQE

GFCKCQCGFDTRNDLKGFEGLNDPDKVAAFNIAKRGFEDLQKYK

>OGX23684.1 hypothetical protein A2Y03_07010 [Omnitrophica WOR_2 bacterium GWF2_38_59]

MKNGINLFKTKTTKTKGVDMEKYQITKTIRFKLLPDNAHEIVEKVKSLKTSNVDELMDEV

KNVHLKGLELLFALKKYFYFDGNQCKSFKSTLEIKARWLRLYTPDQYYLKKSSKNSYQLK

SLSYFKDVFNDWLFNWEESVSELAIIYEKYKICQHQRDSRADIALLIKKLSMKEYFPFIS

DLIDCVNDKNSNKTFLMKLSEELSVLLEKCNSRALPYQSNGIVVGKASLNYYTVSKSEKM

LQNEYEDVCQSLDKNYDITEMKVILYKEKLDNLNFKDVTIANAYNLLKENKALQKRLFSE

YVSQGKVLSLIKTELPLFSNINDNDFEKYKEWSNEIKKLADKKNTFCKKTQQDKIKDIQN

KISELKKKRGALFQYKFTSFQKHCDNYKKVAVQYGKLKARKKAIEKDEIEANLLRYWSVI

LEQEDKHSLVLIPKNNAKDAKQYIETINTKGGKYIIHHLDSLTLRALNKLCFNAVDIEKG

QMVRENTFYQGIKEEFERNKINCDNQGVLKIQGLYSFKTEGGQINEKEAVEFFKEVLKSN

YAREVLNLPYDLESNIFQKEYTNLDQFRQDLEKCCYALHSKIGKDDLDEFTRRFEAQVFD

ITSIDLKSKKEKTKTTGEMKKHTQLWLEFWKGAIEQNFATRVNPELSIFWRAPKSSREKK

YGKGSDLYDPNKNNRYLYEQYTLALTITENAGSHFKDIAFKDTSKIKEAIKEFNMSLSQS

KYCFGIDRGNAELVSLCLIKNEKDFPFEKFPVYRLRDLTYQGDFKDKHDQMRYGVAIKNI

SYFIDQEDLFEKNNLSAIDMTTAKLIKNKIVLNGDVLTYLKLKEETAKHKLTQFFQGSSI

NKNSRVYFDEDENVFKITTNRNHNPEEIIYFYRGEYGAIKNKNDLEDILNEYLCKMETGE

SEIVLLNRVNHLRDAISANIVGILSYLIDLFPETIVALENLAKGTIDRHVSQSYENITRR

FEWALYRKLLNKQLAPPELKENILLREGDDKIDQFGIIHFVEEKNTSKDCPNCRKTTQQT

NDNKFKEKKFVCKSCGFDTSKDRKGMDSLNSPDTVAAYNVARKKFES

>MBR4264557.1 transposase [Bacteroidales bacterium]

MDNTKYQITKTVRFKLEPMFDEKKLVKDVKTNEDKRNTDNVSLLDDFITLIKGVVTSFKN

CVFFHNVDPQTGECFKPYDDESKYKKDNTIYVIDKATNLHVWNKLLEVKYTWLRRYMQDE

FYSNRAETIGKLPKSYGVNQLEYLKKYFELDWLIKIEEVISNLEEIKNKPQENHSRNAAV

AVEISKLKRRTYFEFMKTFVFALTSTNKPDIDNLVKKIKNELDDCESKLNNIYSYFLPLQ

SKGAELLSGSLNYFTVNKKTTDFDEDEQAANNFLDGDIDLSVYPNFNEWKTLPLSGKSKD

EAYWIVKEWKAKNKSKFLECLRNENYGEAFGIEPFTHTQTYDFESLKRETRNQRLCSDIL

SIKNFSVDKLTKEQQTFITENIATLNLDSDSDEVVTEAVKQKNKEYKHLISEFFKIEDAS

NQFEIKTYNCIPAYEQLCNDYKQLALKRGQSIAKLAAIDKERQTIERLKYWSLIVDCNNE

QFLYMIERTEEDKIQKAYKHISDLPTIKNDTEVKYRIHFFESLTLSALRKLCFNKVNKNF

IKAVKKNASKNEDDYNKLNVQYEQAMETDKDKINFYKCVLTYTSTLNTQSFELQSLLKGN

YNTLNDFEIALNTICYSKTIKTDVDIDSFLVNDCKALKFKLTSLDIKNNKVFEAYYAKNE

AEKYINHKSNTMTKLWKKFWSNKNLESNFEVRLNPELRVFWRDAKQSRIEKYGADSKKNN

RYLRPHFTLALTMCENANSQKINYAFKSTEDKGHEIKAFNGEINKTICKNFALGIDAGSI

ELATLSVVERKNNLLLPGLFDVYRIIYSQRDYEKEGYLKDGTPKQYKLILNPSYYLNRDL

YIKTFFKKNDATSLEVKIDDFENTKKLLFEKVSVASIDLTTAKVISGVIVTNGDYKTFEN

LRILHAQRIINQVLRKDLTALLEPHQGEKAKDCYIKLKSKVYDGTIYHWRGNHGVITSFE

DAKVKLTNNFNEYKDFYSDPNGYMRSHKDKFLFDETRLDDNINNTRRSMVANMVGVIKYV

YDQFKEKYGLGHICMENFDPITYESHRKEFEGDVLRPLEIALYKKFQNDCIVPPIGELLK

QRSYTTKDKDGFTQFGIIKFVDKEGTSSTCPKCGKKAYANDDQLADAKLNKGEFRCPHCG

FVVYKNNNKQSDRCTDMFGLLDDNDKIAAFNIGKRAISSEAIEITPEIQKVIDEETKKSL

KKESIQISSSIVNQKSVEKTTKSDKKLKNTYQNNKPSKGGNKKHRR

>PIZ64637.1 hypothetical protein COY15_04620 [Candidatus Roizmanbacteria bacterium CG_4_10_14_0_2_um_filter_39_12]

MDRFKNLYEVKKTVRFELKPSRKKIFEGGDVIKLLKDFKKVQRLFLEIFVYKKKDSKLEF

KKKREIKYTWLRTHTKNEFYNWSRRSDIEKNYVLSKIDFLPEEILRWLDEWQKLTESLED

ITQTEEHKQKRKSNIAFALRKFSKRQNLPFIKDFFNAVIDIQGKQGNESDVRIKEFRGRL

KEIEKNFNACSKKYLPTQSNGVLLHKASFNYYTLNKTPKEYEDLKKEKELELNGVLSWAI

YKRERVIDRRTEQTEILFDCDIDWLKKIGLGDEIQNWVLDEAYQKMKIWKANQKSDFIEA

VARDKLTFQNFRKKFPLFDASDENFEICYKLTKELDKKPKDAKKVAQERGKFFNAPKEKI

QTKNYFELCELYKRIALKRGKIIAEIKGIENEEVQSQLLTHYAMVAEEENKKFIVFIPRK

NGEELENHKNAHKFLQGKEKKESGDVKVYHFKSLTLRALEKLCFKKTKNTFAPEIEKETN

PKIWFPKYKQQWNKTPEKLIEFYKKVLQSDYSKKYLDLVDFGNLNTFLKTDFTTLEEFES

VLEKTCYTKVPVYFSKKEFETFKDEFEAEVFEITTRSVSTGSARKENAHAKIWKDFWSRK

NETAGHTIRLNPEVSIFYRDEIKEMSNASRQNRTSDVNNRFSDPRFTLATTITTHADKKK

PNLAFKKIEDIKNHIDSFNTVFNRDFSGVWVYGIDRGLKELATLNVVKFSDVKNKFGVLQ

PKEFAKISIYKLSDEKAILKDVGGKSLKNAKGEDRKVIDNISEVLEEGKEPDPILFENRI

VSSIDLTQAKLIKGHIITNGDQKTYLKLKETSAKRRIFELFSNAQIDKNTTISGNKTIML

GKNNTIYWLCEWQRQNPWRDKKESLMQSLKDYLQNLDIKNHFKDIETIEQINHLRDSITA

NMVGLLFHLQNKLKIHGVIALENLDTVQKKSDSKMIDEHFEQSNEDISRRLEWALYRKFA

NTGEVPPQIKESIFLRGEFKVCQMGILKFVEVGGTSRGCPNCQSEWNQECGNQCNPKCNK

CTENDSYKKNKRKNVYICKDDNQCKFSTEESRNLLEQNLHNSDDVAAYNIAKRGLEINSI

>GHV42792.1 hypothetical protein FACS1894180_0100 [Bacteroidia bacterium]

MEIYKTTKAIRFKLSVNKDNDLRNHIPENQELNLSQLGKLGFEIINQVENYIWFEKKDKV

SEKPQTAMQIAFAKANIATNINQKIAKELSKANSIKTNFLTNFCKRDFYEWIGNEKPKKQ

YSLSKLKFLDEKLKNVIQNINQKLEIITNYADKRTRKEHHNLERKKEINLLVIELENKQH

FKFIKSFVENLILKNQSKDELDSLLKEFEKELKIATDYYRPSQSSGLIIAKASFNYYTLN

KNNKDLKIEENIKGELIKLEKQRTQENLPQEFTNIDFADLEKSKECTEKQQQEKFDKVST

LLANFIRYVNIGSKSNQPFSYDLSLKMKNDPNIKFHQEKENLSNYELMYMLMKLYKAKAK

ALFIAGISKKENLNMSFEDLQQNYPLFVVNNKEDFTTFKKWNEKVVKQTSDDKATTERGK

YFQKDGIYQQFVKKYTSISKKYGELKSKLAGIDKEKVESQLLQYWALILQEKEQHKLVLI

HREKAKSFKEKVEKLPPATNESPQLIYFESLTYRSLQKLCFGNSDTSTFAPEIKKEMNVP

IGEYEFQGNEQQKICFYKTVLQTDHARKVLNLDWSKIDDDIIKPHFEKLDDFKIALEKVT

YNKHVKIPEDINLFKDNNALFFDIVSQDLNLKQENKEYHKNKKHTEIWNNFWSDDKNNNY

NIRLNPEIAISYRQAKPSRIDKYQDGIVNGKQMRNRYLGEQFTLITTISENCLSKENNLS

FKTNKDLTDYIVDFNTQFNNKCQNVQYSLGIDVGTKDLAYATLIGKSDKHNLGYLPKTFT

AYELIKLDYAKPNINSKEYKAINNLSYFLNENLYNKSFADNKFQECFNEIFKEKQLSSFD

LTISKVIDGKLIVNGDYHALLNLKILHAKRQIYQSLIENIKQEFSDDFFRGIYAHRNEFE

NISFLKKDDIRIVATIENIKHELEDYFQEQKQIRQSDDNKEKEDKLLESYINKARNALVA

NMIGVIDFIRKQYQCYIVLEDLNQSVIESHRTEFNGDITRFLEFKLYQKMQNYGFVPPIK

GIAELREREGKKLKQVGIVRFVDEEETSLICPNCGKKAYSKSDDYVYNSDKGKGIFKCKK

YEILNFTKQTEEIIHTGKYYPKDIHKPCEYHNRDNQMQFSDLITNDAIAAFNIAKRGFEN

FLKQQ

>KKQ38176.1 hypothetical protein US54_C0016G0017 [Candidatus Roizmanbacteria bacterium GW2011_GWA2_37_7]

MEIQELKNLYEVKKTVRFELKPSKKKIFEGGDVIKLQKDFEKVQKFFLDIFVYKNEHTKL

EFKKKREIKYTWLRTNTKNEFYNWRGKSDTGKNYALNKIGFLAEEILRWLNEWQELTKSL

KDLTQREEHKQERKSDIAFVLRNFLKRQNLPFIKDFFNAVIDIQGKQGKESDDKIRKFRE

EIKEIEKNLNACSREYLPTQSNGVLLYKASFSYYTLNKTPKEYEDLKKEKESELSSVLLK

EIYRRKRFNRTTNQKDTLFECTSDWLVKIKLGKDIYEWTLDEAYQKMKIWKANQKSNFIE

AVAGDKLTHQNFRKQFPLFDASDEDFETFYRLTKALDKNPENAKKIAQKRGKFFNAPNET

VQTKNYHELCELYKRIAVKRGKIIAEIKGIENEEVQSQLLTHWAVIAEERDKKFIVLIPR

KNGGKLENHKNAHAFLQEKDRKEPNDIKVYHFKSLTLRSLEKLCFKEAKNTFAPEIKKET

NPKIWFPTYKQEWNSTPERLIKFYKQVLQSNYAQTYLDLVDFGNLNTFLETHFTTLEEFE

SDLEKTCYTKVPVYFAKKELETFADEFEAEVFEITTRSISTESKRKENAHAEIWRDFWSR

ENEEENHITRLNPEVSVLYRDEIKEKSNTSRKNRKSNANNRFSDPRFTLATTITLNADKK

KSNLAFKTVEDINIHIDNFNKKFSKNFSGEWVYGIDRGLKELATLNVVKFSDVKNVFGVS

QPKEFAKIPIYKLRDEKAILKDENGLSLKNAKGEARKVIDNISDVLEEGKEPDSTLFEKR

EVSSIDLTRAKLIKGHIISNGDQKTYLKLKETSAKRRIFELFSTAKIDKSSQFHVRKTIE

LSGTKIYWLCEWQRQDSWRTEKVSLRNTLKGYLQNLDLKNRFENIETIEKINHLRDAITA

NMVGILSHLQNKLEMQGVIALENLDTVREQSNKKMIDEHFEQSNEHVSRRLEWALYCKFA

NTGEVPPQIKESIFLRDEFKVCQIGILNFIDVKGTSSNCPNCDQESRKTGSHFICNFQNN

CIFSSKENRNLLEQNLHNSDDVAAFNIAKRGLEIVKV

>MBI9016532.1 hypothetical protein [Phycisphaerae bacterium]

MNKDIKQNSVSEYSYNKALRFKLEHLAGPLPAIPQGQANSKNLIETGNLLADEFEKFIYI

VNDDGTLNRRNGQKVLTKHISIKKQWLRQYFKNDFYAFGRQGKDSSLADYKYIQTPLCKW

FEDFREQLSSLQQVVSYADHQKSRRSEIANILTVINSKGYFVFVKDFCEYANHKANQSLQ

NNLQKLVNEFEQQLTAGINEYLPAQSAGLVIAQGSMNYYTVNKKPKNYQKEIDGLGIKLD

ELEHKYTEYKENKANQKSKLYELINTHASKERIKEECDLFGDNCDDNCLEKVTELTLQIE

KLNENKDSSNNLSLELKKEIKNLKQQRGHYFIKQTRGKTHDMYFKTYWKYCEEYKKIAID

RGKMLAKINGIKREQIESGNLKYWSLIFDNGSEKHLWLIPKDKMQEFHDSITADKQSLCI

LHTPHSLTMRALHKLCFAEESSFVENMPGELKSLQSGVKKATNDQNNRNFNADRSKDVLA

LEFLKKLMQSKYAHDILDLQHFDLSELQNTDDLNKFEAALEQACYYMEPIALSAESADNA

INTYNILIFKITSYDLEKRNQNTHQTAETKGKTHTREIWDAFWMGDANIRLNPEIKIRFR

QKNEELTEYMENKGFDLSKVKNRFLQEQYTAGFTFTLNAGKRYPRLAFAKTEDICQEINN

FNQKFNEQNWQGTYKYGIDRGNIELSTLCIAKFDQNDTYNYNGKSYLKPSFPQGQQDIKV

YELKKQWYDQKQKSDMENIPYENRKEKRIIANLSYFIDKIEDASWFDKKNCSCIDLTTAK

VIKDKIILNGDVLTFLKLKKEAAKRVLFELVADGKITDQNKELKWASDNTDSFKSIELRT

EQGNTIYFFEGKHGRNFEGLLVKDDIVYTRDNIKDNLQSYLDNLLEEKSKQSSNPYKHVP

PIAKINNLRKAIAANMVGVVCYLQKQYPGIIVLEDLEDDTIKKHFSQMNINISRPLERAL

IDKFQTVGKVPPHIKDIIEIRERFRRDGKAKSSQLGALVFVDKSETSAQCPYCGESWTWD

KSRKDKLKFLEHRYICGENEPCGFDTENIEGKTDFLKEINDPDKIAAYNVAKKIQDYKNI

EQLKILTGENKKQYKGTNGKSKPQTEYRQPKKYQGFSLEIPKKDKAQKNSQGTEIDKLIK

LLLNKKAKEVSNILKQLKAKDLSRSQIEQIEKILNKRHDAGQADYNPQVKKYFEIFNEIK

SGN

>MBP7967128.1 transposase [Candidatus Woesebacteria bacterium]

MRNKYEIRKTIRFKLDPKTKVIQPSVQTSLDMNVVQKNFIDAYKTVINSFEKLLFIQKFG

EDCLNRLITVKHNWLRIYTKHEFYEFKDQIIKYNRNGQKIINKINIADPHVDFLYTYFKE

WIINNRNAIENIQYLYARKDHELKRRSDLAYWITFISKRNNFESIFELFDGNIVHSESNV

DIDLIKEHLVNSRALLNQLMQYIAPNAVFGIEIERASLNYFTVNKRFCDYQTEILTKEKQ

LEQVFSVGELDALYRRDRFNELPSKFLYYVGFIGYIQQDNKSNNMSIAAWHEKLKLYKAE

QKKLFYERVAKGMTYEELKNSSDLPLLKDISPANFYSFCSGPKKRSSHFQYSFSNYKKYC

DIKKKLAMVYGKMKAEVKSLEKARIDAERTQSWALIHEKDGQKYLLTIQRDANGTMKKAK

KHLDGFLSDRNEYPNKLYLFESLTLRALDKLCFGLDKNTFLPSIQEELYKKCPSLFYGNT

LKRKDQLSKDNSELLLLYQTVLGLTATKNTLAVDSFIGFEALTKEKFNSLSDFEQKLKQA

CYIKKQITMNENDFDAFINSYQVRIYKITSYDLQKNDPDFLNSLSKKKILDRSHPEAHTK

IWLDFWTKENEQNNHNIRINPEIKISYRQAEKKDISLEQKSKPFNRAYHDQYTLSTTITL

NAHERSINTSYLKKREIEEFIQTYNTDFNKNIDQSKIYYYGLDRGEKELITLGVFSCKEF

DTSSASALQPQEITNLEVYTIPKDQYLTVKNGYTLYRSPAEFIDDHEYITPNGARSYLDL

TCAKLLKGKIVTNGDIATYLKLKEINAKRMIYELVRKNRLTHAQVLYDKVKNKIILDQGG

DSRYRYKDIYFCDDEYQQIAPPTSIEKDLQGFLTEILHGKGIIPTILEINRMRDAICANA

VGIISFLQTKYPGFLVLEGFNISKKEGFIQKTMEHLASRIEWKLFQKFQSTSQVPPVYRE

LMSIIGDNTVGLSEAEVYQLGIMLYVSEKNTSTCCPHCETISTKNKDAKFSSHQFVCVNC

GYIGNSDSIAAYNIAKRGLLAMKNKF

>WP_213348974.1 hypothetical protein [Candidatus Vampirococcus lugosii]

MTNIKNLQNLYEVNKTVRFELKPYFDEELIKPKYEGNLDENLKKFVEIYEKILSIFPEIV

FNFNKEKLNKKLKIKFSFLKNYTKREFYDKNIIFLKKGNNQIEIGESKKLDYLLDRFKSF

ENDNNFFLGKIKNLGFRNQEDKARNSDLNYFFYQISKITNFNFLKELFNNINIEEAEISK

KINDIKEKIKKTEKLIINIEKHLLPKNSGQVIEKASFNYFTVNKKPKNYDKNILEKEKEN

NKKLNEFTYFNKKERKIKNIFNIYNIKNSGFCKYITSDIGDLEIKRAYELMKEYKTEQKS

NFLKYLEAKAKGENIDLNISLYNDIFDKDQYKNILDLTKAILILSTAKSDILNIKKAKNK

EQKNEEIKYNLEKYNNEFGEKFGEIEKENFFEKITELKKERGQYFFEGNRKGKFKFYKFK

NFCNEYKKIATEYGKIKALIKSIEKEKIEAEKTNSWALILENKNNKYLLTIPRDTKKEIS

KYKTNLNQSKYFIDNLQNENNGEWILYKFESFTLRALDKLCFGKEVENIHGKLKEVDNSF

RKDLYNSFRKEVDDSFRKNLNKNICKYKYKIFFNKGKLKDKFEFKKQNGENDEKLLIDFY

KTVLNLESTKKQISIKYFDGFDDFIKKDFENLEKFEEELKKVCYKREKIIISDKTKQEII

ENFEAKLYKITNYDLKKYDEEEIKKLETKKEFKREKPNNYTKVWWNFWTEDNEKEKFPIR

INPEIKISFIEKDKNFEKKYSGLKFNRKLYDRYILTTSITQNSFNKSIDLNFKETDEILN

FYNKYNKDFNKENSFKYFYGLDRGENELVSLGIFDFSKENNKGVKIKVYELNEKGLKATN

EKGTPIYKNLSYYTFDTHPDFYDEKEVSCIDLTKAKFINGRIYLNGDLSTYLNLKIASAK

RKIYDLVYAKKIKDYKILDDDYNITINCIEKKEFLYYYKENVICSKEEIIEELQEYLNKN

KFDENENNISILKINNLRNAICANMVGIIIYLQELYPGMIFFEDFELSKKQNDFEKNNTT

LGSRIEEKLLQKFSSLGKVPPNYKQVFSFQSDKKLDQLGIIGYINKYNTSSACPVCNEKL

FGHGKGPKFENSMHHYKENNGIDYGNNLKKAKKTENNTCNYHMKNKRHGFDFINSGDDLA

TYNIAKKGMEYIKKQINS

>RAL57849.1 hypothetical protein BLD25_00640 [Candidatus Gracilibacteria bacterium GN02872]

MLESFKDLYEVRKNASFRLEPQENAATNENLYNQDVDLSEILSKYKNFVYDLKNLLYIEG

EKVTEKEKDNDLSVDKLDSSNVVHINHKYLSKYFSKEFFENYDRIIKRNKHGAKTKNWTT

IGNNDFLEVFFRSFFNNAFENINKLQELIDTKENEQKRKADIAFEIRRLLSRNHLGKIFP

LFKGGYLTHKNDIKTIPELTNQAKELKELLDNAKIYFLEDSNFGVQVEHISLNYYTRNKT

QKEYDELIKGKKKLLDSPYNKDISELYGIDRNLPIKQLYEEMKMFKARQKSAFFQQLQSK

TPFKKFKEPFTFTDNEGIKHKEFRTELFFDVEEEYYNKMLDLTKKIEKSGSNKFKKERGD

FLLFQDKGTKSFKGWSGFCSEYKKVAMEYGKRKAESLSLEREKILAEQERGWGVFSKVNN

EFYINTFPIDKVKDAYQSLSKKKNDGELSYFILSSITLRALNKLCFSKDSKFINKEILIK

LDNEFLKEKEYNNKTKEIVLRKKSDLTELKLIKLYYNVLELQNSLRISYKNSDSFDKLKS

SKNLEEFELNLKNETYKLEEFKTTEDEFNKILKSNEGKSYKIINKEKGNFEKWWKNFWNS

SKDGEIRINPELNISLRKGNEDYNDKISFHRKKDDVYLLNVNFSHFANRSYIDSSFIEDD

KRKENLKKFNDLYNKNIKLNYFYGLDKGTNELITLGLFKKEGGKIKSVNISKEIPVYRIT

RKGLLHSKVTFKKAPKEGQNPKTIHTLYKNPSLFIDELDNEEIFEKVNIESCLGNLNSAK

LIKGNIILNGDILTNLNLHKLSAKRKLYIAITEGKTIIKNIGFDEKEQENGAFFYEYINK

GKTEKEPVFYINEEFLTIVSLDELKNELNEYIKYIKNEPNEYKKYMGNNALDTDISMLAI

NKIKNALCANIIGIIMKLQEKFPGYVCFESLSNDNNNRDMEKEHSFLGNLINDKLYSKLN

INASVPPILKKFRSELKNSYYQYGQVIYVNEDSTSSACPVCNDKFIKGINKKGNLEEYKK

NKLYGHLTESESSMHHLTDDEYNKKYDNNDKFREKHTNDKGKKKDNIYRSGNGCNYHMKD

NPKGFDFIKSGDDLATYNIAKKALEYLEYLESQNNDETK

>MBB1565335.1 hypothetical protein [Candidatus Gracilibacteria bacterium]

MNKFTKPYEVRKSIKFKLIPEIQKRPHYTPDYDLEFSELSDSYKELIENLINFFYSEKEI

VDKEDKTSKFSNLKGFKEAFANFDFDEAEEKVFDLKNKEFSKMPRLKHKFFKENFKEDWY

AIKDRLIKKNKTKNYEILDSLPFFQKVLKDFFEKSEQLYKELDEIISSPDNASKRKSDLR

FVLQKISGNEVFKKIIKLFSLGFVDHKNDVEIVGKISEELGEFSEKIQNAIELLRVNDSF

GFPVEYLSLNYYTINKTPKEYDKEITEKQNNLEKLYEGGLSELYGIDKNQTIEQLRNDMK

MFKARQKSAFLEFLQMKNEFSELEELKTYLNDIEKKYKQKKGSKYNFYEELQNNTFELDS

SDVDTDIDKVFKAFSLILFLETEKDDYEKLLNLTKKIEKSSTLSERTSYKIERGEFFKGK

NENGIFSEDRDTSYDGFCSEFKKVAISYGRLKAEKLSLEREREQARFERGLAILSKDLDG

NYYINSFSNNDSKKAFEKLKNINSTSGDYSYFIMKSITLRALQKLCGKENFKKTVVNKIS

EKFTKVNENDTGRQVRKFKSFEEIGDNIVEFYVDILDKQRTLDVSYRSGYEEGLKKLKGA

KDLDELEILLKQETYTLEEHKISKDDFEGILEDYKGNSYKITSEDLEKNIETNNFTKWWN

IFWTIENKEHKYISRLNPELVISLRAGEKVFKDRKKHRKSDKSFLLTMSYSHYTDKSYID

DAFIKDEERKSSLEQFNKHYNENHKFDYIYGLDKGTNELVTLGIFKKVGDKLEKVNISEK

IPVYRITEKGLKYSKKYSTRVDGLGGEREIFLYKNPSLFIDEIDNEELFEKVNIDSCIGN

LNCAKLIKGNIILNGDINGFLSLNKIAAKRHLYDAITKGFMLKDKITFDEVKNNFYYEHT

LAGKKFNKVVFYLDSFESITTKDEIEKELNNYIKLVIEHKSEEDVSIDGINKKRVAISGN

IVGIILKLQEKFPGYIVWEALNLDQNNKNISSNHAFLGNLINDKIYQKLMISLDVPPILK

KYRSELTKDTLTQHGKVFYINENLTSKACPSCNDTLTIEVNGTLKEKDPNKHKVKVYQLF

GHLTEYENEMHHLTDEEYDNLYDNDKNFKKFHSNDKGNKKQNSYLAGENCDYNMKHNPKG

FDFIRSGDDLATYNIAKKAKEYLEFLEKQKNNNS

>MBP5368392.1 hypothetical protein [Bacteroidales bacterium]

MDNTKYQITKTVRFKLEPVFDETNTKVEFDKEFNPIVYSQIDLDSFSNDLKELLQKLKKF

LLEYEGDSPKSIDKPNNKFVYVFKKNIEIRYDFLMAFIKKDFFAARIKEEKLKFYSVNKY

GFLAQRITEVLLDIEDVNRKLSDFANLDLEEQEQRAVIAYWINRLDNRFHFHFIHELFKS

LNDKLFADTSYKERLSELDSFKDEIKVMQKSYLPAQSNGLQVATSSLNYYTVEKTPKPLD

SLDGSDEEGYSLINEKMKAQNANQSKIFEPLYRGFVLANDYISYVNYSKWDSIDQKISFK

KGNFQIKGDVENVNIYGQCESLIQNAINKKYPFINVYSYFKNLQKSESDGQNELFINLYA

SQNNKDIKEVAAFFIDFESDNILDDKYRKLFPDDFGKRLHNLYTTKQSYRIDELISHFSL

EETYLFLKCWRKSIVKSYFTQNVYNYLSDKDEYKDKEAIKIALKAFDLYKLSDDDLTQYI

ELTKQIKVISNQINDAIKNFENVDSLQKQLQNICDERNTILTNQPLYGNFCDFYMKVAIA

YGKNKALIKSIEKENNTALQLNYWSFISEIDNTRYVYFIPRNSSNNYKAAKKILFDDKHK

VSGKPNLFYFESLTLRALRKLCFNNINDNTFRDSLTSCKLPACEQIKKIPGQKPIPTTDF

DKILMYVRVLNDGGSIKNQLNTYSKLKKKVLNIRYNRDTASFADFETKLNLYCYKVLGYC

CDVDALLKNNNIPFFKFKIELPRKNNKEKDYTKLWNEFFKSGNETNDYPIRLNPEVKLFR

RDANVDFGSWYDCGKSGAERKETRFEKPRYTLALTFTENADSTKVDYSFSSDTRDYGKVD

RKVINIQHKIDDFTNEYKKDKNCKFALGIDVGDNECLACLTACDENGNPILIDVMEVNDD

TKQGKNDIAKKNGNVAEFMHNPSYYLNKDLYDNSFDTPYSRDKYVVHKKVCSIDLTTTKV

FALDNIGKNKGIVLDSDYISHLNLKKINAQVKMSECRKENPECEFVLGTFRGNTRIFALS

KDVAQRYSATSSEILLDKQPSIYDFNCKYDNVKNINEIYNEIKILERQLGKENLEENINK

AKNTIVANMVGVIYYVYNQLQKRFGADSYGMIVFEGLSEKEIAAHRDSNKADITLTLRNA

LLKKFQNDLCVPPYKEVSKLSDGLKFTIKKSKNNPKTEKDFGIVRFVNQDNTSQCCPRCK

AKSSVIHPEIFDCCSITLNNWKATPNMSDEEILLYESMDINDKVAAFNIAKRAFGLF

>KIM12007.1 MAG: hypothetical protein KU37_03970 [Sulfuricurvum sp. PC0866]

MLHAFTNQYQLSKTLRFGATLKEDEKKCKSHEELKGFVDISYENMKSSATIAESLNENEL

VKKCERCYSEIVKFHNAWEKIYYRTDQIAVYKDFYRQLSRKARFDAGKQNSQLITLASLC

GMYQGAKLSRYITNYWKDNITRQKSFLKDFSQQLHQYTRALEKSDKAHTKPNLINFNKTF

MVLANLVNEIVIPLSNGAISFPNISKLEDGEESHLIEFALNDYSQLSELIGELKDAIATN

GGYTPFAKVTLNHYTAEQKPHVFKNDIDAKIRELKLIGLVETLKGKSSEQIEEYFSNLDK

FSTYNDRNQSVIVRTQCFKYKPIPFLVKHQLAKYISEPNGWDEDAVAKVLDAVGAIRSPA

HDYANNQEGFDLNHYPIKVAFDYAWEQLANSLYTTVTFPQEMCEKYLNSIYGCEVSKEPV

FKFYADLLYIRKNLAVLEHKNNLPSNQEEFICKINNTFENIVLPYKISQFETYKKDILAW

INDGHDHKKYTDAKQQLGFIRGGLKGRIKAEEVSQKDKYGKIKSYYENPYTKLTNEFKQI

SSTYGKTFAELRDKFKEKNEITKITHFGIIIEDKNRDRYLLASELKHEQINHVSTILNKL

DKSSEFITYQVKSLTSKTLIKLIKNHTTKKGAISPYADFHTSKTGFNKNEIEKNWDNYKR

EQVLVEYVKDCLTDSTMAKNQNWAEFGWNFEKCNSYEDIEHEIDQKSYLLQSDTISKQSI

ASLVEGGCLLLPIINQDITSKERKDKNQFSKDWNHIFEGSKEFRLHPEFAVSYRTPIEGY

PVQKRYGRLQFVCAFNAHIVPQNGEFINLKKQIENFNDEDVQKRNVTEFNKKVNHALSDK

EYVVIGIDRGLKQLATLCVLDKRGKILGDFEIYKKEFVRAEKRSESHWEHTQAETRHILD

LSNLRVETTIEGKKVLVDQSLTLVKKNRDTPDEEATEENKQKIKLKQLSYIRKLQHKMQT

NEQDVLDLINNEPSDEEFKKRIEGLISSFGEGQKYADLPINTMREMISDLQGVIARGNNQ

TEKNKIIELDAADNLKQGIVANMIGIVNYIFAKYSYKAYISLEDLSRAYGGAKSGYDGRY

LPSTSQDEDVDFKEQQNQMLAGLGTYQFFEMQLLKKLQKIQSDNTVLRFVPAFRSADNYR

NILRLEETKYKSKPFGVVHFIDPKFTSKKCPVCSKTNVYRDKDDILVCKECGFRSDSQLK

ERENNIHYIHNGDDNGAYHIALKSVENLIQMK

>MCK9562686.1 hypothetical protein [Bacteroidales bacterium]

MENIKNNYQLSKTLRFGLTQKQNGNSSNTDNVYHSHSALKELVDISENRIKKNVSTEGAT

EMQLSIESIRKCMIMIEQFIKDWKRVYYRSDQISLDKDFYKKLSKKIGFEAFWFERNKKT

QQRIKKPQSCIIALSELSKRDNFGKERQEYIVEYWENNLLKSTERYEEVSEKLEQFELAL

KINRTDNRPNEVELRKMFLSLVNMVREVVEPLCLGQISFPKLEKLADNSKNKQLRKFATD

YQSKSDLLTQISELKKYFEENGGNVPYCRATLNPLTAVKNPKSTDSSILDEIKKLKLDVI

LRDYQSVALFDNSIRDLTASQKMQLLNQNNEGLIKRGLLFKYKPIPAIVQYEIAKVLSAE

LNKDEQELRNFLRDIGQVKSPAKDYAELQDKKDFDINHYPLKVAFDFAWESLAKSVYHPD

IDFPEEQCKTLLREVFQVDENNENFKFYAQLLELRSLLATLEHGKPTEVITIENEVKKIL

ENIDWSKFGDRGKNYKSAIENWIHNRNKKDFKGDYFKKAKQQIGLTRGRQKNLIKKYDEI

TKSYKDIAMKMGKTFAEMRDKITGAAELNKVSHYAMIVEDTNQDKYVLLQEFVENNKDRI

YAKSTPQNEDFKAYSVNSVTSSAIAKMIRKIRIDKLQANERNNNRQQAPELSETQKEARN

IKEWKDFIAEKRWNYEFDLKLDNKNFEQIKKEIDSKCFKLETKYMSEEVLVDLVKNQNCL

LLPIINQDLAKKIKSESNQFTKDWNAIFAQNTPWRLTPEFRVSYRKPTPNYPKSERGDKR

YSRFQMIGHFLCDFIPKTADYISNREMIANFKDDEKQKQTIINFHERLNPKSENEKMNML

LAKFGNKNSNQSKETKKEEKFYVFGIDRGQKELATLCVIDQDKKIIGDFNIYTRSFNTQN

KQWEHQLLDKRHILDLSNLRVETTIVIDGKPDVRKVLVDLSEVKVKDKNGNYTKTDKMQV

KMQQLAYIRKLQFQMQTNPDTVLEWYNKNQTKEAILNNFVDKPNGEKGLVSFYGSAVEEL

KDTLPIERIEKMLQQFKSLKNEEKEGKDVKAEIDKLIQLEPVDNLKAGVVANMVGVIAHL

LKEFNYQVYISLEDLSNPFGSHVIDGTTGTHSKTNKGEGKRADVEKYAGLGLYNFFEMQL

LKKLFRIQQDSQNILHLVPAFRAVKNYENIIAGKDKIKNQFGIVFFVDANSTSKMCPVCN

STNETNREYPNAKKGTSKDDKEVWVERDKSNGDDIIRCFVCGFDTTKKYEENPLKFIKSG

DDNAAYIISAFGIKAYELAKSVIDNK

>MBC6415218.1 hypothetical protein [Bdellovibrionales bacterium]

MGDNKKPTNYQYTRGVRFKAEPVDNTKLLFEKEETKNVNLTALEEDLSKFHKDLTDLLYN

KEKKIKNKSDKKFKTTITINKLWLKCWHKDIFYGQIKKNSNKKGKYNLKDLNSLPFKEKL

EQWDKACKEIKKFSSAPQDSQYRRSDFAEWIKLLLNNWKYFNDFLRELHAKSPEDDKKIT

QLKEDSDKIHKNLKISEKSYLSFQSSGVEIAKASLNYYTVNKKPKEYDKELKEAEKDLKE

DSFSSIVEEKYRYQLKSKSAIFTFKSKQEKEWIERYCKEEFNKNLKENDIGLSLDKTYSM

MKAFKAEQKSIFYELASRVPLKANDNLRNKSSTSLFTRFQEKVSHTNSDKKEDYINNNKK

HFLFNYKFLCESFSSIDKLSESFSLFKDIKGNKKFHYEEFVKLCKKIKNPKNKKEGSSNT

KKLAEKRGKYLKKTQAYFKEYTDFCDDYEKIAQKRGKLIAQIKGIEKEKRESSEINYWAL

IYTKSGKRQLWLIPKKPSQTDKSEQNNSSDTNLQSAKKCIDKKPQASAEANSYLSSFESL

TMRALHKLCFAEQSSFVKEMPDELKKQQKEVKTFRTQGDKEKLKQKEKKEINFLKDLLKD

NYTKSKLKLNNFNLTKVHKAENKQDFELALEKDCYYEKKVSFNDKEKEEFIKKFHVSTFK

ISSYDLQGRNKNTHQSPESENRIHTDWWNEFWKKIKNNQEESIVKEFKIGAIRLNPEIKI

RHRKVDKNLEAYFYKRKFPKEFKHRNLQEQFTAHFTLSLNAGKKYEDLAFSKPEEILEKI

ENFNQRLNKSRNFETAWKYGIDRGNIELATLCLSRFNREDFYEVKGIKILKPTFTKTEKD

TPCYSLKNYDLKETYQTKTQGEKTRFAVQNVSYFLEEKYLNDTNFKPENLTCLDLTTAKV

IKGKIITNGDIMTYLKLKKAVAKRSLFELYTKREINDFSKLQWIDYEHGDKNRKMSDGVL

NIKTKDGEKTIYWYCRKYENILINSNKNIKYSKKSILCSLSSYLGNLKESNNQHTPSILK

INHLRDALTANMVGVICFLQKKYPGFIILEDLNRGIIDRHFFQHNENISRRLENALYNKF

QTLGLVPPHVKDIIQLRENNRKKANKEKEKIEEQIGAIVFVSEENTSKTCPYCEEISKKQ

NKELKNDLKFRQHRFICDSCGFDTYYFYENPVKEPTPEIDETKKKKEFEIVRDIDDPDKV

ASFNVAKKSMKESS

>MDE0119592.1 hypothetical protein [Bdellovibrionales bacterium]

MNQNNQAITFKGNPYQYIRGVRFQARIDGRVSESFKTKHKAIGNEEVSKSNLSELAGLLS

GFHTELKSLLYDTGSETEKEEFRKNLSVSKTWLKKWHKELFYSQIKKDNNRQGKYALKDL

KGILHFFEQKLVSWGEHSEKIKKSAEASKNERFRHSDIATAIRALSSRELLDYIGDFLME

VHTKSTELDNRVKKLKEDLENLREQLKTAEQEYLSFESPGIEIAKASFNYYTLNKRPKEY

YEKEPQKAKDELCKNHFSKIQKTKNREYSWEFVNPRSEIFQFKSDQEKKWIEKYCEKHLK

DDLKTEINLTLDQTYSAMKDFKAEQKSIFYEVITHIASNQNNSYEDKNSNHLLKGWQFPY

SQQNVQGVNTAFSLFQFKEKEIKKNQWTIKNILSLYKNESNKVTADKAFEVFIHLTKQIQ

QSGDKKNKKTQTKKLLQHPKHRFLARRQNRLKNDSSLGFYNNMDTIPYKSQNPKKDRGAF

LFGKGCYFKEYEKFCEEYKKIAQKRGRLNAQIKGIEKEKREALQTGFWAFIYTTQTKKQL

WLVPKDKCEAAKSFIYSPDRKNGSSEDSQCLSCFESLTMRALHKLCFAEQSSFVKDMPYD

LKTLQQSAKEFKTKDENKDLQEQKMKEKSQKELVFFKKLLKNEYAKEKLLLKNFNLQECY

KAENLTEFEKSLEKACYHEKKISLTEGEKQDFLKKQDVTVLNISSYDLEGRNKNTYQTPV

SANRYHTDLWQEFWKGVGVPSNRDRTVKGFKIGEIRLNPEVKIRYRKADEGLKKYFDKKK

FPSKFKHRRLQEGFTVHFTLALNAGKRYEGLAFARSEELLSKINDFNQKLNQEINFKTAW

KYGIDRGQKELATLCLVRFNPDKDFYQVDNKNILKPTFPKDDKDSETIKCWTLKNCNYKE

EYTTKKGEKRERWAIKNLSYFVDERYLNNRELFEEENIACLDLTTAKVIKGKIITNGDVL

TYLKLKKTVAKRQLYELFQNGKINGNASLEPSTQENGSKEEERHRPDGVLNIKVSNGEEK

TIYHYRKGFEDILINKEKNIKYNQQSIKDSLNHYLNELRENNNSHTPTILKINHLRDAII

ANMVGVICHLQKTYPGFVILEDLSKSNIDKHRFKNEENISRRLENALYNKFQSIGLVPPH

VKNIIQLREQQQTKRKQEKTSKESMEKEIKNMEKKIEDVKKGINSARDAEKKEELEKKLK

DTKEKISYLEKKKNRLESNSLMKENQNQFSQIGAIVFVPKEDTSKTCPCCPEKTPGNSDL

KFRQHRFLCNNKACGFDTYYFKKEEEKVRDPNPSVKEEGYKEQFNLFKDINDNDKVAAYN

VANYNNKVIKKMK

>WGK68195.1 hypothetical protein P0082_06830 [Candidatus Haliotispira prima]

MSDSIGAGNGQYSYTRAIRLKGILQSGSLPKPNDTLEVSLEELAEHLEEISLLFEKIVFR

RNEKNEQESRKGIRINKSWLKRYHKALFYLLIKNNKNRDGKYDLKTELRHVQEDIRERWL

TDFEHYQKNLTSAARRPKASKLRRSEIAESVRGLLHPSILHYALDLFSYNGITTKDTEID

GQITKLRGRLETVREGLLSVQRKYLPAEATGIQVAKASFNFYTVNKKPKEYTVESLEKEL

YEKPYLTISIKTTKKGNKQVEFVGLSKPLSDSQIEQEIVWFEKYLEANIKPVSENKWELS

LEETYKLLKAFKSQQKDIFYKFMGHLAQNKKESFMVKNPDKPLDNYCFLPQKEGQNNIDF

VNSLFSLFKFRNLKKESEEKNNRFGHPSIAEENYKKFISYYKDIKEDKKERGKYLFGKNC

YFLGYSKLIQKYEEIARNRGKLVAQIKGVTREKQEAEEIGYWSFIYRTSEQSKLWLVPKT

EDLSALKKFLDEQVDAGAEDKEYLCYFRSLTMRALHKLCFAEGSSFAKGLSSELQVLLSE

LKELLKAAKEYPTTDERSRKEKKQKQLVFFKKLVQSQEKLGTLELEGFDLEKVKNSQTLQ

DFELALEKVCYNVRKIRLNETKQQRIISDYKIVTFDITSYDLEDRKRTGEDRRHTAIWRT

FWGYPGDGAVSKVQGFDLGEVRLNPEVKIRYRQADDEWKKSQTEERGNTVFKPHRKLQEQ

YTLGFSFALHAGRIHEELAFAKPEDLLAKINQFNADINEKMDFDHAWRYGIDRGYIELAT

LCIAKFQPEKKYKYAGKDIPIPEFPGTALKAYSFKGEKYDYTELYVTQVGGKQRRAILNL

SYFLEKDEAPRADLFEEESITCLDLTTAKVIKGHIITNGDVMSYLKLKKVAAKRQIYTLY

HQARSIGKDDTLTWGKENELSLPGLEKVIYYFLPKYEAILSKDEIKDSLNSYLDELRRLE

KARQGQSSEKENPHTPAVQEINHLRDAITANMVGVICHLQKTYSGYVMLEDLDDVKKKHF

EEHQENISSRLELALYNRFQTLGFVPPHVKDILQLRKENKPESCQIGTILFVDKTKTSKD

CPYCEQTDKKGEKDADRAKLVEHRFSCTPKENSCGFDTYRFKGPQDRVEGSVEVEDTDRD

FSFLKEIDDPDKVAAFNVSKKIKTVKDIPPWDMSNKPEEKQDQEEIVNKKKKHPHKNIKK

ESPETFTIGDSMQEKRHKRR

>HOI18830.1 hypothetical protein [Candidatus Woesearchaeota archaeon]

MLIQFKNHYSYNKSIRFKLEHKNGKLPKLESDNVDLNKLVDIGNSLKDIFEELVYTKNNY

NKLNSLVSIKKQWLKIYFKNEFYSNGKIQNYSLSNFSYLPNKLIEWLNNWQNNLKALIEL

TKQQDFNKTKKSEIAYILSLFNGKYSFSFVKDFSTCINHKNSQEQILKLQGVVENFEKVL

NLCIQEYLPSKSAGVVIAQGSMNYYAINKEPKRYDNILADLNQKFEELDKEYIAMKQYKS

SQKSRLFEFIRKGFSKDQILSEFKKKENNEVSFVYNNQIIIRIYTQELFKDSYCLGEVIK

LTKKIEELNESKDSNNNLPEETKKEITKLKKEIGFYFIRRTRGKSHNNYFKSYYGFCNDK

FKKKAQERGRLLTKIKAIRKEKIESQNLRYWSLILDDGKDKFLWLVPKENMQEFRRELSK

IHPSGESSLFLFHSLTMRALHKLCFAQESDFVKEMPKVLKEEQLNCEKASNDTETNKRIK

RNFGLNYIKTKDELTLSFLKKLIISEYAHERLDLNHFDLSKLQVATTLNEFEEYLEDACY

YLEKISISSSMIKELLEEYNILNFRITSYDLEKRNKNTYQTPESDIKRHTKEIWNKFWEG

DRFIRLNPEIKIRYRQKNQNIEDYLKEKGFDLTKIKNRFLQEQYSVSFTFALNAGKKYPK

LAFVKTEEILEKIEEFNDEFNKQYFDNSYKYGIDRGNIELATLCITKFNKNDTYEYKGKK

YLKPNFPTSQEDIKTYELKNEWYKRTAISNIETKPKNKKTPKRIIANISYFIDNVENEEW

FNKKTCTSIDLTTAKVIKGKLILNGDVLTFLKLKKEAAKRILFELVAQNKLTAKNKELKW

KSDDGNNSDSVRLICDVLDNETNSIYFYEDSKYGRGFEGLLTTDKTAYSKEGIRINLQNY

LNHLISEKENKSNKAYSHVPSIEKINHLRDALVANMVGVISYLQAYYPGIVVLEDLNHKL

LIKHFEDLNINISNRFEHALIEKFQTLGMVPPHIKDYLEIRSSFRMSRNDSSQFGALIFV

SKEGTSKECPYCEKKWNWGKEKEIELKFSKKQYICGKENSCGFDTKHIQNTFEFLSEIND

PDKIAAYNIAKRGFKSFINKSSIKKQ

>MDD3976412.1 hypothetical protein [Candidatus ainarchaeum sp.]

MSELKFLGDYSYNKAIRFKLEHKCGELPKFESNNSNPLDLIDLGLKIVDDFKDLIYLKTE

KNELKLDKNKSPILTKFVSIKKQWLRYYTKLEFYDGSKNKSVDFSITNFPYLSDSFKKWI

VDWTDAINHLNNLLQDKENLMIRRSEIAYYLLLISSKNYFNYIKDFSFYSNDKNTTKVAE

FQKLILIFEEKLNSCVIKYAPAQNAGLLIASGSMNYYTINKKPKDYNFELSQLKKEYEIK

QEEYNLFKVYKSTQKSLLFQYINSGLDKDEILKKLADNYNFYKENNKKIKTFYSGDSGND

LAYKYSRLFADDRALTSIINLTDKINELNLKLQTTNNNKDQIKLEISALKQKRGFYFIKR

HPRKSGDPDFYISNEKNWHDRYTFCFSNYTNFCKIFKVIAQKTGKLKAKIRAIEKEKIES

QTLKYWSFVVDRSDEKELWMVPKENRTKFKDLLKSDNSTDVNIYQLTSLTMRALNKLCFA

EESSFVKDMPDYLKDLQKKVKEATDDEQKIKNQRKKINFSTDIKTKSELKLSFFKELIKS

EYAKSLLDLSKFNLEDLQKVKDLKEFEMLLEKNCYVLEKIGLSKESCEKLIKECNILIFK

ITSYDLEDRPKNTYQTKESDYKRITSEIWNTFWENPGINLRINPEVKIRYREKNEEHIKY

LEEKGFDLTKIKNRFIVDQYTCYFTFTLNPGKLYPELVFAKTEELIKKIQEFNDQFNKIN

LEGTYKYGIDRGSIELSTLCLAKFDKNDVYTYTVDDKTKTILKPKFPNGEEDIKVYELKK

ELYNKKAISDLENLPMEKRKPKRIIANISYFIDKINDPNWFNKKTVTCIDLTTAKVIKDK

IIVNGDLLTFLKMKKEAAKRVLFELVSEGKITPDHKELKWGSENLEDPSSRELRYYPYTS

DKIVYHFENNFGRDLEGLLIKDDLIYSKENVLSNFQVYLTELLDNKLVDKNEPNTHTPTI

PKINNLRRALVSNMVGVITYLQKYYPGYVVLENLKSSEELNKFERNYINTSIPFEKALIQ

KFQTMGLVPPHIKNLWEIREEKTKKNLLVNKQNFQLGAILFVNKDRTSKDCPYCEKPYTW

DKEKADELKFKLHQYKCGPDPSCGFDTKELLPKFDFLKEISDPDKIAAFNVAKRWKDIVE

NDAKK

>MDD3794022.1 hypothetical protein [Candidatus Gracilibacteria bacterium]

MDKENSFKGFTNLYEVRKTVRFGLTQPNKKGELKTHLEFDDLINKSFENIKKDVKSRDKP

NFKEKELIEKINQFINGLEKQLGNWKQIYERYDVISVNKDYYKILARKAKFDAFKKDKKP

QASQIKLSSLQKDNRKDNIIRYWGNIITRSDYLINIFKPKLEQYLNAVNNPNNSSHTKPD

LIDFRKVFLQFLKVNEEYLQPLFDKSIQFETGKKENSEEIKKINTFSGDENNKEINYLID

LGKEIREYFEANGSQVPYGKVSLNYYTALQKPNNFGEDIRKGVENLGIIKFLNKSEEDIK

NYLKQNSKEKINLLNNAKNHYFIELIHLFKPKTIPFSVKYNLAKYLEKNFNLKYEDILNK

FDLLGKSVDIGKDYLECKEKEKFSLEKYPIKSAFDYSWENLARNLKRDVDFPKSVCEKFL

KDNFDIIINNSSFNLYANLLFIAENLATIEYGNPNNENEIIESIKNTFDDIKFESNKQEY

DGYKKEILNILNQEKSKRNYKNILTAKQRLGLLRGQQKNKISKYYNLTQSFKKIASFIGK

TLATIREGLKEENELNKITDYGIIIEDKNQDKYILTLKLDGKDIREKIKSKLWDGEYKVF

EINSFTSRALNKFIKNPLGEDSKKFHGDYKYKHKEVSIYKDVKWIGYKEEFLIHLKDSLV

NSQIAKEQNWKAFGWNFDNFNTYEKIEKEIDKKGYKLIKNSISKENLEYLINEEKCLLFP

LINQDISSKKEQNKNEFTKDFNKAFLGIGYRIHPEFSIFYRQPDEENKKINKSGIINRFG

RLQLLANIGIEYIPQNNDYKTRKEQNKISLDQTNQNELVQNFNKEKVNKYFDSLDDYYIF

GIDRGIKQLATLCITNKNGIIQSYEIYTKYFNNNSKKWEYKKNRIEGILDLTNLKIESDK

DGNKFLVDLSLFEAKDENGNSTGTNKQNIKLKQLAYIRKLQYQMSSNEKGVLNFLKKYQT

KEERQNNIKELITPYKEGHHFEDLPVNIFEEMFENYEKLKNDKTLSEIEKQNLMKLTIEL

DSSEDLKKGVIANMIGVIVYLMKKYDYKVKIAVENLNQSFMGQNDGLNNSYISIKTNFKD

QENGALAGMGTYHFFENQLLRKLYKVSVEEGILHLVPFFNSLDNVNKLNFEKEKILWVQT

ENYRKFGIVSFVRPHNTSKRCPICKSINVKRKDNITTCSDCGFITGKDNNIVIKKYKKEG

LNLDLIKNGDDNGAYNICCKIGL

>MBB1543285.1 hypothetical protein [candidate division SR1 bacterium]

MVNLESFKNLYEVRKTVRFGLNQPNKKSNINKTHGQLKDLVDLSFEREKKLINNEKNQVL

IDSEKALIEKLQQYVNGLEVQLENWEGTYQRYDLIAINKDYYKILARKAKFDAMWEISKF

DKKSNKYIKVKQPQASQISLSSLKIGNRSDSIIQYWGEIIEKTDYLLNIFKPKLEQYERA

INDANSTHIKPDSIDFRKIFLQLLKLTKEFLQPLLDRSIIFEFSKKKVSQEIEKISEFAG

EKNNTKIYNVLKNGEELRQYFEANGSQVPYGRVSLNYYTAVQKPNNFDQEIKKAIDDLGI

INFLKKRDSQIIDYLHQGSKQKIKLLLTSKSPYSIELLQLFKVKPIPFSVKYNLAKFIEK

NYKNEINLSYEEILDKLNLLGRAIDIANDFKNSNNQNNFSLDEYPVKLAFDYAWENTARS

LKRTIPFPKEVCKQFLKDNFDVDVNVDNADFKLYANLLFIADNLATIEYNNPNNEAELIN

EIKQVFDSIDFSFDKERYGGYKNDVLVLLNKAKPQRDYSTILKAKQELGLLRGGLKNKIK

KYRDLTQRLIDKKDSHFGIASFVGKTLAKIRDRLKEENELNKISHYGVILEDKNQDKYLL

ISQLDGKDTREKISQKFGNGDIKVYQVNSFTSKALNKFIKNPLSEDAKKFHGDFRYKHKE

VSIYDEKGKWTGYQESFLSHIKKCLIDSEISREQNWEAFGWNFAGCNTYEEIEKEVDSKG

YQLTENFISIGNLESLEKDEGCLLFPIINQDISSQKQENKNIFTLDLEKVFEGKECRIHP

EFSIFYRKPMEEHKKENKSGIINRFGRLQLLANLGIEFIPRNSSFKTKKEQNRIAIDQKK

QNQLVQEFNQEKVNTYFEGLDNYYIFGIDRGIKQLATLCITNKNGVIQDFDIYTKHFNSE

SKKWEYKFHRKDGILDLTNLKIESDKSGNKYIVDISLFQAKDEDGNPTGTNKQNIQLKKL

AYIRKLQYQMSANEEGVLSFLEKYKNKEEREQNMKELITPYKEGKNFVDLPMDIFQEMFE

NYYRLKTDQNLSESEKKNLMKITTELDASESLKKGVVANMIGVIYYLMKKYEYKVKISLE

DLSNAWFFSKDGLSGDVVLNTKNDGTMDLKKQDNLALAGVGTYHFFEMQLLKKLFKISTE

EGILHLVPSFGSVKNYTEIFKDKGKYVYKQFGIVYFVDPRNTSKKCPVCLNTQTTGKKEG

IPIINRNYKKSNIFYCERCGFQSIHSHCQEENIKDSNGFHYSVEEVERIEMKNKEAIEKY

KKQGKNLHFIKNGDDNGAYNIGEKIRELPKKSDVKNT

>MDR2345186.1 transposase [Planctomycetaceae bacterium]

MKNLTQFVNLYQLSKTLKFGLTLRNKIRKNGFEGEIYESHTELQELIKISEQKIIKETTD

KNKEIMTFTELPLDEIRKCLDDMHKYLDDWEQFYNRYDQIAVLKDYYRKLERKARFDGFW

REKNIKNKIQNNESETIKKPQSQVIKLSSLNNEYENKKRRDYITDYWNENIQKAKRKFYE

VNSVLKQFEVANEQNRDDKKLNEVVLRKLFLSFTNLINDTLEPLCNGSICFPDIEKLTNS

KTDEQLQRFVFDDGFKKILSEQIENLKIYFAINGGYVQYGRVTLNKYTALQQPNKVDEDI

KNIIKELGLLEFVKKYENTEQIINYIKNIKDKKQELNANNLSLIEKVQLFKYKTIPAGVQ

PSLIAYLARTEKKDKKTLRELFYAIGQPQSPSKDYKELQNKTDFNLYKYPLKVAFDYAWE

SLAKSKYNPHIDFPDVKCKEFLKDIFGTDISVNDNFKLYSALLFVRENLATLDHGNPNDK

NIHVNKVENTFKEIKDRLAKKEYKKEYKALEIICKWHKNSATIEQSEYEAAKQTIYEAAK

QTIGQLRGRQKNQISKFKELTDSFKKLAPKFGKAFANLRDKFNEEYEINKISHCGVIVED

RNNDQYLLLSQLNDNRENASDIFELEADPNGELKIYQVKSLTSKTLLKFLKNKKGSNTGF

HINENWTFPKGKWDVINKDKIFLNYVIQRITNSSMAKEQKWSNFKWDFRRCDSYEAIAKE

VDAKGYILESVNISKLTLNKLITEKKCLLLPIVNQDLTRQDKKTKNQFTKDWIKIFESNN

CYRLHPEFKISYRYPTPNYPKPEEKRYSRFQMISHLLCEYIPQNDNYKSRKEQVKIFNDK

VAQKESVEQFNQQFEITDDYYIFGIDRGIKQLATLCILNKNGQIQGDFEIYTREFDKVNK

QWKHTILEKRNILDLSDLRVETTVEGKKVLVDLSKVKLHSGNENKQTIKLKRLAYIRMLQ

YQMQHEQDKVLRFINQYKTIDEIEKNIRDLISPFKEGKQYADLPTEKIKDMLIQFGELSK

NDSDKSKKELCTLCELDAVDDFKTGAVANMIGVIAYLLEKYKYNVYISLEDLTRAFRLQR

DRLTNNILQSTNKDNTVDFKDQENLVLAGLGTYHYFEIQLLRKLFRIQRNSEGDILHLVP

TFRSVDNYEKIVRRDKKTDNDKYVNYPFGIVRFVDPKYTSKKCPICDKTNTTRKDNVLIC

NSCNAVSGEYETDNENRHYITNGDDNGAYHIALKALSLRKSKNLEKKK

>MDR0982475.1 transposase [Culturomica sp.]

METLNQFTGLYSLSKTMRFGLTLKEKKPKNDSIAVESLYQSHQDLKELVELSDKRIIEEK

KPEPPVENLGNPPIEKLRDCLNSMQKYLNDWRKVYTRYDQLAVLKDFYRKLERKARFDGF

WKDKKGQNQPQSQEIKLSSLKHKSGEKEIKDCIVTYWGENIRKANEKWHQVDSVLKQFEE

AKRKNRDDKKLNQVELRKLFLSLANLVNDTLVPLCQRSITFPNADKLSDNARDKSVLDFI

GDNEIREHLLDKITKLKEYFQDNGGYVPFGRVTLNQYTAMQKPNKTDKEIEDAIKNLGLS

IIKSQNFDAFEHIEEATDKVERLNTVSLPLVERAQYFKDKTIPVGVRDSLAKYLAKDDTA

KEKELIDLFEKIGMPKRPAKDYSDPTLKEKFDLRKYPLKVAFDYAWETVASKELHDDILK

NKCKKYLKDIFDVDTDKSIFFNIYSDLNYMKIILSRIEYPTQNQLSKDNFLEWNRKVITI

LDGDDFSHFNKNADGSTDKKMNTAKTYVKTWLDKLEANIEQFDGQDFKKFYEDFKKKNKN

SCKDFDDAKRDIGLKRGGLKQIIEETETFTDKKTGKQKPKYKDSKYKELTEAFKSIAVDF

GKHFATLRDKFNEENEINKIEYYGVIVEDENADRYLLLSKLSESREEIKNIFPDKAEGLK

TYKVKSLTSKTLTKLVKNKGAYKDFHISDMRVDFKKIKEEWSAYKNDQAFLKYLKKCLTD

SSMAQAQNWSEFGLDFDKCNTYEEVEKELDGKAYLLQETRLSKATITNLVKNKGCYLLPI

INQDLAREDRTAKNQFTKDWKQIFENKKHYRLHPEFNMAYRQPTPNYPNSEIGDKRYSRF

QMIANFMCEIVPQSTSYATRKEQIQTFNDNNKQQKAVKDFDSKFKLSDSYFIFGIDRGIK

QLATLCVLDQGGVIRGGFEIYTRHFDGNKKQWVHTSLERRNILDLTNLRAETTIDGKKVL

VDLSKVEIKNQTDNKQNIKLKQLAYIRKLQYQMQTNPEKVKNMSDEDIENDLKDIITPYK

EGTHYADLPIENIKAMLDRFKVLYGKTDQQSKQELKELCELDAADNLKGGIVANMVGVIA

HLMEQYNYRVKISLENLTTSFVNQSDGLNEYFISRGMDFKEQENAALAGLGTYQFFEMQL

LKKIFRIQQDDGNVLHLVPAFRSKEDYEKIIRRDKNDGDEYVNYPFGLVTFVDPRYTSRK

CPICGKTDVKRNDNIITCKKCGAVSGKYSFDDKNRQFITNGDENGAYHIALKTRKEVHNE

N

>MDR2755657.1 transposase [Planctomycetaceae bacterium]

MLQQFEKAKQQNRDDKKLNEVELRKLFLSLANLINETLEPLCNGSICFPDSEKITDREND

KKLQNFVFDNNFKEKLSKQIENLKIYFRQNGGYVPLGRVTLNKHTAQQKPNKVDNDINKL

IEKIKLQKFIKEYTNDKQLNDFIKTTKNKSEKIKDNELSLIEKIQLFKYKTIPVGVRHEL

SEYLFKICKIDKEQIKKLLQEIGKPQSPAKDYKDLQNKEEFDLHKYPLKVAFDYAWESLA

KNEYHTDIEFPKEKCQEFLREVFNINIGNNKDFKLYSALLFIRENLATLDHSKPNNIDIY

IKNVETTFKKIGLEKHTKTINTIINWHKKLSNIQQLDYDKAKQTIGQLRGGLKNTIKCFK

DLTEYFKVLAVEFGKKFAELRDKFNEEYEINKISHYGIIVEDTNADRYFLLTELSESREE

VENLFTNENKGFKTCTVKSLTSKALWKLLHNNKGISEDFHTKNWNCPKKRENLSHEENLQ

YIIECLKESAMAKVQNWAEFGWNFDKCKTYEDVEKEVDRKGYKLQESGRISKETIESLIK

DKNCLLLPIVNQDIARKDREAKNQFTKDWEKIFATNNGYRLHPEFKISYRHPTPNYPKPE

EKRYSRFQMIAYLMCEYIPKNNTYTSRKEQIKFFNDKEKQKDSVEKFNNMIETKNDYFIF

GIDRGIRQLATLCVLNKNGQIQGGFEIYTREFDYVCKQWKHTLLEKRNILDLSNLRVETT

IDGKKVLVDLSEVAVKIKDEHGKYIKDEEGKYKTDKNNQQKIKLKQLAYIRKLQYKMQTE

QERVLSFIKKYKTDQDIEQNIKELITPYKEGKNYTDLPIEKIKDMFIQFTELTSKNDDKS

EKELREFCELDAADDLKTGVVANMIGVIVYLLEKYNYNIYIALEDLTRAFYEQKDGLSDV

ILSNTNKDKTVDFKDQENLVLAGLGTYQYFEMQLLKKLFRIQQNNGNILHLVPTFRSVDN

YEKIVRQDSKTVGNKYVNYPFGIVQFVDPKYTSEKCPKCGKTNTTRKENILTCKECAFKT

PCNQAGEQNIHYIQNADDNGAYHIALKVLENLEKPQKRV

>MDR2342477.1 transposase [Campylobacteraceae bacterium]

TKLKLNEVNLVLKQFEEAKQQNRNDKKLNEVALRKLFLSLTNLIKDTLEPLCNGSICFPD

IEKITNSENDEKLQSFVFDNGLKEELSKQIESLKIYFEQNGGYVPYGRATLNKYTALQKP

NKVDDKIEVLVKKLKLLDFIKKYKNSKQLNDFIKTIKNKSEKIKDSELSLIEKVELFKYK

TIPIEVQNNLSEYLSKTSRINEEQIKKLFGEIGKSQSPAKDYKDLENKEKFDLCKYPLKV

AFDYAWESLAKSKCHHDIEFPKEKSQEFLEKVFDIDVNNNENFKLYSALLLIQENLATLD

HSKPNDRNVHIANIENIFENISLKEKDFKAAKTIINWHKKLPNINQSDYDKAKQTIGQLR

EGLKNKISKFKDLTNAFQELSQKFDKAFAYFRDKLNERYEINKISHYGIIVEDANNDHYF

LLHKLSENREEVEFLFPNEAEEFKTYTVKSLTSKALWKFLHNKKGISENFHTNQNWHCPK

KREHLSDERNLKYITNCLQESEMSKEQNWAEFGWNFDNYKTYEEIEKEVDSKGYKLQISH

LSQETISFLINSKNCLLLPIINQDISREDKKAKNQFTKDWEKIFADDNGYRLHPEFKISY

RHPTPNYPKPEEKRYSRFQMIAHLICEYIPQNSNYISRKEQIKTFNNKDIQKESVENFNK

QLDIKDDYYIFGIDRGIKQLATLCILNKDKQIQGDFEIYTREFDKDKKQWKHSLIEKRNI

LDLSNLRVETTVDNKKVLVDLSEVRVKIKGEDGKYVKDNNGEYKKDKNNRQNIKLKELAY

IRKLQYAMQTESENVLRLANQCKTEQDITKENMENLITAYKEGRMADDIPKSKIFDMLRQ

FKQFSDNKDEKSKRELIELDPTGNLKTGIVANMIGVVTHLLEKYGYQVYISIEDLTKCFY

VQKDGLSRADLTRDMSFKVQENAVLAGLGTYHYFEMQLLRKLFRIQQNNGDILHLVPAFR

SVDNYEKIVRRDKQTNGDEYVNYPFGIVQFIDPKYTSKRCPNPKCEKTNTTRIKNILTCK

ECGFKTPWDQVNEQNINYIQNGDDNGAYNIALKALKNFEKQK

>MDR0305996.1 transposase [Chitinispirillales bacterium]

MNQEVQGKQYQFSKTLRFGLTSTNQNLYSEETMRLLKVSQEKIEKQVKKENNNTDKTNQL

RNCLVQIKEYLKTWDNTYPQIDFLAITKDYYKVISRKARFDFDKGNGSEIKLSSLQSMYN

NKKRYQYITDFWKENLHKTENLYRKSDDLLRIFEEAEKQNREDKKLNKVELRKTFLSLFN

LVNESLKPLIEGNLFIVNDEKIDEQNPKHNYVSDFILKAEARKPLYNCIGNLQNYFKDNG

GYVPFGRVTLNKWTALQKSNNRDTKINRIIKELKINSFLIKNINYKYNEFTSNFKEKKDK

KGKIVKNKDGDIVWELEPNDKSVIELCQFFKYKKIPINACLNLAKRLIKENKLEKEKENT

FLSELGVSKSPALDYKKDQSNFSLTNYPLKVAFDYAWENCAKAKYEDIPFPKKQCEKYLR

DVFDLDIETNADFAKYALLLRFKILIGRIKVEETTRIENIATIKEFFNDVKSNLTKEKDK

TVAEINNWLTFKENQTDKKAKYSNQDEFSEAMKTIGEERGGLKSKISRYKALTDMFKVCS

SKFGKQFADLRDYFNEAYEVDKIKYRAWIIEDDKKNRFVLLADKGKEVGLTSGNGDLYFY

EVKSLTSKSLVKFIKNKGAYPDFHNKKSEDGFCQIYLNSENKENKDRFIDDVKIHWSTYK

NDQEFLKKLKECLKNSKMAIEQNWNEFNFDFSECDNYEKLEKEIDRKGYKFERKAISLTD

ITDLVENKECLLLPIVNHDINKEKQTENQSQFTKDWFAIFKNKKHLHPEFNIFYRFQTKD

YLKTKFKNGTEKTKRYSRFQMLAHFGCEVIPQGDYLSKKEQIAIFNDDEKQKKEVENFKE

NISSDFDYVIGIDRGIKQLATLCVLDKKGVIQGDFQIFTRKFNDITKKWEHKELEKRNIL

DLSNLRVETTIAGEKVLVDLASIKTKKGENQQKIKLKELAYIRELQYAMQTRKDELLDFA

NKINSADDITEDSIKNFISPYKEGTRYADLPKSEFFNRLTEWKNADDKGKLKVAELDSAD

NLKSGIVANMIGVIAFLCEKYKYKVRISLEDLTRAYGIQKDALSGTAIYQNDEDFKEQEN

RRLAGVGTMQFFEMQLLRKLFKIQIDEKLCLIPSFRSVANYEKIVRRDRKSSGDKFVNYP

FGIVCFVDPSYTSQKCPYCDNKHKKNDKETGKKAFYRDKGENKNSLLCKQCGVSTIKGQE

KPSNKNDSKKQFNIHFITNGDENGAYHIAKKTLNNLIPNNKNNKNQPSDFPIGTCT

>GHT58552.1 hypothetical protein AGMMS50239_03340 [Bacteroidia bacterium]

METNKTTKAINEYQTQKTIRFGLTVTNNNLYSENIVKLLKCSEEKIKEQLKKTQTDDLQN

QRLRCCLIEIKEYLKTWNNVFSQIDFLAITKDYYKVLSRKAKFDYDKGNGSEIKLSSLQS

KQSKYNDKKRYQYILDFWHENFIKVENLYRKSDDLLKVFEEAANQNQDDKKLNKVDLRKT

FLSLFNLVNETLKPLIEGNLFIVNDDKIDEHNSKHNFVSDFIVKTEERKQLHDCITDLQD

LFKANGGYVPFGRATINKWTALQKSNHKDDEIKRIIRELKIENISMQNIDYKYKYDSFAE

NFKQIYNKEGEKVWVLQFDANSVIKVCQYFKYKKVPINARLNIAERLIKEKSWQREKKND

FLSEFGISKSPALDYKNDKENFNLANYPLKVAFDYAWENCAKAIYETTTFPKEHCEKYLK

EVFDLDIANNACFTKYALLLRFKILICRIKSEETTQIQNIEAVRGILDEINKNISGRQDF

SKAKIITEINNWLSFKEKQTDKKEKYSNQDNFSLAMQIIGQERGGLKSRIEKYKTLTDMF

KVCASKFGKQFADLREYFQEAYEVDKIKYRAWIIEDEKQNRFVLFANKEREIDLTSEEGN

LYFYEVKSLTSKSLVKFIKNRGAYADFHKLKNNFNYEKIKRDWQYYKNDKYFIQNLKDAL

RNSKMAIDQNWAEFKFDFTKFNTYEDIEKEIDRKGYKLVCKTVSLNTLKDFVENKGCLLL

PIINQDINKDDKQAKNQFTKDWNSIYDNKKRLHPEFNLFYRFPTQDYPNTKFSNGTEKTK

RYSRFQMLAHFGCESVPKGDYLSKKEQIAIFNDDAKQKDAVEKFNNSIASDFEYIIGIDR

GIKQLATLCVLNKNGQIQGDFEIYTRTFENKQWKHTLSEKRNILDLSNLRVETTIDGNKV

LVDLASITTKNGENQQKIKLKQLAYIRELQYSMQTRRDDLLDFAKGLQSADDILKDIRNF

IVPFKEGGQYADLPNERIYNLLKEWRDADDEAKRKIAELDPAQDLKSGIVANMIGVVAFL

CEKYGYKVRISLEDLTRAFGIQKDALSGIAIAPNDEDFKEQENRRLAGVGTYQFFEMQLL

KKLFKTQVDKNLHLVPAFRSVDNYEKIVRRDKKTNGDEYVNYPFGIVRFIDPKYTSKRCP

KCGKTDVNRNQKTNIVKCNNCEYETKAGNSSEANNIHFITDGDQNGAYHIAQKALKIQKE

Q

>MDD4848569.1 hypothetical protein [Bacteroidales bacterium]

MKNITNKYQITKTLRFGLSQKGKTKKEGFDGEIYQSHQEFNKLVSVSEARIKKSVTTEQK

TELALSIDNVARCLNNISDFLINWQRVYYRTDQIALDKDYYKIMCKKIGFEGFWFETNRR

TQQKIKKPQSRIISLSALDKKDGLGKERKQYILDYWKENLLSAAEKYEVVSEKLKQFQDA

LNINRTDNKPNEIELRKLFLSLTHIVYDILQPLCYGQICFPKIEKLDNTKEDNKKLIEFA

SDYQSKSDLLSEIAELKQYFEENGGNVPFCRATLNPKTLVKNPKSTDNSINEEIKDLGLK

EILKTYKDVLNYNNYLESLSAKQKLQLLNDRNTSIITRSLLFKYKPISANVQFDIAKTLS

PEVGKGEEDLRAFLRGIGQPKSPAKDYADLQNKSDFNIEAYPLKVAFDFAWESLARAIYH

ADSDLPMDACKNFLQDNFKVKNDDTNLKLYAQLQELKAVLSTLENGNPNNAAAFRLKATN

LLNEIPWKTVGNYGQQNKDEISKWLNNGKNKDDYKKAKQQIGLFRGRLKNNIQGFDNITQ

TNKNIAMKMGRTFATMRDKITGAAELNKVSHYAMIIEDRNTDRYVLLQPFTENEQDRIYS

QTDYNNGDYTTYEVNSITSGAIAKMLRKARIDELSKNDNNRNLTSQPELTEEKKEKRNIK

EWKNFIENKRWDLEFQLKLNEKNFEQIKKEVDTKCYNLRTKKINKTTLEDLVNKSDCLLL

PIVNQDLAKEEKTNGNQFTKDWNSIFAQNTPWRLTPEFRVSYRKPTPDYPISDKGDKRYS

RFQMIGHFLCDYIPKSDKYISNREQILNYKNDELQKKAVKDFHEDLKGKTEEENQNESMN

ALMAKFGNVNKKQKATTVEKPKEKFYVFGIDRGQKELATLCVIDQDKKIVGDFDIYTRSF

NSERKEWEHTFFEKRHILDLSNLRVETTASIDGKAEKKKVLVDLSEIKVKDKNGNYSKPD

KMQIKMQQLAYIRKLQFQMQTNPEGVLAWFKENSTKDLIINNLVDKKNGEKGLISFYGSA

IEKMEDTLPVDRIEEMLQKFAALKKQEKEGEDVKLSIDQLVQLEPVDNLKNGVVANMVGV

IAYLLQKFNYQVYISLEDLSNPFGSQITGGIAGVPLKQGKDEGRRMDVEKYAGLGLYNFF

EMQLLKKLFRIQQDSCNILHLVPAFRAQKNYDHVAVGKEKVKGQFGIVFFVDANATSKTC

PVCGTTNNKPNNQKYPNAKKGLSADGKEVWLERDKSNGNDIIRCFVCNFDTTKEYTENPL

KYIKSGDDNAAYLISAAGIKAYELATTLINNQ

>MDX1959256.1 hypothetical protein [Leptospiraceae bacterium]

MEKYQITKTVRFGLTATNSNLYSDELKDLIETSEIKIKESLKNKSHNSLQIEQLRSCLNG

VKEYLKTWNNVYSQIDFLGISKDYYKVISRKARFDFDNKGLGSEVKLASLQSKYNSKKRI

QYILDFWEDNFQKTEILYRKSDELLKVFEEAEKQKRDDKKLNEVELRKTFLSLFNLVNES

LKPLVEGNLFTINDDKIDSRNQNHEVIADFISNTKVRTELYESITELQNFFRDNGGYVPF

GRATFNQWTALQKADKNGEREIDKIIKQLNLETVSMANIDYKYNTFTKNFEQGGQVWKIK

QNAKSVIELCQFFKYKKVSITTRLNLAKRLNKTNNFLSEFGISKSPALDYKKDKENFNLA

NYPLKVAFDYAWENCAKAKHESITFPELQCKDYLHNVFGVDANKDKNGKIKNEELNKYAD

LLQFKILLGRLKAEFHKAAEETNKNNIRKLKNIFENLDYSGVQDFNKNKIKEIVEVWFAN

KEKNIGKKKEEMIPLTEKKKDDFSKAMQIIGQERGGLKSRIKKYKTLTEMFKVCASRFGK

LFADLRDYFNEAHEVDKIKYRSWILEDGKQNRFVLLVDKAKDLELENEENGELKLYEVKS

LTSKSLIKFIKNKGAYPDFHSLNSFNSDEIKKNWTNHKANINFLKNLKSALENSLMAINQ

NWKEFNFDFSRCDTYEQIEKEIDRKGYILKQQNISLNTIKKSINEEKSEKINNSKKLPSL

LFPIVNQDINREAKQEKNQFTKDWFEIFAEENNLHKKRLHPEFHLFYRFPTKNYPNTKFK

NGKEKSKRYSRFQMLAHFGLEVFPQGDYISKKEQIEIFNDDKKQKEAVEKYNNSIVSEVE

YIIGIDRGIKQLATLCVLNKNGVIQSGFQIYTPSFNHDTKQWEHSFLGKRNILDLSNLRV

ETTIKNEKVLVDLASIQTKKGENQQKIKLKQLAYIRELQYSMQTRQVELLEYAKTLNSAE

DITEEKIKIFISPFKEGSHYEHLPKQEIYNLLNEWQNADETRKRKIQELDPTDSLKSGIV

ANIVGVIAFFCEKYNYKVRISLEDLTRAFSIQKDALTGTPIHRNDEDFKEQENRRLAGVG

TMQFLEMQLLKKLFKLQSEKNKHLIPAFRSVANYEKIVRRDKENGGDEFVNYPFGIVTFV

DPRNTSQKCPYCNNIARKEDDAFYRNAGENKNSLLCKKCGLSTIKGKENKSNQDDSKNQF

NIHFITDGDQNGAYHIALKTLENLHRLNTPKVTKHTKTKWKK

>MDD6002420.1 zinc ribbon domaincontaining protein [Bacteroidales bacterium]

MEQYQLTKTIRFGLTKVRKEKKHLSHEELDELVMVSEERIKKEHPQAENQLDEQSFVKKI

GDCSKAIKEYLLSWGKVCRRIDKITVRKEFFKILARKTFFKYESKVGKKRTPLPSEVKLS

GQKGNNYYDEPINEGISQFWQNRVSKALKLHSQLESMLFDYKKAIETEKHNQENPKENND

QFDKLHLVDFRKMFLSVCSLVMDSLRPIVNELIIVSDNVSKDEDKYILDFVNDKSKQWDL

FKQIEDLQNLCKDNGGNIPFGKATFNKYTSEQAPNHRDNDIHKVIRELKIEEFVSDFIGL

EQEDIYRKIYQSTQHSLVNLNKPSISPIIRAQFFKYKPIPVLVRFGLANYLNKQQGKKYS

NRLKDIQELFRIFGTSKSPELDYSDKNNRTEFSLDKYPIKVAFDYAWERCARSKYAQKPV

DFPKEICVTFLETYFEYNSKEKNREAFEIYAHLLKVNECLATLEHFENEPPKDIKSLWQD

VQNHLDKVGKLCSNEDRKAITQWYEEYKNLWAKGNYKKLKKWIESKSYTVTNFTQAKMHL

GQKRGSQKTMVLKSYFHPTYGKIKDGNRFINSNVTEVFKNIASTFGKSFATIRDYFNEES

EVNKIEYGAVIIKDKKGDKYLLLQKKNEGGIDMPVFNESDGNGDYDVYQVQSLTYKTVNK

IYNSTKYPEFFAINGEKAIYAPNRPQRFKDDQEKNTFNEKKLQSLKKCLTESDFMTNTTE

NYLQKFNWTEEINNCTDFEPLAKIVDQKGYYLKSYKISKEQIAELVNNQNCYLLPIVNQN

ITAKTKNDTNQFTKDWNKIFNDEYKDYRLHPEFTMFYRYPTPDYPCPGEKRYSRFQMNVN

FLMEVIPSEGEYVSRKEQIESFNTPKQDKEDNDNENSQAKKVAQFNDNINTKKPSYIIGI

DRGINELATLCVINSEGKIVAVDENGLIKDEFDIYVKHFDKDNKCWIHNIKPKTATDNKP

RTILDLSNLRVETTIDGKQVLVDLSSDENGIQVNSKQIVHLKRLYYLRCLSYLLQSSDYK

SIILEKLKDVNNMTDDAIYEVFKNDKFIDSYKGGVQYTDLPYDEIRHLISTYQEIDQSNK

TDSEKQSELNTLCQLDATESLKKGVVANMIGVVVYILKKLNYDAYISLENLCRALYFSKD

SLSGYTIENTSVNPDLDFKDQENAKLAGLGTYSYFEIQLLKKLFKLQIDEKQFLVPGFRS

VENYEKIVKLGKVKHSIYQFGVVHFVEPANTSLKCPICGANGKRIKYNPNYDEDELVCKK

CGFRSNISKIQNSKIMEDSVIKTYYDNHNYKAIISGDTNAGFNIALRLLMNLNTQIENAI

NHLHKTGKNYHSVNK

>HHU47362.1 hypothetical protein [Bacteroidales bacterium]

MKQIKNQYQLSKTLRFGLTQKNKTKKENYAGEIYKSHSELSDLVEISEQRIKDSVSTNKN

SESSLPVDAIGKCLNQISEFLKGWQQVYQRTDQIALDKDYYKILCKKIGFDGFWFDKKNG

RKTKKPQARIISLLELEKKDDKETERKQYILDYWQENFINAVEKYNVVSEKLKQFEVALK

INRTDNKPNEVEFRKLFLSLVNIICDTLKPLCFQQICFPRLEKIDNSKIDNKNLIDFAID

YQSKNELLSLISKLKSYFEENGGNVPYCRATLNPKTAVKNPESTDNSIESEIKKLGLDKI

IKNNKDAFSFSYNLYNNTAEDKKSKLKDDENGGLIERSLLFKYKSIPATVRFEIAKTLSK

PDGKTEEEILEFLRDIGQLESPAKDYADLKEKDNFNIEKYPLKVAFNFAWEGLARAKYHP

EAVFPTEICKQYLKNHFKITEDNKDFVMYAKLLELNAVLSTLEKAKPTDEKKFSVAAKKL

LEEIEWEKVGKNGSKNKEAITKWLQTKSKTDKNFKSAKQEIGLFRGRIKNNIRIKNNIKS

EYSEITNVFKNIAEEMGKTFAEMRDKISGAAESNKISHYAMIIEDNNKDKYVLLQEFVEN

KNERIYAKSDSQKSDFKAYSVNSITSGAIVKMLKKIRTDKLKESNNFANTQPELTSKEKE

KRNIKEWKKFINEKGWNLEFGLKLENKTLEEIKKEVDAKCYKFDIKYFDKETLSDLVKNK

NCLLLPIVNQDLAKKEKNESNQFTKDWNAVFPQDTPWRLTPEFRISYRKPTPNYPKSDKG

DKRYSRFQMIGHFLCDYIPKTDSFISNRQQIENYKDDERQELAVKKFNAALRGRTKNEEY

KEQLNELAAKYSKNGQQKINVKTNEKFYVFGIDRGQKELATLCIIDQDKKIIGPHKIYTR

SFNSEKKQWEHKFLEERHILDLSNLRVETTVFIDGKPEKTKVLVDLSEVKVKDKVTGEYT

KPDKMQIKMQQLAYIRKLQFKMQNEPEAVLAWYEKNSTEDLILKNFVDNEDGTNNGLVSF

YGAAIEELKETLPIERIVDMLKEFKTIKKEEGKLTKEDEEGREKNKRKMDKLVQLEPVDN

LKNGVVANMVGVIAFLLQKFDYQVYISLEDLSKPFSSKIISGIDGVPIRVEKEEGRRADV

EKYAGLGLYNFFEMQLLKKLFRIQQDSENILHLVPAFRAMKNYDHIAVGKGKVKNQFGIV

FFVDAEATSKTCPRCGSTNQKPNKKDYPNAQQARLSNDKEGWIDRDKSNGNDIIRCFVCG

FDTTKEYTENPLKYINSGDDNAAYLISAEGVKAYELATTLADNI

>MBR4327012.1 hypothetical protein [Bacteroidales bacterium]

MAGKEVDAKYLSHEELDDLVMRSEYNLIKRNVLEWSKKKESDKSNYIFDELDKRFEEIIE

IEDRDQRNAEYVEFLNEFFKEVNHQTINQLDESTFINKIGDCSKAIKEYLLSWGKVCRRI

DKITVRKDYFKILARKTFFKYESKVGKKRTPLPSEVKLSGQKGNNYFDEPINEGISQFWQ

NRVAKALNLHSQLESMLFDYKKAIETEKHNQENPKGDNGSFDKLHLVDFRKMFLSVCSLV

MDSLRPIVNELIIVSDNVSKDEDKYILDFVNDKKTQWDLFQQIENLQTICKDNGENIFFG

KATFNKYTSEQAPNHRNNDIAKVLRELKIEKFVSDYIDLDQEAINRKIYQSTQSRLENLN

NPQISPIIRAQYFKYKPIPTLVRFGLAKELAKQQGKKYSDRLKGIQELFRIFGSSKSPAL

DYKNNRTDFSLDNYPIKVAFDYAWEMCARSEYAQKPVDFPKSICEKFLEKFFECKSNEKY

QQSFVTYARLLKINEDLATLEHFENEPPKDIESIYQDAQRYLDEVGNLCSNEDRAAIAKW

YEEYNKLWTKGDHKKLKEWIVSKSSIITNFTQAKMHLGQKRGSQKTFVLKSYFHSSYGKI

RDNNRFVNSNVTEVFKTIASTFGKSFATIREYFNEESEVNKIEYGAVIIKDKNGDKYLLL

QKKNEGGIDMPIFNKSDENGDCDLYQVKSLTSKTVRKIIASPNKYNDFFVNNDGKKIIYP

DKTDFKYKINPYDKEEVKKRKRELYNNDLVRPIIYSLTQSKFANKQNFEKYFDWTKALKQ

CSNIEQLYKTIDQKGYSLNPSKISKEQIADLVNNMNCYLLPIVNQNITAKTKNDTNQFTK

DWNKIFNEVDKDYRIHPEFTMFYRYPTPDYPKFGEKRYSRFQMNVNFLMEVIPADGEYCS

RKEQIEIYNAPKDNENCQKNVVERFNNKIKALKPSYFIGIDRGINELATLCVIDKEGKIV

GDFEIYKREFDSNLKRQKYTSIETRDILDLSYLRVEKDENGESRLVDLSESEVWIDSLDH

DKGKRANKQIVHLKHLYYLRCIAHLLQSSDYKSIVLEKLKDCNNLKDEEIKKVFEKDKFV

DSYKGGEAYTDLPYDEIRKLISDYQEIEQSNQTESEKSKALNTLCQLDASEYLKKGVVAN

MIGVVVYVLKKYNYDAYISLENLCYAYGYSKDTLSGYSITSTKEDPYLDFKDQENAKLAG

LGTYSFFEVQLLKKLFKLQIEENTELIPAFRSVDNYEKIFLLKNIDNKIYQFGIVYFVDP

KYTSLCCPICGEHGKKNVDRKKHTKKYDEDELVCKQCGFHTNLSHIETRVMEDKTIKNSY

DECNLKAIVSGDANAAYNIAIRLGKNIYSTIADKVKDLHHEGKKYIIVKG

>MEE1259578.1 hypothetical protein [Paludibacteraceae bacterium]

MGKNENKYQLSKTLRFGLTLKEKISNNEKRPYQSHSQFRDLIILSENKIREGISTPQNRD

LPSFIHRIQNCTDFINDFIHDWWMILMHTGQIELDKDYYKSLTKKVGFVGFWYKENKKKG

GKTKQPQARNIPMGELRHLCPQNTKECATYITDYWKDLLITAANKLHESSEQQKKFIKAM

EQNRTDNKPNEIDLKKSFLSLVSVTMELLNPILNGQILFNKMDRLDMSKKSDNDFIDFVN

DHETVRELNNDIEEIIADFKENGNNVNYCKATLNPDTALKQHNNNIPNDIATDLEELMMD

SIVGNYDDVNSFMDNCVSNLSAKEKIRIIKDSNISLIYRAILFKYKMIPANVRRDIAQGM

AKKINKDEDNIYSFLCEFGTLRTPQKDYADLKDKNSFNLNDYPLKVAFDFAWEGLAKAQH

HDQSDFPFDLCRNFLQENFDVNLEENQDFLLYADLIELNALLSTLDKGNPADPDSIIKKA

LDMVEQINWDSFDEKRGNEYKKRGNEYKRVIINRLNFSKDDEKYERIKKEISMSRGRLKN

KIKKYDDLTSQYKRIAMDLGKKFASLRDKIIDANEDNKVTHYAMILEDSNCDKYLLLQKV

SNNNIYHCMSYDSSDPKAYYVDSITSSAIAKMIRKGTDTSKIKEYAKLEEKERERRNVDD

WCRFISKKEYDRRYQLNINNGLSFEALKKEIDSKSYILVKKNISVDSIRELVENEGCLLF

PIVNKDLTKERKTTEDNQFTKDWNMIFSGSETNWRLPPEFRVTYRNPVPGYPNDKFGSKR

YSRFQMNAHFVCDFIPSSNSYTSNREQIAIFKDENEQKKRVEEFNRTLSNINQKFYVIGI

DRGQKELATLCVVDQDKKIHGDFKIYTRKFNSERKQWEHYSLEGEKGTRNILDLSNLRVE

TTIMIDGKPERRQVLVDLSEVLVKDKDGNYTKPNKMQIKMQQMAYVRKLQFQMQANPTEV

LEWYEQNPTEELIIKNLVDKENGEKGLISFYGAALVELDQTLPVSKIKEMLEEFKILKQR

ESKKENVQKELNNLTQLEAVDSLKAGIVANMVGVISYILKTLDYNAYISLEDLSTAQSSP

EFASGISGAITKMSREEGRRIDVEKYAGLGLYNFFEMQLLRKLHHIQTDSGNILHLVPAF

RAQKNYDHIMVGKEKIKNQFGIVFFVDAAATSIKCPRCGAVNKEDFHPDKQKYPDAEKGP

NLRNGKEQSGKKVWVTRDKGDEDRIKCYCCGFDTKQKNEGNPFMYIKSGDDNAAYLISDL

GVESYRKAYELAATVVEDRKKH

>WP_324313362.1 hypothetical protein [Xanthomarina sp.]

MNHIYNNYQVSKTLRFGLTQKQKIRRPGYTGELYESHKVLKELVKISEEKVKNLIVPAKN

EELLSSLDSVKWTLTEIREFLDQWRYIYNKSNQIALDKSYYLILSKKLGFNDEKKSRVIK

MIEIKDDIKEKIINYWAFNLNESNQKLLMVNEMVNTQLKALEINRTDHKINEIELRKALQ

SLFNTVLDILKPLVYREISFINLEKIEKDSKNSLLEKFATDFQRKIDLLEKIRSLKTHFS

ENGGNVSFCRATFNPKTAIKNPKSNDNSILKEIKKLGIKDILENNENVFYFEKKLAEITA

KEKLEYIIKDSESFLIRSLLFKYISIPAFLHHGIATELAPIISKEKNDLINFMISIGQIK

SPAKDYADIPNKNDFNVNAYPIKVAFDYAWETVAKSQYHHDINAPVSMCKTFLDENFENC

TKTKYFTLYSDLLELHTLLSTLDYGNPSMEDSIIDKANKIIAKIDNKEHKTKDKDLDKDI

DKYKETIKNRLNHKNFNDKQRYSDAKKELSQFRGKLKNENDIYRKLTESYKKIAMNTGKI

FAEMRDKISNASEQNKISHHALIIEDHNKDRYLFLQEFTTDKEKQIESICNDQAGQYIVY

WVNSITSKSISKMLSKKRIEKLKQKKIINNSIKTSILSDAEKEARDIKEWVSFIKEKGWD

IDFNLDLQNKNLEEIKKEVDAKAYKLKETLISQKTLSNLVKEGNCLLFPIINKDLVKKVK

TEKNQFTKDWNSIFKKDNLWRLTPEFRVSYRQATPGYPTSDIGTKRYSRFQMTAHFLCDF

LPQGTKYISNREQIENYKSSEKQKEAVEIFHQQIENDNNNVISTQSLNHLARHFGSKNIK

KKHNTIEKKFYVFGIDRGQKELATLCIIDQDKKIEGPFKIYTRSFNTKTKQWEHQFYEER

YILDISNLRVETSISIDGKPDQQKILVDLSYYKEGEKFIKLPKMQVKLQQLAYIRKLQYQ

MQRNPETVLDWSYKNTDDKSILENFVDKPNGEKGLVSFYGAAVIELKDTLPLSEIKDMLE

RFKELKGKEKNGEDVSQQLNELTQLKSVDHSKYGVVANMVGVIAHLLERYDYKAYISLED

LTKPYSAIDGITGQKTDAKSISGKQQDVEKYAGLGLYNFFEIQLLKKLFRIQKDSQNTLH

LVPAFRATKNYENLIAGEDKVKNRFGIVYFVDPKSTSIMCPSCGKTNNSSNKEKRVVRDK

KNGNDIIYCEFCGFDTRNDYKENPLKFIKSGDDNAAYIISTHTAKKAYELAKSIL

>GFI53642.1 CRISPRassociated endonuclease Cas12a [Alistipes sp.]

MKLEDFTNLYSLSKTLRFELRPIGKTRENIENGGLLRQDEDRAEKYVHIKKLIDEYHKAY

IDKQLSGLVLQYADIGKANSLEEYYHSTRKSKDSDKDKIVKIQDNLRKQIVKRLKDSDEF

KRIDKKELIQSDLAEFIKPAEDRALIAEFKNFTTYFTGFNENRQNMYSDKAISTAIAYRL

IHENLPKFIDNIETFDRIAGITELYDQTSSDAEIFRLEHFSETLSQKQIDAYNSVMGRYN

MLINEYNQTHKQSRLPKFKMLYKQILSDREHPSWLPEQFESDTAVLTAIRECYDDLRIPM

ANLKTLLEGLGNYDPSGIFLRNDQHLSQISKRLTGDRSSIERSVTEDLLTSRRLNKRKSR

TTDEEESRKLFKQKGSLSIGYIADTAKIDVERYFAKLGAINTVTEQSENLFAKAENARTT

ADELLANDYPAGKRLVQSNDDIALLKNLLDALMELQWFVKPLLGTGDEAGKDERFYGEFA

QIWEQLDRITPLYNMVRNYVTRKPYSTDKFKLNFESAALLGGWDKNKEPDCLSVILRKDE

QYYLGIINKNHKKIFENDILPCEGECYDKMVYKLLPGANKMLPKVFFSASRIAEFAPSDE

VKRIYNDKTFQKGEKFDLNDCRTLIDFYKASIDKHEEWNKFGFEFSDTNNYEDISGFFRE

VDRQGYKMSFRPVAASYIETLVEEGKLYLFQIYNKDFSAYSKGTPNMHTLYWRMLFDERN

LSDVVYQLNGGAELFFRRKSLQNGRPTHPANIPIKNKNSRNDKKESLFDYDLIKDRRYTV

DKFQFHVPITLNFKSDGAGRINERVREYLRSADDVHVIGIDRGERNLLYLVVTDMDGNIC

EQFSLNEICNTDYHSLLDEREHKRMQERQSWQAIEGIKELKEGYLSQVVHRIATLMVKYR

AIVVLEDLNFGFMRSRQKVEKSVYQKFEHMLIDKLNYLVDKKANPTTPGGLLKAYQLTDK

FESFQKLGKQSGFLFYVPAWNTSKIDPATGFVNMLDLGYESIDKAKTLLCKFDSIRYNAC

KDWFEFALDYDKFGSKATGTRTKWTVCTYGQRIDTYRNKDSQWVSRDVDLTNELKSLFSE

HGIDIYSNLKDAIVAQNDKEFFANMQRILKLTMQMRNSKTGTDTDYIVSPVADANGRFFD

SRQADATMPKDADANGAYNIARKGIMLVQQIKQSDDLRTMKFDISNKSWLRFAQHTNQAD

E

>MBK6607653.1 hypothetical protein [Leptospiraceae bacterium]

MEKYQITKSISFRLNKVKAPILEEQVEKIGDVKEAEASLLRLFSVGQDLAGLLKQYIYKD

NQKLKSSVTVHFRWLRDYTKDKFYNWKTESRRDFSREKRFKLSEVDYLKEVFEHFISDWD

AIIQNLGSEISRKQEALSRKAEIGKIIKRIGTRNVFLFFENLIIESKDKNEIHLEAQLNE

KIKVFKENLIKAEQWALPAQSAGIELAKASFNYYTINKKPKDFELEKRNLEKQLHNYDER

FMKDIAQRENKLFGELRIIEKDDATNTYRFLYELIQNGEGKQVLEKTDKGLTVDLDSLYK

NLKVFRAEQKKSFQEAVSKGSSYDDTAKINILFSSVDQKNNEKNLKAFNKYKEFTDTIQK

LADQKNKVERDSLNSRKLQEEISRIRQYRGKLLFQGPSKRGEPEERFDKYLKFCKVYKDI

ALKRGRLVARLKGIEKEKIDSQRLKYWTFIIEKDKQHQLVLIPKSQSQKVYQKISSYESL

GAGSSDIYYFESLTYRALKKLCFGVNGNTFIPEISEELRKAGKWNYEERDFGEHIFKIRS

GVEEIRDEKRLIAFYKDVLGTDYIRKNREGGKIVLPSSFDEEVLIPEFSTEQEFHAALEK

CCYLRKKKIPKDDIEKIFSECSAQIFTITSYDLEKQDKTNLKAHTKLWFDFWTGENEAQR

FPLRLNPEIRILWREPKATRIEKYGKDSKLFDPKKRNRYLHPQFTLTTSFIENALHPEIN

YRFLDAKQKKDNVKKFNLDINAKLKPLPKAWYYGIDTGIVELASLCLIQKNGIPQTFEVL

NLKEDKMNYDKTGYLKDGTKKKYKAIQNLSYFLNEDLYKRTFQDESFLDTFAEIFEKKEV

SALDLSVSKIISGHIVMNGDITARLNVALANARRKIVDALIKNPSVQLSEQDYNIKVGDE

IIFKGRIEFNAIKSWEDIQAIIHSSFEKDKENLARVQEDINKYRQVIAANVTGVIHFLYK

KFPGLITIEYLDQSEVESHRKDFEGIIDNPVERALYRKFQTEGLTPPVSELWSIKNKNEQ

IGIIYFADPKDGKKGAHGIRCPKCGEKAYPRSEDDGYKKDKNNKIFQCKQCGFHNLNNSM

GLLGLDSNDKVAAYNIAKRGLEILK

>HRG77638.1 hypothetical protein [Leptospiraceae bacterium]

MEKYQITKSISFRLNKVKAPILEEQVEKIGDVKEAEASLLRLFSVGQDLAGLLKQYIYKD

NQKLKSSVTVHFRWLRDYTKDKFYNWKTESRRDFSREKRFKLSEVDYLKEVFEHFISDWD

TIIQNLGSEISRKQEALSRKAEIGKIIKRIGTRNVFLFFENLIIESKDKNEIHLEAQLNE

KIKVFKENLIKAEQWALPAQSAGIELAKASFNYYTINKKPKDFELEKRNLEKQLHNYDER

FMKDIAQRENKLFGELRIIEKDDATNMYRFLYELIQNGEGKQVLEKTDKGLTVDLDSLYK

NLKVFRAEQKKSFQEAVSKGSSYDDTAKINILFSSVDQKNNEKNLKAFNKYKEFTDTIQK

LADQKNKVERDSLNSQKLQEEISRIRQYRGKLLFQGPSKRGEPEERFDKYLKFCKVYKDI

ALKRGRLVARLKGIEKEKIDSQRLKYWTFIVEKDKQHQLVLIPKSQSQKVYQKISSYESL

GAGSSDIYYFESLTYRALKKLCFGVNGNTFIPEISEELRKAGKWNYEERDFGEHIFKIRS

GVEEIRDEKRLIAFYKDVLGTDYIRKNREGGKIVLPSSFDEEVLIPEFSTEQEFHAALEK

CCYLRKKKIPKDDIEKIFSECSAQIFTITSYDLEKQNKTNLKAHTKLWFDFWTGENEAQR

FPLRLNPEIRILWREPKATRVEKYGKDSKLFDPKKRNRYLHPQFTLTTSFIENALHPEIN

YRFLDAKQKKENVKKFNMDIYAKLKPLPKAWYYGIDTGIVELASLCLIQKNGIPQTFEVL

NLKEDKMNYDKTGYLKDGTKKKYKAIQNLSYFLNEDLYKRTFQDESFLDTFAEIFEKKEV

SALDLSVSKIISGHIVMNGDITARLNVALANARRKIVDALIKNPSVQLSEQDYNIKVGDE

IIFKGRIEFNAIKSWEDIQAIIHSSFEKDKENLARVQEDINKYRQVIAANVTGVIHFLYK

KFPGLITIEYLDQSEVESHRKDFEGIIDNPVERALYRKFQTEGLTPPVSELWSIKNKNEQ

IGIIYFADPKDGKKGAHGIRCPKCGEKAYPRSEDDGYKKDKNNKIFQCKQCGFHNLNNSM

GLLGLDSNDKVAAYNIAKRGLEI

>HIZ28593.1 type V CRISPRassociated protein Cas12a/Cpf1 [Candidatus Adamsella sp.]

MEKFYDKFNNLFSVSKTLRFELEPIGKTKEFFEKYILKEDERKSDLFKKVKKYCDEYHKL

FISECLKNFDDKDFENLLKEYFALVNLSNPSEKETKRITDILKVLRAKISDRFTKNPKYK

GLFGKELINDYLKDYYKNNKNILDEITPFKDFTSYFTGFNTNRKNMYSKEEKHTAIAYRL

IDENLSTYIKNIKAFNIILENIPDIKIQIKENLNLDCDEFFSNIENYTNVLTQEQIETYN

LAISGKSQENSKIKGINEIVNLYKQKNKIKIPKLKELYKQLLSDTTSASFKFDIIEYDYQ

IVEMINSYYETFKKNFIDDNGCQSALSNIQNYDIKNIYINNDVSLTSLSQNIYNDWNYIN

SLLGNEYDQNYKGKAKFGTEKYSEKKEETLKKIKEISLLKIEELIKNYDKENNNSEKIFE

YFKNAIEENLKKIDIEYKNTKTILQKEYDKTSTDLLKDEQSIEKIKNFLDSIKDLQILIK

YLIPKNNTLETDLEFYNALNYDILSEIIPVYNKTRNYLTKKTYSLEKFKLNFDCPTFLNG

WDLNKEEANLGTLFRKDGNYYIGIMNTNSRKSFIEYNQAEFKNEKKYQKIEYKLLPNPNQ

MLPKVFFSESRIKEFNPSEELLSKYERGYHKKGADFDLNFCHQLIDFFKSSINKHEDWSK

FNFKFSETNKYNDISEFYKEVELQGYKISFTDISEEYINKLVEEGKLYLFRIYNKDFSSY

SKGTPNLHTLYWKALFDEDNLKNIIYKLNGGAEIFYRKKSIEAKITHPKNQPIDNKNPNN

PKKQSIFEYDLIKDKRFTLDKFQFHVPITLNFKAKDTHKLNDLVNNKIKNTEDINIIGID

RGERHLLYACVINKNGEILEQYSLNTIGQTNYHYLLSKKEDDRLKQRQAWGTIQNIKELK

EGYMSQVIHKLTKLMVQYNAVIVLEDLNSGFKNSRKKVEKSVYDKFETALINKLSYLVDK

SIQNKFEQGGIMNAYQLANADIKSAKQNGVIFYIPAWCTSKIDPTTGFVNLFDLKKIDKE

FVKKFNSIKFNNTENYFEFDFDYSNFSDKSSGDRTNWTLCSFGNRIRTFKNPKKNNNYDS

QTVELTQEFKKLFDCYNLDCSNLRDEILNKSDSKFFNATPEKNNFYGFALLFKLLLQLRN

SITGSEEDYILSPIKNKKGLFYDSRTANGVLPENADANGAYNIARKGLLILNRIKNSEEN

TKTDYTIKNADWLTYVQQQDR

>MBS1644465.1 type V CRISPRassociated protein Cas12a/Cpf1 [Bacteroidota bacterium]

MTNFQSFTQKYALSKTLRFELKPIGKTKDHIEAKGLLAQDTNRASAYQKVKKIIDNYHKY

FIDLAMQDVKLSKLSDFQELYFATADKKKDESYKIEVEKVQTDLRKEIVAGFKADRVKSI

FDKLFKKELITEILPSWLDENGGSKELVAEFKTFTTYFTGFHENRKNMYSDKEQSTAIAY

RLIHENLPKFLDNLKVFEKLKGVPELYAKCTQLYTDIEEYLNVRTIDEAFELAYFNEVLI

QKQIDVYNLIIGGRTAVEGTTRIQGLNEYINLYNQTQQDRTSRVPKLKPLYKQILSDRES

ISFLQDKFDSSQEVLASINEYYHANVISFQSAENEGAENVLTRIKELLDDLNRYNLAQIY

IRNDRSLTDISQALFGDFSIIKEALKFSFISTLEIEKKVLSKKQEESIEKYLKQSYFSIV

EIETALQQYKGENDALKDLGENVIASYFKNNFKTKVNDKDYDLVANVTAKYSCVKGILES

YPDDKKLNQEQKDIDNIKNFLDSLMLVLHFVKPLMLPKDSTLEKDANFYNTILPYYEQLQ

HLIPLYNKVRNFATQKAYSTEKIKLNFENSTLLDGWDVNKETDNTSILFRKGNDYFLGVM

DKKHNNIFRNPPKANTDNTYSKVNYKLLPGASKMLPKVFFSAKNIGFYNPSEEIQNIRNH

GTHTKSGTPQNGFEKKDFNLLDCRKMIEFFKDSIAKHPEWKQFGFNFSNTDSYDSIDGFY

REVEAQGYTISYTEIDDEYINQLVEDGKLYLFQIYNKDFSEFSKGKPNMHTLYWKALFEP

ENLNDVIYKLNGQAEIFYRKKSIHDDKKIIHKANEKVANKNPLNTKKESLFAYDLVKDKR

FTVDKFQFHVPITLNFKAKGNDFINQDVLSFLQNNPDINIIGLDRGERHLIYLTLINQKG

EILQQFSLNDIVNEHNGETYKTSYKVLLDKKEKERADARENWGTIENIKELKEGYISQVV

HKIAKMMVEYNAIVVMEDLNMGFKRGRFKVEKQVYQKLEKMLIDKLNYLVFKDKAPDEAG

GLYKALQLANKFESFQKMGKQSGFLFYVPAWNTSKIDPTTGFVNLFFTKYESVEKAQKFF

ENFDSICFNMNEKFFEFAFDYNKFTERAAGTKTKWTVCTQGERILTFRNPETNNQWDNKE

ITLSEQFEDLFGKYNISYGDGTCIKAQIVAQNEAKFFKELMDLFKLSLQMRNSITNSELD

YLISPVKNESGVFYDSRNADETLPKDADANGAYHIAKKGLMWLRQIHEFEGNDWKNLKFE

NTNKGWLNFVQNKNEKL

>MBQ4618127.1 type V CRISPRassociated protein Cas12a/Cpf1 [Clostridia bacterium]

MTKRDGFINQYSISKTLRFSLLPVGKTEEHFNHKLLLEKDKTRAQEYQKVKEYIDRYHKY

YIESVLKKAVLSNVSEYAELYYKSAKTENDFKTMEKLETAMRKQIAGWLTKTDTYKLLDK

REMIEEILPEFLKDADERATVEMFRGFATYFSGFYENRKNMYSAEPQSTAIAYRVVHENL

PKFLDNAAAFAKVKNALPAETMDELNATYQGLYGVSVEDVFSVEYFSFVLAQSGIDRYND

ILGGYTCEDQTKVKGLNEHINLYNQQVAGNDRSKRLPLMKALYKQILSERETVSFLPEQF

ASDNAALLAVKRFYEDVMDSSAQDAQTLFAELETFDPDGIFVRSGPAVNELSNAVFGSWS

AVVGAWEEEYRLAHPLKGKADPEKYEEKMKKAYKAIASFSLAKVQALGAQSATDEHSDKT

VVEYYKQAVAQALEDVKTAYEQAKGLLDSDYEATHVKKLCANDEAIDCVKTLLDAIKNVE

HLLKSLRGSGKEESKDSVFYGRFTPLYETLSSIDRLYDKVRNYVTKKPYSKDKIKLNFDN

PQLLGGWDRNKERDYRTILLRKNGMYYLAIMDKSNNKAFMSLPSTAGVEHYEKMEYKLLP

GPNKMLPKVFFAASNIDTFAPSPEILAIRKNETFKKGDKFKLEDCHQFIDFFKASIERHE

DWKQFGFRFSPTEKYRDISEFYNEVKEQGYSLRFQDVDAAYINGLVESGQLYLFQIYNKD

FSPHSKGRPNLHTMYFKMLFDERNLKDVVFQLNGGAEMFYREASIRKEEQIVHPANQPID

NKNPNNPKTQSTFVYDLIKDRRFTKRQFALHLPITLNFKADGKSYLNEEVRAAVKNAEDT

YVIGIDRGERHLLYICVINGKGEIVEQRSLNQILGEHGHKVDYHALLDKKEAERDAARKS

WGTIENIKELKEGYLSIVVHEICQLMLKYDAIVALEDLNFGFKRGRFKVEKQVYQKFENM

LIQKLNFLVDKTAEPTENGGVLRAYQLTNKVEGVHRGKQNGFLFYVPAWMTSKIDPTTGF

VDLLKPRYTSVADAKGFISVIDVIRYNQGEDLLEFDVDLDKFPRCTADFRKRWTVCSNAD

RIETFRNPQKNNMYDNRRVVLTEEWKALFAQYGVPLSEDMKEAILGVKETEFYRRFMKLM

GLTLQMRNSITGDSSVDYLTSPVRNRDGLFFDSRNYSDTAALPANADANGAYHIARKALQ

MIDVLKQTPEHELAKADLTISNADWLAYTQK

>MCQ4773245.1 type V CRISPRassociated protein Cas12a/Cpf1 [Lacrimispora saccharolytica]

MEDKQFLERYKEFIGLNSLSKTLRNSLIPVGSTLKHIQEYGILEEDSLRAQKREELKGIM

DDYYRNYIEMHLRDVHDIDWNELFEALTEVKKNQTDDAKKRLEKIQEKKRKEIYQYLSDD

AVFSEMFKEKMISGILPDFIRCNEEYSEEEKEEKLKTVALFHRFTSSFNDFFLNRKNVFT

KEAIATAIGYRVVHENAEIFLENMVAFQNIQKSAESQISIIERKNEHYFMEWKLSHIFTA

DYYMMLMTQKTIEHYNEMCGVVNQQMKEYCQKEKKNWNLYRMKRLHKQILSNASTSFKIP

EKYENDAEVYESVNSFLQNVMEKTVMERIAVLKNSTDNFDLSKIYITAPYYEKISNYLCG

SWNTITDCLTHYYEQQIAGKGARKDQKVKAAVKADKWKSLSEIEQLLKEYARAEEVKRKP

EEYIAEIENIVSLKEVHLLEYHPEVNLIENEKYAIEIKDVLDNYMELFHWMKWFYIEEAV

EKEVNFYGELDDLYEEIKDIVPLYNKVRNYVTQKPYSDTKIKLNFGTPTLANGWSKSKEY

DYNAILLQKDGKYYMGIFNPIQKPEKEIIEGHSQPLEGNEYKKMVYYYLPSANKMLPKVL

LSKKGMEIYQPSEYIINGYKERRHIKSEEKFDLQFCHDLIDYFKSGIERNPDWKVFGFNF

SDTDTYQDISGFYREVEDQGYKIDWTYIKEADIDRLNEEGKLYLFQIYNKDFSEKSTGRE

NLHTMYLKNLFSEENIREQVLKLNGEAEIFFRKSSVKKPIIHKKGTMLVNRMYMEEVNGN

SVRRNIPEKEYREIYNYKNHRLKGELSTEAKKYLEKAVCHETKKDIVKDHRYSVDKFFIH

LPITINYRASGKETLNFVAQRYIAHQNDMHVIGIDRGERNLIYVSVINMQGEIKEQKSFN

VVNKYDYKEKLKEREQIRDEARRNWKEIGQIKDLKEGYLSGVIHEIARMMIKYHAIIAME

DLNYGFKRGRFKVERQVYQKFENMLIQKLNYLVFKDRPADEDGGVLRGYQLAYIPDSVKK

MGRQCGMIFYVPAAFTSKIDPTTGFVDIFKHKVYTTEQAKREFILSFDEICYDVERQLFR

FTFDYANFATHNVTLARNNWTIYTNGTRAQKEFGNGRMRDKEDYNPKDKMVELLESEGIE

FESGQNLLPALKKVSNAKVFEELQRIVRFTVQLRNSKSEENDVDYDHVISPVLNEEGKFF

DSSKYENKEEKKESLLPVDADANGAYCIALKGLYIMQAIQKNWSEEKALSPDVLRLNNND

WFDYIQNKRYR

>RGD65337.1 type V CRISPRassociated protein Cpf1 [Lachnospiraceae bacterium OF096]

MENKRDENIYGRYEEFIGLNSLSKTLRNSLIPVGSTLKHIQEYGILEEDSLRAQKREELK

GIMDDYYRNYIEMHLRDVHDIDWNELFEALTEVKKNQTDDAKKRLEKIQEKKRKEIYQYL

SDDAVFSEMFKEKMISGILPDFIRCNEEYSEEEKEKKLKTVALFHRFTSSFNDFFLNRKN

VFTKEAIATAIGYRVVHENAEIFLENMVAFQNIQKSAESQISIIERKNEHYFMEWKLSHI

FTADYYMMLMTQKAIEHYNEMCGVVNQQMKEYCQKEKKNWNLYRMKRLHKQILSNASTSF

KIPEKYENDAEVYESVNSFLQNVMEKTVMERIAVLKNSTDNFDLSKIYITAPYYEKISNY

LCGSWNTITDCLTHYYEQQIAGKGARKDQKVKAAVKADKWKSLSEIEQLLKEYARAEEVK

RKPEEYIAEIENIVSLKEVHFLEYHPEVNLIENEKYAIEIKDVLDNYMELFHWMKWFYIE

EAVEKEVNFYGELDDLYEEIKDIVPLYNKVRNYVTQKPYSDTKIKLNFGTPTLANGWSKS

KEYDYNAILLQKDGKYYMGIFNPIQKPEKEIIEGHSQPLEGNEYKKMVYYYLPSANKMLP

KVLLSKKGMEIYQPSEYIINGYKERRHIKSEEKFDLQFCHDLIDYFKSGIERNPDWKVFG

FNFSDTDTYQDISGFYREVEDQGYKIDWTYIKEADIDRLNEEGKLYLFQIYNKDFSEKST

GRENLHTMYLKNLFSEENVKEQVLKLNGEAEIFFRKSSVKKPIIHKKGTMLVNRTYMEEV

NGNSVRRNIPEKEYQEIYNYKNHRLKGELSTEAKKYLEKAVCHETKKDIVKDYRYSVDKF

FIHLPITINYRASGKEMLNSVAQRYIAHQNDMHVIGIDRGERNLIYVSVINMQGEIKEQK

SFNIINEFNYKEKLKEREQSRGAARRNWKEIGQIKDLKEGYLSGVIHEIARMMIEYHAII

AMEDLNYGFKRGRFKVERQVYQKFENMLIQKLNYLVFKDRSADEDGGVLRGYQLAYIPDS

VKKLGRQCGMIFYVPAAFTSKIDPTTGFVDIFNHKAYTTDQAKREFILSFDEICYDVERQ

LFRFTFDYANFATHNVTLARNNWTIYTNGTRTQKEFVNRRVRDKEDYNPKDKMVELLESE

EIEFKSGQNLLPALKKVSNAKVFEELQKIVRFTVQLRNSKSEENDVDYDHVISPVLNEEG

KFFDSSKYKNKEEKKESLLPVDADANGAYCIALKGLYIMQAIQKNWSEEKALSPDVLRLN

NNDWFDYIQNKRYR

>MDD5770391.1 hypothetical protein [Candidatus Gracilibacteria bacterium]

MKNLQNLYEVRKTVRFLAQGKTQEIINFLEIDKKQGNDKKYDLLGSFLLKFESLVSLNNK

LFFYENSGNLKDSFKIKYKFLQNHTKYDFYDMLKKPEYIREKPKDYSFKDLNFIEKFLKE

YFELLDNIYNNLNNYQSAKKEKQNRYREIGFYLQKLSGKNISILKDLFEFLLKTNKKYID

DKIEEIKKIISEFEKDLVELLKEFGPNNGLEIAKGSFNYYILNKSSKSYFDDEILKLKNK

QKYNLPKDIFTLKERDKQDVYLNETFFQNIGLSKNELENMTLENAYNFLKDFKAKRKSAF

NQFVGNGYTHYDLTNKTKGKYKYKQNGELKEEEMSVELFDDIGETHFEDFKKLTDEINGL

GKEINDILQPLDKKRKDYEKEITKKKIEERTEEENNFMKLSKKVQDKAKERGKYFNIPNE

KIYTKNYKSLCEFYKNIALKLGLNKSKILGYEKQIVDSEKLKFFSVIVEKGNDKFLVLIP

KGGENHHKKAYDFLNDEKNNNSNGNINIYSFKSLTLRALQKLCFKAEDKEVLIGEENLNS

KERNTFIKNIIDELSGNQETQNKKYLYNGKIKRFDEFKIYDKYEKDKTKWQFNEKLLIEF

YKDVLKTKSAKKVLDLKDFGLDNILNENFETLDEFELTLEKVAYVKNITKLDENGYKNFI

EKNKAFEYKINSQDLSINDENHKNKKHTNYWLEFWTDHNKNKDYDIRINPEFSLYYTLSD

EDLKDKKENGLLKDGLKVGFKNRKLENQLIFTTNFSLNANEKLQNLAFLETPDLKENINK

FNENLETHISKNDLWYFGIDRGINELATLSITKFLDTKNENQVNKFEFAKINCLKLKDEY

DKILDIKVDEKGNTKYEFKKLEFEDKNGYKREFAASKNISYFLPKIEIFESITTGIIDLT

QAKLIGDKIVLNGDKSTLLKLKELSAKRRLFEYHREIDKNSEFEVVNKNKNSNGGLFIKL

INGNYIQIYWFNENLSITSQEKKELKNYKENLQSQLIEIFKKYLTFLSEENKFEDLETIL

KINHLRDAISSNIVGIINFLMIKKGYFGKIVLENLDETIGFTGLERNKTKNGKFENLKEY

TGEELKMIDGHFYQSNTDISRRLEWSLYRKFQETGFVPPEIKQSIFLKDDFGINKFGIIE

FVKTGGTSSTCPYCQNCNENFGNKDGLKKHHEKNDCGFTKQSGEYLEFKNKDNSINYDSI

ASFTIGKFAFEGKFGEFEKNKKIISYGEYKESHNKFESKNKTHIIEFKRKK

>MDR1615103.1 transposase [Campylobacteraceae bacterium]

MEKYKITKTIRFKLDYDKEESIKKLEKQVENLKKEQDFDLYIFVTECHKFIPRLKEYLYC

KDKNDKITNKLKKTIYIKKSWIRLYAQDTYYKSMQIDNKIREFSIGTIKEMNELIKKWFE

RFEELVQLLTEKANENEHNLSKHSAIGLLIKKMFARNNFPFISSFVDNSNDKSEIEVLSL

QLKNIKEKIESMLETGLKRYLPSQSNGFPLVKASFNYYTIFKNAVDFGKEIKKQKGNLII

NIQKLTSYFNITTNNQKIIECVIDDIKYIVQDKTLLLGDTPNFSNDNTVSLRQILKNIKS

VQKKTFNELMQKENFSYDNLKNNHKLYLFENISKNEFETYKNITQEITEKATKRDLSNSE

EYKRQLKSEMEELKQKRGKAINGADKSTKQQFIGYKSFCNFYGNVAHKHGIILTLIKGIE

KESVESQQLKYWAVIAEEENKHLLILIPKENNKAKECRDFIINKQNDEGNKKAYWFESLT

LRSLRKLCFGYVENKIKTNEFYLKIEEEFKGTKYQYFSNNKFLQGEYELKTEEKKIKFYK

DVLDSNYAKKVLNLPFEQLQNEVLNHDFNSLEDFAIALEKICYRRKVIYENDIIQKMQST

YDAQIFEIASLDLKNKEKINLKTHTKLWKSFWENENEQNGFNVRLNPEITISWRDAKESR

VKKYGKESELYDAKKKNRYRDPQYTLITTISENSNLPTKNISFIEQDEFKKQIDSFNENI

KKDDIKFIIGIDNGIVELSSLCVYIPEFKTHESISNQDKISNLETTKHKFPVLTITEGGL

GFSKYRNDKEYKIIKNPSYFLNKEFFVKTFKTTTLEYDEVKNKYFKEGSLLTIDLTMAKI

INGNIVTNGDISSFLSLKLMDAQKIIYELNDHIKEETAQKIIIKSGLELTEDEIDEINKN

RKEDKKYNNEKEGFNFFALVFIADHLSEIKEVLRNRIYFKYLIEQEQIKERLERYNVKRE

ITNEKLSLKINHLRKAIVSNAIGVIDFLYKHYKSILGGDGLIIKEGFDTQKTENDLKLFQ

GNIYRMLEWGLYQKFQNYGLVPPIKNLLSVRGEGIDDNDKNKIMRLGNIGFVSQAGTSQE

CPICGEKNKDKNKILENKKKKIFICENQNCTFSSISIMHSNDGIAAFNIAKRGFQNFMNE

RN

>MEG1642078.1 type V CRISPRassociated protein Cas12a/Cpf1 [Synergistaceae bacterium]

MAKSLLDFSGKYLLSKTLRFELKPIGKTEDYISKRNLISEDEARRENYKKVKKIIDRYHK

KFISNVLNAIPETGANLDWTELEKAIKNKTDDKYRENLEKEQKKKREEIIECFSGKSGAK

TKTLSEEEKNKEKEKQEFFKQIFSEKLFSEILVSEKNLLTEEEVEQLKTFNKFSTYFTGF

HENRKNMYSSEEKSTSVAYRIVNENFPKFLTNIEIYNKIKEIAPEIITETENELKDILNG

TKLDEIFNIKAFNKALSQEGIDYYNLILSGRTEKAGERKIQGINEKANLFRQQHKDNMPK

GTSFKMLLLFKQILSDRETLSFIPEKFLDDKEVLDTINALGDDFSKTNLYSEIRNLLSNI

DEYDTTKIYVENKNREKLSPELLGNWEALKNAQIAYKENEIGSPEKSTNLKKIEKYLKQK

TYSLKEINEAISFYMGEEKDYTKQYIDNLLSDISAIEKYQNKFNELKDKETNLLENENEI

EIVKNYLDAIEKIFFKIKMLQNGVEIEIDVDFYSQYDKLFEQLKQIIFVYDKVRNYVTQK

PYSEEKFKLTFKCPTLANGWDKNKEKDNLSILLLKDGNYYLGIMNKDKKLSEKELIGDES

VNQYRKITYKFLPGPNKMLPKVFLSKKGIETFNPSKEILENYKNGMHKGGDTFDLEFCHK

LIDFFKDSISKHEDWKNFGFEFSKTETYENISKFYKEVAEQGYKITFDNISEQRINKFVE

DGKLYLFQIWNKDFANGATGKKNLHTLYWESLFSEENLKDVIYKLNGEAELFFRKKSIEN

PIRHEIGSKKVNKRDSDGNAIPDEIYKEIYLHANGKKAINEINTTAREYIEKGNVRITEV

KHELIKDKRYTENKYLFHVPITINFKLPDKVIVNNEINKFLKDNKDINIIGIDRGERNLI

YVTLINRKGELLEQKDYNNINKTDYHAKLDTKERSRDEARKNWKTIEKIKELKEGYLSQV

VHEIVKMAIEYNAIIVMEDLNFGFKRGRFHVEKQVYQKFEKMLIDKLNYLVFKEKATTEP

CGILKGLQLTEKFDSFKNIGKQNGILFYIPASYTSKIDPVTGFANLFNLSKTTTQEKQKE

FMSKIDSIKYVKEEDKFVFEVDYDKFDTYQTSYKKKWSIYTNGKRIVGHKENGKWQQQDT

MPTEEMKKVLFSLNVPYEKGEELKDYIENREKEFAAKLYYIFKNTLQLRNSNAQTGEDYI

ISPVKDKKGKFFDSRTAGSLLPKDADANGAYHIALKGLCVLDKIDENLDENGEISYKEMT

ISTPDWFEFAQTRDIK

>KIE18642.1 hypothetical protein DS62_10740 [Smithella sp. SC_K08D17]

MEKYKITKTIRFKLLPDKIQDISRQVAVLQNSTNAEKKNNLLRLIQRGQELPKLLNEYIR

YSDNHKLKSNVTVHFRWLRLFTKDLFYNWKKDNTEKKIKISDVDYLSRVFEDFFNEWETV

IERINTDCNRPEESKTRDAEIAFSIKKIATKQMFPFIKSFVYNSNYKNSEETKSKLTALL

NEFETVLKICEQNYLPSQSAGIVIAKASFNYYTINKKQKDYKGYTDDIEKIEKGMNSKFH

YERKYDQLLEELNLIALKELPLIEFYSKIKSYKSTRKIEFSEAVSKGLAFADLKSKFPLF

QTESNKYAEFLELTGRITQISTAKSLLSKDNPEAQKLRDEIKKLRINRGEYFKNNFHKYI

SLCNLYKKIADKKGRLKGQVKGIENERIDSQRIQHWALVLEDNLKHSLILIPKEKVTEVY

RKVRASKADSTSSSSSLYYFESMTYRALHKLCFGVNGNTFLPEIQKELPEYNPNKQSDFG

EFCFHKSNTDKEIDEPKLISFYQSVLKTNYVKDNLNLPQSVFDEATVQTFETRQDFQIAL

EKCCYAKKTIISETLKKEILEDNNVQIFQITSLDLQRSEQKNLKAHTKIWNRFWTKQNET

ANYDLRLNPETAIVWRKPKKTRIDKYGAGTSLYDPKKRNRYLHEQYTLCTTVTDNALNNE

ITFAFEDTKKKGTEIVKYNEKINQTLKKEFNKNQLWFYGIDAGEIELATLALMNKDKEPQ

LFTVYELKKSDFFKHGYIYNKERELVIREKPYKAIQNLSYFLNEELYEKTFRDGKFQETF

NELFKEKHVSAIDLTTAKVINGKIILNGDMITFLNLRILHAKRKIYEELIINPQAELKEN

EKEYYLYFDKEGTEKVEKIYRSRLDFEHIKPYQEIRNDLNAYFKNVQKNEAKVEDQINQT

RRALVGNMIGVIYYLYQKYRGIISIEDLKQTKVESDRNKFEGNIERPLEWALYRKFQQEG

YVPPISELIKLRELEKFPLKDVKQPKYENIQQFGIIKFVSPEETSTTCPSCEKKAYELQK

EKKGEEKPAENKRYEADKKAGVFCCPKCGFHNRTNPMGYESLDSNDKVAAFNIAKRGFEQ

NFRQ

>Cas12h

MKVHEIPRSQLLKIKQYEGSFVEWYRDLQEDRKKFASLLFRWAAFGYAAREDDGATYISP

SQALLERRLLLGDAEDVAIKFLDVLFKGGAPSSSCYSLFYEDFALRDKAKYSGAKREFIE

GLATMPLDKIIERIRQDEQLSKIPAEEWLILGAEYSPEEIWEQVAPRIVNVDRSLGKQLR

ERLGIKCRRPHDAGYCKILMEVVARQLRSHNETYHEYLNQTHEMKTKVANNLTNEFDLVC

EFAEVLEEKNYGLGWYVLWQGVKQALKEQKKPTKIQIAVDQLRQPKFAGLLTAKWRALKG

AYDTWKLKKRLEKRKAFPYMPNWDNDYQIPVGLTGLGVFTLEVKRTEVVVDLKEHGKLFC

SHSHYFGDLTAEKHPSRYHLKFRHKLKLRKRDSRVEPTIGPWIEAALREITIQKKPNGVF

YLGLPYALSHGIDNFQIAKRFFSAAKPDKEVINGLPSEMVVGAADLNLSNIVAPVKARIG

KGLEGPLHALDYGYGELIDGPKILTPDGPRCGELISLKRDIVEIKSAIKEFKACQREGLT

MSEETTTWLSEVESPSDSPRCMIQSRIADTSRRLNSFKYQMNKEGYQDLAEALRLLDAMD

SYNSLLESYQRMHLSPGEQSPKEAKFDTKRASFRDLLRRRVAHTIVEYFDDCDIVFFEDL

DGPSDSDSRNNALVKLLSPRTLLLYIRQALEKRGIGMVEVAKDGTSQNNPISGHVGWRNK

QNKSEIYFYEDKELLVMDADEVGAMNILCRGLNHSVCPYSFVTKAPEKKNDEKKEGDYGK

RVKRFLKDRYGSSNVRFLVASMGFVTVTTKRPKDALVGKRLYYHGGELVTHDLHNRMKDE

IKYLVEKEVLARRVSLSDSTIKSYKSFAHV

>Cas12h2

MATRSFIRTGNLKAKNTAEEVMQWYADLQSDYRSFLNLFFGWMAIGYGTNAEDEVFYTSK

EESERLRSLTIGDAKKEQLAVSFIELLLKGGENASSCYNVFYRNYKSLGKAKLTQKKNDF

LSALPLLDENKIKEYFKTDEQLSQICIEEWLEYGVKNLPLPEIWAEVSPRLASIERSLGV

DLRLAFGLSCIRSRDCNYCRILIEMVGRDLRSIFEKYNNHLLETEKIKLSMNDKQGPVYD

SICCFAAELESKNSGLTKYVLTKGIDHVKKGTGEKTDIRLAVKELKKNKYRILIESSYSE

IMSAYSCWRTKKQLEKRKLYPCFDPNRNDYKVPVGQGSLGNFTVSVEDSGDVLIEIVGVG

VIRCAASCYFSGIVFDEIRNKNGRTGYSLNFCHKSISKGKKAVKAASHTGDKISGVLKEI

GLRNTDSGFFVSLPYSIHHDEKNFKIAEFFMSACPKKENVENLPDKIVVGAIDLNVSNPV

AAVKAVVYRDDKSGQLNALDYGSGNLIKKPFMLVANGPRIKNLIEIRDDARRVIGAIREF

KVSNAVKEHVGEDTRDFLILCGDTKSSSTRYLIQSWVKKINSRLRKIKFEMRSGGYRDCA

DNIRLIEAMDQCASMAESYNRIHLKSGEKLVKVAKFDKSRANFRNFVLRQLASKIANEMK

DCNVVFGEDLDFIFDSDKNNNALLRLFSAATLLKYIIEALEKIGVGFVKVAKNGTSQSDP

VTSNPGWRDDKNKSRLYVVRDKQLGWIDSDLAATMNILIQGLNHSVCPYKFYVKEYENKP

NSTQDSINAIKKPEEAIGKRIKRFFNLKYGSSVPKFVSDDRGRVTFAKKIDSTQTRLINQ

FVYAHSSCIVTCELHNEMVNKIKQLAVEKPNCQEFDVTCDPDGRYNNFALPEVHDSSKDV

GAKALTTKDVDFKTILKDHTA*

>Cas12l 8DC2_A Chain A, CasLambda [uncultured virus]

MASHKKTESNQIIKTFSFKIKNANGLSLDVLNDAITEYQNYYNICSDWIKDHLTMKISEL

YKYIPNEKKNSGYALTLISDEWKDKPMYMMFKKGYPANNRDNAIYETLNTCNTEHYTGNI

LNFSDTYYRRFGYVASAISNYVTKISKMSTGSRSKNISNDSDVDTIMEQVIYEMEHNGWT

SVKDWENQMEYLESKTDSNPNFVYRMTTLYEFYKSHIDEVNSKMETMSIDSLIKFGGCRR

KDSKKSMYIMGGSNTPFDITQIGGNSLNIKFSKNLNVDVFGRYDVIKDNTLLVDIINGHG

ASFVLKIINDEIYIDINVSVPFDKKIATTNKVVGIDVNIKHMLLATNILDDGNVKGYVNI

YKEVINDSDFKKVCNSTVMQYFTDFSKFVTFCPLEFDFLFSRVCNQKGIYNDNSAMEKSF

SDVLNKLKWNFIETGDNTKRIYIENVMKLRSQMKAYAIVKNAYYKQQSEYDFGKSEEFIQ

EHPFSNTDKGIEILNKLDNISKKILGCRNNIIQYSYNLFEINGYDMVSLEKLTSSQFKKK

PFPTVNSLLKYHKILGCTQEEMEKKDIYSVIKKGYYDIIFDNDVVTDAKLSAKGELSKFK

DDFFNLMIKSIHFADIKDYFITLSNNGTAGVSLVPSYFTSQMDSIDHKIYFVQDNKSGKL

KLANKHKVRSSQEKHINGLNADYNAARNIAYIMENTDCRNMFMKQSRTDKSLYNKPSYET

FIKTQGSAVAKLKKEGFVKILDEASVGSSGHHHHHH

>Cas12l DAE37039.1 MAG TPA: hypothetical protein [Bacteriophage sp.]

MAHKKNVGAEIVKTYSFKVKNTNGITMEKLMNAIDEFQSYYNLCSDWICKNLTTMTIGDL

DQYIPEKAKGNTYATVLLDEAWKNQPLYKIFSKKYSSNNRNNALYCALSSVIDTNKENVL

GFSKTYYTRNGYILNVISNYASKLSKLNTGVKSRAIKETSDEATIIEQIIYEMEHNKWES

IEDWKNQIEYLNSKTDYNPTYMERMKTLSAYYSAHKSEVDAKMQEMAVENLVKFGGCRRN

NSKKSMFIMGASNTNYTISYIGDNSFNINFANILNFDVYGRRDVVKNGEVLVDIMANHRD

SIVLKIVNGELYADVPCSVTLNKVESNFEKVVGIDVNMKHMLLSTSVTDNGSSDFLNIYK

EMSNNAEFMALCPEEDRKYYKDISQYVTFAPLELDLLFSRISKQGKVKMEKAYSEILEAL

KWKFFANGDNKNRIYVESIQKIRQQIKALCVIKNAYYEQQSAYDIDKTQEYIETHPFSLT

EKGMSIKSKMDKICQTIIGCRNNIIDYAYSFFEKNGYSIIGLEKLTSSQFEKTKSMPTCK

SLLNFHKVLGHTLSELETLPINDVVKKGYYTFTTDNEGKITDASLSEKGKVRKMKDDFFN

QAIKAIHFADVKDYFATLSNNGQTGIFFVPSQFTSQMDSNTHNLYFENAKNGGLKLAPKY

KVRQTQEYHLNGLPADYNAARNIAYIGLDETMRNTFLKKANSNKSLYNQPIYDTGIKKTA

GVFSRMKKLKRYEII

>Cas12l 7YOJ_A Chain A, CasPi [Armatimonadota bacterium]

MAKATKEVKSKRVEALRQVAYQRLERLERKAQKIGAHLRKPGKAADLQSLHYLLHKVEVE

YHDIARNLEKDPTWTPKPKMRREKRAIVPESGPAAPLPTTAKGEPGRPANRHIPPPVPLD

SARIPEDQQSMGQGSGGRSWCSAPFVEVKLPPTQWSNVREKLLKFRIEDDADIVRRWAEA

KFGSIETARDGLRASAEIGTSPDVWRSFISRAISNGKKDFEPLLSLDDDELTADATAERV

VRRWHQIDWVGRMLDSILETVPSGVSKDTFRSRVESRLKTFHSSVNSFELKKRKDGTVER

KRKHTNPQFPYLSPSAVSIDPDVVTMEAVELLQMQPEERFAKDPNDANGRMRLRVLQAEL

GKARREALGRRGEKAPPWSGRKVFRGTTTRKREACLVWDKEAQADGLYFALVMSGGPKID

DKRFVYMDGQPLQSDWQLHNGVAGKAKSCRAMPLILKHDFLRWYHRHIKNHDVNAPLEKR

CVHTTTQFVFVEPDEKKGLQPRLFIRPVFKFYDPVYEVPDSHSIDKKPDCRYLIGIARGV

NYPYRAAVYDCETNSIIADKFVDGRKADWERIRNELAYHQRRRDLLRNSRASSAAIQREI

RAIARIRKRERGLNKVETVESIARLVDWAEENLGKCNYCFVLADLSSNLNLGRNNRVKHI

AAIKEALINQMRKRGYRFKKSGKVDGVREESAWYTSAVAPSGWWAKKEEVDGAWKADKTR

PLARKIGSYYCCEEIDGLHLRGVLKGLGRAKRLVLQSDDPSAPTRRRGFGSELFWDPYCT

ELCGHAFPQGVVLDADFIGAFNIALRPLVREELGKKAKAVDLADRHQTLNPTVALRCGVT

AYEFVEVGGDPRGGLRKILLNPAEAVI

>Cas12l RIJ98969.1 MAG: hypothetical protein DCC46_10025 [Armatimonadota bacterium]

MSPLERSLRKVGENRLERLRVREEKIRKHIEQHPRGKNDHQALHFLLHQIEVERNDLYRN

LKDPEYVPKPAKQRRERRQINVAKPPTRPKKEKGPQPESTKYVIRPPVPGKNLPAFASKY

EARDTRDDSYQDGRSWTSAPYVEVELPILGADKVIQKLMKFVQKDERSIVRDWATKTYSS

IEAAREALLVGAQVSEDVSVWRGLLAETKNAQNFAALSDDQIEAAMSKEAKGADLRPRRA

ALLVAQRHWVDQTVKAIKESAPSGVDKDTLDRRLRAGLRGFHTAANSGKHTNPQFPYLTA

EKPVVPMESVVQSVLAFLDDPDDQRYTKDKEDDKKRHRVTVLQKELGKARPRKRLELQTP

KWAGRPTVKGTISKRRDAALVWDTSKEANGLCLALPIGGMPKIDVEQFIYQDGTSLLSDC

QIASKTTKKGAACAVLPLKPKHDFLRWFTKHVENHNPDAPLERRCLHNTTQFVIVDPEGP

RPRLFVRPVFKFYDPGKTVPNTHETWKKPDCRYLVGIDRGINYVLRAVVVDTEEKKVIAD

IGLPGRKHEWRMIRDEIAYHQQMRDLARNTGKHASVVAKHVRALALARKKDRALGKFATV

EAVAELVKKCEQDYGSGNYCFVLEDLDMGAMNLKRNNRVKHMAVMEEALVNQMRKQGYAY

DGRRGRVDGVRHEGAWYTSQVSPFGWWAKRDEVEEAWKRDKTRPIGRKVGNWYEMPEPGQ

DGDRPDTYRKGYWSKPKNAEGKPYGRNRFSVEPGDEKPDAERRFCWGSELFWDPNVKSFK

GKEFPEGVVLDADFVGALNIALRPLVNDGQGKGFKAEDMAREHTILNPQFKIACQIPVYE

FVEEDGDKWAALRRIML

>Corynebacterium glutamicum Cas12n

MTSTTLAPEEPLMVFRGARFRLDPTGEQQGILSQQAGAARVAYNMMCTLNKDILEARSQL

YSTLIKDGKTKDKAKKELKAAAKEDPSLAIVWARDFDKNYITPERNRHKHAAQRIAAGEN

PVDVWNPDEERFNEPWLHTANRRVLRSGQKQYEQALDNFFKSQNGSRAGQKMGKPRFKTK

IRSTDSFTIDAVDVSSSTTLIRDIGPKDHARYKTGEASTGIIADYRHVRLSHLGTFRVFG

STKALVRQLDRGGRIKSCTVSRSADRWYVSFLVELPIEIARSTPTKKQYKAGAVGIDLGV

KSLAALSTGEIIPNPRFLRTADKKIKKLQRKIARCQKGSKNSIRLKRRLARCHHELALQR

AGYLNELTSMLASSFSAIALEDLNVAGMTSSARGTVENPGKNVKQKAGLNRSILDISPGR

IRTLLEYKCTDRGVELQVIDRFFPSSQLCSSCGSKTTIPLAQRIYHCDVCGGVIDRDVNA

AINIVYEAKRLVEQKCSEHSAPEGAEDKRPWSVHLPSTQYVDGLYTRKRQGPSGHSS

>Nocardiopsis metallicus Cas12n

MKKDAHQRWRDEVDILITTGLSEKDARTRLKGTSKIPNKPDVYKAFQHLRGDAAQGIDGI

APWHAEIPTYVFQSAFQDADRAWKNWLDSYTGKRAGRRVGYPRFKKKFRSRDSFRLHHDA

KKPKPALRLEGYRRLRLGGALKTVRLHGSAKPLHRLVSSGRAVVQSVTVSRGGTRWYASV

LCKVETDVPDPTHRQKANGRIGLDWGLNHLAALSTPLDGHHLVDNPRHLRHASKRLTKAQ

RALSRTQKGSARRRRAAARVGKLHHLVAEQRATFLHTLTKRLTTTFACVAIEDLNVAGMT

RSARGTVQLPGKNVSSKAGLNRSILDAAPAELRRQLEYKTSWYGSHIAILDRWFPSSKTC

SGCGWRNPSLPLSEREFVCAECGLRLDRDLNAARNIAAHAEVPASGTGAPGRGESVNARG

GCVSPRLLRESGQHPSKREDAAPSGPAPPRRSNPPTFP

>Rothia dentocariosa Cas12n

MATTDKKDEENLRAYKFRLDPNQAQTTALYQAVGAARYTYNMLTAYNLEVNRLRDDYWKR

RHDEDISDADIKKELNALAKEDKRYKQLNYGAFGTQYLTPEKKRHEQAEHRIENGEDPSV

VWNQETERSANPWLHTANQRVLVSGLQNASDAWDNFWASRTGKRAGRLVGTPRFKKKGVS

RDSFTVPAPEKMGAYGTAYLRGEPAYKQGRRKITDYRHVRLSYLGTIRTFNSTKPLVKAV

VAGAKIRSYTVSRNADRWYVSFLVKFSEPIRRSATKRARAAGSVGVDLGVKYLASLSDSE

APQRFPNLKFVEGLPSLENPRWSEASSRRLHKLQRALARSQKGSNRRSRLVKQIARLHHM

TALRRESNLHQLTKKLATEYTLVGFEDLNVSGMTASAKGTVENPGKNVAQKSGLNRVVLD

AAFGVFRNQLEYKAVWYGSAFEKVDRYFASSQTCSECGRKAKTKLTLRDRVFDCAYCGNM

MDRDLNAAVNICREAQRLFDEKLASEDRESLNGRGSRGALRGAETVEASRPPASHRRGSP

>Mycobacterium heckeshornense Cas12n

MVRSVGDDLRAYRFALDLRPAQLRAVAEHAGAARWAYNHALGVKFAALKQRQTVIAELVT

AGVDAQAAARRAPRIPTRPQIQKDLNATKGDSRSGVDGLCPWWWTASTYAFQSAMIDADR

AWHNWMSSVTGQRAGRRVGRPRFKSKHRCRDSFRIHHNVKQPTIRPDTSGYRRLIVPRLG

SLRTHDSTKRLRRALDRGAVIQSVTISRGGHRWYASVLVKTPDAPAVGPSRAQRKAGIVG

VDIGVHSLAALSTGELVVNPRHLNSSCDRLGAAQRALARCQKGSNRRRRAARLLGRRHHE

LAERRATALHQLTKRLATCWSVVAVEDLHVAGMIRSARGTVDKPGRNVRAKSALNRAILD

VAPGELRRQLAYKTVWYGSSLVVCDRFYPSTQTCSACGAKAKLTLADRVFRCTACGFGPT

DRDINAARNIAANAAVASGAGETLNARRADQQHLPQVGPCRRPAMKREGHHQSKTDSGHL

SRATG

>Saccharomonospora piscinae Cas12n

MRAYRFTLDPTRTQLDTLAQHAGAARWAYNHALATKLDALRRRQLQIDELVTLGLTEQQA

RRDATVTVPSKPQVQKRWNQLKGDTTRGGDGICPWWRAVSTYAFQSAFLDADRAWKNWMD

SLTGKRAGRRVGAPRFKKKGRCRDSFRLHHAVNDPTIRLEGYRRLRMPRLGSIRLHDSGK

RLARALARGGRVQSVTVSRGGHRWYASVLVDEPDHTPGRETQHRPSRAQHTAGGVGVDVG

VHHLAALSTGETLDNPRHLHHAHTRLVKAQRALARTQKGSHRRRRAAERVGRLHHQLAER

RASHLHTITKRLATQHALVAVEDLNVQGMTRTARGTLTQPGRNVRAKAGLNRAILDAAPG

ELRRQLEYKASWYGSTLAVCDRWAPTSKTCSTCGTVKTKLPLSTRVYRCDTCGMVCDRDI

NAARNILKDADPVAPGRGETLNACGGPVSPSAPHAPTARPSEAGRPGHPARSPRGNDPPS

TPKTE

>Thiopseudomonas Cas12n

MPLNGWTMLAQEEVRAMTTTTGTDLVPRLRAFKHRLDPNPAQATLLAQYAGAARVAYNML

TAHNRAALAASAARRTELAETGLAGPELAARMKAERAADPTLRVASYQSYSTTHLTPLIR

RHREAAAAIAAGADPAEAWTDERYAEPWMHTVPRRVLVSGLQNAAKATENWMASASGTRA

GARVGLPRFKKKGRSRDSFTIPAPEVIGAAGTPYKRGEPRRGVITDHRHLRLASLGTIRT

YDKTSRLVRACRRGAQIRSMTISQAGGRWYASILVADPTPIRTGPSRRQRANDAVGVDLG

VKHLAALSTGEVIDNGRPGARQAARLTRLQRAYARTQPGSNRRERVRRQIAALHHGIALR

RAGLLHQVSTRLAMDFAVVALEDLNVAGMTRSARGTLEAPGRNVAAKSGLNRAILDAGLG

MLRRQLDYKTSWAGSQVKMIDRFAPSSKACSRCGTVKSTLSLAERTFECEACHLVIDRDV

NAAINIRAWAVQEERGAGVGLARGRRESRNGRGAAVSGPPSGGAAGQGRGSVKPAPQGVG

MSSRATGWSSQPPSTEGESAERGASALAR

>ISDra2TnpB

MIRNKAFVVRLYPNAAQTELINRTLGSARFVYNHFLARRIAAYKESGKGLTYGQTSSELT

LLKQAEETSWLSEVDKFALQNSLKNLETAYKNFFRTVKQSGKKVGFPRFRKKRTGESYRT

QFTNNNIQIGEGRLKLPKLGWVKTKGQQDIQGKILNVTVRRIHEGHYEASVLCEVEIPYL

PAAPKFAAGVDVGIKDFAIVTDGVRFKHEQNPKYYRSTLKRLRKAQQTLSRRKKGSARYG

KAKTKLARIHKRIVNKRQDFLHKLTTSLVREYEIIGTEHLKPDNMRKNRRLALSISDAGW

GEFIRQLEYKAAWYGRLVSKVSPYFPSSQLCHDCGFKNPEVKNLAVRTWTCPNCGETHDR

DENAALNIRREALVAAGISDTLNAHGGYVRPASAGNGLRSENHATLVV

>WP_077795519.1 RNAguided endonuclease TnpB family protein [Shigella sonnei]

MKRLQAFKFQLRPGGQQEREMRRFAGACRFVFNRALARQNENHEAGNKYIPYGKMASWLV

EWKNATETQWLKDAPSQPLQQSLKDLERAYKNFFQNRAAFPRFKKRGQNDVFRYPQGVKL

DQENSRIFLPKLGWMRYRNSRQVTGVVKNVTVSQSCGKWYISIQTESEVSTPVHPSASMV

GLDAGVAKLATLSDGTVFEPVNSFQKNQKKLARLQRQLSRKVKFSNNWQKQKRKIQRLHS

CIANIRRDYLHKVTTTVSKNHAMIVIEDLKVSNMSKSAAGTVSQPGRNVRAKSGLNRSIL

DQGWYEMRRQLEYKQLWRGGQVLAVPPAYTSQRCACCGHTAKENRLSQSKFRCQVCGYTV

NADVNGARNILAAGHAVLACGEMVQSGRPLKQEPTEMIQATAQRQLSRKVKFSNNWQKQK

RKIQRLHSCIANIRRDYLHKVTTAVSKNHAMIVIEDLKVSNMSKSAAGTVSQPGRNVRAK

SGLNRSILDQGWYEMHRQLEYKQLWRGGQVLAVPPAYTSQRCAYCGHTAKENRLSQSKFR

CQVCGYTANADVNGARNILAAGHAVLACGEMVQSGRPLKQEPTEMIQATA

>WP_052982496.1 RNAguided endonuclease TnpB family protein, partial [Shigella sonnei]

GNKYIPYGKMASWLVEWKNATETQWLKDAPSQPLQQSLKDLERAYKNFFQNRAAFPRFKK

RGQNDVFRYPQGVKLDQENSRIFLPKLGWMRYRNSRQVTGVVKNVTVSQSCGKWYISIQT

ESEVSTPVHPSASMVGLDAGVAKLATLSDGTVFEPVNSFQKNQKKLARLQRQLSRKVKFS

NNWQKQKRKIQRLHSCIANIRRDYLHKVTTTVSKNHAMIVIEDLKVSNMSKSAAGTVSQP

GRNVRAKSGLNRSILDQGWYEMRRQLEYKQLWRGGQVLAVPPAYTSQRCACCGHTAKENR

LSQSKFRCQVCGYTVNADVNGARNILAAGHAVLACGEMVQSSRKVKFSNNWQKQKRKIQR

LHSCIANIRRDYLHKVTTAVSKNHAMIVIEDLKVSNMSKSAAGTVSQPGRNVRAKSGLNR

SILDQGWYEMHRQLEYKQLWRGGQVLAVPPAYTSQRCAYCGHTAKENRLSQSKFRCQVCG

YTANADVNGARNILAAGHAVLACGEMVQSGRPLKQEPTEMIQATA

>WP_192936035.1 RNAguided endonuclease TnpB family protein [Flavonifractor plautii]

MEKAYKFRLYPTAKQEELIRKTIGCSRFVYNQTLAARKGAYASGSSIHMTFKESHSGYDC

VKLLPGLKDTYPWLREVDSTALQASVLNMDHAYKNFFSGRKGRRKVGFPKFKAKHHSKAS

YTSKVVGHNIQASDRAVKLPKLGWVKAKVSTTVQGRILNATVSMSRSGKFFVSLCCTEVD

IPQLLSTGAAVGLDVGIKDLVITSDGQKYDNPKYLQKAEKKLATLQRRLSRKPKGSCNRE

KARLRLARQHEYIANCRQDYXVSMSRSGKFFVSLCCTEVDIPQLLSTGAAVGLDVGIKDL

VITSDGQKFDNPKYLRKAEKKLATLQRRLSRKPKGSCNREKARLRLSRQHEYIANCRQDY

LHKLTTQLVRDYDIICVESINVGGMLKNHKLAKAIADASWGELARQLKYKAAWQHKMLVE

VGTFFPSSQLCSCCGYQNKEVKDLSVREWTCPQCGAHHDRDQNAAQNLLIEGLRLLPHDF

QRKSQASA

>WP_150190461.1 RNAguided endonuclease TnpB family protein [Bacillus cereus]

MTKQNKAYKFRLYPTEDQAHLMRKTFGCVRFVYNRMLAERKEAYEKHKDDKDQLKKQKLP

TPAKYKAEFEWLKEVDSLALANAQLNLQTAYKNFFRGQNDFPTFKSKKDRKSYTTNVVNG

NIMLLNGHIKLPKLKMVRIKQHREIPQDHIIKSCTIFMTPTGKYYVSILTEYEKEIVQKK

VETVVGLDFAMDQLYVSSEDERANYPKFYREMLDRLAKAQRVLSRRTKGSGRWNKQCIRV

AKLHEKVANQRKNFLHHKSKELATHFDVVAIEDLNMKGISQALHFGKSIADNAWGMFTSF

LAYKLNEQGKQLVKIDKWFPSTKTCSSCGXNFLHHKSKELATHFDVVAIEDLNMKGISQA

LHFGKSIADNAWGMFTSFLAYKLNEQGKQLVKIDKWFPSTKTCSSCGNVKNMSLSERVYS

CICGVNLDRDYNAAINIKNEAIRLLALA

>WP_023070664.1 RNAguided endonuclease TnpB family protein [Leptolyngbya sp. Heron Island J]

MSQKAFRYRFYPTPEQENLLRRTMGCARLVYNRALAARSDAWYKHQKRVGYKETSAMLTL

WKKEDSLNFLNEVSCVPLQQCLRHLQKAFTNFFAGRAKYPNFKKKHNGGSAEFTKSAFKF

RDNQIYLAKCSEPLPIRWSRPLPEGAKPSTVTVKLDPSGRWFVSLLVDVGIETLPKSPNK

IGVDLGVTSLIADSNGGKVANPKTFRKKHKRLALAQKRLAKKQKGSNNRHKARLKVARIQ

AEIADTRRDFLHKLTTRLIRENQVIAVEDLAVKNMKKNHKLALSISDASWGELVRQLEYK

ADWYGRTVVKIDRWFPSSKRCSNCGHISPKMLLNVRDWVCPECGHSHDRDINAAINILAA

GLAVEVCGATVRPEQSKSVKASATKQKTLKREPGNPPL

>Cas12m 8PM4_A Chain A_Transposase [Gordonia otitidis NBRC 100426]

SNAGGGGMTRVTVQTAGVHYKWQMPDQLTQQLRLAHDLREDLVTLEYEYEDAVKAVWSSY

PAVAALEAQVAELDERASELASTVKEEKSRQRTKRPSHPAVAQLAETRAQLKAAKASRRE

AIASVRDEATERLRTISDERYAAQKQLYRDYCTDGLLYWATFNAVLDHHKTAVKRIAAHR

KQGRAAQLRHHRWDGTGTISVQLQRQATDPARTPAIIADADTGKWRSSLIVPWVNPDVWD

TMDRASRRKAGRVVIRMRCGSSRNPDGTKTSEWIDVPVQQHRMLPADADITAAQLTVRRE

GADLRATIGITAKIPDQGEVDEGPTIAVHLGWRSSDHGTVVATWRSTEPLDIPETLRGVI

TTQSAERTVGSIVVPHRIEQRVHHHATVASHRDLAVDSIRDTLVAWLTEHGPQPHPYDGD

PITAASVQRWKAPRRFAWLALQWRDTPPPEGADIAETLEAWRRADKKLWLESEHGRGRAL

RHRTDLHRQVAAYFAGVAGRIVVDDSDIAQIAGTAKHSELLTDVDRQIARRRAIAAPGML

RAAIVAAATRDEVPTTTVSHTGLSRVHAACGHENPADDRYLMQPVLCDGCGRTYDTDLSA

TILMLQRASAATSN

>H5TRP0.1 RecName: Full=CRISPRassociated DNAbinding protein Cas12m; Short=GoCas12m [Gordonia otitidis NBRC 100426]

MTRVTVQTAGVHYKWQMPDQLTQQLRLAHDLREDLVTLEYEYEDAVKAVWSSYPAVAALE

AQVAELDERASELASTVKEEKSRQRTKRPSHPAVAQLAETRAQLKAAKASRREAIASVRD

EATERLRTISDERYAAQKQLYRDYCTDGLLYWATFNAVLDHHKTAVKRIAAHRKQGRAAQ

LRHHRWDGTGTISVQLQRQATDPARTPAIIADADTGKWRSSLIVPWVNPDVWDTMDRASR

RKAGRVVIRMRCGSSRNPDGTKTSEWIDVPVQQHRMLPADADITAAQLTVRREGADLRAT

IGITAKIPDQGEVDEGPTIAVHLGWRSSDHGTVVATWRSTEPLDIPETLRGVITTQSAER

TVGSIVVPHRIEQRVHHHATVASHRDLAVDSIRDTLVAWLTEHGPQPHPYDGDPITAASV

QRWKAPRRFAWLALQWRDTPPPEGADIAETLEAWRRADKKLWLESEHGRGRALRHRTDLH

RQVAAYFAGVAGRIVVDDSDIAQIAGTAKHSELLTDVDRQIARRRAIAAPGMLRAAIVAA

ATRDEVPTTTVSHTGLSRVHAACGHENPADDRYLMQPVLCDGCGRTYDTDLSATILMLQR

ASAATSN

>Cas12m VBA31969.1 hypothetical protein LAUMK4_05640 [Mycobacterium persicum]

MAVTVQTMGVHYRWPLPDVLRAQLRLAHDLREDLVTLQLEYEETLKAIWSSYPLVAAAEE

TLQFAEAAAETAAQAVSAERLRQRTKRIVGPIADRLTAARVEVKRARQQRRDAIVAVRDH

AADRIHQAATQLKTDHKRLYAQYCQQKGLYWATYNDVLNQHNTAVKLIKRARAAGRKSQL

RHHRYDGTGSIAVQLRRQSHQPARTPIVLADPAGKYRNALVLPWIEPTRWQAMSRAQQRH

RGRVTVRMRCGSQNGQPTWIEIPVQQHRMLDADADIIGARLTVTRTAGHLTARLSVTAKL

PDPPEVDSGPVVAIHLGWRDTDNGTQVATWRSTSPLDIPMELRDQLIASPDAPTGAVVVP

HRITERITHSDELRSQRDLALDAVRAKLAAWLAEHGPVPHPTRPEATVEGGDVARWRSPA

RFAALAQAWRVAPPPGGQDIAGVLESWRRSDRALWERQEHGRGKAVRHRNNLYRQIAAIF

ADQAGRIVVDDTSVSAIAAAPTDLPTEVATRIARRRVVAAPANLRAAITSAATREGVPVS

VVPAAGLTRIHAHCGYQNPADGRHIARPVLCDGCGSSYDPDASATLLMLQRVNAYPAPAT

RTK

>Cas12m WP_122526562.1 hypothetical protein [Mycobacterium persicum]

MGVHYRWPLPDVLRAQLRLAHDLREDLVTLQLEYEETLKAIWSSYPLVAAAEETLQFAEA

AAETAAQAVSAERLRQRTKRIVGPIADRLTAARVEVKRARQQRRDAIVAVRDHAADRIHQ

AATQLKTDHKRLYAQYCQQKGLYWATYNDVLNQHNTAVKLIKRARAAGRKSQLRHHRYDG

TGSIAVQLRRQSHQPARTPIVLADPAGKYRNALVLPWIEPTRWQAMSRAQQRHRGRVTVR

MRCGSQNGQPTWIEIPVQQHRMLDADADIIGARLTVTRTAGHLTARLSVTAKLPDPPEVD

SGPVVAIHLGWRDTDNGTQVATWRSTSPLDIPMELRDQLIASPDAPTGAVVVPHRITERI

THSDELRSQRDLALDAVRAKLAAWLAEHGPVPHPTRPEATVEGGDVARWRSPARFAALAQ

AWRVAPPPGGQDIAGVLESWRRSDRALWERQEHGRGKAVRHRNNLYRQIAAIFADQAGRI

VVDDTSVSAIAAAPTDLPTEVATRIARRRVVAAPANLRAAITSAATREGVPVSVVPAAGL

TRIHAHCGYQNPADGRHIARPVLCDGCGSSYDPDASATLLMLQRVNAYPAPATRTK

>8HHL_A Chain A, Cas12m2 [Mycolicibacterium mucogenicum]

GGMTTMTVHTMGVHYKWQIPEVLRQQLWLAHNLREDLVSLQLAYDDDLKAIWSSYPDVAQ

AEDTMAAAEADAVALSERVKQARIEARSKKISTELTQQLRDAKKRLKDARQARRDAIAVV

KDDAAERRKARSDQLAADQKALYGQYCRDGDLYWASFNTVLDHHKTAVKRIAAQRASGKP

ATLRHHRFDGSGTIAVQLQRQAGAPPRTPMVLADEAGKYRNVLHIPGWTDPDVWEQMTRS

QCRQSGRVTVRMRCGSTDGQPQWIDLPVQVHRWLPADADITGAELVVTRVAGIYRAKLCV

TARIGDTEPVTSGPTVALHLGWRSTEEGTAVATWRSDAPLDIPFGLRTVMRVDAAGTSGI

IVVPATIERRLTRTENIASSRSLALDALRDKVVGWLSDNDAPTYRDAPLEAATVKQWKSP

QRFASLAHAWKDNGTEISDILWAWFSLDRKQWAQQENGRRKALGHRDDLYRQIAAVISDQ

AGHVLVDDTSVAELSARAMERTELPTEVQQKIDRRRDHAAPGGLRASVVAAMTRDGVPVT

IVAAADFTRTHSRCGHVNPADDRYLSNPVRCDGCGAMYDQDRSFVTLMLRAATAPSNP

>Cas12c_0_sd2019_length_1145

MTKLRHRQKKREVLGSNGKVTEFRKAFSAYARATKGNMFTHSFPFKTKPSLHQCELADKA

YQSLHSYLPGSLAHFLLSAHALGFRIFSKSGEATAFQASSKIEAYESKLASELALSIQNL

TISTLFNALTTSVRGKGEETSADPLIARFYTLLTGKPLSDGPERDLAEVISRKIASSFGT

WKEMTANPLQSLQFFEEELHNVSLSPAFDVLIKMNDLQGDLKNRTIVFDPDAPVFAEDPA

DIIIKLTARYAKEAVIKNQNVGNYVKNAITTTNANGLGWLLNKGLSLLPVSTDDELLEFI

GVERSHPSCHALIELIAQLEAPELFEKNVFSDTRSEVQGMIDSAVSNHIARLSSRNSLSM

DSEELERLIKSFQIHTPHCSLFIGAQSLSQQLESLPEALQSGVNSADILTYQRTLNRINY

LSGVAGQINGAIKRKAIDGEKIHLPAAWSELISLPFIGQPVIDVESNEFDTLIKNFDLNF

NKALLNRTQHFEAMCRSTKKNALSKPEIVSYRDLLARLTSCLGSLVLRRAGIEVLKKHKI

FESNSELREHVHERKHFVFVSPLDRKAKKLLRLTPDLLHVIDEILQHDNLENKDRESLWL

VRSGYLLAGLPDQLSSSFINLPIITQKGDRRLIDLIQYDQINRDAFVMVTSAFKSNLSGL

QYRANKQSFVVTRTLSPYLGSKLVYVPKDKDWLVPSQMFEGRFADILQSVWKDAGRLCVI

DTAKHLSNIKKSVFSSEEVLAFLRELPHRTFIQTEVRGLGVNVDGIAFNNGDIPSLKTFS

NCVQVKVSRTNTSLVQTLNWFEGGKVSPPSIQFERAYYDQIHEDAKRKIRFQMPATELVH

ASDDAGWTPSYLLGIDPGEYGMGLSLVSINNGEVLDSGFIHINSLINFASKKSNHQTKVV

PRQQYKSPYANYLDSAAGDIAHILDRLIYKLNALPVFEALSGNSQSAADQVWTKVLSFYT

WGDNDAQNSIRKQHWFGASHWDIKGMLRQPPTEKKPKPYIAFPGSQVSSYGNSQRCSCCG

RNPIEQLREMAKDTSIKELKIRNSEIQLFDGTIKLFNPDPSTVIERRRHLGPSRIPVADR

LEFKELITIVSRSIRHSPEFIAKKRGIGSEYFCAYSDCNSSLNSEANAAANVAQKFQKQL

FFEL*

>Cas12c_1_sd2019_length_1251

MNARDWRKHVGVLAQQHKETTRTYTFPLDTTGSAIDFDAALQAYNAVEGVGYGSLLGLAC

AVHLSGFRLFSTGKEAATFRNRARYPNAAFQAALRKELGTTITTLTPETLDRLFSSRPKR

RNGVPLPWNQDSIRDRLYTNWVKPRPGDTPDAVLFQIATGIAQEITEDVSSWTDLAKNSD

RGLKAAHRYFARVGGFPAFDNLTPPATVQPTDTTIDYDPNAPFHLVSHADQTLIHQSISL

CAHRIRQEDPALDPNKSGFIKQLQNNFLSQTFYGLSWLFGAGYVHFRECTANDLAIQYGI

PNNCRDGIHQIKSFADAILPNTFFEKKHYRKDSRSVGKKADQVHRDNLHQLKNRLPLDLR

RPQALNKISGGVPDVAKSIRGLETQLDQVLKERRSHFGRLTKWAKECGITLDPLQPLIES

EKQRVAERGSAHDAKELAIRLLLQRIGRLGHRLSPTNATAIQEKSWISNYWQRLLQLQTW

VDHTWVTLPQELTEAQFKPLFRGLLVDAVELMAIAERLPQRLADCRDSLDCLMGKGPQAA

TKNDVEIVEKVREEIESFVGQIEQLGNQLRHQLENENNLLRPVFAVKREFNLFFHNHMGA

LYRSPYSTSRHQPFQINVDVAHGTDWIGTIETLIQNLFTQIQDDALLRDLVQLEGFVFSH

KLRALPGVIPSELARPNNLQQMGLPALLLVLLQADQVHRETVLRVFNLYGSAINGYLFQA

LRPGFIVRAGFQRLETKKLRYVPKAQSWQYPDRLHHAKSAIKNSLSAGWIKKNHQGAILP

QKTLTALVKQKSLKDTGVPEYLVQAPHDWYVPIDLRGPAIPIEGLTVGTEGPELTQLGPM

KDDCAFRAIGPSSFKSKIDAGLLPQDVKYGDMTLIFDQHYQQSISFNGTFSIQYQPTSLQ

VKAAIPVVDKRPRDTRNNSHLYDRIVAIDLGERKIGYAIFDLKQVLKSEQLEPMREDGKP

LIGSISIRSIRGLMKAVQTHRNRRQPNYRIDQTYSKALMHYRESVIGDVCNAIDTLCARY

GGFPVLESSVRNFEVGSAQLKTVYGSVSRRYTWSAVDAHKNQRQQYWLGGTIWTHPYLMT

REWDEKNSKWSNRSKPLKMHPGVEVHPAGTSQICHQCKRNPIGALWNVADTVVLDDQGQL

DLDDGTIRLNSGYIDTTEIKRARRKKIRLPENKPLTGSHKTSHVRAVARRNLRQPPKSTR

AKDTTQSRYTCLYVDCGHECHADENAAINIGRKYLQERIHIEASRQALST*

>Cas12c_4_sd2019_length_1242

MKKFELKQNFRNNYSGKTLRNFRQTLAQIANKKSSDSILTIKFKLDCSKTGKLPKYENLI

SLYDTIEDIKKGTLSYYLFTLIVSGFKFFGSASQAKAFSTKDIFKDNDFYNQFKIQSHLD

LPDFVPSKIYQRLKKNVRSTNGKDNAFKASVIVAEYRKEIGKLSSEHQCEELFKKIGTAL

ETRFSSWQDLINNCSTGCEIIDEILDSFGTLPSIKKMVLASTTQSSDDGIAIAYDPDSTF

IKSDELLNPYFAVATILKSMPPEIQQDKKSAYVKANLTTPTHNALSWIFGKGLTLFQEST

EKLCAMFNVSDKVQDAAKAVKLPAELDLNHCTLKFQDFRSSLGGHLSELKKKIEKNEKWD

IWKNNLKKIPKLNKLSGGVPDAWKEIREIEQKFHEISENQKKHFTEVMEWIDAGNGTIDI

FESRFKYDELLKKSKKNNLQSADELAFRSVLNKLGRFARQGNDLVCEKIKNDSWTTNYLK

RLDELNDLLLNLPKNLSLPDIFMIDGKDFIEYSGCNRDEIQQMIDFVVNEQNRIKLQESL

NALLGKGNNQICSDDISTVKDFSEIVLHSFVQQIDLEQSSNEANSIFWFKEQNIFDSSKD

FNRYFINQKGFIFKHPSSKKDNSPYNLSANLLEKRYEVTNTVGALLEQCESDPVNDPFSM

RSLVEFRALWFSINISGISKEIPTKIAQPKLDDSTYQESVSPTLKYRLEKEQITSSELNS

IFTVYKSLLSGLSIRLSRNSFYLRTKFSWIGNNSLIYCPKETTWKIPAAYFKSDLWNEYK

DKQILIVNEEYDVDVVKTFESVYKIVKSKKNRILPLLKQLPHDWMFKLPFGASNAEKCKV

LKLEKNNKKFKPLSVSKDSLARLSGPSTYFNQIDEIMMNDESELSEMTLLADEPVRQQMS

NGKIEIIPDDYVMSLAIPITRSLKKGNTESFPFKNIVSIDQGEAGFAYAVFKLSDCGNER

AEPIATGLIPIPSIRRLIHSVKKYRGKKQRIQNFNQKFDSTMFTLRENVTGDICGLIVAL

MKKYNAFPILEKQVGNLESGSKQLMLVYKAVNSKFLAAKVDMQNDQRRSWWYQGNSWNTP

ILRISNPNQSNNKNVKNINGKKYEELKIYPGYSVSAYMTSCICHVCGRNALELLKNDDST

GKVKKYQINQDGEVTIGGEVIKLYRKPDRLTPVKNLARERTYASINERAPMSKDTTQSRY

FCVFKNCPCHNKEQHADVNAAINIGRRFLKDCILDDNKEKD*

>Cas12c_6_sd2019_length_1228

TKLRHRQKKLTHDWAGSKKRLGSNGKLQNPLLMPVKKGQVTEFRKAFSAYARATKGTDGR

KNMFTHSFPFKTKPSLHQCELADKAYQSLHSYLPGSLAHFLLSAHALGFRIFSKSGEATA

FQASSKIEAYESKLASELALSIQNLTISTLFNALTTSVRGKGEETSADPLIARFYTLLTG

KPLSRDTQGPERDLAEVISRKIASSFGTWKEMTANPLQSLQFFEEELLDANVSLSPAFDV

LIKMNDLQGDLKNRTIVFDPDAPFEYNAEDPADIIIKLTARYAKEAVIKNQNVGNYVKNA

ITTTNANGLGWLLNKGLSLLPVSTDDELLEFIGVERSHPSCHALIELIAQLEAPELFEKN

VFSDTRSEVQGMIDSAVSNHIARLSSSRNSLSMDSEELERLIKSFQIHTPHCSLFIGAQS

LSQQLESLPEALQSGVNSADILLGSTTNSLVEESIATYQRTLNRINYLSGVAGQINGAIK

RKAIDGEKIHLPAAWSELISLPFIGQPVIDVESDLAHLKNQYQTLSNEFDTLISALQKNF

DLNFNKALLNRTQHFEAMCRSTKKNALSKPEIVSYRDLLARLTSCLYRGSLVLRRAGIEV

LKKHKIFESNSELREHVHERKHFVFVSPLDRKAKKLLRLTDSRPDLLHVIDEILQHDNLE

NKDRESLWLVRSGYLLAGLPDQLSSSFINLPIITQRLIDLIQYDQINRDAFVMLVTSFKS

NLSGLQYRANKQSFVVTRTLSPYLGSKLVYVPKDKDWLVPSQMFEGRFADILQSDYMVWK

DAGRLCVIDTAKHLSNIKKSVFSSEEVLAFLRELPHRTFIQTEVRGLGVNVDGIAFNNGD

IPSLKTFSNCVQVKVSRTNTSLVQTLNRWFEGGKVSPPSIQFERAYYKKDDQIHEDAAKR

KIRFQMPATELVHASDDAGWTPSYLLGIDPGEYGMGLSLVSINNGEVLDSGFIHINSLIN

FASKKSNHQTKVVPRQQYKSPYANYLEQSKDSAAGDIAHILDRLIYKLNALPVFEALSGN

SQSAADQVWTKVLSFYTWGDNDAQNSIRKQHWFGASHWDIKGMLRQPPTEKKPKPYIAFP

GSQVSSYGNSQRCSCCGRNPIEQLREMAKDTSIKELKIRNSEIQLFDGTIKLFNPDPSTV

IERRRHNLGPSRIPVAKNISPSSLEFKELITIVSRSIRHSPEFIAKKRGIGSEYFCAYSD

CNSSLNSEANAAANVAQKFQKQLFFEL*

>Cas12c_8_sd2019_length_1251

MAHKHKEQLDFVSGQKARQFRKALSGSASDPDKRETVRTLAFPIDISPLGQRAFNEAAQL

GNIVDGVASGTLYGLLTTLLCTGFGVYSSAAERKKAEISSRRPDDPFQEQLKASTGLLIQ

GMSPLELFTEVSTRRRSNNKSDLRGLIDRWIKDKDDGRSVEFVVEMQSYLVAAMQTGGMT

VFNLPERIDTIGRLVDSFLNAKGFSLPSFETQIPDLTSESENFANSTVAFDPSAPRQAGL

DPLLRMHFIVARFWGEARRHGNGEKQLDKYIQSRVTTANGNGFSWLFNEGLGYLRDTSEE

TLIADFKMAAEAISIVQNVQSVARSIPGTALFSTYGPLHNSRSRVQGKLDSWISNYVSRL

GELVDSVSKVGKLIFPEDLEGDAYRRFFLRSGVNVDALKVYGSELPMQAEKIRGSLEILQ

GAIPFDSTAVESVDRFNRFLGELAGSLSAINKRIEDAREAEEADVPRDLPRHDWLKAGKK

LNSISGGTPEYRLELCRLAESFRRLLAARADHFERVRRWASEQSIPFDNAFVAQEEYERR

QETEGREAAERRGSVAERARRQVLERLSRVCRNGSPRLRRIAMGTYLEDGVCSRTILNRY

FEAIQGGFYVAYRSRSRHDPYVVGSGFLRTFDPLAFIGKLQGRLRSEGAGDYNSDLLSLE

NLAIALRLGNVPGPVPGDLNRCVELASIAFRLPYWANDPTRSITSLQLQYLFNMYHSNLS

GLATQLTRPGFRIAVSLSPYKDQARIQYSPKQGSWMPPRHYLHAGGSLEAPLKELGVPTE

PGVEKTARAIVSTLEKNPSSLERRKLATILSQLPHDWVLHADLSGLGESRWTVKVQKGVI

KSGASREGCIRLGGPPSYKTALDRALLDGEISPPTIVIEREVRQTLDEGKVIFKDVSVRG

FVNIPLAEPASTPPLLPRLRNVLGVDLGETGIGFAVIEGTQLPNYEYSRAKLISSAFLPI

RSLRRLINEVNVYRRRTQPRQRYRDAFASPLKALREAAIADTVAAIDGLCAKYEAIPILE

SSVGAFETGSQRLKVIYESVLKYYRADNVDAHQAVRQHHFMGAWKWVHPFFRRTEVRASH

TRDQLQLRPLALFPGAVIHPAGTSRICSACKRNSFQALGRYLTSHSEGAVPITGKGCLSL

DDGVVQFERPASVDPIERARLNRARLRRSFAPIDAGLYPRGQLFSLLAQSVRRPNPSRTA

KDTTQSIFVCPYVDCAASRNADENAAENIARRWLERQGENQRAFDQWDSL*

>Cas12c_10_sd2019_length_1271

MQTKKTHLHLISAKASRKYRRTIACLSDTAKKDLERRKQSGAADPAQELSCLKTIKFKLE

VPEGSKLPSFDRISQIYNALETIEKGSLSYLLFALILSGFRIFPNSSAAKTFASSSCYKN

DQFASQIKEIFGEMVKNFIPSELESILKKGRRKNNKDWTEENIKRVLNSEFGRKNSEGSS

ALFDSFLSKFSQELFRKFDSWNEVNKKYLEAAELLDSMLASYGPFDSVCKMIGDSDSRNS

LPDKSTIAFTNNAETVDIESSVMPYMAIAALLREYRQSKSKAAPVAYVQSHLTTTNGNGL

SWFFKFGLDLIRKAPVKSLQELFSVPDDKLDGLKFIKEACEALPEASLLGELLGYQDFRT

SFAGHIDSWVANYVNRLFELIELVNQLPESIKLPSILTQKNHNLVASLGLQEAEVSHSLE

LFEGLVKNVRQTLKKLAGIDSSSPNEQDIKEFYAFSDVLNRLGSIRNQIENAVQTAKKDK

IDLESAIEWKEWKKLKKLPKLNGLGGGVPKQQELLDKALESVKQIRHYQRIDFERVIQWA

VNEHCLETVPKFLVDAEKKKINKESSTDFAAKENAVRFLLEGIGAAARGKTDSVSKAAYN

WFVVNNFLAKKDLNRYFINCQGCIYKPPYSKRRSLAFALRSDDTIEVVWFETFYKEISKE

IEKFNIFSQEFQTFLHLENLRMKLLLRRIQKPIPAEIAFFSLPQEYYDSLPPNVAFLALN

QEITPSEYITQFNLYSSFLNGNLILLRRSRSYLRAKFSWVGNSKLIYAAKEARLWKIPNA

YWKSDEWKMILDSNVLVFDKAGNVLPAPTLKKVCEREGDLRLFYPLLRQLPHDWCYRNPF

VKSVGREKNVIEVNKEGEPKVASALPGSLFRLIGPAPFKSLLDDCFFNPDKDLRECMLIV

DQEISQKVEAQKVEASLESCTYSIAVPIRYHLEEPKVSNQFENVLAIDQGEAGLAYAVFS

LKSIEAETKPIAVGTIRIPSIRRLIHSVSTYRKKKQRLQNFKQNYDSTAFIMRENVTGDV

CAKIVGLMKEFNAFPVLEYDVKNLESGSRQLSAVYKAVNSHFLYFKEPGRDALRKQLWYG

GDSWTIDGIVTRERKEDGKEGVEKIVPLKVFPGRSVSARFTSKTCSCCGRNVFDWLFTEK

KAKTNKKFNVNSKGELTTADGVIQLFEADRSKGPKFYARRKERTPLTKPIAKGSYSLEEI

ERRVRTNLRRAPKSKQSRDTSQSQYFCVYKDCALHFSGMQADENAAINIGRRFLTALRKN

RRSDFPSNVK*

>Cas12c_12_sd2019_length_1215

MTKHSIPLHAFRNSGADARKWKGRIALLAKRGKETMRTLQFPLEMSEEAAAINTTPFAVA

YNAIEGTGKGTLFDYWAKLHLAGFRFFPSGGAATIFRQQAVFEDASWNAAFCQQSGKDWP

WLVPSKLYERFTKSPREVTKKDGSKKSFTQENVANECHVSLVGASITDKTPEDQKEFFLK

MAGALAEKFDSWKSANEDRIVAMKVIDEFLKSEGLHLPSLENIAVKCSVETKPDNATVAW

HDAPMSGVQNLAIGVFATCASRIDNIYDLNGGKLSKLIQESATTPNVTALSWLFGKGLEY

FRTTDIDTIMQDFNIPASAKESIKPVVESAQAIPTMTVLGKKNYAPFRPNFGGKIDSWIA

NYASRLMLLNDILEQIEPGFELPQALLDNETLMSGIDMTGDELKELIEAVYTWVDAAKQG

LATLLGRGGNVDDAVQTFEQFSAMMDTLNGTLNTISARYVRAVEMAGKDEARLEKLIECK

FDIPKWCKSVPKLVGISGGPPKVEEEIKVMNAAFMDVRARMFVRFEEIAAYVASKGAGMD

VYDALEKRELEQIKKLKSAVPERAHIQAYRAVLHRIGRAVQNCSEKTKQLFSSKVIEMGV

FKNPSHLNNFIFNQKGAIYRSPFDRSRHAPYQLHADKLLKNDWLELLAEISATLMASEST

EQMEDALRLERTRLQLQLSGLPDEYPASLAKPDIEVEIQTALKMQLAKDTVTSDVLQRAF

NLYSSVLSGLTFKLLRRSFSLKMRFSVADTTQLIYVPKVCDWAIPKQYLQAEGEIGIAAR

VVTESSPAKMVTEVEMKEPKALGHFMQQAPHDWYFDASLGGTQVAGRIVEKGKEVGKERK

LVGYRMRGNSAYKTVLDKSLVGNTELSQCSMIIEIPYTQTVDADFRAQVQAGLPKVSINL

PVKETITASNKDEQMLFDRFVAIDLGERGLGYAVFDAKTLELQESGHRPIKAITNLLNRT

HHYEQRPNQRQKFQAKFNVNLSELRENTVGDVCHQINRICAYYNAFPVLEYMVPDRLDKQ

LKSVYESVTNRYIWSSTDAHKSARVQFWLGGETWEHPYLKSAKDKKPLVLSPGRGASGKG

TSQTCSCCGRNPFDLIKDMKPRAKIAVVDGKAKLENSELKLFERNLESKDDMLARRHRNE

RAGMEQPLTPGNYTVDEIKALLRANLRRAPKNRRTKDTTVSEYHCVFSDCGKTMHADENA

AVNIGGKFIADIEK*

>Cas12d_7_sd2019_length_1186

MQKVRKTVHKNPYGTKVRNAKTGYSLQIERLSYTGKEGMRSFKIPLENKNKEVFDEFVKK

IRNDYISQVGLLNLSDWYEHYQEKQEHYSLADFWLDSLRAGVIFAHKETEIKNLISKIRG

DKSIVDKFNASIKKKHADLYALVDIKALYDFLTSDARRGLKTEEEFFNSKRNTLFPKFRK

KDNKAVDLWVKKFIGLNKDKLNFTKKFIGFDPNPQIKYDHTFFFHQDINFDLERITTPKE

LISTYKKFLGKNKDLYGSDETTEDQLKMVLGFHNNHGAFSKYFNASLEAFRGRDNSLVEQ

IINNSPYWNSHRKELEKRIIFLQVQSKKIKETELGKPHEYLASFGGKFESWVSNYLRQEE

EVKRQLFGYEENKKGQKKFIVGNKQELDKIIRGTDEYEIKAISKETIGLTQKCLKLLEQL

KDSVDDYTLSLYRQLIVELRIRLNVEFQETYPELIGKSEKDKEKDAKNKRADKRYPQIFK

DIKLIPNFLGETKQMVYKKFIRSADILYEGINFIDQIDKQITQLLPCFKNDKERIEFTEK

QFETLRRKYYLMNSSRFHHVIEGIIKKRENSELKTFSDKKGKKYENEVYYTFYINPKARD

QRRIKIVLDINGNNSVGILQDLVQKLKPKWDDIIKKNDMGELIDAIEIEKVRLGILIALY

CEHKFKIKKELLSLDLFASAYQYLELEDDPEELSGTNLGRFLQSLVCSEIKGAINKISRT

EYIERYTVQPMNTEKNYPLLINKEGKATWHIAAKDDLSKKKGGGTVAMNQKIGKNFFGKQ

DYKTVFMLQDKRFDLLTSKYHLQFLSKTLDTGGGWWKNKNIDLNLSSYSFIFEQKVKVEW

DLTNLDHPIKPSENSDDRRLFVSIPFVIKPKQTKRKDLQTRVNYMGIDIGEYGLAWTIIN

IDLKNKKINKISKQGFIYEPLTHKVRDYVATIKDNQVRGTFGMPDTKLARLRENAITSLR

NQVHDIAMRYDAKPVYEFEISNFETGSNKVKVIYDSVKRADIGRGQNNTEADNTEVNLVW

GKTSKQFGSQIGAYATSYICSFCGYSPYYEFENSKSGDEEGARDNLYQMKKLSRPSLEDF

LQGNPVYKTFRDFDKYKNDQRLQKTGDKDGEWKTHRGNTAIYACQKCRHISDADIQASYW

IALKQVVRDFYKDKEMDGDLIQGDNKDKRKVNELNRLVPIINKNL*

>Cas12d_8_sd2019_length_1128

MQKVRKTVHKNPYGTKVRNAKTGYSLQIERLSYTGKEGMRSFKIPLENKNKEVFDEFVKK

IRNDYISQVGLLNLSDWYEHYQEKQEHYSLADFWLDSLRAGVIFAHKETEIKNLISKIRG

DKSIVDKFNASIKKKHADLYALVDIKALYDFLTSDARRGLKTEEEFFNSKRNTLFPKFRK

KDNKAVDLWVKKFIGLNKDKLNFTKKFIGFDPNPQIKYDHTFFFHQDINFDLERITTPKE

LISTYKKFLGKNKDLYGSDETTEDQLKMVLGFHNNHGAFSKYFNASLEAFRGRDNSLVEQ

IINNSPYWNSHRKELEKRIIFLQVQSKKIKETELGKPHEYLASFGGKFESWVSNYLRQEE

EVKRQLFGYEENKKGQKKFIVGNKQELDKIIRGTDEYEIKAISKETIGLTQKCLKLLEQL

KDSVDDYTLSLYRQLIVELRIRLNVEFQETYPELIGKSEKDKEKDAKNKRADKRYPQIFK

DIKLIPNFLGETKQMVYKKFIRSADILYEGINFIDQIDKQITQLLPCFKNDKERIEFTEK

QFETLRRKYYLMNSSRFHHVIEGIIKKRENSELKTFSDKKGKKYENEVYYTFYINPKARD

QRRIKIVLDINGNNSVGILQDLVQKLKPKWDDIIKKNDMGELIDAIEIEKVRLGILIALY

CEHKFKIKKELLSLDLFASAYQYLELEDDPEELSGTNLGRFLQSLVCSEIKGAINKISRT

EYIERYTVQPMNTEKNYPLLINKEGKATWHIAAKDDLSKKKGGGTVAMNQKIGKNFFGKQ

DYKTVFMLQDKRFDLLTSKYHLQFLSKTLDTGGGWWKNKNIDLNLSSYSFIFEQKVKVEW

DLTNLDHPIKPSENSDDRRLFVSIPFVIKPKQTKRKDLQTRVNYMGIDIGEYGLAWTIIN

IDLKNKKINKISKQGFIYEPLTHKVRDYVATIKDNQVRGTFGMPDTKLARLRENAITSLR

NQVHDIAMRYDAKPVYEFEISNFETGSNKVKVIYDSVKRADIGRGQNNTEADNTEVNLVW

GKTSKQFGSQIGAYATSYICSFCGYSPYYEFENSKSGDEEGARDNLYQMKKLSRPSLEDF

LQGNPVYKTFRDFDKYKNDQRLQKTGDKDGEWKTHRGNTAIYACQKC*

>Cas12d_12_sd2019_length_1197

MAESKQMQCRKCGASMKYEVIGLGKKSCRYMCPDCGNHTSARKIQNKKKRDKKYGSASKA

QSQRIAVAGALYPDKKVQTIKTYKYPADLNGEVHDRGVAEKIEQAIQEDEIGLLGPSSEY

ACWIASQKQSEPYSVVDFWFDAVCAGGVFAYSGARLLSTVLQLSGEESVLRAALASSPFV

DDINLAQAEKFLAVSRRTGQDKLGKRIGECFAEGRLEALGIKDRMREFVQAIDVAQTAGQ

RFAAKLKIFGISQMPEAKQWNNDSGLTVCILPDYYVPEENRADQLVVLLRRLREIAYCMG

IEDEAGFEHLGIDPGALSNFSNGNPKRGFLGRLLNNDIIALANNMSAMTPYWEGRKGELI

ERLAWLKHRAEGLYLKEPHFGNSWADHRSRIFSRIAGWLSGCAGKLKIAKDQISGVRTDL

FLLKRLLDAVPQSAPSPDFIVSISALDRFLEAAESSQDPAEQVRALYAFHLNAPAVRSIA

NKAVQRSDSQEWLIKELDAVDHLEFNKAFPFFSDTGKKKKKGANSNGAPSEEEYTETESI

QQPEDAEQEVNGQEGNGASKNQKKFQRIPRFFGEGSRSEYRILTEAPQYFDMFCNNMRAI

FMQLESQPRKAPRDFKCFLQNRLQKLYKQTFLNARSNKCRALLESVLISWGEFYTYGANE

KKFRLRHEASERSSDPDYVVQQALEIARRLFLFGFEWRDCSAGERVDLVEIHKKAISFLL

AITQAEVSVGSYNWLGNSTVSRYLSVAGTDTLYGTQLEEFLNATVLSQMRGLAIRLSSQE

LKDGFDVQLESSCQDNLQHLLVYRASRDLAACKRATCPAELDPKILVLPAGAFIASVMKM

IERGDEPLAGAYLRHRPHSFGWQIRVRGVAEVGMDQGTALAFQKPTESEPFKIKPFSAQY

GPVLWLNSSSYSQSQYLDGFLSQPKNWSMRVLPQAGSVRVEQRVALIWNLQAGKMRLERS

GARAFFMPVPFSFRPSGSGDEAVLAPNRYLGLFPHSGGIEYAVVDVLDSAGFKILERGTI

AVNGFSQKRGERQEEAHREKQRRGISDIGRKKPVQAEVDAANELHRKYTDVATRLGCRIV

VQWAPQPKPGTAPTAQTVYARAVRTEAPRSGNQEDHARMKSSWGYTWSTYWEKRKPEDIL

GISTQVYWTGGIGESCPAVAVALLGHIRATSTQTEWEKEEVVFGRLKKFFPGETIF*

>Cas12d_14_sd2019_length_1193

MAESKQMQCRKCGASMKYEVIGLGKKSCRYMCPDCGNHTSARKIQNKKKRDKKYGSASKA

QSQRIAVAGALYPDKKVQTIKTYKYPADLNGEVHDSGVAEKIAQAIQEDEIGLLGPSSEY

ACWIASQKQSEPYSVVDFWFDAVCAGGVFAYSGARLLSTVLQLSGEESVLRAALASSPFV

DDINLAQAEKFLAVSRRTGQDKLGKRIGECFAEGRLEALGIKDRMREFVQAIDVAQTAGQ

RFAAKLKIFGISQMPEAKQWNNDSGLTVCILPDYYVPEENRADQLVVLLRRLREIAYCMG

IEDEAGFEHLGIDPGALSNFSNGNPKRGFLGRLLNNDIIALANNMSAMTPYWEGRKGELI

ERLAWLKHRAEGLYLKEPHFGNSWADHRSRIFSRIAGWLSGCAGKLKIAKDQISGVRTDL

FLLKRLLDAVPQSAPSPDFIASISALDRFLEAAESSQDPAEQVRALYAFHLNAPAVRSIA

NKAVQRSDSQEWLIKELDAVDHLEFNKAFPFFSDTGKKKKKGANSNGAPSEEEYTETESI

QQPEDAEQEVNGQEGNGASKNQKKFQRIPRFFGEGSRSEYRILTEAPQYFDMFCNNMRAI

FMQLESQPRKAPRDFKCFLQNRLQKLYKQTFLNARSNKCRALLESVLISWGEFYTYGANE

KKFRLRHEASERSSDPDYVVQQALEIARRLFLFGFEWRDCSAGERVDLVEIHKKAISFLL

AITQAEVSVGSYNWLGNSTVSRYLSVAGTDTLYGTQLEEFLNATVLSQMRGLAIRLSSQE

LKDGFDVQLESSCQDNLQHLLVYRASRDLAACKRATCPAELDPKILVLPAGAFIASVMKM

IERGDEPLAGAYLRHRPHSFGWQIRVRGVAEVGMDQGTALAFQKPTESEPFKIKPFSAQY

GPVLWLNSSSYSQSQYLDGFLSQPKNWSMRVLPQAGSVRVEQRVALIWNLQAGKMRLERS

GARAFFMPVPFSFRPSGSGDEAVLAPNRYLGLFPHSGGIEYAVVDVLDSAGFKILERGTI

AVNGFSQKRGERQEEAHREKQRRGISDIGRKKPVQAEVDATNELHRKYTDVATRLGCRIV

VQWAPQPKPGTAPTAQTVYARAVRTEAPRSGNQEDHARMKSSWGYTWSTYWEKRKPEDIL

GISTQVYWTGGIGESCPAVAVALLGHIRATSTQTEWEKEEVVFGRLKKFFPS*

>Cas12e_2_sd2019_length_987

MEKRINKIRKKLSADNATKPVSRSGPMKTLLVRVMTDDLKKRLEKRRKKPEVMPQVISNN

AANNLRMLLDDYTKMKEAILQVYWQEFKDDHVGLMCKFAQPASKKIDQNKLKPEMDEKGN

LTTAGFACSQCGQPLFVYKLEQVSEKGKAYTNYFGRCNVAEHEKLILLAQLKPEKDSDEA

VTYSLGKFGQRALDFYSIHVTKESTHPVKPLAQIAGNRYASGPVGKALSDACMGTIASFL

SKYQDIIIEHQKVVKGNQKRLESLRELAGKENLEYPSVTLPPQPHTKEGVDAYNEVIARV

RMWVNLNLWQKLKLSRDDAKPLLRLKGFPSFPVVERRENEVDWWNTINEVKKLIDAKRDM

GRVFWSGVTAEKRNTILEGYNYLPNENDHKKREGSLENPKKPAKRQFGDLLLYLEKKYAG

DWGKVFDEAWERIDKKIAGLTSHIEREEARNAEDAQSKAVLTDWLRAKASFVLERLKEMD

EKEFYACEIQLQKWYGDLRGNPFAVEAENRVVDISGFSIGSDGHSIQYRNLLAWKYLENG

KREFYLLMNYGKKGRIRFTDGTDIKKSGKWQGLLYGGGKAKVIDLTFDPDDEQLIILPLA

FGTRQGREFIWNDLLSLETGLIKLANGRVIEKTIYNKKIGRDEPALFVALTFERREVVDP

SNIKPVNLIGVDRGENIPAVIALTDPEGCPLPEFKDSSGGPTDILRIGEGYKEKQRAIQA

AKEVEQRRAGGYSRKFASKSRNLADDMVRNSARDLFYHAVTHDAVLVFENLSRGFGRQGK

RTFMTERQYTKMEDWLTAKLAYEGLTSKTYLSKTLAQYTSKTCSNCGFTITTADYDGMLV

RLKKTSDGWATTLNNKELKAEGQITYYNRYKRQTVEKELSAELDRLSEESGNNDISKWTK

GRRDEALFLLKKRFSHRPVQEQFVCLDCGHEVHADEQAALNIARSWLFLNSNSTEFKSYK

SGKQPFVGAWQAFYKRRLKEVWKPNA*

>Cas12e_3_sd2019_length_979

MQEIKRINKIRRRLVKDSNTKKAGKTGPMKTLLVRVMTPDLRERLENLRKKPENIPQPIS

NTSRANLNKLLTDYTEMKKAILHVYWEEFQKDPVGLMSRVAQPAPKNIDQRKLIPVKDGN

ERLTSSGFACSQCCQPLYVYKLEQVNDKGKPHTNYFGRCNVSEHERLILLSPHKPEANDE

LVTYSLGKFGQRALDFYSIHVTRESNHPVKPLEQIGGNSCASGPVGKALSDACMGAVASF

LTKYQDIILEHQKVIKKNEKRLANLKDIASANGLAFPKITLPPQPHTKEGIEAYNNVVAQ

IVIWVNLNLWQKLKIGRDEAKPLQRLKGFPSFPLVERQANEVDWWDMVCNVKKLINEKKE

DGKVFWQNLAGYKRQEALLPYLSSEEDRKKGKKFARYQFGDLLLHLEKKHGEDWGKVYDE

AWERIDKKVEGLSKHIKLEEERRSEDAQSKAALTDWLRAKASFVIEGLKEADKDEFCRCE

LKLQKWYGDLRGKPFAIEAENSILDISGFSKQYNCAFIWQKDGVKKLNLYLIINYFKGGK

LRFKKIKPEAFEANRFYTVINKKSGEIVPMEVNFNFDDPNLIILPLAFGKRQGREFIWND

LLSLETGSLKLANGRVIEKTLYNRRTRQDEPALFVALTFERREVLDSSNIKPMNLIGIDR

GENIPAVIALTDPEGCPLSRFKDSLGNPTHILRIGESYKEKQRTIQAAKEVEQRRAGGYS

RKYASKAKNLADDMVRNTARDLLYYAVTQDAMLIFENLSRGFGRQGKRTFMAERQYTRME

DWLTAKLAYEGLPSKTYLSKTLAQYTSKTCSNCGFTITSADYDRVLEKLKKTATGWMTTI

NGKELKVEGQITYYNRYKRQNVVKDLSVELDRLSEESVNNDISSWTKGRSGEALSLLKKR

FSHRPVQEKFVCLNCGFETHADEQAALNIARSWLFLRSQEYKKYQTNKTTGNTDKRAFVE

TWQSFYRKKLKEVWKPAV*

>Cas12g_0_sd2019_length_767

MNRTRYREERTITRGMRRLPGEERKSFKAKVITLRRNFEQFNTDVSEICQWLMSIRPNGK

HNIPNTEPFWDFILEPHNFVVNQEETNIDSVRLVVFEMAVGWRQVTDVANFELERQLLMS

LESIQSVPRTIAAKRMLQRIKNYEFQHKMVLLRSAVEWINTRFIRTYKNWEMNIKEFLEK

KKVWENDHPKLTEEIRNTFNKVFDELEISKKNPNICRWSHLKKNRDNCNYAGVRIKVGGE

YNNHSEKCKRYQDFLKKHSAHKKYFAANAMMYINIRKKRRDLTKREAIKVLLDKIPQARS

WFPQAWDNYLEYLGLNEISLINKFDGQLPHCLRLDTECIYNVHTQSCRKYYVLLKDLPDK

YLSLEETYREWRKYFLREPRKPVFAYPSTRQRTVSKIFGRDYFEADYDNSIIKLRLDDMA

EGQFLSFGFKPWPVDYDVQPIDTEITSVLVHFIGTRARVGFRFKMPHRPSRINIKQDELD

ELRSRSRLIQEKDQALLEKVRLRLRDGFIGIFDKELRVLAVDLGTSSCATAFFVGRQFQE

SSRLQIVKYDRVYKSNYEIKKRRNNKGIDKQKQLLFKEKGLNQYHIKVHLDKLAEQNKQI

IKKREASGNPTPTEQDMRRLSLHIGWMHRDWVRINASQIIKSAKKLRADLIVFESLRDFR

PMMFNEFDIDKKRRLAFFPFGLIRHKVIEKAVESGMRVVTVPYMFSSQFCGACGRQQNDK

KRLQKNKTDKRFICEYNDCAFEGDPDENAARVLGGVFWGNIGLPLS*

>Cas12g_1_sd2019_length_794

MHPSRYKTARTLVRRLCRLPGEDRSAFRSKVGLLRGHFEQFNVDVSELCQWLMSLRKRNK

VPENPATALGDFLLQPGLPGEETDEKEADRLRLAVFDAVAGFRMLEDRLAASIPASLSDA

IRDVRAAGKPSGLARVLARLEACAPAQRLVLLKSAAEWIVARFLRGTENWMRQRAEWEKE

KAAWEAAHPHLTPEVRAQFNKIFESLHDPENSGKPGVSRKNPRICPWDRLKQNLDNCCYG

EKGHSALCWRYQDFLKQRMGENRRKKNFSATAMDLAQICREWKIQHSRNALNNPRVLDRL

FAEKANPKADYLFKAHWKAYLEHMKLNDTTVLERGCLPHCLSIKKNGKESTCKWNKHTEL

CLEYKRSLAPLPDSVLELEPEYREWRRLYLHGPGRPHFRYPSAGELPLPKVFGEGFHQVD

LDRSIVRLRLEGAAEGEWLEFGFIPWPRGYQPSRREVLITSVQVHFVGTRPRAGFRFDVS

HRTSRFGCSQDELDELRSRRYPRQAQDKEFLAAARAQLIQTFEGGAARQQMRVMSVDLGE

GGACASIYEGRTHQKDESLKVIKIDRRYDQHPEVLEKDVGAAKPQKFEKSDPRGVRKEHV

ARHLNRIAAGASAIAEHRRKERSDAECSVGELQEHDFRSLKRHIAWMIRDWVRLNAAQII

DVAKQHCCDLIVFESQRGFRLPGYDELDRGKKQRFAILAFGRIRRKVVEKAVEHGMRVVT

VPYFASSQVCSACKRVQENRGSWRENKKKRVFACEFCKLKLNSDANASRVLARVFWGEIE

LPEPTRAHLPSKA*

>Cas12g_2_sd2019_length_793

MLPTRYKPARTLVRPLGRLPHEPRKEFVEKCRRVRMHFEQFNIDVADLCQWLMSLRPNTR

IGDAQSTVFWDFFLNPSILTVEADEKERDRWRLAAFDELLQIRFGHDPNAPPWSEEFRSA

IRHVAQRPKSATAQRLFDRLRSLTAPHRLVLLKSAAEWIIARYQRGMENWQRQFAEWQRE

KEEWEAAHPNLTPEVRDAFTRVFKNLFEPDGDGKIGVRRKNPRICSWERLKLNKDNCVYA

GQKGHGPLCWEFSKFVKAQKNAGTIKTFFVDVANKYLHVRRNLSKPGKLKKSPRQEAFKR

LYNQKGMEKARNWFTDAWSGYLTALNLNEKTILDHGCLKHCGAIGAEFEKSLCQFNPHTH

LCVQYRNALESLEPAIRELEGDYREWRRLFLAPPRKPSFRYPSSRRLPMPKIFGEHFHQI

DFDQSILRLRLEDMAEGEWIEFGFKPWPKDYRPGKDEVRVTSVHVNFHGNRMRAGFHFEA

PAKPSRFACTQDELDDLRSKQFPRQSQDRQLLEVARRRLLESFDGMLESDLRILAVDLGE

KGAAAAVYQGHGHEADVAIPIVKIDRLYDHVPDVLDVESARVPPPKFDDSRDPRGVRKEH

VGRHLGQLQRGAQTLAQHRQQDESAPAALRRHDFRSLTRHIRWMIRDWTRHNAAQITAAA

ETHRCHLIVFESLRGFKPRGYDQMDFAQKARLAFFAYGRVRRKVVEKAVERGLRVVTVPY

GFTSQICSECGHRQRNKGRLRKNKYQRRFVCECGTCRLQLGSDVNAARVLARVFWDEIVL

PTREEMREPAVD*

>Cas12g_4_sd2019_length_720

MSIRPDAKKPDKETKSFWDFFLNPESFFDPTINNVDIIRLNLFKVITGRESEANLIRYNL

PLLLYESIILLKKQEPSDTARRLFARLKKMEPVHVMILLKAAAEWVYARYQRLMDNHEYQ

YKVWHDEKSAWENKHPELTPEIREKYNSIFKELGRKQGVTRKNPRICNWEKLEENKDNCG

YNGKRIQFGDKWKAHSMLCIEYRNFLRDNKITGKRIGFFATHAYNYLKLRAHQPRLTKDE

AFKRIFKSAPNGIYWFPKAWKNYLQFMNLNELNLIRKYNANLPHCLEFKGDKDCQYNKHT

ELCQEYKTLLLEKFTEDELKLEGLYREWRKQYLSGPSKPAFRYPSCSKLPTPKIFGKRFH

EIDFENSIVRLRLDDMPDGEYLTFKFKPWPNDYQPQPEEAEISSVHVHFVGTRARVGFRF

KIAHKQSRFKTSQDEIDELRSRKYPRQAQDADFLKAAREKLLQSFKGENPTKEIKIMAVD

LGEYRGYISVYKGENIEISEPLSILKIDKLYDSLESAGVDKTDLAKYIKDHKGLIKEHVD

SHLKVISEKANEITKHRPAGKKTGASNLKDYDLRSLTAHTGWMIRDWVRLNVSQIIRIAE

KHEVDLIVLESLRGWKAPGYDEFDLRKKRWLAFFSYGRIRHKLKEKAVERGMMVVTVPYY

KSSQICSKCGKEQENKGLWKKNKNERLFICDYPGCGHRDNSDANAAKVLAKIFWGEIVL*

>Cas12g_6_sd2019_length_749

RPRPRYREERTLVRKLLPRPGQSKQEFRENVKKLRKAFLQFNADVSGVCQWAIQFRPRYG

KPAEPTETFWKFFLEPETSLPPNDSRSPEFRRLQAFEAAAGINGAAALDDPAFTNELRDS

ILAVASRPKTKEAQRLFSRLKDYQPAHRMILAKVAAEWIESRYRRAHQNWERNYEEWKKE

KQEWEQNHPELTPEIREAFNQIFQQLEVKEKRVRICPAARLLQNKDNCQYAGKNKHSVLC

NQFNEFKKNHLQGKAIKFFYKDAEKYLRCGLQSLKPNVQGPFREDWNKYLRYMNLKEETL

RGKNGGRLPHCKNLGQECEFNPHTALCKQYQQQLSSRPDLVQHDELYRKWRREYWREPRK

PVFRYPSVKRHSIAKIFGENYFQADFKNSVVGLRLDSMPAGQYLEFAFAPWPRNYRPQPG

ETEISSVHLHFVGTRPRIGFRFRVPHKRSRFDCTQEELDELRSRTFPRKAQDQKFLEAAR

KRLLETFPGNAEQELRLLAVDLGTDSARAAFFIGKTFQQAFPLKIVKIEKLYEQWPNQKQ

AGDRRDASSKQPRPGLSRDHVGRHLQKMRAQASEIAQKRQELTGTPAPETTTTLQPFDLR

GLTVHTARMIRDWARLNARQIIQLAEENQVDLIVLESLRGFRPPGYENLDQEKKRRVAFF

AHGRIRRKVTEKAVERGMRVVTVPYLASSKVCAECRKKQKDNKQWEKNKKRGLFKCEGCG

SQAQVDENAARVLGRVFWGEIELPTAIP*

>Cas12g_8_sd2019_length_770

MNRTRYREERTITRGMRRLPGEERKSFKAKVITLRRNFEQFNTDVSEICQWLMSIRPNGK

HNIPNTEPFWDFILEPHNFVVNQEETNIDSVRLVVFEMAVGWRQVTDVANFELERQLLMS

LESIQSVPRTIAAKRMLQRIKNYEFQHKMVLLRSAVEWINTRFIRTYKNWEMNIKEFLEK

KKVWENDHPKLTEEIRNTFNKVFDELEISKKNPNICRWSHLKKNRDNCNYAGVRIKVGGE

YNNHSEKCKRYQDFLKKHSAHKKYFAANAMMYINIRKKRRDLTKREAIKVLLDKIPQARS

WFPQAWDNYLEYLGLNEISLINKFDGQLPHCLRLDTECIYNVHTQSCRKYYVLLKDLPDK

YLSLEETYREWRKYFLREPRKPVFAYPSTRQRTVSKIFGRDYFEADYDNSIIKLRLDDMA

EGQFLSFGFKPWPVDYDVQPIDTEITSVLVHFIGTRARVGFRFKMPHRPSRINIKQDELD

ELRSRSRLIQEKDQALLEKVRLRLRDGFIGIFDKELRVLAVDLGTSSCATAFFVGRQFQE

SSRLQIVKYDRVYKSNYEIKKRRNNKGIDKQKQLLFKEKGLNQYHIKVHLDKLAEQNKQI

IKKREASGNPTPTEQDMRRLSLHIGWMHRDWVRINASQIIKSAKKLRADLIVFESLRDFR

PMMFNEFDIDKKRRLAFFPFGLIRHKVIEKAVESGMRVVTVPYMFSSQFCGACGRQQNDK

KRLQKNKTDKRGACFICEYNDCAFEGDPDENAARVLGGVFWGNIGLPLS*

>Cas12g_10_sd2019_length_822

MLPTRYKPARTLVRPLGRLPHEPRKEFVEKCRRVRMHFEQFNIDVADLCQWLMSLRPNTR

IGDAQSTVFWDFFLNPSILTVEADEKERDRWRLAAFDELLQIRFGHDPNAPPWSEEFRSA

IRHVAQRPKSATAQRLFDRLRSLTAPHRLVLLKSAAEWIIARYQRGMENWQRQFAEWQRE

KEEWEAAHPNLTPEVRDAFTRVFKNLFENPDGDGKIGVRRKNPRICSWERLKLNKDNCVY

AGQKGHGPLCWEFSKFVKAQKNAGTIKTFFVDVANKYLHVRRNLSKPGVKLKKSPRQEAF

KRLYNQKGMEKARNWFTDAWSGYLTALNLNEKTILDHGCLKHCGAIGAEFEKSLCQFNPH

THLCVQYRNALESLEPAIRELEGDYREWRRLFLAPPRKPSFRYPSSRRLPMPKIFGEHFH

QIDFDQSILRLRLEDMAEGEWIEFGFKPWPKDYRPGKDEVRVTSVHVNFHGNRMRAGFHF

EAPAKPSRFACTQDELDDLRSKQFPRQSQDRQLLEVARRRLLESFDGMLESDLRILAVDL

GEKGAAAAVYQGHGHEADVAIPIVKIDRLYDHVPDVLDVESARVPPPKFDDSRDPRGVRK

EHVGRHLGQLQRGAQTLAQHRQQDESAPAALRRHDFRSLTRHIRWMIRDWTRHNAAQITA

AAETHRCHLIVFESLRGFKPRGYDQMDFAQKARLAFFAYGRVRRKVVEKAVERGLRVVTV

PYGFTSQICSECGHRQRNKGRLRKNKYQRRFVCECGEPKKSANKTAAPDRSATVSPCTCR

LQLGSDVNAARVLARVFWDEIVLPTREEMREPAVDSAPPSK*

>Cas12g_12_sd2019_length_721

MSIRPDAKKPDKETKSFWDFFLNPESFFDPTINNVDIIRLNLFKVITGRESEANLIRYNL

PLLLYESIILLKKQEPSDTARRLFARLKKMEPVHVMILLKAAAEWVYARYQRLMDNHEYQ

YKVWHDEKSAWENKHPELTPEIREKYNSIFKELGRKQGVTIRKNPRICNWEKLEENKDNC

GYNGKRIQFGDKWKAHSMLCIEYRNFLRDNKITGKRIGFFATHAYNYLKLRAHQPRLTKD

EAFKRIFKSAPNGIYWFPKAWKNYLQFMNLNELNLIRKYNANLPHCLEFKGDKDCQYNKH

TELCQEYKTLLLEKFTEDELKLEGLYREWRKQYLSGPSKPAFRYPSCSKLPTPKIFGKRF

HEIDFENSIVRLRLDDMPDGEYLTFKFKPWPNDYQPQPEEAEISSVHVHFVGTRARVGFR

FKIAHKQSRFKTSQDEIDELRSRKYPRQAQDADFLKAAREKLLQSFKGENPTKEIKIMAV

DLGEYRGYISVYKGENIEISEPLSILKIDKLYDSLESAGVDKTDLAKYIKDHKGLIKEHV

DSHLKVISEKANEITKHRPAGKKTGASNLKDYDLRSLTAHTGWMIRDWVRLNVSQIIRIA

EKHEVDLIVLESLRGWKAPGYDEFDLRKKRWLAFFSYGRIRHKLKEKAVERGMMVVTVPY

YKSSQICSKCGKEQENKGLWKKNKNERLFICDYPGCGHRDNSDANAAKVLAKIFWGEIVL

*

>Cas12i_0_sd2019_length_1001

MMSDNIILPYNSKLAPDERKQRLLNDTFNWFDMCNEVFFDFVKNLYGGVKHEKPKKVSNS

KKPKKKDVDNACATFWFRLQAKSTLDQSVQTAEERIRRFRDYAGHEPSSFAKSYLNGNYD

PEKTEWVDCRLLYVNFCRNLNVNLDADIRTMVEHNLLPVLPGQDFKTNNVFSNIFGVGNK

EDKGQKTNWLNTVSEGLQSKEIWNWDEYRDLISRSTGCSTAAELRSESIGRPSMLAVDFA

SEKSGQISQEWLAERVKSFRAAASQKSKIYDMPNRLVLKEYIASKIGPFKLERWSAAAVS

AYKDVRSKNSINLLYSKERLWRCKEIAQIDNTQVAEAQQILVNYSSGDTNSFTVENRHMG

DLTVLFKIWEKMDMDSGIEQYSEIYRDEYSRDPITELLRYLYNHRHISAKTFRAAARLNS

LLLKNDRKKIHPTISGRTSVSFGHSTIKGCITPPDHIVKNRKENAGSTGMIWVTMQLIDN

GRWADHHIPFHNSRYYRDFYAYRADLPTISDPRRKSFGHRIGNNISDTRMINHDCKKASK

MYLRTIQNMTHNVAFDQQTQFAVRRADNNFTITIQARVVGRKYKKEISVGDRVMGVDQNQ

TTSNTYSVWEVVAEGTENSYPYKGNNYRLVEDGFIRSECSGRDQLSYDGLDFQDFAQWRR

ERYAFLSSVGCILNDEIEPQIPVSAKKKFSKWRGCSLYSWNLCYAYYLKGLMHENLANNP

AGFRQEILNFIQGSRGVRLCSLNHTSFRLLSKAKSLIHSFFGLNNIKDPESQRDFDPEIY

DIMVNLTQRKTNKRKEKANRITSSILQIANRLNVSRIVIENDLPNASSKNKASANQRATD

WCARNVSEKLEYACKMLGISLWQIDPRDTSHLDPFVVGKEARFMKIKVSDINEYTISNFK

KWHANIATTSTTAPLYHDALKAFSSHYGIDWDNLPEMKFWELKNALKDHKEVFIPNRGGR

CYLSTLPVTSTSEKIVFNGRERWLNASDIVAGVNIVLRSV*

>Cas12i_1_sd2019_length_1047

MVSESTIRPYTSKLAPNDPKLKMLNDTFNWLDHAYKVFFDVSVALFGAIEHETAQELIGE

KSKFDADLLCAIMWFRLEEKSDNPGPLQTVEQRMRLFQKYSGHEPSSFTQEYIKGNIDSE

KYQWVDCRLKFIDLARNINTTQESLKIDAYTLFMNKLIPVSKDDEFNAYGLISQLFGTGK

KEDRSIKASMLEEISNIIEDKKPNTWEEYHDLIKKTFNVDNYKELKEKLSAGSSGRDSSL

VIDLKEEKTGLLQPNFIKNRIVKFREDADKKRTVFLLPNRMKLREFIASQIGPFEQNSWS

AVLNRSMAAIQSKNSSNILYTNEKEERNNEIQELLKKDILSAASILGDFRRGEFNRSVVS

KNHLGARLNELFEIWQELTMDDGIKKYVDLCKDKFSRRPVKALLQYIYPYFDKINAKQFL

DAASYNTLVETNNRKKIHPTVTGPTVCNWGPKSTINGSITPPNQMVKGRPAGSHGMIWVT

MTVIDNGRWIKHHLPFHNSRYYEEHYCYREGLPTKNKPRTKQLGTQVGSTISAPSLAILK

SQEEQDRRNDRKNRFKAHKSIIRSQENIEYNVAFDKSTNFDVTRKNGEFFITISSRVATP

KYSYKLNIGDMIMGLDNNQTAPCTYSIWRVVEKDTEGSFFHNKIWLQLVTDGKVTSIVDN

NRQVDQLSYAGIEYSNFAEWRKDRRQFLRSINEDYVKKSDNWRNMNLYQWNAEYSRLLLD

VMKENKGKNIQNTFRAEIEELICGKFGIRLGSLFHHSLQFLTNCKSLISSYFMLNNKKEE

YDQELFDSDFFRLMKSIGDKRVRKRKEKSSRISSTVLQIARENNVKSLCVEGYLPTSTKK

TKPKQNQKSIDWCARAVVKKLNDGCKVLGINLQAIDPRDTSHLDPFVYYGKKSTKVGKEA

RYTIVEPSNIKEYMTNRFDDWHRGVTKKSKKGDVQTSTTVLLYQEALRQFASHYKLDFDS

LPKMKFYELAKILGDHEKVIIPCRGGRAYLSTYPVTKDSSKITFNGRERWYNESDVVAAV

NIVLRGIIDEDEQPDGAKKQALARTK*

>Cas12i_2_sd2019_length_1052

YRSNVMVSESTIRPYTSKLAPNDPKLKMLNDTFNWLDHAYKVFFDVSVALFGAIEHETAQ

ELIGEKSKFDADLLCAIMWFRLEEKSDNPGPLQTVEQRMRLFQKYSGHEPSSFTQEYIKG

NIDSEKYQWVDCRLKFIDLARNINTTQESLKIDAYTLFMNKLIPVSKDDEFNAYGLISQL

FGTGKKEDRSIKASMLEEISNIIEDKKPNTWEEYHDLIKKTFNVDNYKELKEKLSAGSSG

RDSSLVIDLKEEKTGLLQPNFIKNRIVKFREDADKKRTVFLLPNRMKLREFIASQIGPFE

QNSWSAVLNRSMAAIQSKNSSNILYTNEKEERNNEIQELLKKDILSAASILGDFRRGEFN

RSVVSKNHLGARLNELFEIWQELTMDDGIKKYVDLCKDKFSRRPVKALLQYIYPYFDKIN

AKQFLDAASYNTLVETNNRKKIHPTVTGPTVCNWGPKSTINGSITPPNQMVKGRPAGSHG

MIWVTMTVIDNGRWIKHHLPFHNSRYYEEHYCYREGLPTKNKPRTKQLGTQVGSTISAPS

LAILKSQEEQDRRNDRKNRFKAHKSIIRSQENIEYNVAFDKSTNFDVTRKNGEFFITISS

RVATPKYSYKLNIGDMIMGLDNNQTAPCTYSIWRVVEKDTEGSFFHNKIWLQLVTDGKVT

SIVDNNRQVDQLSYAGIEYSNFAEWRKDRRQFLRSINEDYVKKSDNWRNMNLYQWNAEYS

RLLLDVMKENKGKNIQNTFRAEIEELICGKFGIRLGSLFHHSLQFLTNCKSLISSYFMLN

NKKEEYDQELFDSDFFRLMKSIGDKRVRKRKEKSSRISSTVLQIARENNVKSLCVEGYLP

TSTKKTKPKQNQKSIDWCARAVVKKLNDGCKVLGINLQAIDPRDTSHLDPFVYYGKKSTK

VGKEARYTIVEPSNIKEYMTNRFDDWHRGVTKKSKKGDVQTSTTVLLYQEALRQFASHYK

LDFDSLPKMKFYELAKILGDHEKVIIPCRGGRAYLSTYPVTKDSSKITFNGRERWYNESD

VVAAVNIVLRGIIDEDEQPDGAKKQALARTK*

>Cas12i_3_sd2019_length_1041

MSSAIKSYKSVLRPNERKNQLLKSTIQCLEDGSAFFFKMLQGLFGGITPEIVRFSTEQEK

QQQDIALWCAVNWFRPVSQDSLTHTIASDNLVEKFEEYYGGTASDAIKQYFSASIGESYY

WNDCRQQYYDLCRELGVEVSDLTHDLEILCREKCLAVATESNQNNSIISVLFGTGEKEDR

SVKLRITKKILEAISNLKEPKNVAPIQEIILNVAKAETFRQVYAGNLGAPSTLEKFIAKD

GQKEFDLKKLQTDLKKVIRGKSKERDWCCQEELRSYVEQNTIQYDLWAWGEMFNKAHTAL

KIKSTRNYNFAKQRLEQFKEIQSLNNLLVVKKLNDFFDSEFFSGEETYTICVHHLGGKDL

SKLYKAWEDDPADPENAIVVLCDDLKNNFKKEPIRNILRYIFTIRQECSAQDILAAAKYN

QQLDRYKSQKANPSVLGNQGFTWTNAVILPEKAQRNDRPNSLDLRIWLYLKLRHPDGRWK

KHHIPFYDTRFFQEIYAAGNSPVDTCQFRTPRFGYHLPKLTDQTAIRVNKKHVKAAKTEA

RIRLAIQQGTLPVSNLKITEISATINSKGQVRIPVKFDVGRQKGTLQIGDRFCGYDQNQT

ASHAYSLWEVVKEGQYHKELGCFVRFISSGDIVSITENRGNQFDQLSYEGLAYPQYADWR

KKASKFVSLWQITKKNKKKEIVTVEAKEKFDAICKYQPRLYKFNKEYAYLLRDIVRGKSL

VELQQIRQEIFRFIEQDCGVTRLGSLSLSTLETVKAVKGIIYSYFSTALNASKNNPISDE

QRKEFDPELFALLEKLELIRTRKKKQKVERIANSLIQTCLENNIKFIRGEGDLSTTNNAT

KKKANSRSMDWLARGVFNKIRQLAPMHNITLFGCGSLYTSHQDPLVHRNPDKAMKCRWAA

IPVKDIGDWVLRKLSQNLRAKNIGTGEYYHQGVKEFLSHYELQDLEEELLKWDRKSNIPC

WVLQNRLAEKAVVYIPVRGGRIYFATHKVATGAVSIVFDQKQVWVCNADHVAAANIALKG

IGESDEENPDGSRIKLQLTS*

>Cas12i_4_sd2019_length_1094

MSNKEKNASETRKAYTTKMIPRSHDRMKLLGNFMDYLMDGTPIFFELWNQFGGGIDRDII

SGTANKDKISDDLLLAVNWFKVMPINSKPQGVSPSNLANLFQQYSGSEPDIQAQEYFASN

FDTEKHQWKDMRVEYERLLAELQLSRSDMHHDLKLMYKEKCIGLSLSTAHYITSVMFGTG

AKNNRQTKHQFYSKVIQLLEESTQINSVEQLASIILKAGDCDSYRKLRIRCSRKGATPSI

LKIVQDYELGTNHDDEVNVPSLIANLKEKLGRFEYECEWKCMEKIKAFLASKVGPYYLGS

YSAMLENALSPIKGMTTKNCKFVLKQIDAKNDIKYENEPFGKIVEGFFDSPYFESDTNVK

WVLHPHHIGESNIKTLWEDLNAIHSKYEEDIASLSEDKKEKRIKVYQGDVCQTINTYCEE

VGKEAKTPLVQLLRYLYSRKDDIAVDKIIDGITFLSKKHKVEKQKINPVIQKYPSFNFGN

NSKLLGKIISPKDKLKHNLKCNRNQVDNYIWIEIKVLNTKTMRWEKHHYALSSTRFLEEV

YYPATSENPPDALAARFRTKTNGYEGKPALSAEQIEQIRSAPVGLRKVKKRQMRLEAARQ

QNLLPRYTWGKDFNINICKRGNNFEVTLATKVKKKKEKNYKVVLGYDANIVRKNTYAAIE

AHANGDGVIDYNDLPVKPIESGFVTVESQVRDKSYDQLSYNGVKLLYCKPHVESRRSFLE

KYRNGTMKDNRGNNIQIDFMKDFEAIADDETSLYYFNMKYCKLLQSSIRNHSSQAKEYRE

EIFELLRDGKLSVLKLSSLSNLSFVMFKVAKSLIGTYFGHLLKKPKNSKSDVKAPPITDE

DKQKADPEMFALRLALEEKRLNKVKSKKEVIANKIVAKALELRDKYGPVLIKGENISDTT

KKGKKSSTNSFLMDWLARGVANKVKEMVMMHQGLEFVEVNPNFTSHQDPFVHKNPENTFR

ARYSRCTPSELTEKNRKEILSFLSDKPSKRPTNAYYNEGAMAFLATYGLKKNDVLGVSLE

KFKQIMANILHQRSEDQLLFPSRGGMFYLATYKLDADATSVNWNGKQFWVCNADLVAAYN

VGLVDIQKDFKKK*

>Cas12i_5_sd2019_length_1034

MMSDNIILPYNSKLAPDERKQRLLNDTFNWFDMCNEVFFDFVKNLYGGVKHEHLILVNFA

EKPKKVSNSKKPKKKDQEVNIHVEPNQAEWVDNACATFWFRLQAKSTVQLDQSVQTAEER

IRRFRDYAGHEPSSFAKSYLNGNYDPEKTEWVDCRLLYVNFCRNLNVNLDADIRTMVEHN

LLPVLPGQDFKTNNVFSNIFGVGNKEDKGQKTNWLNTVSEGLQSKEIWNWDEYRDLISRS

TGCSTAAELRSESIGRPSMLAVDFASEKSGQISQEWLAERVKSFRAAASQKSKIYDMPNR

LVLKEYIASKIGPFKLERWSAAAVSAYKDVRSKNSINLLYSKERLWRCKEIAQILVDNTQ

VAEAQQILVNYSSGDTNSFTVENRHMGDLTVLFKIWEKMDMDSGIEQYSEIYRDEYSRDP

ITELLRYLYNHRHISAKTFRAAARLNSLLLKNDRKKIHPTISGRTSVSFGHSTIKGCITP

PDHIVKNRKENAGSTGMIWVTMQLIDNGRWADHHIPFHNSRYYRDFYAYRADLPTISDPR

RKSFGHRIGNNISDTRMINHDCKKASKMYLRTIQNMTHNVAFDQQTQFAVRRYADNNFTI

TIQARVVGRKYKKEISVGDRVMGVDQNQTTSNTYSVWEVVAEGTENSYPYKGNNYRLVED

GFIRSECSGRDQLSYDGLDFQDFAQWRRERYAFLSSVGCILNDEIEPQIPVSAEKAKKKK

KFSKWRGCSLYSWNLCYAYYLKGLMHENLANNPAGFRQEILNFIQGSRGVRLCSLNHTSF

RLLSKAKSLIHSFFGLNNIKDPESQRDFDPEIYDIMVNLTQRKTNKRKEKANRITSSILQ

IANRLNVSRIVIENDLPNASSKNKASANQRATDWCARNVSEKLEYACKMLGISLWQIDPR

DTSHLDPFVVGKEARFMKIKVSDINEYTISNFKKWHANIATTSTTAPLYHDALKAFSSHY

GIDWDNLPEMKFWELKNALKDHKEVFIPNRGGRCYLSTLPVTSTSEKIVFNGRERWLNAS

DIVAGVNIVLRSV*

>Cas12i_6_sd2019_length_1092

MFTLLLSDISQQNFNKFLKNFFFTRNKTVVHCSSEIRHKGYRSNVMVSESTIRPYTSKLA

PNDPKLKMLNDTFNWLDHAYKVFFDVSVALFGAIEHETAQELIGEKSKFDADLLCAIMWF

RLEEKSDNPGPLQTVEQRMRLFQKYSGHEPSSFTQEYIKGNIDSEKYQWVDCRLKFIDLA

RNINTTQESLKIDAYTLFMNKLIPVSKDDEFNAYGLISQLFGTGKKEDRSIKASMLEEIS

NIIEDKKPNTWEEYHDLIKKTFNVDNYKELKEKLSAGSSGRDSSLVIDLKEEKTGLLQPN

FIKNRIVKFREDADKKRTVFLLPNRMKLREFIASQIGPFEQNSWSAVLNRSMAAIQSKNS

SNILYTNEKEERNNEIQELLKKDILSAASILGDFRRGEFNRSVVSKNHLGARLNELFEIW

QELTMDDGIKKYVDLCKDKFSRRPVKALLQYIYPYFDKINAKQFLDAASYNTLVETNNRK

KIHPTVTGPTVCNWGPKSTINGSITPPNQMVKGRPAGSHGMIWVTMTVIDNGRWIKHHLP

FHNSRYYEEHYCYREGLPTKNKPRTKQLGTQVGSTISAPSLAILKSQEEQDRRNDRKNRF

KAHKSIIRSQENIEYNVAFDKSTNFDVTRKNGEFFITISSRVATPKYSYKLNIGDMIMGL

DNNQTAPCTYSIWRVVEKDTEGSFFHNKIWLQLVTDGKVTSIVDNNRQVDQLSYAGIEYS

NFAEWRKDRRQFLRSINEDYVKKSDNWRNMNLYQWNAEYSRLLLDVMKENKGKNIQNTFR

AEIEELICGKFGIRLGSLFHHSLQFLTNCKSLISSYFMLNNKKEEYDQELFDSDFFRLMK

SIGDKRVRKRKEKSSRISSTVLQIARENNVKSLCVEGYLPTSTKKTKPKQNQKSIDWCAR

AVVKKLNDGCKVLGINLQAIDPRDTSHLDPFVYYGKKSTKVGKEARYTIVEPSNIKEYMT

NRFDDWHRGVTKKSKKGDVQTSTTVLLYQEALRQFASHYKLDFDSLPKMKFYELAKILGD

HEKVIIPCRGGRAYLSTYPVTKDSSKITFNGRERWYNESDVVAAVNIVLRGIIDEDEQPD

GAKKQALARTK*

>Cas12k_0_sd2019_length_727

MSLITFPFRLDALGKTPKDREQVRRYFWELMTQQHTPLINRLNLLVAEHPKFANWQIKNT

VDADDLLCLWRSLKKQPQFQGIPDRACVSARLVTHNTYESWLALQADRQEKLTKLLNSVE

ILKPDVELIAIGSCNLDVIRNRAKEVLSEVNTQISNDSEKKKSQKAIEGGIYKILYDKHR

KTDDLVEKCAIAYLIKNRFKINSDETEEDVKKLNARRHTKQKEIEIIQKQLHSRLPKGRP

WAGRDIFGEIASIDVRETEWNSIESALLKQQTFMPHPILFESSDDLIWAEPEKFQLKDLQ

VEAGKSGSTELDEFQNENFQGEQANSVESRKRVCVGFKSFEEKYAFEVAGDYRHIHAVWQ

ALKERKIYDKNIDENTSALFLVRSATLIWRDYKKNENRIVRRRKANNKRSKREGGTAGTK

PDSLRAPAFYDPEFPWNRYQLFLHCTIETLYVSKEGTELKLEMQKKPITKALQTLEKNIT

ESEEKGESTKNRKDRHSRQSGTLRRMECYDDNYERPSKSLYTGQPHIVTGIALGASGLVT

ATIVDTSSGKILECRGRKALLGKQDRLVNRRQFQRQLNMRRRTQNQRRGANNQFGEANLG

DTIDRHIANAVVDFAKTYQSGCIVLPDMEDYRRRKQSEMAALAERECSGWKGVEKKFAKA

QNVKIHSWSYGRSIEYITNQAEKEGMLVKIGRQPIHGSSQEQAKQMAIEVYQDKTKFKQS

LSNQSA*

>Cas12k_1_sd2019_length_679

MSRDRQKKSTSPIHRTIRCHLHASEDVLRKVWEEMTQKNTPLIVQLLKSVSEQPEFEANQ

EKGTISKKEITKLRKALTNDSDIQQQSGRLGSSADSLVTEVYTSWLTLSQKIKKQKEGKE

YFLNNILKSDVELVEESNCDLQTIRCKAQDILSQPKEFLEKIINNDAVLNQTKSARKKVQ

NSSNEINASKQSENSDLKENVDKNIPQTLTEILYKIHKITQDILTQCAVAYLIKNHNQVS

DIEEDIKNLKKRRTEKQVQIKRLEEQIHNKKLPNGRDITGERYNQAFDNLINQVPQDNEE

FAEWIASLSTKVSHLPYPIDYLYSDLTWYKNEQEKICVYFNGWAKFHFQICCNKRQLHFF

KSFLEDYKALKESEKGETKLSGSLVTLRSVQLLWQQGEGAGAPWKVNKLALHCTYDARLL

TAEGTEDVRQEKTDTTQKQVTKAEANENIDSDEQKNLNRNISSLSRLNNSFARPSKPIYR

GQSNIIVGVSFHPVELVTLAVVDIITKEKIICKTVKQLLGDAFSLLSRRRRQQVHFRKER

KKAQKKDSPCNIGESQLGEYVDKLLAKRIVEVAKEYQAICIVLPTLKDTREIRTSVIQAK

AETKFPGDVNAQQLYVKEYNHQIHNWSYSRLQESIKSKAAELKISIEFSIQASYDTLQEQ

AINLALSAYQCRINTIGR*

>Cas12k_2_sd2019_length_693

MILVTQIEKGVFAGDIVTMSRDRKKGSKSPSLRTIRCHLHTKEDVLRKVWEEMTQKNTPL

IVELLKSVSEQPEFETNKENGKITKKEITNLRKSLTQDSGRLYSSVDNLVQEVYSAWLTL

YQKRKKQKEGKEYFLNNILKSDIELVEESNCDLQILRAKAQEIISNPQDILRQITIDNPK

DKPTKSIQKRVRKNINASNSDTAKNNILNSQEKTTEENISKSVIEILYEIHKTTQDTIIR

CAVTYLIKNYTKISDTEEDLNKLKERRAEKEIEIKRLEKQIQDSRLPNGRDITGARYLEA

FDKLINQVPKNNEEFANWIADASRKISNLPYPIDYLYSDLTWYKNQDGKIFVYFNGWSKY

HFQICCNKRQRHFFERFLEDHKAWKESEKGEVKLSGSLVTLRCVQLLWQQGEGKGEPWKV

NKLSLHCTYDTRLWTAEGTEEARKEKINKIQRQVEQAEETENLDEQQQKQLKKNKSSLSR

LNNSFNRPIQPIYQGLSNMIVGVSFHPVELATVAVVDTTTQKVIAYKTINELLDNAFHLL

SRMRRQQIHFRKERKKAQKKDSPCNLGESKLGEYVDKLLAKRIVEVAKEYQAICIVLPKL

KDMKEIRTSVIQAKAETKFPGNVNAQKLYVKEYNRQVHNWSYNRLQESIKSKAAELKISI

EFGIQLSYDTLQAQARDLALSAYQCRIHTIDR*

>Cas12k_3_sd2019_length_675

MSRDRKKGSKSPSLRTIRCHLHTKEDVLRKVWEEMTQKNTPLIVELLKSVSEQPEFETNK

ENGKITKKEITNLRKSLTQDSGRLYSSVDNLVQEVYSAWLTLYQKRKKQKEGKEYFLNNI

LKSDIELVEESNCDLQILRAKAQEIISNPQDILRQITIDNPKDKPTKSIQKRVRKNINAS

NSDTAKNNILNSQEKTTEENISKSVIEILYEIHKTTQDTIIRCAVTYLIKNYTKISDTEE

DLNKLKERRAEKEIEIKRLEKQIQDSRLPNGRDITGARYLEAFDKLINQVPKNNEEFANW

IADASRKISNLPYPIDYLYSDLTWYKNQDGKIFVYFNGWSKYHFQICCNKRQRHFFERFL

EDHKAWKESEKGEVKLSGSLVTLRCVQLLWQQGEGKGEPWKVNKLSLHCTYDTRLWTAEG

TEEARKEKINKIQRQVEQAEETENLDEQQQKQLKKNKSSLSRLNNSFNRPIQPIYQGLSN

MIVGVSFHPVELATVAVVDTTTQKVIAYKTINELLDNAFHLLSRMRRQQIHFRKERKKAQ

KKDSPCNLGESKLGEYVDKLLAKRIVEVAKEYQAICIVLPKLKDMKEIRTSVIQAKAETK

FPGNVNAQKLYVKEYNRQVHNWSYNRLQESIKSKAAELKISIEFGIQLSYDTLQAQARDL

ALSAYQCRIHTIDR*

>Cas12k_4_sd2019_length_680

MSSDRKKKSTIPVHRTIRCHLDASEDILRKVWEEMTQKNTPLILKLLKSVSEQPEFEANK

EKGEITKKEIVKLRKNVTKNPELEEQSGRLRSSAESFVKEVYSSWLTLYQKRKRQKEGKE

YFLKNILKSDVELIDESNCDLETIRSKAQEVLSQPEEFIKQLTINDEDVKPTKSARKRVN

KNINNKSTDAEQRKDSSSTNNVDKNKLETLTNILYEIHKQTQDILTRCTVAYLIKNHNKI

SNLEEDIQKLKKRRNEKIVQIKRLENQIQDNRLPSGRDITGERYSEAFGNLINQVPKNNQ

EWEDWIANLSKKISHLPYPIDYLYGDLSWYKNDVGNIFVYFNGWSEYHFKICCNKRQRHF

FERFLEDYKAFKVSQKGEEKLSGSLITLRSAQLLWQQGEGKGEPWKVHKLALHCTYDSRL

WTAEGTEEVRKEKTDKAQKRVSKAEENEKLDDIQQTQLNKDKSSLSRLKNSFNRPGKLIY

QSQSNIIVGISFHPIELATVAIVDINTKKVLACNTVKQLLGNAFHLLSRRRRQQVHLSKE

RKKAQKKDSPCNIGESKLGEYIDKLLAKRIVEIAKFYQAGCIILPRLKDMKEIRTSAIQA

KAEAKIPGDVNAQKLYVKEYNRQIHNWSYNRLQESIKSKAAEFKISIEFGIQPHYGTLEE

QAKDLAFYAYQSRNHTLGR*

>Cas12k_5_sd2019_length_666

MSSTSGKNLTNPIFRTISCDLSAKEDVLRKVWEEMSQKNAPLIVQLLKSVSEQPEFEANK

ENGTITKREITKLRRDITENSDLKEQSGRLRSSADSLVTEAYSSWLRLYQVRKNQKEGKE

YFLKNILKSDVELVEQSNCDLQTIRCKAKEILSQVEEFIEQIKNKPKINQTKLAKKKNNK

SNKSNKAIDKILNRFFIGNLDKTLTNTLYEIHRKSPDSLTQCAVAYLIKNDNKVSEAEEN

FTQLNKRRAAKEIEIKRLETQIQNTRLPIHYGRDITGEKYSQAFEKLVNQVPQDNEEFAE

WIAILLKKVSSLPYPILYCSGDLSWYKDEKGNIFVYFNGWAEYHFQICCDKRQLRFFERF

LKDYKALKPSKKEEEKVSGSLVTLRSAHLLWRQGKGKGEPWKVNKLALHCTYDARLWTAE

GTEEVRQEKTDKAQAKVNKAESNENLNSEQQKELTKNKSSLSRLKNSFARPSKPLYRGQS

NIIVGISFHPVELATLVVVDINTKEILICKTVKELLGDAFPLLSRRRRQQVHFRKEREKA

QKKDSPCHLGESKLGEYVDRLLAKRIVEVAKEYQASCIVLPGLKGIREIRTSAIQAKAET

KFPGDINVQELYVKEYNRQIHNWSYSRLQESIKSRAAELKITIKFGKQPSHSTLQEQAIN

LALSA*

>Cas12k_6_sd2019_length_672

MSRNSKKDFKSPILRTIRCHLYASEDVLRKVWEEMTQKNTPLIVQLLKSVSEQPEFEANK

ENGKITKKEITELRKSLTQDSDLDKQSGRLRSSADTFVTEVYSSWLTLYQKRKSQKERKE

YFLNNILKSDFELVEESNCDLQTIRFKAQEILSQPEKLLQQIIIENGNKAEINSSRKKGI

KNSNNQNSNTKENANEGNSKTLTDILYEIHKKTRDIVTQCAVAYLIKNYNQVSELEEDIQ

KLKKRRSQKEIEIKRLEKQIKNNQLPNGRDISGKTYIQALDNLINQVPNDNEEYANWLAC

LSKKISALPYPIDYLYGDLSWYKNEEEKIFVYFNGWADYHFQIHCNKRQRHFFERFLEDH

EAFKKNEKSEEKLSGGLITLRSAQLLWQQGEGKGEPWKVHKLALHCTYDARLWTAEGTEE

VREEKTDKTQKQVIKGEENENLDKKEQTQLKKNKSSLSRLNNSFTRPSKLVYRGQPHIIV

GVSFHPVELATVTVVDVNTKKVLTNKTVKQLLGDDFDLLSRRRRQQVHLRKEREKAQKKD

SPCNIGESKLGEYVDKLLAKRIVEVAKYYQASCIVLPTLKDTREIRTSVIQAKAETKFPG

DVNAQKLYVKEYNRQIHNWSYTRLQESIKLKAAESKISIECGIQPHYDTLQEQARDLALS

AYQCRINTIGR*

>Cas14a_1_sd2019_length_482

MKSDTKDKKIIIHQTKTLSLRIVKPQSIPMEEFTDLVRYHQMIIFPVYNNGAIDLYKKLF

KAKIQKGNEARAIKYFMNKIVYAPIANTVKNSYIALGYSTKMQSSFSGKRLWDLRFGEAT

PPTIKADFPLPFYNQSGFKVSSENGEFIIGIPFGQYTKKTVSDIEKKTSFAWDKFTLEDT

TKKTLIELLLSTKTRKMNEGWKNNEGTEAEIKRVMDGTYQVTSLEILQRDDSWFVNFNIA

YDSLKKQPDRDKIAGIHMGITRPLTAVIYNNKYRALSIYPNTVMHLTQKQLARIKEQRTN

SKYATGGHGRNAKVTGTDTLSEAYRQRRKKIIEDWIASIVKFAINNEIGTIYLEDISNTN

SFFAAREQKLIYLEDISNTNSFLSTYKYPISAISDTLQHKLEEKAIQVIRKKAYYVNQIC

SLCGHYNKGFTYQFRRKNKFPKMKCQGCLEATSTEFNAAANVANPDYEKLLIKHGLLQLK

K*

>Cas14a_3_sd2019_length_530

MAKNTITKTLKLRIVRPYNSAEVEKIVADEKNNREKIALEKNKDKVKEACSKHLKVAAYC

TTQVERNACLFCKARKLDDKFYQKLRGQFPDAVFWQEISEIFRQLQKQAAEIYNQSLIEL

YYEIFIKGKGIANASSVEHYLSDVCYTRAAELFKNAAIASGLRSKIKSNFRLKELKNMKS

GLPTTKSDNFPIPLVKQKGGQYTGFEISNHNSDFIIKIPFGRWQVKKEIDKYRPWEKFDF

EQVQKSPKPISLLLSTQRRKRNKGWSKDEGTEAEIKKVMNGDYQTSYIEVKRGSKIGEKS

AWMLNLSIDVPKIDKGVDPSIIGGIDVGVKSPLVCAINNAFSRYSISDNDLFHFNKKMFA

RRRILLKKNRHKRAGHGAKNKLKPITILTEKSERFRKKLIERWACEIADFFIKNKVGTVQ

MENLESMKRKEDSYFNIRLRGFWPYAEMQNKIEFKLKQYGIEIRKVAPNNTSKTCSKCGH

LNNYFNFEYRKKNKFPHFKCEKCNFKENADYNAALNISNPKLKSTKEEP*

>Cas14a_4_sd2019_length_501

MEVQKTVMKTLSLRILRPLYSQEIEKEIKEEKERRKQAGGTGELDGGFYKKLEKKHSEMF

SFDRLNLLLNQLQREIAKVYNHAISELYIATIAQGNKSNKHYISSIVYNRAYGYFYNAYI

ALGICSKVEANFRSNELLTQQSALPTAKSDNFPIVLHKQKGAEGEDGGFRISTEGSDLIF

EIPIPFYEYNGENRKEPYKWVKKGGQKPVLKLILSTFRRQRNKGWAKDEGTDAEIRKVTE

GKYQVSQIEINRGKKLGEHQKWFANFSIEQPIYERKPNRSIVGGLDVGIRSPLVCAINNS

FSRYSVDSNDVFKFSKQVFAFRRRLLSKNSLKRKGHGAAHKLEPITEMTEKNDKFRKKII

ERWAKEVTNFFVKNQVGIVQIEDLSTMKDREDHFFNQYLRGFWPYYQMQTLIENKLKEYG

IEVKRVQAKYTSQLCSNPNCRYWNNYFNFEYRKVNKFPKFKCEKCNLEISADYNAARNLS

TPDIEKFVAKATKGINLPEK*

>Cas14a_5_sd2019_length_508

MEEAKTVSKTLSLRILRPLYSAEIEKEIKEEKERRKQGGKSGELDSGFYKKLEKKHTQMF

GWDKLNLMLSQLQRQIARVFNQSISELYIETVIQGKKSNKHYTSKIVYNRAYSVFYNAYL

ALGITSKVEANFRSTELLMQKSSLPTAKSDNFPILLHKQKGVEGEEGGFKISADGNDLIF

EIPIPFYEYDSANKKEPFKWIKKGGQKPTIKLILSTFRRQRNKGWAKDEGTDAEIRKVIE

GKYQVSHIEINRGKKLGDHQKWFVNFTIEQPIYERKLDKNIIGGIDVGIKSPLVCAVNNS

FARYSVDSNDVLKFSKQAFAFRRRLLSKNSLKRSGHGSKNKLDPITRMTEKNDRFRKKII

ERWAKEVTNFFIKNQVGTVQIEDLSTMKDRQDNFFNQYLRGFWPYYQMQNLIENKLKEYG

IETKRIKARYTSQLCSNPSCRHWNSYFSFDHRKTNNFPKFKCEKCALEISADYNAARNIS

TPDIEKFVAKATKGINLPDKNENVILE*

>Cas14b_0_sd2019_length_611

LDLITEPIQPHKSSSLRSKEFLEYQISDFLNFSLHSLFFGLASNEGPLVDFKIYDKIVIP

KPEERFPKKESEEGKKLDSFDKRVEEYYSDKLEKKIERKLNTEEKNVIDREKTRIWGEVN

KLEEIRSIIDEINEIKKQKHISEKSKLLGEKWKKVNNIQETLLSQEYVSLISNLSDELTN

KKKELLAKKYSKFDDKIKKIKEDYGLEFDENTIKKEGEKAFLNPDKFSKYQFSSSYLKLI

GEIARSLITYKGFLDLNKYPIIFRKPINKVKKIHNLEPDEWKYYIQFGYEQINNPKLETE

NILGIDRGLTHILAYSVFEPRSSKFILNKLEPNPIEGWKWKLRKLRRSIQNLERRWRAQD

NVKLPENQMKKNLRSIEDKVENLYHNLSRKIVDLAKEKNACIVFEKLEGQGMKQHGRKKS

DRLRGLNYKLSLFDYGKIAKLIKYKAEIEGIPIYRIDSAYTSQNCAKCVLESRRFAQPEE

ISCLDDFKEGDNLDKRILEGTGLVEAKIYKKLLKEKKEDFEIEEDIAMFDTKKVIKENKE

KTVILDYVYTRRKEIIGTNHKKNIKGIAKYTGNTKIGYCMKHGQVDADLNASRTIALCKN

FDINNPEIWK*

>Cas14b_1_sd2019_length_633

MSDESLVSSEDKLAIKIKIVPNAEQAKMLDEMFKKWSSICNRISRGKEDIETLRPDEGKE

LQFNSTQLNSATMDVSDLKKAMARQGERLEAEVSKLRGRYETIDASLRDPSRRHTNPQKP

SSFYPSDWDISGRLTPRFHTARHYSTELRKLKAKEDKMLKTINKIKNGKIVFKPKRITLW

PSSVNMAFKGSRLLLKPFANGFEMELPIVISPQKTADGKSQKASAEYMRNALLGLAGYSI

NQLLFGMNRSQKMLANAKKPEKVEKFLEQMKNKDANFDKKIKALEGKWLLDRKLKESEKS

SIAVVRTKFFKSGKVELNEDYLKLLKHMANEILERDGFVNLNKYPILSRKPMKRYKQKNI

DNLKPNMWKYYIQFGYEPIFERKASGKPKNIMGIDRGLTHLLAVAVFSPDQQKFLFNHLE

SNPIMHWKWKLRKIRRSIQHMERRIRAEKNKHIHEAQLKKRLGSIEEKTEQHYHIVSSKI

INWAIEYEAAIVLESLSHMKQRGGKKSVRTRALNYALSLFDYEKVARLITYKARIRGIPV

YDVLPGMTSKTCATCLLNGSQGAYVRGLETTKAAGKATKRKNMKIGKCMVCNSSENSMID

ADLNAARVIAICKYKNLNDPQPAGSRKVFKRF*

>Cas14b_2_sd2019_length_776

MKALKLQLIPTRKQYKILDEMFWKWASLANRVSQKGESKETLAPKKDIQKIQFNATQLNQ

IEKDIKDLRGAMKEQQKQKERLLLQIQERRSTISEMLNDDNNKERDPHRPLNFRPKGWRK

FHTSKHWVGELSKILRQEDRVKKTIERIVAGKISFKPKRIGIWSSNYKINFFKRKISINP

LNSKGFELTLMTEPTQDLIGKNGGKSVLNNKRYLDDSIKSLLMFALHSRFFGLNNTDTYL

LGGKINPSLVKYYKKNQDMGEFGREIVEKFERKLKQEINEQQKKIIMSQIKEQYSNRDSA

FNKDYLGLINEFSEVFNQRKSERAEYLLDSFEDKIKQIKQEIGESLNISDWDFLIDEAKK

AYGYEEGFTEYVYSKRYLEILNKIVKAVLITDIYFDLRKYPILLRKPLDKIKKISNLKPD

EWSYYIQFGYDSINPVQLMSTDKFLGIDRGLTHLLAYSVFDKEKKEFIINQLEPNPIMGW

KWKLRKVKRSLQHLERRIRAQKMVKLPENQMKKKLKSIEPKIEVHYHNISRKIVNLAKDY

NASIVVESLEGGGLKQHGRKKNARNRSLNYALSLFDYGKIASLIKYKADLEGVPMYEVLP

AYTSQQCAKCVLEKGSFVDPEIIGYVEDIGIKGSLLDSLFEGTELSSIQVLKKIKNKIEL

SARDNHNKEINLILKYNFKGLVIVRGQDKEEIAEHPIKEINGKFAILDFVYKRGKEKVGK

KGNQKVRYTGNKKVGYCSKHGQVDADLNASRVIALCKYLDINDPILFGEQRKSFK*

>Cas14b_3_sd2019_length_778

MVTRAIKLKLDPTKNQYKLLNEMFWKWASLANRFSQKGASKETLAPKDGTQKIQFNATQL

NQIKKDVDDLRGAMEKQGKQKERLLIQIQERLLTISEILRDDSKKEKDPHRPQNFRPFGW

RRFHTSAYWSSEASKLTRQVDRVRRTIERIKAGKINFKPKRIGLWSSTYKINFLKKKINI

SPLKSKSFELDLITEPQQKIIGKEGGKSVANSKKYLDDSIKSLLIFAIKSRLFGLNNKDK

PLFENIITPNLVRYHKKGQEQENFKKEVIKKFENKLKKEISQKQKEIIFSQIERQYENRD

ATFSEDYLRAISEFSEIFNQRKKERAKELLNSFNEKIRQLKKEVNGNISEEDLKILEVEA

EKAYNYENGFIEWEYSEQFLGVLEKIARAVLISDNYFDLKKYPILIRKPTNKSKKITNLK

PEEWDYYIQFGYGLINSPMKIETKNFMGIDRGLTHLLAYSIFDRDSEKFTINQLELNPIK

GWKWKLRKVKRSLQHLERRMRAQKGVKLPENQMKKRLKSIEPKIESYYHNLSRKIVNLAK

ANNASIVVESLEGGGLKQHGRKKNSRHRALNYALSLFDYGKIASLIKYKSDLEGVPMYEV

LPAYTSQQCAKCVLKKGSFVEPEIIGYIEEIGFKENLLTLLFEDTGLSSVQVLKKSKNKM

TLSARDKEGKMVDLVLKYNFKGLVISQEKKKEEIVEFPIKEIDGKFAVLDSAYKRGKERI

SKKGNQKLVYTGNKKVGYCSVHGQVDADLNASRVIALCKYLGINEPIVFGEQRKSFK*

>Cas14b_4_sd2019_length_626

MLALKLKIMPTEKQAEILDAMFWKWASICSRIAKMKKKVSVKENKKELSKKIPSNSDIWF

SKTQLCQAEVDVGDHKKALKNFEKRQESLLDELKYKVKAINEVINDESKREIDPNNPSKF

RIKDSTKKGNLNSPKFFTLKKWQKILQENEKRIKKKESTIEKLKRGNIFFNPTKISLHEE

EYSINFGSSKLLLNCFYKYNKKSGINSDQLENKFNEFQNGLNIICSPLQPIRGSSKRSFE

FIRNSIINFLMYSLYAKLFGIPRSVKALMKSNKDENKLKLEEKLKKKKSSFNKTVKEFEK

MIGRKLSDNESKILNDESKKFFEIIKSNNKYIPSEEYLKLLKDISEEIYNSNIDFKPYKY

SILIRKPLSKFKSKKLYNLKPTDYKYYLQLSYEPFSKQLIATKTILGIDRGLKHLLAVSV

FDPSQNKFVYNKLIKNPVFKWKKRYHDLKRSIRNRERRIRALTGVHIHENQLIKKLKSMK

NKINVLYHNVSKNIVDLAKKYESTIVLERLENLKQHGRSKGKRYKKLNYVLSNFDYKKIE

SLISYKAKKEGVPVSNINPKYTSKTCAKCLLEVNQLSELKNEYNRDSKNSKIGICNIHGQ

IDADLNAARVIALCYSKNLNEPHFK*

>Cas14b_5_sd2019_length_614

MISLKLKLLPDEEQKKLLDEMFWKWASICTRVGFGRADKEDLKPPKDAEGVWFSLTQLNQ

ANTDINDLREAMKHQKHRLEYEKNRLEAQRDDTQDALKNPDRREISTKRKDLFRPKASVE

KGFLKLKYHQERYWVRRLKEINKLIERKTKTLIKIEKGRIKFKATRITLHQGSFKIRFGD

KPAFLIKALSGKNQIDAPFVVVPEQPICGSVVNSKKYLDEITTNFLAYSVNAMLFGLSRS

EEMLLKAKRPEKIKKKEEKLAKKQSAFENKKKELQKLLGRELTQQEEAIIEETRNQFFQD

FEVKITKQYSELLSKIANELKQKNDFLKVNKYPILLRKPLKKAKSKKINNLSPSEWKYYL

QFGVKPLLKQKSRRKSRNVLGIDRGLKHLLAVTVLEPDKKTFVWNKLYPNPITGWKWRRR

KLLRSLKRLKRRIKSQKHETIHENQTRKKLKSLQGRIDDLLHNISRKIVETAKEYDAVIV

VEDLQSMRQHGRSKGNRLKTLNYALSLFDYANVMQLIKYKAGIEGIQIYDVKPAGTSQNC

AYCLLAQRDSHEYKRSQENSKIGVCLNPNCQNHKKQIDADLNAARVIASCYALKINDSQP

FGTRKRFKKRTTN*

>Cas14b_6_sd2019_length_616

METLSLKLKLNPSKEQLLVLDKMFWKWASICTRLGLKKAEMSDLEPPKDAEGVWFSKTQL

NQANTDVNDLRKAMQHQGKRIEYELDKVENRRNEIQEMLEKPDRRDISPNRKDLFRPKAA

VEKGYLKLKYHKLGYWSKELKTANKLIERKRKTLAKIDAGKMKFKPTRISLHTNSFRIKF

GEEPKIALSTTSKHEKIELPLITSLQRPLKTSCAKKSKTYLDAAILNFLAYSTNAALFGL

SRSEEMLLKAKKPEKIEKRDRKLATKRESFDKKLKTLEKLLERKLSEKEKSVFKRKQTEF

FDKFCITLDETYVEALHRIAEELVSKNKYLEIKKYPVLLRKPESRLRSKKLKNLKPEDWT

YYIQFGFQPLLDTPKPIKTKTVLGIDRGVRHLLAVSIFDPRTKTFTFNRLYSNPIVDWKW

RRRKLLRSIKRLKRRLKSEKHVHLHENQFKAKLRSLEGRIEDHFHNLSKEIVDLAKENNS

VIVVENLGGMRQHGRGRGKWLKALNYALSHFDYAKVMQLIKYKAELAGVFVYDVAPAGTS

INCAYCLLNDKDASNYTRGKVINGKKNTKIGECKTCKKEFDADLNAARVIALCYEKRLND

PQPFGTRKQFKPKKP*

>Cas14b_7_sd2019_length_543

MSKTTISVKLKIIDLSSEKKEFLDNYFNEYAKATTFCQLRIRRLLRNTHWLGKKEKSSKK

WIFESGICDLCGENKELVNEDRNSGEPAKICKRCYNGRYGNQMIRKLFVSTKKREVQENM

DIRRVAKLNNTHYHRIPEEAFDMIKAADTAEKRRKKNVEYDKKRQMEFIEMFNDEKKRAA

RPKKPNERETRYVHISKLESPSKGYTLNGIKRKIDGMGKKIERAEKGLSRKKIFGYQGNR

IKLDSNWVRFDLAESEITIPSLFKEMKLRITGPTNVHSKSGQIYFAEWFERINKQPNNYC

YLIRKTSSNGKYEYYLQYTYEAEVEANKEYAGCLGVDIGCSKLAAAVYYDSKNKKAQKPI

EIFTNPIKKIKMRREKLIKLLSRVKVRHRRRKLMQLSKTEPIIDYTCHKTARKIVEMANT

AKAFISMENLETGIKQKQQARETKKQKFYRNMFLFRKLSKLIEYKALLKGIKIVYVKPDY

TSQTCSSCGADKEKTERPSQAIFRCLNPTCRYYQRDINADFNAAVNIAKKALNNTEVVTT

LL*

>c2c4_0_SD2019_length_601

MTSIPTGAVTVHTFGVHYRWELPPIIETQLRLAHEAREEFVALHLAYDADVKALWSSYPG

VASAETELEQAETDCAAVTEQAKALRVQRSSRRTVPEIQAQLRAAKNRLSEARQARRDAI

TAVRDVSAQRRRDRLKQLRADQKALYTKYTGQGLYWATVNDIAAQHKALVQRINKQRREG

RPAQLRHRRFDGTGTLTVQIQRRDAAPIRTPATLANTQGRYSYFAVLPWIAPQRWEAMAP

AERRAAGRVSVRMRVGSTGEGQPQWVDIPVQAHRWLPAQADITHVRLSVKRIAARRTAQV

LVTARTDQQHALASDRAAVAVHLGWRQTESGIQVASWRSTAPLQIPPALESVMIADPGAT

TGHIVVPPAITARLDRTDQLGSMRAESLPHVRDALAAWLHAHGPVQSQGRALTADVVRQW

RSPAQFAAVAKAWRDSEAPDGIGATLEAWRKVDVRLWDRQEHGRRRALGHRDDLYRQVSA

AIIGQARMVVVDDMDLSELARVANSDPGEPVALQRSAERRHRAAPGRFREAVEAAARRAG

LRCEAVSPKGIARIHAACGYQNPGDGRFASSLVTCEGCGEQYEVDASTTLLMLRGAGVLS

*

>c2c4_1_SD2019_length_603

MMAVTTYIIGIPYGPSGWEIPDQVRSQLRLAHDLREDLVTLQLDYEQAVKDLWSSFTDVV

SVEKVLVAAETRATELAAQVAAERSRQGITRITGPLASELTSARAEVARLRAERRSAIAE

IKADADPRLAELTEALEASRKVLYAKYCRSGDLYWGTFNAVKNHHAATVRRISAARKQGR

SAQLRHHRYNGTGTLAVELPRESKDPARTPALVAEPGGKYRNVIQIPWTEPERWEQLSRA

GRRHAGRDTVRMRCGSAGGVAQWVDIPVQMPNRMLPADADITGVQLTVTRTADTYRARLA

VTARIPDPKPVAPTDGPTVAVHMGWRKTSRGITVATWRSTSRIHVPPAWQHVMRVDRGGV

TGTIEFSNSAIRRRRTTDRLASIRDNALTGIQDKLMTWLGHHGPKPHPLREGDQIDAATV

ANWRSSARLAAVALRWRDESPADGAGLAAHLETWRRVDRRRWERQAHGLRKAVRRRDDMW

CNIAAILAGQAGRIVLDDTSIADTARRRPQLPTHVEHAIGRQLAIAAPRTLRDTIQAACV

RRGVTVTVVPATGLSRTHARCGHQNPADERYKAPPVRCEGCKRQYDPDSSATVIMLRRAR

GR*

>c2c4_2_SD2019_length_621

MTMASDDEPVQPGPVTPEGAITVHTMGVHYRWQIPEVVRQQLRLAHSLREDLVTLQLAYD

EDIKAIWSSYPTVAAAEAHVAATEAEAIAANEALKAARSAARGKRVNTAITERLHDARTA

LKAARHARREAIADVRQDAAQRRRDRGAQLAAEQKALYRTYCQHGDPTLFWATFNSVMDE

HKAAVRRIQQQRAQGRPAALRHHRFDGTGTLAVQLQRTAGAPPRTPMLLADPAGRYHNVL

HLPWVDPDQWASMTRAEQRHAGRITVRMRCGSLGGQPQWIDLPVQAHRWLPQDADITRAK

LTVTRLGADLRARLSITARLPHPTVPRRGDLPTLAIHLGWHAIDEGVVVAHWRSDRPVAI

PSELADVIVPIVEGISGRIIAPGSVGTRLERYAEIASARDAALNDIRDRLSTWLADQGPR

PHPIRPDEQITAADAARWRSPARFAALALAWRGSDLEIAQPLEAWRRADKLLWQQQAHGR

ARALGHRTDLWRRVASALVSQCGRLVVDDTNISAVIRNTMEHTRIPADLQRQIDRRRDHA

APGALRSSMVAAATRDGVPVIVVPAAGLSRIHAGCGYENRVESRRRRRKIICAGCGRTYD

PDLSATALMLARAAQPPVQP*

>c2c4_3_SD2019_length_604

MAITVHTAGVHYRWTDNPPEQLMRQLRLAHDLREDLVTLQLDYETAKAGIWSSYPAVAAA

ETELADAESAAEQAAAAVSEERTKLRTKRITGPLAQKLTAARKRVREARSTRRAAISEVH

EEAKGRLVDASDALKAQQKALYKTYCQDGDLFWATFNDVLDHHKAAVKRIGQMRAAGQPA

QLRHHRFDGTGSIAVQLQRQAGQPQRTPELIADVDGKYGRVLSVPWVQPDRWERIPRRER

RMIGRVTVRMRAGQLSGEPQWLDIPVQQHRMLPLDADITGARLTVTRTAGTLRAQISVTA

KIPDPEPVTDGPDVAVHLGWRNTDTGVRVARWRSTEPIEVPFDFRDTLTVDPGGRSGEIF

VPEAVPRRVERAHLIASHRADRMNELRARLVDYLAETGPRPHPSREGEELGAGNVRMWKS

PNRFAWLARVWADDESVSTDIREALAQWRHQDWISWHHQEGGRRRSAAQRLDVYRQVAAV

LVSQAGRLVLDDTSYADIAQRSATTKTEELPNETAARINRRRAHAAPGELRQTLVAAADR

DAVPVDTVSHTGVSVVHAKCGHENPSDGRFMSVVVACDGCGEKYDQDESALTHMLTRAVQ

SAA*

>Cas12j_1

MADTPTLFTQFLRHHLPGQRFRKDILKQAGRILANKGEDATIAFLRGKSEESPPDFQPPV

KCPIIACSRPLTEWPIYQASVAIQGYVYGQSLAEFEASDPGCSKDGLLGWFDKTGVCTDY

FSVQGLNLIFQNARKRYIGVQTKVTNRNEKRHKKLKRINAKRIAEGLPELTSDEPESALD

ETGHLIDPPGLNTNIYCYQQVSPKPLALSEVNQLPTAYAGYSTSGDDPIQPMVTKDRLSI

SKGQPGYIPEHQRALLSQKKHRRMRGYGLKARALLVIVRIQDDWAVIDLRSLLRNAYWRR

IVQTKEPSTITKLLKLVTGDPVLDATRMVATFTYKPGIVQVRSAKCLKNKQGSKLFSERY

LNETVSVTSIDLGSNNLVAVATYRLVNGNTPELLQRFTLPSHLVKDFERYKQAHDTLEDS

IQKTAVASLPQGQQTEIRMWSMYGFREAQERVCQELGLADGSIPWNVMTATSTILTDLFL

ARGGDPKKCMFTSEPKKKKNSKQVLYKIRDRAWAKMYRTLLSKETREAWNKALWGLKRGS

PDYARLSKRKEELARRCVNYTISTAEKRAQCGRTIVALEDLNIGFFHGRGKQEPGWVGLF

TRKKENRWLMQALHKAFLELAHHRGYHVIEVNPAYTSQTCPVCRHCDPDNRDQHNREAFH

CIGCGFRGNADLDVATHNIAMVAITGESLKRARGSVASKTPQPLAAE*

>Cas12j_2

MPKPAVESEFSKVLKKHFPGERFRSSYMKRGGKILAAQGEEAVVAYLQGKSEEEPPNFQP

PAKCHVVTKSRDFAEWPIMKASEAIQRYIYALSTTERAACKPGKSSESHAAWFAATGVSN

HGYSHVQGLNLIFDHTLGRYDGVLKKVQLRNEKARARLESINASRADEGLPEIKAEEEEV

ATNETGHLLQPPGINPSFYVYQTISPQAYRPRDEIVLPPEYAGYVRDPNAPIPLGVVRNR

CDIQKGCPGYIPEWQREAGTAISPKTGKAVTVPGLSPKKNKRMRRYWRSEKEKAQDALLV

TVRIGTDWVVIDVRGLLRNARWRTIAPKDISLNALLDLFTGDPVIDVRRNIVTFTYTLDA

CGTYARKWTLKGKQTKATLDKLTATQTVALVAIDLGQTNPISAGISRVTQENGALQCEPL

DRFTLPDDLLKDISAYRIAWDRNEEELRARSVEALPEAQQAEVRALDGVSKETARTQLCA

DFGLDPKRLPWDKMSSNTTFISEALLSNSVSRDQVFFTPAPKKGAKKKAPVEVMRKDRTW

ARAYKPRLSVEAQKLKNEALWALKRTSPEYLKLSRRKEELCRRSINYVIEKTRRRTQCQI

VIPVIEDLNVRFFHGSGKRLPGWDNFFTAKKENRWFIQGLHKAFSDLRTHRSFYVFEVRP

ERTSITCPKCGHCEVGNRDGEAFQCLSCGKTCNADLDVATHNLTQVALTGKTMPKREEPR

DAQGTAPARKTKKASKSKAPPAEREDQTPAQEPSQTS

>Cas12j_3

MEKEITELTKIRREFPNKKFSSTDMKKAGKLLKAEGPDAVRDFLNSCQEIIGDFKPPVKT

NIVSISRPFEEWPVSMVGRAIQEYYFSLTKEELESVHPGTSSEDHKSFFNITGLSNYNYT

SVQGLNLIFKNAKAIYDGTLVKANNKNKKLEKKFNEINHKRSLEGLPIITPDFEEPFDEN

GHLNNPPGINRNIYGYQGCAAKVFVPSKHKMVSLPKEYEGYNRDPNLSLAGFRNRLEIPE

GEPGHVPWFQRMDIPEGQIGHVNKIQRFNFVHGKNSGKVKFSDKTGRVKRYHHSKYKDAT

KPYKFLEESKKVSALDSILAIITIGDDWVVFDIRGLYRNVFYRELAQKGLTAVQLLDLFT

GDPVIDPKKGVVTFSYKEGVVPVFSQKIVPRFKSRDTLEKLTSQGPVALLSVDLGQNEPV

AARVCSLKNINDKITLDNSCRISFLDDYKKQIKDYRDSLDELEIKIRLEAINSLETNQQV

EIRDLDVFSADRAKANTVDMFDIDPNLISWDSMSDARVSTQISDLYLKNGGDESRVYFEI

NNKRIKRSDYNISQLVRPKLSDSTRKNLNDSIWKLKRTSEEYLKLSKRKLELSRAVVNYT

IRQSKLLSGINDIVIILEDLDVKKKFNGRGIRDIGWDNFFSSRKENRWFIPAFHKAFSEL

SSNRGLCVIEVNPAWTSATCPDCGFCSKENRDGINFTCRKCGVSYHADIDVATLNIARVA

VLGKPMSGPADRERLGDTKKPRVARSRKTMKRKDISNSTVEAMVTA*

>Cas12j_4

MYSLEMADLKSEPSLLAKLLRDRFPGKYWLPKYWKLAEKKRLTGGEEAACEYMADKQLDS

PPPNFRPPARCVILAKSRPFEDWPVHRVASKAQSFVIGLSEQGFAALRAAPPSTADARRD

WLRSHGASEDDLMALEAQLLETIMGNAISLHGGVLKKIDNANVKAAKRLSGRNEARLNKG

LQELPPEQEGSAYGADGLLVNPPGLNLNIYCRKSCCPKPVKNTARFVGHYPGYLRDSDSI

LISGTMDRLTIIEGMPGHIPAWQREQGLVKPGGRRRRLSGSESNMRQKVDPSTGPRRSTR

SGTVNRSNQRTGRNGDPLLVEIRMKEDWVLLDARGLLRNLRWRESKRGLSCDHEDLSLSG

LLALFSGDPVIDPVRNEVVFLYGEGIIPVRSTKPVGTRQSKKLLERQASMGPLTLISCDL

GQTNLIAGRASAISLTHGSLGVRSSVRIELDPEIIKSFERLRKDADRLETEILTAAKETL

SDEQRGEVNSHEKDSPQTAKASLCRELGLHPPSLPWGQMGPSTTFIADMLISHGRDDDAF

LSHGEFPTLEKRKKFDKRFCLESRPLLSSETRKALNESLWEVKRTSSEYARLSQRKKEMA

RRAVNFVVEISRRKTGLSNVIVNIEDLNVRIFHGGGKQAPGWDGFFRPKSENRWFIQAIH

KAFSDLAAHHGIPVIESDPQRTSMTCPECGHCDSKNRNGVRFLCKGCGASMDADFDAACR

NLERVALTGKPMPKPSTSCERLLSATTGKVCSDHSLSHDAIEKAS*

>Cas12j_5

MSSLPTPLELLKQKHADLFKGLQFSSKDNKMAGKVLKKDGEEAALAFLSERGVSRGELPN

FRPPAKTLVVAQSRPFEEFPIYRVSEAIQLYVYSLSVKELETVPSGSSTKKEHQRFFQDS

SVPDFGYTSVQGLNKIFGLARGIYLGVITRGENQLQKAKSKHEALNKKRRASGEAETEFD

PTPYEYMTPERKLAKPPGVNHSIMCYVDISVDEFDFRNPDGIVLPSEYAGYCREINTAIE

KGTVDRLGHLKGGPGYIPGHQRKESTTEGPKINFRKGRIRRSYTALYAKRDSRRVRQGKL

ALPSYRHHMMRLNSNAESAILAVIFFGKDWVVFDLRGLLRNVRWRNLFVDGSTPSTLLGM

FGDPVIDPKRGVVAFCYKEQIVPVVSKSITKMVKAPELLNKLYLKSEDPLVLVAIDLGQT

NPVGVGVYRVMNASLDYEVVTRFALESELLREIESYRQRTNAFEAQIRAETFDAMTSEEQ

EEITRVRAFSASKAKENVCHRFGMPVDAVDWATMGSNTIHIAKWVMRHGDPSLVEVLEYR

KDNEIKLDKNGVPKKVKLTDKRIANLTSIRLRFSQETSKHYNDTMWELRRKHPVYQKLSK

SKADFSRRVVNSIIRRVNHLVPRARIVFIIEDLKNLGKVFHGSGKRELGWDSYFEPKSEN

RWFIQVLHKAFSETGKHKGYYIIECWPNWTSCTCPKCSCCDSENRHGEVFRCLACGYTCN

TDFGTAPDNLVKIATTGKGLPGPKKRCKGSSKGKNPKIARSSETGVSVTESGAPKVKKSS

PTQTSQSSSQSAP*

>Cas12j_6

MNKIEKEKTPLAKLMNENFAGLRFPFAIIKQAGKKLLKEGELKTIEYMTGKGSIEPLPNF

KPPVKCLIVAKRRDLKYFPICKASCEIQSYVYSLNYKDFMDYFSTPMTSQKQHEEFFKKS

GLNIEYQNVAGLNLIFNNVKNTYNGVILKVKNRNEKLKKKAIKNNYEFEEIKTFNDDGCL

INKPGINNVIYCFQSISPKILKNITHLPKEYNDYDCSVDRNIIQKYVSRLDIPESQPGHV

PEWQRKLPEFNNTNNPRRRRKWYSNGRNISKGYSVDQVNQAKIEDSLLAQIKIGEDWIIL

DIRGLLRDLNRRELISYKNKLTIKDVLGFFSDYPIIDIKKNLVTFCYKEGVIQVVSQKSI

GNKKSKQLLEKLIENKPIALVSIDLGQTNPVSVKISKLNKINNKISIESFTYRFLNEEIL

KEIEKYRKDYDKLELKLINEA

>Cas12j_7

MSNTAVSTREHMSNKTTPPSPLSLLLRAHFPGLKFESQDYKIAGKKLRDGGPEAVISYLT

GKGQAKLKDVKPPAKAFVIAQSRPFIEWDLVRVSRQIQEKIFGIPATKGRPKQDGLSETA

FNEAVASLEVDGKSKLNEETRAAFYEVLGLDAPSLHAQAQNALIKSAISIREGVLKKVEN

RNEKNLSKTKRRKEAGEEATFVEEKAHDERGYLIHPPGVNQTIPGYQAVVIKSCPSDFIG

LPSGCLAKESAEALTDYLPHDRMTIPKGQPGYVPEWQHPLLNRRKNRRRRDWYSASLNKP

KATCSKRSGTPNRKNSRTDQIQSGRFKGAIPVLMRFQDEWVIIDIRGLLRNARYRKLLKE

KSTIPDLLSLFTGDPSIDMRQGVCTFIYKAGQACSAKMVKTKNAPEILSELTKSGPVVLV

SIDLGQTNPIAAKVSRVTQLSDGQLSHETLLRELLSNDSSDGKEIARYRVASDRLRDKLA

NLAVERLSPEHKSEILRAKNDTPALCKARVCAALGLNPEMIAWDKMTPYTEFLATAYLEK

GGDRKVATLKPKNRPEMLRRDIKFKGTEGVRIEVSPEAAEAYREAQWDLQRTSPEYLRLS

TWKQELTKRILNQLRHKAAKSSQCEVVVMAFEDLNIKMMHGNGKWADGGWDAFFIKKREN

RWFMQAFHKSLTELGAHKGVPTIEVTPHRTSITCTKCGHCDKANRDGERFACQKCGFVAH

ADLEIATDNIERVALTGKPMPKPESERSGDAKKSVGARKAAFKPEEDAEAAE*

>Cas12j_8

MIKPTVSQFLTPGFKLIRNHSRTAGLKLKNEGEEACKKFVRENEIPKDECPNFQGGPAIA

NIIAKSREFTEWEIYQSSLAIQEVIFTLPKDKLPEPILKEEWRAQWLSEHGLDTVPYKEA

AGLNLIIKNAVNTYKGVQVKVDNKNKNNLAKINRKNEIAKLNGEQEISFEEIKAFDDKGY

LLQKPSPNKSIYCYQSVSPKPFITSKYHNVNLPEEYIGYYRKSNEPIVSPYQFDRLRIPI

GEPGYVPKWQYTFLSKKENKRRKLSKRIKNVSPILGIICIKKDWCVFDMRGLLRTNHWKK

YHKPTDSINDLFDYFTGDPVIDTKANVVRFRYKMENGIVNYKPVREKKGKELLENICDQN

GSCKLATVDVGQNNPVAIGLFELKKVNGELTKTLISRHPTPIDFCNKITAYRERYDKLES

SIKLDAIKQLTSEQKIEVDNYNNNFTPQNTKQIVCSKLNINPNDLPWDKMISGTHFISEK

AQVSNKSEIYFTSTDKGKTKDVMKSDYKWFQDYKPKLSKEVRDALSDIEWRLRRESLEFN

KLSKSREQDARQLANWISSMCDVIGIENLVKKNNFFGGSGKREPGWDNFYKPKKENRWWI

NAIHKALTELSQNKGKRVILLPAMRTSITCPKCKYCDSKNRNGEKFNCLKCGIELNADID

VATENLATVAITAQSMPKPTCERSGDAKKPVRARKAKAPEFHDKLAPSYTVVLREAV*

>cas12O.01

MPSYKSSRVLVRDVPEELVDHYERSHRVAAFFMRLLLAMRREPYSLRMRDGTEREVDLDE

TDDFLRSAGCEEPDAVSDDLRSFALAVLHQDNPKKRAFLESENCVSILCLEKSASGTRYY

KRPGYQLLKKAIEEEWGWDKFEASLLDERTGEVAEKFAALSMEDWRRFFAARDPDDLGRE

LLKTDTREGMAAALRLRERGVFPVSVPEHLDLDSLKAAMASAAERLKSWLACNQRAVDEK

SELRKRFEEALDGVDPEKYALFEKFAAELQQADYNVTKKLVLAVSAKFPATEPSEFKRGV

EILKEDGYKPLWEDFRELGFVYLAERKWERRRGGAAVTLCDADDSPIKVRFGLTGRGRKF

VLSAAGSRFLITVKLPCGDVGLTAVPSRYFWNPSVGRTTSNSFRIEFTKRTTENRRYVGE

VKEIGLVRQRGRYYFFIDYNFDPEEVSDETKVGRAFFRAPLNESRPKPKDKLTVMGIDLG

INPAFAFAVCTLGECQDGIRSPVAKMEDVSFDSTGLRGGIGSQKLHREMHNLSDRCFYGA

RYIRLSKKLRDRGALNDIEARLLEEKYIPGFRIVHIEDADERRRTVGRTVKEIKQEYKRI

RHQFYLRYHTSKRDRTELISAEYFRMLFLVKNLRNLLKSWNRYHWTTGDRERRGGNPDEL

KSYVRYYNNLRMDTLKKLTCAIVRTAKEHGATLVAMENIQRVDRDDEVKRRKENSLLSLW

APGMVLERVEQELKNEGILAWEVDPRHTSQTSCITDEFGYRSLVAKDTFYFEQDRKIHRI

DADVNAAINIARRFLTRYRSLTQLWASLLDDGRYLVNVTRQHERAYLELQTGAPAATLNP

TAEASYELVGLSPEEEELAQTRIKRKKREPFYRHEGVWLTREKHREQVHELRNQVLALGN

AKIPEIRT

>cas12O.02

MPSHKSSRVLLHDVPPELVTHYEASHRVARFLMELLLAMRQTPYVRRETNGELHEVTPDE

IDEFLRRYTGERLEAVRPLLKSFAEAVLHEDDKENRAFAKPENAALLLCNSETESGTQYF

KKPGYQLLKQAIEKKWPWKRLKQELVDEKGNITKKFAALSIEEWRDFFESEDLDALGKEL

LRRTQAEGMRAGRRLREEGVFPVRLPDELDIRSSKAALASVSERLKSWIDCNRRAAEQKA

ERKQRFERLRDALESSRYDLFKKFATDLQEIDYSVTARLVQALRHFPKRQPPELQPALAV

LKLDKYRPLWENCGELGRTYLAEQRWKSHSGRAAVSFCDPDRSPIKVRFGLTGRGRPFLL

SAEGGRFFVTLKLACGDIGLRALPSRYFWNPKVTAHRKNGKSEFNVEFTKCTTENRRFAA

HVKELSIVRHKQRYYCFVDYGFEPVPISEAANTACNFFRAPLTQSQPKPKEPVSVMGIDL

GVNPAFAYAVCTLGQQKANQITVPVAKMKDTSFHATGVGGGVHDRKLHADLKELADTCFY

GSKYIGLSKRLRDRGTLNELQRKILEERYIPGFNIVHVEDDHDQRRRNIGARVRELKSEF

KRLRHVFYERQHGRRRRPAPLICSETFQMLFAVKNLRSVLKAWNRYHWTRGDGESRGRDP

NELKSYIDYYKNLRLDTLKKLTCAIVRTARSHGVEIVALEDIKRVDYDDQVKRAKENSLL

SLWAPGMILERIEQELANEGIRTWRIDPRHTSQTACITDEFGYRPVRGKENLFFEANGEL

LRVNSDVNAAINIARRFLTRYRKLTQLWAQPLEDGDYLISVKRQFEAAFLMAETRQPAAV

LVPEGEGIYRLKGISGERETELREQLMRRRSEKFYRHGEHWFTVKRHRKAIDTLRDQVLE

RGARLIREIPT*
